# Emerging single cell endothelial heterogeneity supports sprouting tumour angiogenesis and growth

**DOI:** 10.1101/2021.06.09.447719

**Authors:** Ken Matsumoto, Florian Rambow, Fabio Stanchi, Thomas Mathivet, Junbin Qian, Wolfgang Giese, Liqun He, Diether Lambrechts, Bin Zhou, Christer Betsholtz, Jean-Christophe Marine, Holger Gerhardt

## Abstract

Blood vessels supplying tumors are often dysfunctional and generally heterogeneous. The mechanisms underlying this heterogeneity remain poorly understood. Here, using multicolor lineage tracing, *in vivo* time-lapse imaging and single cell RNA sequencing in a mouse glioma model, we identify tumour-specific blood endothelial cells that originate from cells expressing the receptor for colony stimulating factor 1, *Csf1r*, a cytokine which controls macrophage biology. These *Csf1r* lineage endothelial cells (CLECs) form up to 10% of the tumour vasculature and express, besides classical blood endothelial cell markers, a gene signature that is distinct from brain endothelium but shares similarities with lymphatic endothelial cell populations. *in silico* analysis of pan-cancer single cell RNAseq datasets highlights the presence of a comparable subpopulation in the endothelium of a wide spectrum of human tumours. We show that CLECs actively contribute to sprouting and remodeling of tumour blood vessels and that selective depletion of CLECs reduces vascular branching and tumour growth. Our findings indicate that a non-tumour resident Csf1r-positive population is recruited to tumours, differentiates into blood endothelial cells to contribute to vascularization and, thereby, tumour growth.

## INTRODUCTION

Endothelial cells form a single cell layer lining the inner walls of blood vessels and play critical roles in organ homeostasis and disease progression. Once formed, following embryonic and early post-natal development, blood vessels retain a high level of adaptability to meet changing metabolic or hemodynamic requirements or enable further tissue growth including in tumours. How local vascular networks respond to these adaptive challenges, rapidly expand, remodel and reestablish homeostasis remains a timely research topic, with many open questions still unresolved. In particular in pathologies, knowledge of the precise nature and origin of new vessel formation, endothelial activation and differentiation steps, and of the mechanisms driving vessel dysmorphia, or preventing effective revascularization in ischemic diabetic complication is critical for new therapeutic approaches. New blood vessel formation in the adult was long believed to arise exclusively by sprouting and proliferation of endothelial cells from local blood vessels without any de novo differentiation from progenitor cells. However, Asahara and coworkers isolated mononuclear cells from human peripheral blood and identified circulating endothelial progenitor cells (EPCs) ^1^. These cells were shown to derive from the bone marrow and contribute to endothelial cells in blood vessels in hindlimb ischemia and tumour xenograft mouse model ^2, 3^. By fluorescent in situ hybridization with sex chromosome-specific probes in patients with cancer after bone marrow transplantation, Peters and coworkers demonstrated that bone marrow-derived stem cells contribute to endothelial cells in human tumour endothelium ^4^. The resulting concept of vasculogenesis in the adult raised the prospect of novel therapeutic approaches for ischemic vascular disease and for targeting tumour angiogenesis ^5, 6^. Interestingly, EPCs also contributed to newly forming lymphatic vessels ^7, 8^. A vast number of publications since have supported or challenged these initial discoveries, and the occurrence, origin and significance of EPCs continues to be a matter of debate ^9–12^. Whereas EPCs and other circulating progenitor cells were long thought to origin from the bone marrow, recent studies challenged this idea and suggested the existence of a different yet unknown vascular niche for endothelial progenitor cells that can contribute to blood vessel formation^13^.

## RESULTS

### A Csf1r cell lineage contributes to blood vascular endothelial cells during glioma growth in mice

In order to study dynamic macrophage recruitment and blood vessel patterning *in vivo* during glioma growth, we adapted a surgical cranial window model for two-photon microscopy allowing repeated visualization of the same animal over time. We generated spheroids of syngeneic C57BL/6 mouse glioma cells (CT2A or GL261) ^14, 15^ modified to stably express blue fluorescent protein (BFP), and injected them into the mouse cortex, followed by implantation of a glass coverslip ^16^. By crossing transgenic *Csf1r (Colony stimulating factor 1 receptor)* specific tamoxifen-inducible Cre driver mice ^17^ with the *mTmG* Cre reporter mouse line ^18^, we induced membrane-targeted GFP expression in *Csf1r*-expressing cells and used this stable lineage trace to follow the cell population over time (Fig. 1a, b). As expected, the vast majority of tumour associated macrophages were labelled by GFP in line with their *Csf1r*-dependent recruitment and expansion^19^. Unexpectedly, however, through the combination of longitudinal live two-photon imaging and lineage tracing, we discovered GFP-positive cells contributing to the tumour vasculature during glioma growth (Fig. 1c). These *Csf1r* lineage endothelial cells (CLECs) frequently emerged at tip cell positions of the growing vasculature, heading blood vessel sprouts in the mouse glioma (Fig. 1d, Supplementary Movie1). Additionally, CLECs were found bridging adjacent capillaries during anastomosis (Fig. 1e). Fluorescent dextran injection confirmed that CLECs contributed to - and were incorporated in the endothelial lining of - functionally perfused blood vessels including lumenized sprouts headed by endothelial tip cells (Fig. 1f). Counterstaining on fixed samples identified that all cells forming blood vessels including the lineage-traced CLECs expressed typical endothelial markers (CDH5, CD31) and lacked macrophage and myeloid cell markers (F4/80, CD45) (Fig. 1g, h, Extended Data Fig. 1). BFP was not expressed in CLECs, demonstrating that they were not derived from glioma cells (Fig. 1g).

**Fig. 1:**
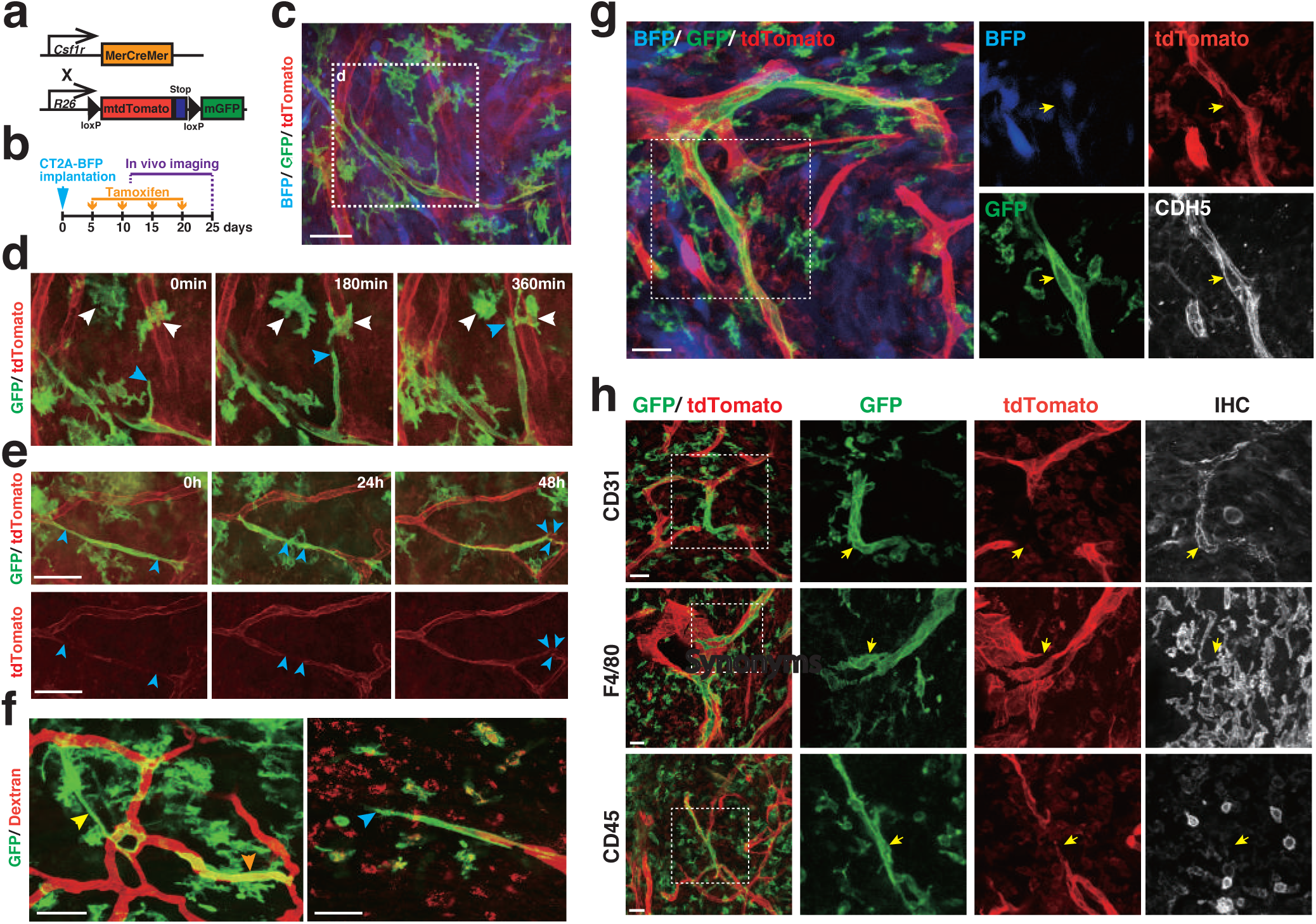
Csf1r lineage endothelial cells (CLECs) contribute to perfused, dynamically sprouting and remodeling blood vessels in mouse glioma. a, Csf1r-Mer2.CremTmG: a mouse model for Csf1r-lineage tracing combining the tamoxifen-inducible Cre-driver under control of the Csf1r-promoter and the mTmG Cre-reporter. b, Timing schedule of glioma implantation, tamoxifen induction and in vivo imaging. c-f, Longitudinal intravital imaging of the tumour vasculature in CT2A mouse glioma of Csf1r-Mer2.CremTmG by cranial window. Scale bar: 50 μm. c, In vivo imaging in 3 weeks glioma. Endothelial cells in red (tdTomato), Csf1r-labeled cells in green (GFP) and glioma cells in blue (BFP). d, Detail image of outlined square in Fig. 1c illustrating tracing of GFP-positive cells. White arrowheads, macrophages; blue arrowhead, CLECs at tip position. e, In vivo imaging of CLECs in 2 weeks glioma. Blue arrowheads, tip of endothelial cells. f, Visualization of blood vessels in 3 weeks glioma by Intravenous injection of dextran with texas red dye. Yellow arrowhead, bridging CLECs; orange arrowhead, tube formation by CLECs; blue arrowhead, CLECs at tip position. g-h, Counterstaining on PFA-fixed 4 weeks CT2A glioma sections of Csf1r-Mer2.CremTmG with indicated antibodies. Yellow arrows, CLECs. Scale bar: 25 μm.

To gain more insight into the nature and significance of CLECs, we isolated and quantified the cell population in the mouse glioma model. By 4 weeks of tumour development (close to the specified humane endpoint of 5 weeks) up to 10% of the tumour vasculature was comprised of cells carrying the *Csfr1* lineage trace. Similar frequencies were found in tumours derived from either of the mouse glioma cell lines, CT2A (Fig. 2a, b), or GL261 (Fig. 2c). Polyclonal antibody staining indicated that *Csf1r* may also be expressed in some brain capillary endothelial cells ^20, 21^. Recent single cell sequencing of adult mouse brain endothelial cells however did not confirm this notion^22^, suggesting that CLECs are unlikely to originate from brain endothelium. Indeed, tamoxifen mediated activation of our transgenic *Csfr1* lineage trace did not label brain endothelial cells in mice lacking tumours, sham operated as well as in the healthy brain parenchyma surrounding implanted tumours (Fig. 2d, e, f). *Csf1r* is important for the embryonic development of microglia and its expression is also detected in adult microglia by single cell RNA sequencing. Surprisingly however, Csf1r-Mer-iCre-Mer mediated recombination of the Cre-Reporter expression was also not observed in microglia of the healthy adult mouse brain. Irrespective of this apparent discrepancy between reported microglial *Csf1r* expression and the lack of Csf1r-Mer-iCre-Mer mediated recombination in microglia, these data also exclude microglia as a potential source of CLECs. Furthermore, we observed CLECs also in tumours originating from B16F1 melanoma cells implanted under the skin; thus, this *Csfr1*-lineage endothelial cell contribution not only occurs in brain tumours with microglia in their environment, but also in other tumour types and locations (Fig. 2g, h). Notably, we also found *Csf1r*-lineage lymphatic endothelial cells in B16F1 skin melanomas indicating that CLECs can contribute to both blood and lymphatic vessels (Extended Data Fig. 2).

**Fig. 2:**
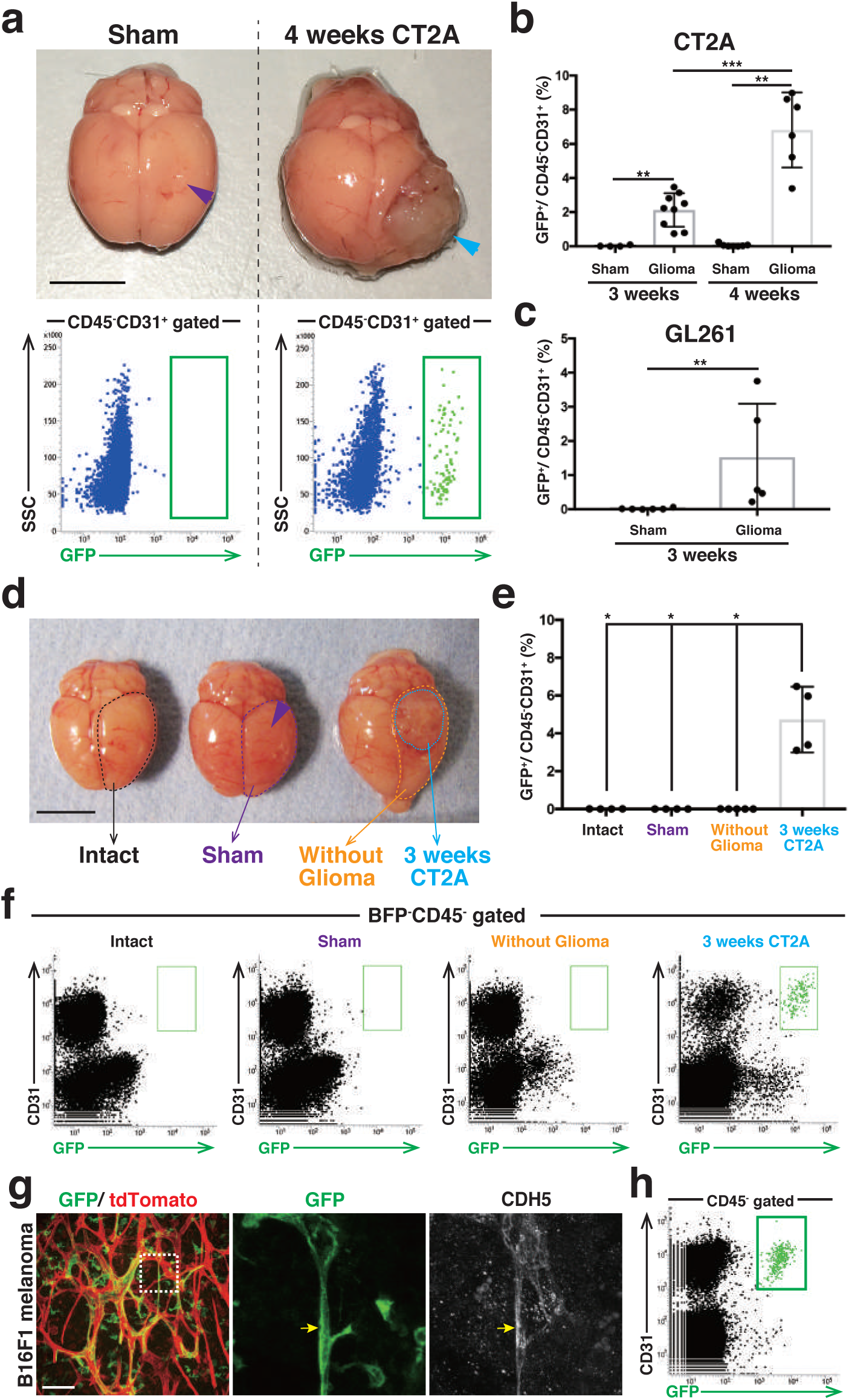
Up to 10% of intra-tumour endothelium is comprised of CLECs not only in glioma, but also in other tumour types and locations. a-i, Quantification of the population of CLECs in glioma of Csf1r-Mer2.CremTmG mice. a, Gross appearance of sham control and 4 weeks CT2A glioma (Upper panel). Purple arrowhead, the area of capillary injection; blue arrowhead, glioma. Scale bar: 5 mm. Flow cytometric analysis of CLECs in sham control and 4 weeks CT2A glioma (Lower panel). b, In 3 weeks CT2A glioma (n= 9), CLECs constitute 2.1 ± 1.0% of total endothelial cells (CD45-CD31+); in 4 weeks CT2A glioma (n= 5), they make up 7.5 ± 1.6% of total endothelial cells. c, In 3 weeks GL261 glioma (n= 5), CLECs constitute 1.5 ± 1.6% of total endothelial cells. d, Gross appearance of normal brain, sham control and 3 weeks CT2A glioma. Purple arrowhead, area of capillary injection. Scale bar: 5 mm. e-f, Quantification of the population of CLECs by flow cytometric analysis in brain and tumour samples. Bars represent mean ± s.d. *p<0.05, **p<0.01, ***p<0.001, ****P<0.0001. Two-tailed unpaired Mann-Whitney’ s U test. i-j, Analysis of CLECs in B16F1 melanoma in Csf1r-Mer2.CremTmG mice. g, Counterstaining on PFA-fixed section of day12 melanoma with CDH5 antibody. Yellow arrows, CLECs. Scale bar: 50 μm. h, CLECs constitute 2.8 ± 0.75% of total endothelial cells in day 12 melanoma (n=4).

### CLECs do not originate from hematopoietic niche in bone-marrow or spleen

Given the abundant labelling of macrophages and the important role of *Csf1r* in the recruitment of bone-marrow derived monocytes and differentiated macrophages, we asked whether CLECs are also derived from the bone-marrow. To test this possibility, we generated bone marrow chimeras. Flow cytometric analysis confirmed 97.3 ± 1.6 % reconstitution of wild-type recipient bone marrow by transplanted *Csf1r-Mer2.Cre^mTmG^* bone marrow after 8 weeks post transplantation (Extended Data Fig. 3). Analysis of glioma blood vessels in these bone marrow chimeras (BM*^Csf1r-Mer2.Cre-mTmG^*) failed to identify bone marrow derived CLECs, although the numbers of tumour macrophage were similar to controls (Fig. 3a). Conversely, chimeras of wild-type donor bone marrow transplanted into *Csf1r-Mer2.Cre^mTmG^* recipients (*Csf1r-Mer2.Cre^mTmG^*::BM^WT^), displayed similar CLECs numbers in glioma vessels as observed earlier in non-chimeric *Csf1r-Mer2.Cre^mTmG^* mice (Fig. 3b). These results were confirmed by intravital imaging in the glioma of *Csf1r-Mer2.Cre^mTmG^*::BM^WT^ (Fig. 3c, Supplemental Movie 2). Thus, the bone marrow chimera and lineage tracing experiments strongly suggest that CLECs do not originate from bone marrow-derived cells, nor do they represent previously suggested transdifferentiation events from bone marrow-derived macrophages. Surgical removal of the spleen also did not affect CLECs numbers, further ruling out spleen-derived macrophages as source of CLECs (Extended Data Fig. 4).

**Fig. 3:**
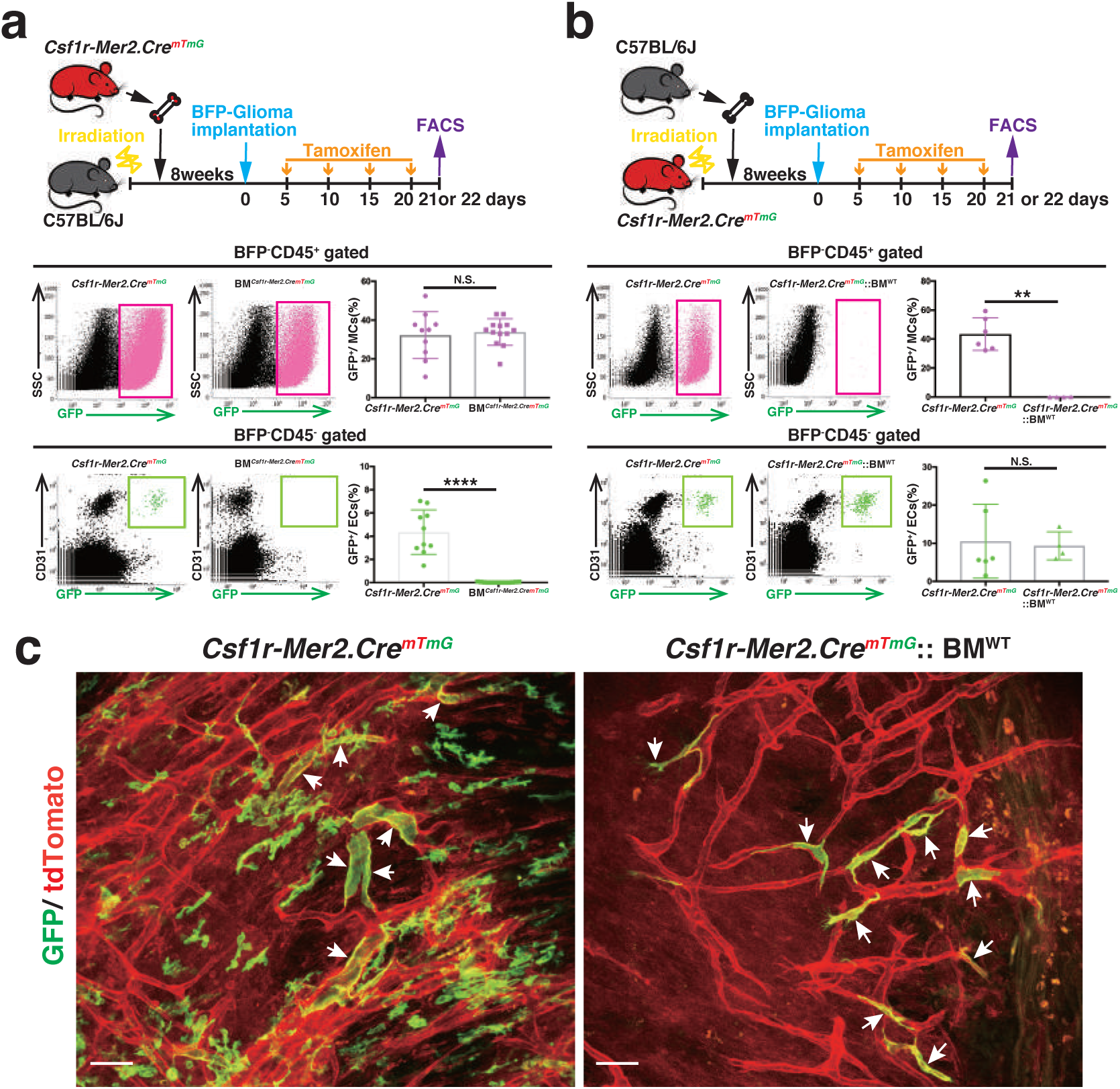
CLECs do not originate from the bone marrow. a-b, Tamoxifen induction timing. b, Quantification of the population of tumour macrophagens (T_MACs: CD45+GFP+) and tumour CLECs (T_CLECs: CD45-CD31+GFP+) by flow cytometric analysis in tumour samples (n=4). Bars represent mean ± s.d. c, Fluorescent images of PFA-fixed 3 weeks CT2A glioma de of Csf1r-Mer2.CremTmG mice within 24 hours after tamoxifen induction. Scale bar: 100 μm. d-e, Flow cytometric analysis of CT2A-glioma in bone marrow chimeras. d, BMCsf1r-mer2.CremTmG: C57BL/6J mice are used as recipients and Csf1r-Mer2.CremTmG mice used as donors. Within the sorted myeloid cell population (BFP-CD45+), 32.3 ± 12.1% are GFP-positive in Csf1r-Mer2.CremTmG mice (n= 10), and 33.8 ± 6.9% are GFP-positive in bone-marrow transplanted BMCsf1r-Mer2.CremTmG mice (n= 12). Among endothelial cell population (BFP-CD45-CD31+), 4.3 ± 1.9% are GFP-positive in Csf1r-Mer2.CremTmG mice (n= 10), whereas no GFP-positive endothelial cells are found in BMCsf1r-Mer2.CremTmG mice (n= 12). e, Csf1r-Mer2.CremTmG:: BMWT: Csf1r-Mer2.CremTmG are used as recipients and C57BL/6J as donors. Of the sorted myeloid cell population, 43.4 ± 11.3% GFP-positive cells are found in Csf1r-Mer2.CremTmG mice (n= 6), whereas 0.1 ± 0.1% GFP-positive cells are found in Csf1r-Mer2.CremTmG:: BMWT mice (n= 4). Of the endothelial cell population, 10.5 ± 9.7% GFP-positive cells are found in Csf1r-Mer2.CremTmG mice (n= 6), and 9.3 ± 3.7% GFP-positive cells are found in Csf1r-Mer2.CremTmG:: BMWT mice (n= 4). Bars represent mean ± s.d. *p<0.05, **p<0.01, ***p<0.001, ****p<0.0001. Two-tailed unpaired Mann-Whitney’ s U test. e, Longitudinal intravial imaging of 2 weeks CT2A glioma of Csf1r-Mer2.CremTmG and Csf1r-Mer2.CremTmG :: BMWT mice. White arrows, CLECs. Scale bar: 50 μm.

### CLECs regulate vascular patterning and support glioma growth

In late-stage gliomas, CLECs preferentially localized to the peripheral tumor area, correlating with the zone of most active angiogenesis (Extended Data Fig. 5). Real-time intravital imaging of mouse gliomas also revealed that CLECs incorporation into vessels is very dynamic. This implied that CLECs might have unique and potentially transient functions during tumour vessel formation. To gain a better understanding of CLECs behaviour during tumour growth, we developed dual-recombination combinatorial genetics as a tool for CLECs tracing ^23, 24^. To achieve selective recombination in CLECs, we used the *Cdh5* endothelial driver for expression of the Dre recombinase, and the same transgenic mouse line as above expressing the tamoxifen inducible form of the Cre recombinase under control of the *Csf1r* transgene. The combined action of these distinct recombinases, Dre and Cre, allowed for deletion of two stop cassettes flanked by RoxP and LoxP sites, respectively, located upstream of a tdTomato expression cassette (Fig. 4a). Using this strategy, we could selectively trace tdTomato-positive CLECs in mouse gliomas following tamoxifen exposure (Fig. 4b). Note that the appearance of labelled CLECs and blood vessel labelling differs from the images of the mTmG line, as the dual Dre/Cre tdTomato reporter used lacks a general label of unrecombined cells. Fluorescent dextran injection instead was used to label vascular lumen, highlighting dtTomato positive CLECs that line the vessel lumen. As expected, counterstaining on fixed samples demonstrated that CLECs express the prototypical endothelial marker, ERG (ETS-related gene) (Fig. 4c). In order to assess whether CLECs contribute functionally to tumour growth and vascular patterning in glioma, we used the same dual recombination strategy to selectively ablate CLECs (Fig. 4d). In this model, CLECs expressed both GFP as well as the diphtheria toxin receptor after tamoxifen induction enabling depletion of GFP-positive CLECs by administration of diphtheria toxin (Fig. 4e, h). Strikingly, CLECs ablation significantly decreased glioma growth (Fig. 4g, i). CLECs ablation also decreased the vascular network length and the number of bifurcations in tumour vessels (Fig. 4j, k, l), indicating that CLECs may promote tumour growth by supporting vessel branching.

**Fig. 4:**
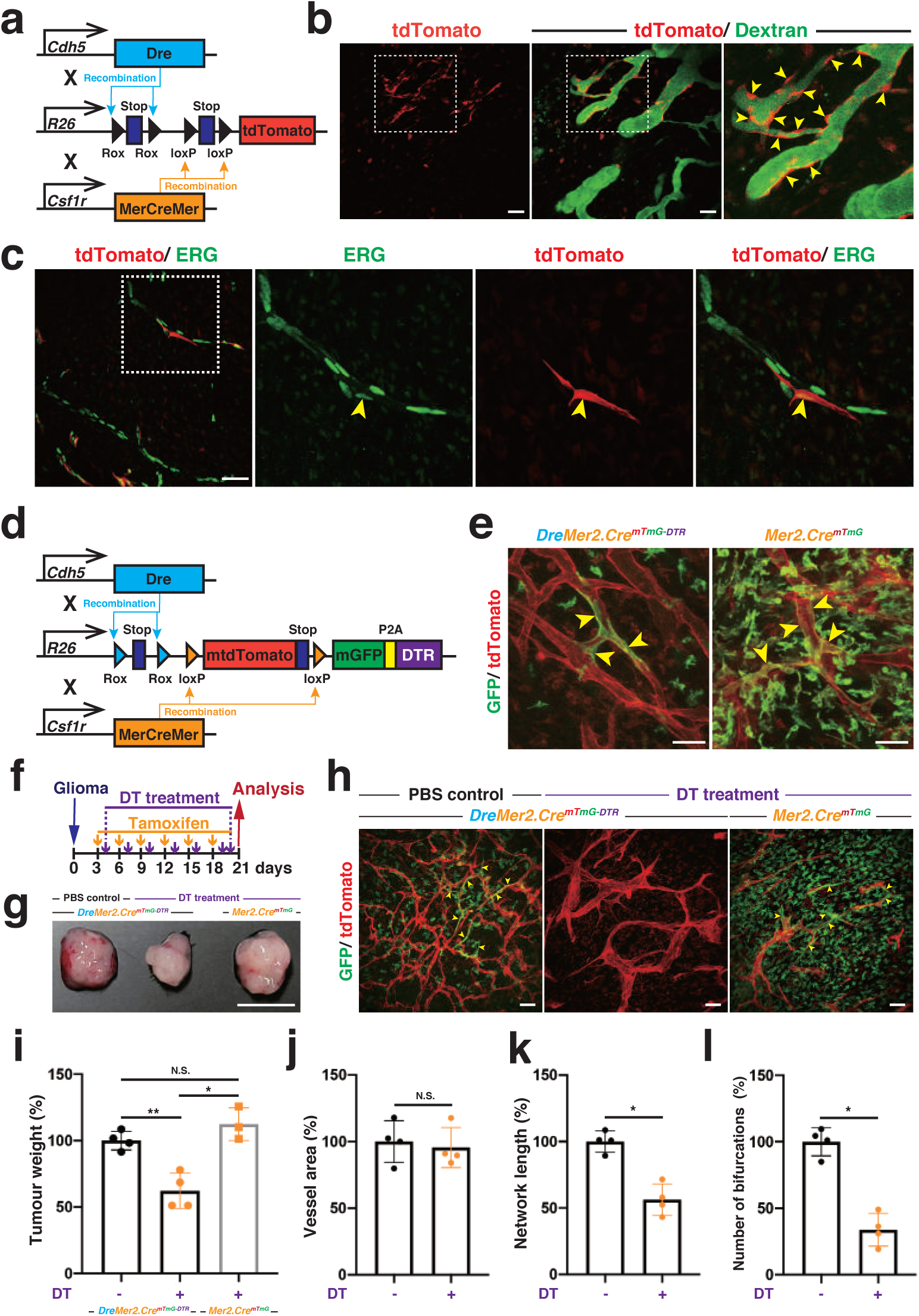
Dual recombination lineage tracing and inducible depletion model identifies a role for CLECs in tumour blood vessel patterning supporting glioma growth. a, Cdh5-Dre: Csf1r-Mer2.Cre: RSR-LSL-tdTomato (DreMer2.CretdTomato): a genetic mouse model for CLEC selective tracing by dual recombination. b-c, Analysis in DreMer2.CretdTomato. b, Longitudinal intravital imaging of the tumour vasculature in 3 weeks CT2A mouse glioma by Intravenous injection of FITC dextran. Yellow arrowheads, CLECs. Scale bar: 50 μm. c, ERG antibody counterstaining on PFA-fixed sections of 3 weeks CT2A glioma. Yellow arrows, CLECs. Scale bar: 50 μm. d, Cdh5-Dre: Csf1r-Mer2.Cre: RSR-mTmG-DTR (DreMer2.CremTmG-DTR): a genetic mouse model for CLEC selective tracing and inducible ablation. DTR, diphtheria toxin receptor. e-m, Analysis in DreMer2.CremTmG-DTR. e, Longitudinal intravital imaging of the tumour vasculature in 3 weeks CT2A mouse glioma. Mer2.CremTmG, Csf1r-Mer2.CremTmG. Yellow arrowheads, CLECs. f, Timing schedule of glioma implantation, tamoxifen induction and diphtheria toxin (DT) treatment. g, Gross appearance of 3 weeks CT2A gliomas. Scale bar: 5 mm. h, Confocal images of PFA-fixed 3 weeks CT2A glioma sections. Yellow arrows, CLECs. Scale bar: 50 μm. i, Tumour weight at 3 weeks CT2A glioma. j-l, Analysis of PFA-fixed 3 weeks CT2A glioma sections. j, Tumour blood vessels area. k, Network length of tumour blood vessels. l, Number of bifurcations of tumour blood vessels. Bars represent mean ± s.d. *p<0.05, **p<0.01. Two-tailed unpaired Mann-Whitney’ s U test.

### CLECs express a unique set of markers

To further understand the nature of tumour CLECs (T_CLECs), we performed single-cell RNA sequencing (scRNA-seq) on isolated T_CLECs in the mouse glioma model. Single-cell transcriptome analysis using Smart-seq2 ^25^ identified that the gene expression pattern of T_CLECs was similar to lineage-negative tumour endothelial cells (T_ECs), but not to tumour macrophages (T_MACs), as determined by UMAP (Uniform Manifold Approximation and Projection) analysis (Fig. 5a). Single-cell differential expression (SCDE) analysis ^26^ confirmed that T_CLECs expressed typical endothelial markers, but lacked macrophage and myeloid cell markers (Fig. 5b). Unexpectedly, however, T_CLECs showed no *Csf1r* expression, despite carrying the *Csf1r*-lineage marker (Fig. 5c). Beyond the overlapping gene expression pattern between T_CLECs and T_ECs, SCDE analysis identified a unique signature that can distinctly define the T_CLECs population; RNAs coding for 24 genes including three types of cell surface proteins (*Aqp1: Aquaporin 1, Fabp4: Fatty acid binding protein 4, Kcnj8: Potassium voltage-gated channel subfamily j member 8*) were significantly enriched in T_CLECs (Fig. 5d; Extended Data Fig. 6; Supplementary Table 1). *Aqp1* is highly expressed in vascular endothelial cells and was reported to contribute to tumour growth and angiogenesis ^27^. Staining of lineage traced T_CLECs on fixed mouse glioma samples confirmed AQP1 expression (Extended Data Fig. 7).

**Fig. 5:**
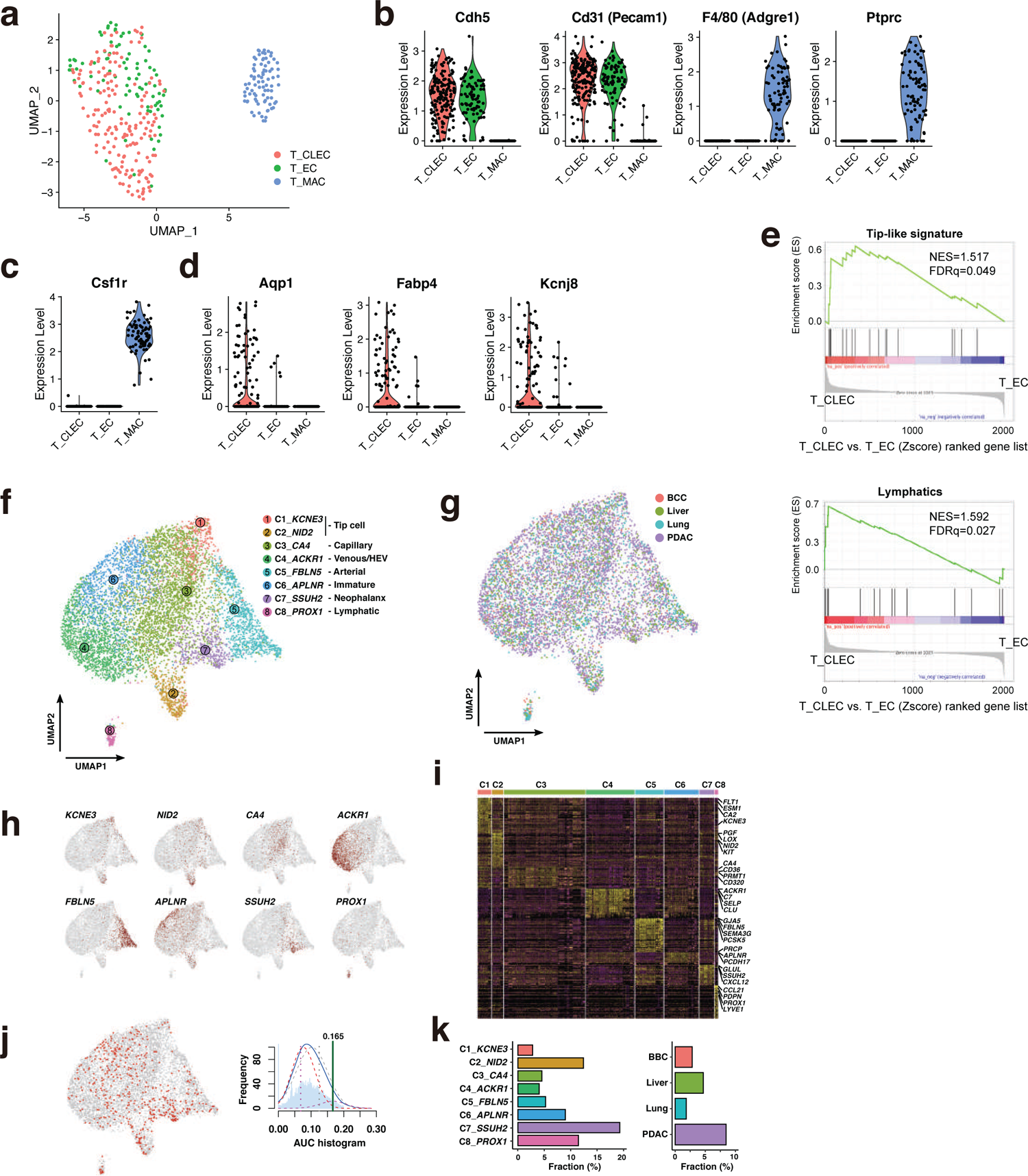
Single-cell analysis identifies a unique signature that defines the tumour CLECs. a-d, Single-cell RNA sequencing (scRNA-seq) analysis of tumour macrophages (T_MACs: CD45+GFP+, 79 cells), tumour endothelial cell (T_ECs: CD45-CD31+GFP-, 78 cells), tumour CLECs (T_CLECs: CD45-CD31+GFP+, 141 cells) in 4 weeks CT2A glioma of Csf1r-Mer2.CremTmG mice. b, Gene expression of typical endothelial cell markers, macrophages and myeloid cell markers. c, Csf1r gene expression. d, Three characteristic genes expressed in T_CLECs. e, Enrichment of Tip-cell signature in T_CLECs compared to T_ECs as shown by GSEA. f-k, Analysis of scRNA-seq data of human endothelial cells from multiple primary tumours. f, UMAP plot of human endothelial cell subtypes. g, UMAP plot of human endothelial cells colour-coded for the tumour type of origins. h, UMAP plots with marker gene expression for each EC subtype. i, Heatmap of marker gene expression for each EC phenotype (each cell as a column and each gene as a row). j-k, AUCell analysis of 24 T_CLECs markers. j, T_CLECs subpopulation indicated (AUC>0.165) by red colored dots. k, Cell type-specific (left) and tumour type-specific (right) populations of T_CLECs. BCC, basal cell carninoma: PDAC, pancreatic ductal adenocarcinoma.

### The T_CLEC state is enriched for “Tip-like” and a lymphatic gene signature

Gene set enrichment analysis (GSEA) identified that the T_CLEC state shows significant enrichment for a Tip-cell-like associated gene expression program (Fig. 5e), thus supporting the morphological observations and their functional importance for branching and tumour growth. Intriguingly, GSEA identified that T_CLECs were also enriched for lymphatic gene signatures (Fig. 5e). AUCell analysis ^28^ of scRNA-seq data from multiple human primary tumours identified subpopulations of endothelial cells that express the T_CLECs marker gene set (Fig. 5f-k, 442 out of 7314 ECs are candidates: 6.0%; Supplementary Table 2). These *bona-fide* T_CLECs were highly enriched in tip cell, neophalanx cell, lymphatic endothelial cell subpopulations and particularly abundant in human pancreatic cancer (Fig. 5k). Neophalanx endothelial cells, which express two pro-angiogenesis markers, *Glul (Glutamine synthetase)* and *Cxcl12 (C-X-C motif chemokine ligand 12)* ^29, 30^ have recently been identified in lung tumour-associated endothelial cell studies ^31, 32^, and were proposed to line neo-vessels after termination of vessel sprouting. Together this data suggests that based on the transcriptional state, T_CLECs might have a supportive role in tumor vessel sprouting and remodeling. Detailed comparison with recent single cell RNA seq data on tumour endothelial cells in mouse and human tumours^31^ confirmed the Tip-like gene signature of T_CLECs, but also identified significant similarities with lymphatics (Fig. 5k). The latter is particularly surprising given that our perfusion experiments with dextran clearly demonstrated that T_CLECs in the glioma model line blood vessels, not lymphatics.

### Acute recombination identifies a CLEC lineage in lymph nodes

The transcriptional similarities between T_CLECs and lymphatic endothelial cells are noteworthy as blood endothelial cells, in particular in the brain environment, and lymphatic EC represent highly differentiated cell types. During vertebrate embryogenesis, lymphovenous specification sees common progenitor cells diverge in their differentiation to establish mature blood endothelium and lymphatic endothelium^33^. The apparent similarities could therefore suggest that T_CLECs represent an immature endothelial cell state with partially overlapping profiles of both immature blood and lymphatic cells. In this case, the expression of lymphatic genes would signify a return to an earlier cell state or potentially some level of transdifferentiation between the two endothelial cell states.

The latter would imply a distant source of lymphatic endothelium as possible origin of CLECs. Indeed, when searching for CLECs in all organs, we observed selective endothelial labelling in lymph nodes, notably restricted to the periphery where specialized lymphatic endothelial cells form the floor of the subcapsular sinus (SCS, Fig. 6a). Bone-marrow transplantation to eliminate the abundant Csfr1-lineage macrophage population highlighted the specific labelling of CLECs in lymph nodes (Fig. 6b).

**Fig. 6:**
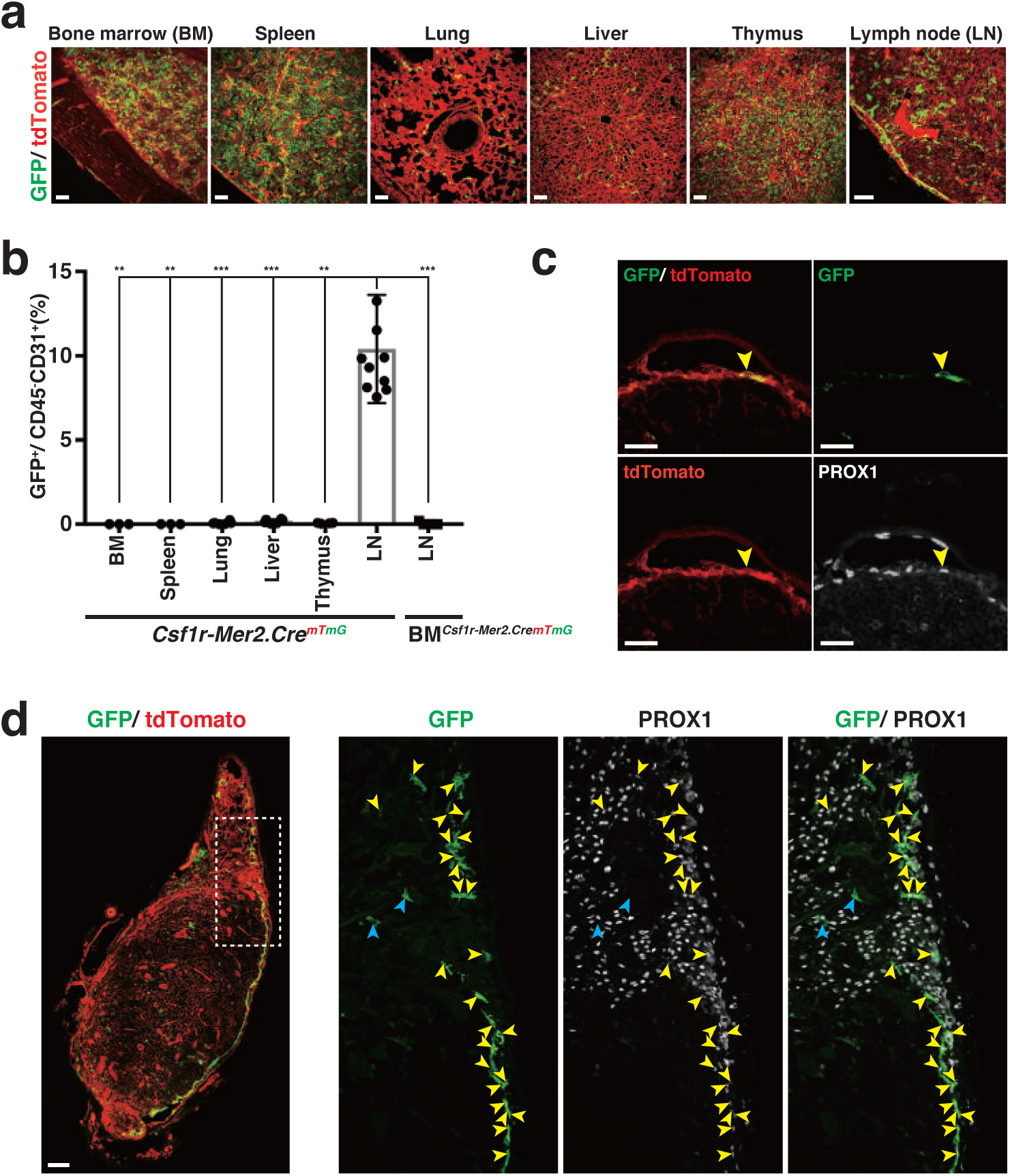
CLECs are linked to Prox1-positive lymphatic endothelium of the subcapsular sinus floor in lymph nodes. a, Fluorescent images of the indicated organs of Csf1r-Mer2.CremTmG mice after multiple tamoxifen induction. Scale bar: 50 μm. b, Quantification of the population of CLECs in Csf1r-Mer2.CremTmG mice by flow cytmetric analysis. Of the endothelial cell population (CD45-CD31+), 9.8 ± 2.1% GFP-positive cells are found in the axillary lymph node of Csf1r-Mer2.CremTmG mice (n= 10). c, Counterstaining on PFA-fixed Inguinal lymph node of Csf1r-Mer2.CremTmG:: BMWT mice using PROX1 antibody within 24 hours after tamoxifen induction. Yellow arrowheads, GFP+tdTomato+PROX1+ cells in the floor of the subcapsular sinus. Scale bar 25 μm. d, Fluorescent images of PROX1 antibody counterstaining on PFA-fixed inguinal lymph node of Csf1r-Mer2.CremTmG::BMWT mice after multiple rounds of tamoxifen induction. Scale bar: 100 μm. Detailed image of square outlined in d. Yellow arrowheads, GFP+Prox1+ cells; blue arrowheads, GFP+Prox1-cells. h-k, Analysis of CT2A glioma in Prox1-CreERT2mTmG mice.

To gain more insight into the emergence of T_CLECs, we asked whether acute recombination could indicate their possible origin. Acute induction experiments (24h after single tamoxifen injection) illustrated that the Csf1r-expressing progenitors of this tumour endothelial cell population do not reside locally in pre-existing brain or the associated brain tumour vasculature (Extended Fig. 8). Close inspection however revealed GFP-positive endothelial cells with significant labelling 24h post-injection in the floor endothelial cells of the SCS of lymph nodes (Fig. 6c). The mTmG Cre-reporter switches from red fluorescent protein expression to mEGFP expression upon recombination. Therefore, cells that acutely recombine, carry both GFP and Tomato expression until the latter is fully degraded. Strikingly, floor SCS endothelial cells exhibited both GFP and simultaneous Tomato expression, demonstrating recent recombination. FACS analysis confirmed the presence of double positive EC in lymph nodes, but not in brain or brain tumours. In fact, double positive cells for GFP and Tomato were not observed in the brain tumours at any stage, excluding the possibility of any acute or continuous local recombination of T_CLECs within the tumour microenvironment. Prox-1 staining confirmed the lymphatic nature of CLECs in the lymphnode, and demonstrated that only a subpopulation of lymphatic endothelial cells in the lymph node carry the Csfr1-lineage trace, the cells in the floor of the SCS (Fig. 6d).

We decided to perform scRNA seq also on endothelial cells isolated from lymph nodes, assess gene expression and perform pseudotime analysis to understand differentiation of T_CLECs, T_EC, LN_CLECs (GFP positive) and LN_ECs (GFP negative) (Fig. 7a-f). We used the same Smart-seq2 platform for comparison to sequence another 384 cells of lymph node origin. After quality control (Fig. 7a), 89 cells EC, not Csfr1 lineage EC, and 104 cells expressing GFP (Fig.7b) in the lymph nodes were selected for further analysis. Comparative analysis with recent mouse lymph node scRNA data^34^ confirmed that the LN_CLEC co-register with identified SCS floor cells (Extended Data Fig. 9a).

**Fig. 7:**
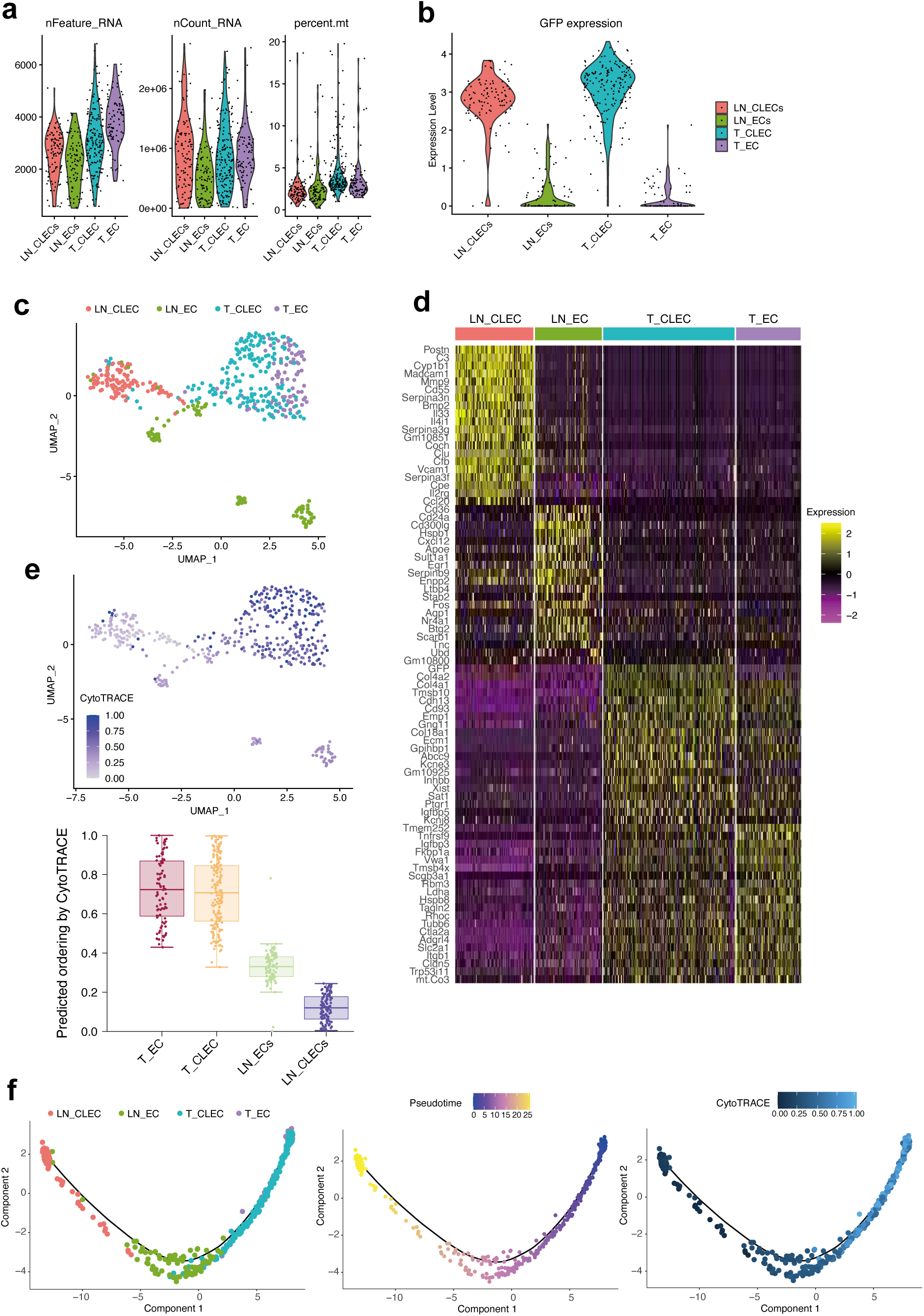
CLEC and EC heterogeneity in tumor and lymph node. a, Quality measures of 4 SMARTseq2 scRNA-seq. libraries: Csf1r-lineage endothelial cells (CLECs) and endothelial cells (ECs) isolated from tumor (T_) and lymph node (LN_). b, Violin plots show GFP expression per single cell. c, Uniform manifold approximation and projection (UMAP) of T_CLECs, T_ECs and LN_CLECs, LN_ECs. d, Heatmap shows the 20 most characteristic markers per cell population. e, UMAP colored by gene expression diversity (CytoTRACE score) and predicted ordering by CytoTRACE. f, Monocle based trajectory analysis of four cell populations, colored by pseudotime and CytoTRACE score.

Unlike T_CLECs, LN_CLECs also expressed Csf1r, in line with our finding of acute recombination in the SCS floor cells.

UMAP based dimension reduction placed T_CLECs and T_EC closely together, as before, and also LN_CLEC and LN_EC co-cluster, although two additional distinct LN_EC clusters appear highly distinct and thus unrelated to any of the other cell types (Fig. 7c). Interestingly, some T_CLEC cluster with the LN_CLEC, suggesting that there is at least some overlap in gene expression between the Csf1r lineage cells in the brain tumour and in the lymph nodes. Looking at relative expression levels of all genes in a heatmap comparing the lineage positive and negative EC from lymph nodes and tumours highlighted that differential expression is more prevalent than any overlap between tumour EC and LN EC (Fig. 7d). However, there are small number of genes like Pde4d, Robo1 and Itga9 that show higher expression in the two lineage positive populations in LN and tumour, compared to the lineage negative cells (Extended Data Fig. 9b). Itga9 is well known to be important in lymphatic development^35^, confirming again that T_CLECs carry some gene expression patterns normally associated with lymphatic EC. Nevertheless, despite the fact that only LN_CLEC appear to acutely recombine, differential gene expression analysis does not provide clear evidence for a lineage relationship between LN_CLEC and T_CLEC.

### T_EC and T_CLECs “co-evolve” during tumour growth, suggesting distinct origin

To gain more insight into the differentiation state and possible route of differentiation, we next performed pseudotemporal ordering using Monocle^36^ (Fig. 7f). The Monocle inferred trajectory suggested equally distributed pseudotime values for T_CLECs and T_EC then LN_EC and finally LN_CLECs. When measuring single-cell gene expression diversity as surrogate for differentiation potential with the CytoTRACE algorithm^37^ (higher score means more undifferentiated, stem-like), we observed again equal scores between T_CLECs and T_EC but lower cytoTRACE values for LN_EC and LN_CLECs (Fig. 7e-f). The equal scores between T_CLECs and T_ECs argue against a role of T_ECs as precursors for T_CLECs, independently confirming that T_CLECs are unlikely emerging through differentiation from the pre-existing brain tumour vasculature. However, the LN_CLECs display the lowest cytoTRACE scores (most differentiated). Based on the cytoTRACE analysis, and its logic of developmental potential, it would seem unlikely that LN_CLEC are the origin of T_CLEC (Fig. 7e-f). Moreover, it appears that both lineage positive and negative EC in the lymph nodes are more differentiated than endothelial cells in the tumour. It is noteworthy that Monocle was designed for developmental questions, which are strictly unidirectional. Interestingly, when plotting CytoTRACE scores onto the Monocle trajectory (Fig. 7f), the right side shows higher scores (T_CLEC and T_EC) and the left side (darker blue) lower scores (LN_CLECs lowest), being in line with pseudotime. This again predicts that LN_CLECs represent an end state rather than the origin. In summary, these data predict that during tumour growth, the T_EC and T_CLECs coevolve from a distinct origin to then share many aspects of their gene expression and biology, yet with a few very distinct gene profiles.

## DISCUSSION

In this study, using inducible *Csf1r* lineage tracing in adult mice intended to study macrophage recruitment during glioma development, we identify a distinct endothelial cell lineage that forms up to 10% of the tumour vasculature in adult mice. Selective experimental depletion of this *Csf1r* endothelial cell lineage demonstrated a role in promoting vascular branching and tumour growth. Through a combination of genetic lineage tracing, longitudinal live two-photon imaging, and single cell analysis, we find that these cells arise during tumour progression, adopt dynamic endothelial phenotypes including tip cell characteristics, and express common endothelial as well as unique markers that can also be found in patient-derived tumour endothelium.

*Csf1r* is a well-known regulator of monocyte/ macrophage differentiation ^38, 39^, but is also important for yolk sac hematopoiesis and progenitor cell differentiation. Klotz and coworkers reported that *Csf1r*-expressing cells in the mouse embryo contribute to *Prox1*-positive lymphatic endothelial cells in the cardiac lymphatic vasculature ^40^. Moreover, a recent report identified that *Csf1r*-expressing erythro-myeloid progenitors from the early embryonic yolk-sac could give rise to intraembryonic endothelial cells in mouse development^41^. *Csf1r* is also expressed in EPCs, such as colony forming unit-Hill cells and circulating angiogenic cells ^42^. In the adult however, *Csf1r* is not expressed in the mature endothelium of blood and lymphatic vessels, but continues to drive recruitment and expansion of myeloid cells in inflammation, ischemia and tumour growth ^43^. Intriguingly, *Csf1r* itself is no longer expressed in the Csf1r lineage cells within the tumour and acute recombination of endothelial cells 24h after tamoxifen injection fails to label cells in the tumour vasculature. These results raise the possibility that a *Csf1r-expressing cell population* of endothelial nature or potential is present elsewhere in the body, recombines upon tamoxifen injection and is recruited to the tumour vasculature where it differentiates into blood endothelium to contribute to tumour angiogenesis. Alternatively, the fact that Csf1r expression is not detected in isolated tumour CLECs could also indicate that the Cre-line is leaky and gives rise to labelled cells through stochastic recombination. In that case, one would expect the labelled cells to populate any part of the vasculature and be otherwise indistinguishable from the unlabelled tumour EC. Single cell analysis however identifies a unique gene expression signature beyond the large number of genes that are co-expressed in T_CLEC and T_EC, and demonstrates that T_CLEC are distinctly enriched in tip-cell and neo-phalanx cell populations. Thus, labelled cells show distinct phenotypes, location and expression patterns, ruling out purely stochastic recombination events. Furthermore, such stochastic recombination within the tumour vasculature would necessitate that we identify acutely labelled cells marked by co-expression of Tomato and GFP. As we failed to identify such cells at any timepoint during tumour growth, and also could detect the emergence of T_CLECs in mice only injected with tamoxifen before tumour implantation, the cumulative evidence would suggest that T_CLECs originate outside of the tumour and also that the progenitor population resides outside of the brain. Despite all our efforts, the present work fails to provide ultimate and definitive proof of the true origin of T_CLECs, but arrives at a clear direction and working hypothesis for future studies. In search of their origin and a potential progenitor population, we imaged and analysed all organs by FACS. We performed bone-marrow transplantation, performed splenectomy and examined the entire mouse by fluorescence stereomicroscopy after clearing to identify regions of recombination. The results of bone-marrow transplantation effectively ruled out EPC or any other known progenitors that may be found in circulation, including transdifferentation from macrophages. Four independent observations placed a spotlight on lymphatics, and the lymph nodes in particular. First, the only *bonafide* Csf1r lineage endothelial cell population that we could identify outside of the tumours, including after complete reconstitution of the bone marrow by wildtype bone marrow, was found in the subcapsular sinus of the lymph nodes. Here the floor cells, specialized lymphatic endothelial cells that we identify as GFP and Csf1r positive line the capsule in lymph nodes in close apposition to the Csf1 expressing ceiling cells. Second, after acute tamoxifen injection the cells in the floor of the subcapsular sinus are uniquely double positive for Tomato and GFP, demonstrating acute and likely local recombination. Third, single cell RNAseq of T-CLECs identified lymphatic markers including the lymphatic master regulator Prox1 co-expressed with blood endothelial markers, and SCDE placed T_CLECs close to lymphatic endothelial cells from various mouse tumour models in recently published datasets. And finally, CLECs can also be found in lymphatics in subcutaneous tumour models and in skin. These independent observations clearly make the SCS floor cells prime candidates for the origin of T_CLECs, but future work will be necessary for a possible direct demonstration of this link.

In developmental studies, lineage relationships have recently been studied by scRNAseq data sets through pseudotime analysis. In particular the CytoTRACE analysis can be useful to estimate the degree of differentiation of cell populations and thereby also predict their stemness ^37^. Our comparative analysis of endothelial cells isolated from tumour and lymph nodes assigned the highest differentiation scores to cells in the lymph node, and lower values to the EC in tumour, including T_CLECs. Accordingly, the cells in the floor of the SCS that carry the lineage label would rather be assumed at the end of a differentiation process rather than progenitors for other populations such as the T_CLECs. Whether however such algorithms are suitable to determine the potential of endothelial cells to become activated and possibly transdifferentiate is unclear. In particular given the altered environmental context of EC within a growing tumour compared to a vascular bed in homeostasis. The combined monocle^36^ and cytoTRACE analysis would fit with a general level of dedifferentiation of endothelial cells within the tumour microenvironment, and therefore a co-evolution of the cell states of both T_EC and T_CLEC. It is likely that pseudotemporal ordering of endothelial cells in a reactivation state will always show the reactivated cells, like in the tumour environment as less differentiated. Indeed, recent work studying the effect of chronic hypoxia on brain endothelium illustrated the wide range of reactivation, in particular noting a strong increase in tip cell population amongst the brain endothelial cells^44^. The current assumption states that these new tip cells arise under hypoxia from pre-existing brain endothelial cells, and possibly emerge from particular endothelial subpopulations. In the absence of lineage tracing, an alternative possibility, raised by our current observation in tumour angiogenesis, could be that new tip cells arise at least in part elsewhere, and are recruited to the hypoxic vessel areas. More work on vascular single cell analysis will be required to establish the fundamental principles of endothelial activation and their interpretation for the different tissue challenges that require vascular adaptations.

Angiogenesis is essential for tumour growth, and contributes to metastasis ^45, 46^. Early hopes that targeting tumour angiogenesis may prove uniquely selective and effective as anti-tumour treatment have, however, been disappointing. Intense research efforts therefore currently focus on understanding reasons and mechanisms for resistance to anti-angiogenic therapy, and identifying new cellular and molecular targets to effectively modify the tumour vasculature ^47^. Intravital imaging shows that T_CLECs arise during tumour progression, adopt endothelial phenotypes with tip cell characteristics and line perfused vascular tubes. Consistent with this observation, AUCell analysis of scRNA-seq data in human samples identifies that T_CLECs signatures are highly enriched in tip cell subpopulation. A recent study using single-cell transcriptome analysis revealed that anti-VEGF (vascular endothelial growth factor) treatment reduces the proportion of endothelial tip-like cells in a tumour xenografts ^48^. How CLECs affect the outcome of anti-angiogenic treatment will be important to investigate in the future. Considering that T_CLECs prominently express *Vegfr3*, even more so than regular endothelial cells in the tumour, we can speculate that T_CLECs are resistant to anti-VEGFR2 treatment, and possibly escape treatment through the activity of VEGFR3. If so CLECs may adversely affect anti-angiogenic therapy targeting VEGFR2 selectively. Tumour blood vessels are generally highly tortuous and the branching pattern is greatly different from normal blood vessels. Our results from the selective targeting of CLECs in the mouse glioma model show that CLECs ablation decreases both vessel branching and tumour growth. Thus, in the absence of CLEC, the dysmorphic vascular phenotype in tumours appears exacerbated, suggesting that CLECs support the formation of a highly branched vascular network, which in turn better promotes tumour growth. Future work will be required to identify the true origin of T_CLECs and to extend this functional analysis to fully appreciate the consequence and opportunities to target or utilize CLECs for therapeutic approaches.

## METHODS

### Mice

Mice expressing both the *Csf1r-Mer-iCre-Mer* transgene (*Csf1r-Mer2.Cre*) ^17^ and the *Rosa^mTmG^* Cre recombination reporter ^18^ were generated by breeding. Mice carrying the. Endothelial specific *Cdh5-Dre* mice ^24^ were mated with *Csf1r-Mer2.Cre* mice and *Rosa26-RSR-LSL-tdTomato* mice ^23^ to obtain *Cdh5-Dre*:: *Csf1r-Mer2.Cre*:: *Rosa26-RSR-LSL-tdTomato* mice (*DreMer2.Cre^tdTomato^*) that allows combinatorial genetics as a tool for CLECs tracing. Endothelial specific *Cdh5-Dre* mice were mated with *Csf1r-Mer2.Cre* mice and *Rosa26-RSR-mTmG-DTR* mice to obtain *Cdh5-Dre*:: *Csf1r-Mer2.Cre*:: *Rosa26-RSR-mTmG-DTR* mice (*DreMer2.Cre^mTmG-DTR^*) that allows for selective tracing and ablation of CLECs. *Rosa26-RSR-mTmG-DTR* mice were designed by Dr. Fabio Stanchi and generated by the team of Dr. Ralf Kühn at the Max Delbrück Center in Germany. All mouse strains were maintained on C57BL/6J background. Animal housing and all experimental procedures were approved by the Institutional Animal Care and Research Advisory Committee of the KU Leuven (085/2016).

### Cell lines

CT2A, GL261 glioma cell lines and B16F1 melanoma cell line were cultured in DMEM (Life Technologies) supplemented with 10% FBS (Life Technologies), 1% penicillin/ streptomycin (Life Technologies) and 1% glutamine (Life Technologies). Spheroids were obtained by seeding the glioma cells for 48h on non-adherent culture dishes. Spheroids of 200-250µm were selected for implantation.

### Intravital imaging on mouse gliomas

Surgery for tumor implantation, installation of cranial windows and *in vivo* imaging were performed as described previously ^16^. Briefly, 8– 12-weeks-old mice were anesthetized with ketamine/ xylazine and a craniotomy was performed on the parietal bone. Spheroids of CT2A or GL261 cells modified to express the blue fluorescent protein TagBFP ^50^ were injected in the exposed parietal brain cortex, which was then sealed by cementing a glass coverslip to the bone surrounding the craniotomy. After surgery, mice were injected intraperitoneally with tamoxifen (65 μg/g body weight) every 5 days. For CLECs depletion, mice were injected intraperitoneally with diphtheria toxin (4 ng/g body weight) the next day after tamoxifen induction. Longitudinal *in vivo* imaging of growing tumors was performed in mice under isoflurane anesthesia using a SP8 upright microscope (Leica Microsystems) equipped with a HCX IRAPO L25x/0.95 water objective and a Titanium: Sapphire laser (Vision II, Coherent Inc.) tuned at 925 nm. Humane end-point of experiments was applied if animals lost 15-20% of their original weight or shown evident signs of distress.

### Immunofluorescence imaging

Mice anesthetized with ketamine/ xylazine were perfused through the heart with 15 ml of ice-cold PBS, followed by 10 ml of 2% PFA in PBS. Tumour tissues or organs were harvested and fixed overnight in 4% PFA. For tumour tissues, sections (200 µm-thick) were prepared with a vibratome 650V (Thermo Scientific). For organ samples, frozen sections (10 µm-thick) were prepared with a cryostat NX70 (Thermo Scientific). For immunofluorescence imaging studies, the PFA-fixed sections were blocked and permeabilized in TNBT (0.1 M Tris, pH 7.4, 150 mM NaCl, 0.5% blocking reagent from Perkin Elmer, 0.5% Triton X-100) for 4h at room temperature. Sections were incubated with antibodies against CDH5 (1:25, AF1002, R&D), CD31 (1:100, ab28364, Abcam), F4/80 (1:100, MF48000, Invitrogen), CD45 (1:100, 550539, BD Pharmingen), CD169 (1:100, 142401, Biolegend), PROX1 (1:200, 11-002, Angiobio), AQP1 (1:4000, AB2219, Millipore), LYVE1 (1:100, 50-0433-80, eBioscience) diluted in TNBT buffer overnight at 4°C, washed in TNT buffer (0.1 M Tris pH 7.4; 150 mM NaCl, 0.5% Triton X-100) and incubated with an Alexa Fluor conjugated antibodies (1:200, ThermoFisher Scientific). Sections were washed and mounted in fluorescent mounting medium (Dako). Images were obtained with a Leica TCS SP8 confocal microscope.

### Flow cytometric analysis

Mice were anesthetized with ketamine/ xylazine, then tumour tissues or organs were harvested and incubated with PBS containing 1mg/ml collagenase I (Gibco), 2 mg/ ml Dispase I (Gibco), 100ug/ml DNase I (Roche) and 2 mM CaCl2 for 30 min at 37 °C. After incubation, the digested tissue was passed through a cell strainer and then washed by PBS including 2% FBS. For red cell exclusion, we incubated samples with red cell lysis buffer (Sigma) for 5 min at 37 °C and then washed by PBS including 2% FBS and 2 mM EDTA. Cells were stained with the following monoclonal antibodies: PE/ Cy7 anti-CD45 (552848, BD), APC anti-CD31 (561814, BD). Data acquisition was performed with BD FACSVerse and analysis was performed with BD FACSuite software, and FlowJo software.

### Mouse melanoma model

Mice (8–12-weeks-old) were anesthetized with ketamine/ xylazine and then 1×10^6^ B16F1 melanoma cells in PBS were implanted into the right flank region of the mice. B16F1 melanomas were removed from mice 12 days after implantation for immunofluorescent imaging studies and flow cytometric analysis. Mice were injected intraperitoneally with tamoxifen (65 μg/g body weight) 4 time/ 2 weeks before tumour implantation and then intraperitoneally with tamoxifen (65 μg/g body weight) every 5 days after tumour implantation.

### Single cell isolation and RNA-sequencing

Single cells were sorted (BD FACSAriaIII) in 96 well plates (VWR) containing 2 μL of PBS including 0.2% Triton X-100 and 4U of RNase inhibitor (Takara) per well. Plates were properly sealed and spun down at 2000 g for 1 min before storing at −80°C. Whole transcriptome amplification was performed with a modified SMART-seq2 protocol as described previously ^25^, using 20 instead of 18 cycles of cDNA amplification. PCR purification was realized with a 0.8:1 ratio (ampureXP beads:DNA). Amplified cDNA quality was monitored with a high sensitivity DNA chip (Agilent) using the Bioanalyzer (Agilent). Sequencing libraries were performed using the Nextera XT kit (Illumina) as described previously ^25^, using 1/4th of the recommended reagent volumes and 1/5th of input DNA with a tagmentation time of 9 min. Library quality was monitored with a high sensitivity DNA chip (Agilent) using the Bioanalyzer (Agilent). Indexing was performed with the Nextera XT index Kit V2 (A-D). Up to 4 x 96 single cells were pooled per sequencing lane. Samples were sequenced on the Illumina NextSeq 500 platform using 75bp single-end reads.

### scRNA-sequencing data analysis

BAM files were converted to merged, demultiplexed FASTQ files, cleaned using fastq-mcf (ea-utils r819), and QC checked with FastQC (0.11.4). Reads were then mapped to the mouse genome (mm18) using STAR (2.4.1b) and quantified with Subread (1.4.6-p2). Cells with less than 100,000 reads and/or 500 genes expressed, more than 20% mitochondrial reads, and less than an average expression level of 3.0 of about 80 housekeeping genes ^51^ were discarded. 469 cells passed these stringent quality criteria. Winsorized Highly Variable genes (HVGs) were identified for tumor derived macrophages (T_MACs), endothelial cell (T_ECs) and *Csf1r* lineage endothelial cells (T_CLECs) ^26^. These cells were then clustered based on HVG expression using non-negative matrix factorization as dimension reduction approach (run=40, rank=10, in MeV 4.8.1). The “best fit” (numbers of clusters) was chosen based on the highest cophenetic correlation coefficient. Next, Single-cell Differential Expression analysis (SCDE) was performed between the different NMF-clusters using the global gene expression matrix^26^. Expression values were library-size-factor-normalized (DESeq) and log2 transformed (log2+1). T_MAC, T_EC, T_CLEC and LN_CLECS, LN_EC, T_CLEC, T_EC cells were projected into a two-dimensional space using Uniform Manifold Approximation and Projection (dims=15, resolution = 0.4) based on 5k most variable features (Seurat pipeline) ^52^. Monocle (2.14) based trajectory analysis was performed on T_CLEC, T_EC, LN_CELC and LN_EC (total=454 cells) taking into account n=790 m3Drop genes. The same data set was subjected to CytoTRACE analysis using the online tool (https://cytotrace.stanford.edu/) ^37^.

### AUCell analysis of human scRNA-seq data in multiple tumours

The droplet-based scRNA-seq data of 4 human tumour types were downloaded from publicly available sources including lung cancer (ArrayExpress: E-MTAB-6149 and E-MTAB-6653)^53^, PDAC (GSA: CRA001160)^54^, liver cancer (GEO: GSE125449)^55^, and BCC (GEO: GSE123814)^56^. These datasets were processed and clustered per tumour type using Seurat (v3.0.4) package, and the endothelial cell clusters were identified using marker genes (*CLDN5*, *VWF*, *PECAM1*). The resulting endothelial cell data were pooled together and aligned using anchor-based canonical correlation analysis (CCA) to mitigate the difference related to distinct tumor types and scRNA-seq technologies. Initial graph-based clustering identified 10 clusters including 3 potential doublet clusters as predicted by DoubleFinder (v2), which were subsequently removed before second round of clustering. The resulting 8 clusters consisting of 7314 endothelial cells were annotated based on known marker genes, and their CLECs signature were calculated using AUCell package (v1.6.1). The threshold of AUC score was chosen at the peak of the right modal, and the percentage of putative CLECs were calculated for different subtypes and their tumour type of origin.

### Generation of bone marrow chimeras

Recipient 8-10-weeks-old mice were lethally irradiated (9.5 Gy) and then intravenously injected with 1×10^7^ bone marrow cells from donor mice 16 h later. Tumour implantation experiments were initiated 8 weeks after bone marrow reconstitution. Reconstitution rate was determined using flow cytometric analysis in the peripheral blood.

### Surgical removal of the spleen

Mice (8–12-weeks-old) were anesthetized with ketamine/ xylazine and then the spleen was gently taken out from the connective tissue, while cauterizing associated vessels. The incision was thereafter sutured. These procedures were performed as described previously^57^.

### Vessel morphology analysis

Confocal images of the tumour vasculature were analyzed for two-dimensional vessel area, network length and number of network bifurcation points. The tdTomato and GFP channels were merged and a Gaussian blur filter as well as Otsu thresholding was applied to obtain binary vessel mask for sectional area estimation. To obtain the vessel network, a semi-automatic multicut workflow from ilastik ^58^ was applied to all images. The segmented vessels were skeletonized and converted into a network graph with nodes and edges by using the Python package ImagePy ^59^. The lengths of network edges were summed up to obtain the total network length for each image, all nodes with a degree of three or more were counted as bifurcations.

### Statistics

Statistical analyses of data were performed with GraphPad Prism 7.0 software. All data are shown as the means ± standard deviations. Differences were assessed using a two-tailed unpaired Mann-Whitney’s U test or two-tailed unpaired Welch’s correction.

### AUTHOR CONTRIBUTIONS

KM led and conceived the project, designed and performed experiments, analysed and interpreted data. FR, FS, TM, WG, LH, JQ performed experiments and analyzed data. DL, BZ, CB, JM and HG analyzed data. HG conceived and supervised the project. KM and HG wrote and edited the manuscript, with contribution by all authors.

## ACKNOWLEDGEMENT

This work was supported by the Belgian Cancer Foundation (Stichting Tegen Kanker, grant 2012-181, 2018-074) and a Hercules type 2 grant (Herculesstichting: AKUL11033). K.M. is a recipient of a Japan Society for the Promotion of Science Overseas Research Fellowship. We would like to thank Dr. Till Acker (Institute of Neuropathology, University of Giessen, Germany) and Dr. Thomas N. Seyfried (Biology department, Boston College, USA) for the gift of GL261 and CT2A cells, respectively. We are grateful for outstanding advice and service by Dr. Ralf Kühn heading the Transgenics Core Facility at the Max-Delbrueck-Center, Berlin, Germany. We would like to thank our colleagues Marly Balcer, Lisse Decraecker, Greet Bervoets and Irene Hollfinger for excellent technical assistance.

## CODE AVAILABILITY

The computational code (R) used for scRNA-seq analysis is based on already published pipelines (see Material and Methods) and available upon request.

## DATA AVAILABILITY

All scRNA-seq data are available at Gene Expression Omnibus under accession number GSE157507.

## STATISTICS

Differences were assessed using either a two-tailed unpaired Mann-Whitney’s or two-tailed unpaired t-test with Welch correction. All planned comparisons were considered stand-alone and were not corrected for multiple testing.

## FIGURE LEGENDS

**Extended Data Fig. 1: CLECs in mouse glioma express endothelial cell markers, but lack macrophage and myeloid cell markers.** Counterstaining on PFA-fixed 4 weeks CT2A glioma sections of Csf1r-Mer2.CremTmG with indicated antibodies. Lower panels show the detail images of outlined squares in upper panels. Yellow arrowheads, CLECs; blue arrowheads, macrophages. Scale bar: 25 μm.

**Extended Data Fig. 2: CLECs contribute to lymphatic endothelium in the tumour environment.** Fluorescent images of counterstaining on PFA-fixed section of day 12 B16F1 melanoma under ear skin of Csf1r-Mer2.CremTmG mice using LYVE1 antibody. Lower panel indicates detail image of yellow outlined square in upper panel. Yellow arrow heads, CLECs. Scale bar: 100 μm.

**Extended Data Fig. 3: Reconstitution rate in bone marrow chimeras**. Reconstitution rate was determined in peripheral blood 8 weeks after bone marrow transplantation. TdTomato-positive cells in blood of: C57BL/6J (0.3 ± 0.1%, n= 6), Csf1r-Mer2.CremTmG (99.2 ± 0.5%, n= 7), BMCsf1r-Mer2.CremTmG (97.3 ± 1.6%, n= 13). Bars represent mean ± s.d. ****p<0.0001. Two-tailed unpaired Mann-Whitney’ s U test.

**Extended Data Fig. 4: CLECs do not originate from spleen.** Surgical removal of spleen (splenectomy) was performed 3 weeks before glioma injection. CLECs isolated from CT2A glioma of sham control Csf1r-Mer2.CremTmG mice (n= 5) constitute 1.5 ± 0.8 % of total tumour endothelial cell population, compared to 1.5 ± 0.5% found in Csf1r-Mer2.CremTmG with splenectomy (n= 4). Bars represent mean ± s.d. Two-tailed unpaired Mann-Whitney’ s U test.

**Extended Data Fig. 5: CLECs locate at peripheral area of mouse glioma**. a, Vibratome section (200 μm-thick) of 4 weeks CT2A glioma of Csf1r-Mer2.CremTmG:: BMWT mice. Scale bar: 1 mm. b, Fluorescent images of area outlined in a. Yellow arrowheads, CLECs. Scale bar: 100 μm.

**Extended Data Fig. 6: 24 tumour CLECs markers in mouse glioma.** SCDE analysis of scRNA-seq data of tumour macrophages (T_MACs: CD45+GFP+, n= 79), tumour endothelial cells (T_ECs: CD45-CD31+GFP-, n= 78), tumour CLECs (T_CLECs: CD45-CD31+GFP+, n= 141) in 4 weeks CT2A glioma in Csf1r-Mer2.CremTmG mice. Bars represent mean ± s.d. *p<0.05, **p<0.01, ***p<0.001, ****p<0.0001. Two-tailed unpaired t-test with Welch correction.

**Extended Data Fig. 7: CLECs in glioma express AQP1**. Counterstaining on PFA-fixed CT2A glioma of Csf1r-Mer2.CremTmG::BMWT mice using AQP1 antibody. Yellow arrowheads, T_CLECs; blue arrowheads, T_ECs. Scale bar: 25 μm.

**Extended Data Fig. 8: CLECs in glioma cannot be labelled by GFP within 24 hours after tamoxifen induction.** a-b, Tamoxifen induction timing. b, Quantification of the population of tumour macrophagens (T_MACs: CD45+GFP+) and tumour CLECs (T_CLECs: CD45-CD31+GFP+) by flow cytometric analysis in tumour samples (n=4). Bars represent mean ± s.d. c, Fluorescent images of PFA-fixed 3 weeks CT2A glioma de of Csf1r-Mer2.CremTmG mice within 24 hours after tamoxifen induction. Scale bar: 100 μm.

**Extended Data Fig. 9: CLEC and EC heterogeneity in tumor and lymph node.** a, Heatmap shows the expression of n=91 fLECs markers in LN_CLEC, LN_EC, T_CLEC and T_EC single cells. b, Violin plots show the expression of 3 CLEC markers: Pde4d, Robo1 and Itga9.

**Supplemental Movie 1: Time-lapse *in vivo* imaging of Csf1r lineage endothelial tip cell in CT2A glioma of *Csf1r-Mer2.Cre^mTmG^* mice.** Imaging was performed over 4 hours (1 frame/ 10 min).

**Supplemental Movie 2: Time-lapse *in vivo* imaging of *Csf1r* lineage endothelial tip cell in CT2A glioma of *Csf1r-Mer2.Cre^mTmG^*::BM^WT^.** Imaging was performed over 50 minutes (1 frame/ 2.5 min).

**Supplemental Table 1: Details of gene expression analysis.** Seurat based differential analysis (non-parameteric Wilcoxon rank sum test) was performed by comparing cell types amongst each other (green columns) and the different Seurat clusters (yellow columns). p_val: p_val (unadjusted). avg_logFC: log fold-change of the average expression between the two groups. Positive values indicate that the feature is more highly expressed in the first group. pct.1: The percentage of cells where the feature is detected in the first group. pct.2: The percentage of cells where the feature is detected in the second group. p_val_adj: Adjusted p-value, based on Bonferroni correction using all features in the dataset.

**Supplemental Table 2: Marker genes for EC subtypes of multiple tumour types.** p_val: p_val (unadjusted). avg_logFC: Average log (fold change). pct.1: Percentage of expressed cells in current cluster. pct.2: Percentage of expressed cells in other cluster. p_val_adj: Adjusted p-value. cluster: Cluster name. gene: Gene name. log2fc: log2 (fold change). Log2pct: log2 ((pct+0.005)/(pct2+0.005)).

## Notes

Conflict of Interest: The authors have declared that no conflict of interest exists.

### Competing Interest Statement

The authors have declared no competing interest.

