## Supplementary Movie 2 for "Emerging single cell endothelial heterogeneity supports sprouting tumour angiogenesis and growth"

### Supplementary Table 1

| gene | p_val | avg_logFC | pct.1 | pct.2 | p_val_adj | cluster | gene |
|---|---|---|---|---|---|---|---|
| Lilrb4a | 1,77E-69 | 2,400385012 | 1 | 0,004 | 2,80E-65 | MAC | Lilrb4a |
| Bcl2a1b | 1,77E-69 | 2,189161117 | 1 | 0,004 | 2,80E-65 | MAC | Bcl2a1b |
| Mpeg1 | 5,11E-69 | 2,010932239 | 0,988 | 0 | 8,10E-65 | MAC | Mpeg1 |
| Tyrobp | 6,88E-69 | 2,918688965 | 1 | 0,008 | 1,09E-64 | MAC | Tyrobp |
| Lilr4b | 2,46E-68 | 2,329919561 | 0,988 | 0,004 | 3,90E-64 | MAC | Lilr4b |
| Fcgr3 | 2,59E-68 | 2,967713703 | 1 | 0,012 | 4,10E-64 | MAC | Fcgr3 |
| Cd52 | 2,59E-68 | 2,278192308 | 1 | 0,012 | 4,10E-64 | MAC | Cd52 |
| Aif1 | 2,59E-68 | 2,103100042 | 1 | 0,012 | 4,10E-64 | MAC | Aif1 |
| Ly86 | 5,88E-68 | 1,946878729 | 0,975 | 0 | 9,32E-64 | MAC | Ly86 |
| Spi1 | 5,88E-68 | 1,790833527 | 0,975 | 0 | 9,32E-64 | MAC | Spi1 |
| Ifi30 | 1,13E-67 | 2,182814707 | 1 | 0,017 | 1,80E-63 | MAC | Ifi30 |
| Apobec1 | 6,67E-67 | 1,931855482 | 0,963 | 0 | 1,06E-62 | MAC | Apobec1 |
| C1qa | 1,13E-66 | 3,788283771 | 1 | 0,025 | 1,80E-62 | MAC | C1qa |
| Fcgr2b | 1,13E-66 | 2,744112754 | 1 | 0,025 | 1,80E-62 | MAC | Fcgr2b |
| Laptm5 | 1,68E-66 | 2,361074799 | 1 | 0,025 | 2,67E-62 | MAC | Laptm5 |
| Csf1r | 3,76E-66 | 2,692010077 | 1 | 0,029 | 5,96E-62 | MAC | Csf1r |
| Pla2g7 | 4,48E-66 | 2,385132456 | 0,988 | 0,021 | 7,09E-62 | MAC | Pla2g7 |
| C1qc | 1,21E-65 | 4,125327152 | 1 | 0,033 | 1,92E-61 | MAC | C1qc |
| Coro1a | 1,37E-65 | 2,157053059 | 1 | 0,029 | 2,18E-61 | MAC | Coro1a |
| Trem2 | 3,33E-65 | 1,787857377 | 0,951 | 0,004 | 5,28E-61 | MAC | Trem2 |
| C3ar1 | 8,20E-65 | 1,837999673 | 0,938 | 0 | 1,30E-60 | MAC | C3ar1 |
| Cd300c2 | 8,20E-65 | 1,436111737 | 0,938 | 0 | 1,30E-60 | MAC | Cd300c2 |
| Pld4 | 1,34E-64 | 2,292815016 | 0,951 | 0,008 | 2,13E-60 | MAC | Pld4 |
| Rgs1 | 3,61E-64 | 3,074974665 | 0,938 | 0,004 | 5,73E-60 | MAC | Rgs1 |
| Itgam | 3,61E-64 | 1,531582192 | 0,938 | 0,004 | 5,73E-60 | MAC | Itgam |
| Cybb | 5,92E-64 | 1,695465599 | 0,951 | 0,012 | 9,39E-60 | MAC | Cybb |
| Ms4a7 | 6,11E-64 | 1,94153161 | 0,951 | 0,012 | 9,68E-60 | MAC | Ms4a7 |
| Cd68 | 1,05E-63 | 2,337275647 | 1 | 0,05 | 1,67E-59 | MAC | Cd68 |
| C1qb | 2,80E-63 | 4,258745352 | 1 | 0,054 | 4,44E-59 | MAC | C1qb |
| Ms4a6c | 6,25E-63 | 1,450614627 | 0,938 | 0,012 | 9,90E-59 | MAC | Ms4a6c |
| Lyz2 | 7,73E-63 | 4,475989832 | 1 | 0,058 | 1,23E-58 | MAC | Lyz2 |
| Ctss | 7,73E-63 | 4,33950261 | 1 | 0,058 | 1,23E-58 | MAC | Ctss |
| Cyp4f18 | 9,52E-63 | 1,741252522 | 0,914 | 0 | 1,51E-58 | MAC | Cyp4f18 |
| Ptprc | 9,52E-63 | 1,496797575 | 0,914 | 0 | 1,51E-58 | MAC | Ptprc |
| Alox5ap | 2,76E-62 | 1,886439132 | 0,963 | 0,025 | 4,38E-58 | MAC | Alox5ap |
| Sirpb1c | 4,34E-62 | 1,470039352 | 0,914 | 0,004 | 6,88E-58 | MAC | Sirpb1c |
| Rgs10 | 1,00E-61 | 1,276072511 | 0,901 | 0 | 1,59E-57 | MAC | Rgs10 |
| Cd53 | 1,57E-61 | 1,85537111 | 0,988 | 0,045 | 2,49E-57 | MAC | Cd53 |
| Ptpn6 | 2,74E-61 | 1,607331114 | 0,914 | 0,008 | 4,35E-57 | MAC | Ptpn6 |
| Clec4a2 | 4,65E-61 | 1,513908845 | 0,901 | 0,004 | 7,37E-57 | MAC | Clec4a2 |
| Lrp1 | 5,38E-61 | 1,668389423 | 0,914 | 0,008 | 8,53E-57 | MAC | Lrp1 |
| Fermt3 | 6,14E-61 | 1,49298526 | 0,901 | 0,004 | 9,73E-57 | MAC | Fermt3 |
| Csf2ra | 7,19E-61 | 1,317595911 | 0,938 | 0,025 | 1,14E-56 | MAC | Csf2ra |
| Ms4a4a | 1,04E-60 | 1,521905739 | 0,889 | 0 | 1,65E-56 | MAC | Ms4a4a |
| Ccl6 | 1,04E-60 | 1,516975938 | 0,889 | 0 | 1,65E-56 | MAC | Ccl6 |
| Fcgr4 | 1,12E-60 | 1,79610778 | 0,914 | 0,008 | 1,77E-56 | MAC | Fcgr4 |
| Fcer1g | 2,14E-60 | 3,44213885 | 1 | 0,083 | 3,40E-56 | MAC | Fcer1g |
| Adgre1 | 4,91E-60 | 1,58175684 | 0,889 | 0,004 | 7,79E-56 | MAC | Adgre1 |
| Wfdc17 | 8,83E-60 | 2,040118947 | 0,901 | 0,012 | 1,40E-55 | MAC | Wfdc17 |
| Clec12a | 1,07E-59 | 1,508464223 | 0,877 | 0 | 1,69E-55 | MAC | Clec12a |
| Rac2 | 2,29E-59 | 1,650161882 | 0,889 | 0,008 | 3,63E-55 | MAC | Rac2 |
| Fam105a | 5,12E-59 | 1,499573611 | 0,877 | 0,004 | 8,11E-55 | MAC | Fam105a |
| Itgb2 | 1,08E-58 | 1,545172129 | 0,864 | 0 | 1,71E-54 | MAC | Itgb2 |
| Lair1 | 1,08E-58 | 1,427215962 | 0,864 | 0 | 1,71E-54 | MAC | Lair1 |
| Cxcl16 | 2,81E-58 | 2,227180165 | 1 | 0,091 | 4,45E-54 | MAC | Cxcl16 |
| C5ar1 | 5,41E-58 | 1,676653978 | 0,864 | 0,004 | 8,58E-54 | MAC | C5ar1 |
| Clec4n | 1,07E-57 | 1,596936596 | 0,852 | 0 | 1,70E-53 | MAC | Clec4n |
| Ctsc | 3,48E-57 | 2,919063102 | 0,988 | 0,103 | 5,51E-53 | MAC | Ctsc |
| Msr1 | 1,05E-56 | 1,516472413 | 0,84 | 0 | 1,67E-52 | MAC | Msr1 |
| Slamf9 | 1,05E-56 | 1,434402142 | 0,84 | 0 | 1,67E-52 | MAC | Slamf9 |
| Arrb2 | 1,37E-56 | 0,863955106 | 0,901 | 0,029 | 2,17E-52 | MAC | Arrb2 |
| Fcgr1 | 5,30E-56 | 1,453640018 | 0,84 | 0,004 | 8,40E-52 | MAC | Fcgr1 |
| Cd300lf | 6,17E-56 | 1,405426153 | 0,84 | 0,004 | 9,78E-52 | MAC | Cd300lf |
| Axl | 8,64E-56 | 1,375333071 | 0,889 | 0,021 | 1,37E-51 | MAC | Axl |
| Acp5 | 1,54E-55 | 1,861950181 | 0,926 | 0,05 | 2,45E-51 | MAC | Acp5 |
| Fcrls | 5,54E-55 | 1,99176459 | 0,827 | 0,004 | 8,78E-51 | MAC | Fcrls |
| Plek | 6,64E-55 | 1,260735668 | 0,827 | 0,004 | 1,05E-50 | MAC | Plek |
| Nckap1l | 7,65E-55 | 1,31726559 | 0,864 | 0,017 | 1,21E-50 | MAC | Nckap1l |
| Cx3cr1 | 9,74E-55 | 1,825864762 | 0,815 | 0 | 1,54E-50 | MAC | Cx3cr1 |
| Sla | 9,74E-55 | 1,434527979 | 0,815 | 0 | 1,54E-50 | MAC | Sla |
| Mrc1 | 1,75E-54 | 1,504749087 | 0,852 | 0,017 | 2,77E-50 | MAC | Mrc1 |
| Cd33 | 3,53E-54 | 1,347542742 | 0,852 | 0,017 | 5,59E-50 | MAC | Cd33 |
| Selplg | 5,30E-54 | 1,472375391 | 0,84 | 0,017 | 8,41E-50 | MAC | Selplg |
| Ccr5 | 2,44E-53 | 1,4336233 | 0,84 | 0,021 | 3,87E-49 | MAC | Ccr5 |
| Tlr13 | 9,56E-53 | 1,074220537 | 0,827 | 0,012 | 1,52E-48 | MAC | Tlr13 |
| Lpcat2 | 3,41E-52 | 1,535071916 | 0,852 | 0,025 | 5,40E-48 | MAC | Lpcat2 |
| Hck | 4,57E-52 | 1,217824949 | 0,79 | 0,004 | 7,24E-48 | MAC | Hck |
| Ccr1 | 5,34E-52 | 1,699117124 | 0,815 | 0,017 | 8,47E-48 | MAC | Ccr1 |
| Evi2a | 8,08E-52 | 1,302358918 | 0,852 | 0,033 | 1,28E-47 | MAC | Evi2a |
| H2-DMa | 1,01E-51 | 1,603167016 | 0,852 | 0,029 | 1,60E-47 | MAC | H2-DMa |
| Ms4a6b | 1,44E-51 | 1,698284467 | 0,864 | 0,037 | 2,29E-47 | MAC | Ms4a6b |
| Mt1 | 2,78E-51 | 1,720681994 | 0,975 | 0,132 | 4,41E-47 | MAC | Mt1 |
| Cd84 | 2,98E-51 | 1,193123543 | 0,827 | 0,025 | 4,73E-47 | MAC | Cd84 |
| Ccl3 | 7,07E-51 | 2,044184471 | 0,765 | 0 | 1,12E-46 | MAC | Ccl3 |
| Slc11a1 | 7,07E-51 | 1,510747551 | 0,765 | 0 | 1,12E-46 | MAC | Slc11a1 |
| Ncf4 | 7,07E-51 | 1,170621712 | 0,765 | 0 | 1,12E-46 | MAC | Ncf4 |
| Gpr65 | 7,07E-51 | 1,131568577 | 0,765 | 0 | 1,12E-46 | MAC | Gpr65 |
| Apoc2 | 7,07E-51 | 1,087753959 | 0,765 | 0 | 1,12E-46 | MAC | Apoc2 |
| Cd86 | 7,07E-51 | 1,022165243 | 0,765 | 0 | 1,12E-46 | MAC | Cd86 |
| Vav1 | 7,07E-51 | 1,008406534 | 0,765 | 0 | 1,12E-46 | MAC | Vav1 |
| Itgb5 | 1,85E-50 | 1,33655926 | 0,79 | 0,012 | 2,94E-46 | MAC | Itgb5 |
| Clec4a3 | 6,31E-50 | 1,132583469 | 0,753 | 0 | 9,99E-46 | MAC | Clec4a3 |
| Bcl2a1d | 6,31E-50 | 0,96999313 | 0,753 | 0 | 9,99E-46 | MAC | Bcl2a1d |
| Hpgds | 9,53E-50 | 0,968983948 | 0,778 | 0,008 | 1,51E-45 | MAC | Hpgds |
| Ncf1 | 1,10E-49 | 1,076307045 | 0,778 | 0,008 | 1,75E-45 | MAC | Ncf1 |
| Gm2a | 1,11E-49 | 1,823513617 | 1 | 0,219 | 1,75E-45 | MAC | Gm2a |
| Ccl2 | 2,04E-49 | 1,96990662 | 0,864 | 0,041 | 3,23E-45 | MAC | Ccl2 |
| Ccl4 | 2,05E-49 | 2,292775707 | 0,765 | 0,008 | 3,25E-45 | MAC | Ccl4 |
| Cd48 | 2,05E-49 | 1,052723344 | 0,765 | 0,008 | 3,25E-45 | MAC | Cd48 |
| Bin2 | 2,83E-49 | 1,287063192 | 0,765 | 0,008 | 4,48E-45 | MAC | Bin2 |
| Cd72 | 4,89E-49 | 1,826717892 | 0,827 | 0,037 | 7,75E-45 | MAC | Cd72 |
| Plbd1 | 5,55E-49 | 1,632109157 | 0,741 | 0 | 8,80E-45 | MAC | Plbd1 |
| Fgd2 | 5,55E-49 | 1,318948008 | 0,741 | 0 | 8,80E-45 | MAC | Fgd2 |
| Aoah | 5,55E-49 | 0,989179868 | 0,741 | 0 | 8,80E-45 | MAC | Aoah |
| Apoe | 1,06E-48 | 5,76904748 | 1 | 0,281 | 1,67E-44 | MAC | Apoe |
| Klra2 | 4,82E-48 | 1,07273413 | 0,728 | 0 | 7,65E-44 | MAC | Klra2 |
| Prune2 | 5,85E-48 | 1,135055499 | 0,84 | 0,041 | 9,27E-44 | MAC | Prune2 |
| Irf5 | 1,08E-47 | 1,501959508 | 0,877 | 0,091 | 1,72E-43 | MAC | Irf5 |
| Ifi207 | 2,25E-47 | 1,035036117 | 0,753 | 0,012 | 3,56E-43 | MAC | Ifi207 |
| Bcl2a1a | 4,14E-47 | 1,005691984 | 0,716 | 0 | 6,56E-43 | MAC | Bcl2a1a |
| Tmem119 | 4,60E-47 | 1,483698869 | 0,728 | 0,004 | 7,29E-43 | MAC | Tmem119 |
| Arhgap45 | 3,50E-46 | 1,004643969 | 0,704 | 0 | 5,55E-42 | MAC | Arhgap45 |
| Tbxas1 | 4,84E-46 | 1,013880169 | 0,728 | 0,008 | 7,67E-42 | MAC | Tbxas1 |
| Sirpa | 6,66E-46 | 1,643392497 | 0,963 | 0,169 | 1,06E-41 | MAC | Sirpa |
| Ctsh | 9,56E-46 | 2,001194247 | 0,988 | 0,252 | 1,51E-41 | MAC | Ctsh |

| gene | p_val | avg_logFC | pct.1 | pct.2 | p_val_adj | Seurat_cluster | gene |
|---|---|---|---|---|---|---|---|
| Lrg1 | 2,39E-28 | 1,416100655 | 1 | 0,601 | 3,79E-24 | 0 | Lrg1 |
| Ly6c1 | 1,45E-27 | 1,251394325 | 0,958 | 0,513 | 2,31E-23 | 0 | Ly6c1 |
| Ly6a | 1,27E-24 | 1,158792599 | 0,989 | 0,693 | 2,02E-20 | 0 | Ly6a |
| Entpd1 | 2,29E-19 | 1,058261784 | 0,926 | 0,816 | 3,63E-15 | 0 | Entpd1 |
| Cyyr1 | 2,98E-19 | 0,932372667 | 0,905 | 0,518 | 4,73E-15 | 0 | Cyyr1 |
| Heg1 | 9,20E-18 | 0,842477285 | 0,968 | 0,575 | 1,46E-13 | 0 | Heg1 |
| Slco2a1 | 2,84E-17 | 1,040811755 | 0,832 | 0,43 | 4,51E-13 | 0 | Slco2a1 |
| Ramp2 | 8,90E-17 | 0,718415995 | 0,958 | 0,596 | 1,41E-12 | 0 | Ramp2 |
| Mgp | 1,78E-16 | 1,47038136 | 0,495 | 0,11 | 2,82E-12 | 0 | Mgp |
| Cd200 | 2,58E-16 | 0,779260323 | 1 | 0,649 | 4,09E-12 | 0 | Cd200 |
| Slc9a3r2 | 4,11E-16 | 0,896849171 | 0,884 | 0,518 | 6,52E-12 | 0 | Slc9a3r2 |
| Insr | 6,19E-16 | 0,942762923 | 0,863 | 0,509 | 9,80E-12 | 0 | Insr |
| Thbd | 9,70E-16 | 0,847426382 | 0,663 | 0,232 | 1,54E-11 | 0 | Thbd |
| Podxl | 7,84E-15 | 0,782148002 | 0,853 | 0,487 | 1,24E-10 | 0 | Podxl |
| Ptprb | 1,51E-14 | 0,780920999 | 0,905 | 0,557 | 2,39E-10 | 0 | Ptprb |
| Calcrl | 2,34E-14 | 0,752624296 | 0,811 | 0,43 | 3,71E-10 | 0 | Calcrl |
| Tsc22d1 | 3,33E-14 | 0,962864556 | 0,863 | 0,482 | 5,28E-10 | 0 | Tsc22d1 |
| Jam2 | 5,39E-14 | 0,938081989 | 0,663 | 0,25 | 8,55E-10 | 0 | Jam2 |
| Cxx1a | 7,79E-14 | 0,580798406 | 0,926 | 0,61 | 1,23E-09 | 0 | Cxx1a |
| Igfbp7 | 1,94E-13 | 0,726377765 | 1 | 0,671 | 3,08E-09 | 0 | Igfbp7 |
| Plpp3 | 3,05E-13 | 0,876499368 | 0,821 | 0,434 | 4,83E-09 | 0 | Plpp3 |
| Lims2 | 6,84E-13 | 0,795972978 | 0,505 | 0,136 | 1,08E-08 | 0 | Lims2 |
| Ptprg | 7,45E-13 | 0,6501828 | 0,695 | 0,285 | 1,18E-08 | 0 | Ptprg |
| Id1 | 9,76E-13 | 0,885006272 | 0,758 | 0,417 | 1,55E-08 | 0 | Id1 |
| Epas1 | 4,14E-12 | 0,855417855 | 0,895 | 0,544 | 6,57E-08 | 0 | Epas1 |
| Eng | 7,84E-12 | 0,600944625 | 0,979 | 0,732 | 1,24E-07 | 0 | Eng |
| Acer2 | 1,80E-11 | 0,81329729 | 0,621 | 0,254 | 2,85E-07 | 0 | Acer2 |
| Cd34 | 4,43E-11 | 0,518219717 | 0,979 | 0,649 | 7,02E-07 | 0 | Cd34 |
| Cbfa2t3 | 7,69E-11 | 0,439752616 | 0,547 | 0,193 | 1,22E-06 | 0 | Cbfa2t3 |
| Slc52a3 | 1,22E-10 | 0,622917456 | 0,411 | 0,105 | 1,94E-06 | 0 | Slc52a3 |
| Plpp1 | 1,34E-10 | 0,732451252 | 0,842 | 0,553 | 2,12E-06 | 0 | Plpp1 |
| Tmem252 | 1,39E-10 | 0,722735567 | 0,832 | 0,539 | 2,20E-06 | 0 | Tmem252 |
| Abcg2 | 1,59E-10 | 0,630096263 | 0,842 | 0,491 | 2,52E-06 | 0 | Abcg2 |
| Clec14a | 1,68E-10 | 0,766830841 | 0,768 | 0,465 | 2,67E-06 | 0 | Clec14a |
| Tmem88 | 1,75E-10 | 0,537650399 | 0,926 | 0,601 | 2,77E-06 | 0 | Tmem88 |
| Tek | 1,96E-10 | 0,658723593 | 0,758 | 0,417 | 3,10E-06 | 0 | Tek |
| Egfl7 | 2,55E-10 | 0,531155839 | 0,989 | 0,658 | 4,05E-06 | 0 | Egfl7 |
| Vegfc | 2,76E-10 | 0,574418063 | 0,337 | 0,07 | 4,38E-06 | 0 | Vegfc |
| Gimap6 | 2,94E-10 | 0,546793709 | 0,937 | 0,618 | 4,66E-06 | 0 | Gimap6 |
| Ctla2a | 3,62E-10 | 0,677275442 | 0,979 | 0,654 | 5,74E-06 | 0 | Ctla2a |
| Grrp1 | 4,09E-10 | 0,63973569 | 0,663 | 0,329 | 6,49E-06 | 0 | Grrp1 |
| Rbp7 | 4,22E-10 | 0,602118922 | 0,305 | 0,053 | 6,69E-06 | 0 | Rbp7 |
| Abcb1a | 5,89E-10 | 0,445443673 | 0,642 | 0,285 | 9,34E-06 | 0 | Abcb1a |
| Crip2 | 1,07E-09 | 0,49691827 | 0,947 | 0,636 | 1,70E-05 | 0 | Crip2 |
| Igfbp5 | 1,26E-09 | 1,163171383 | 0,547 | 0,246 | 1,99E-05 | 0 | Igfbp5 |
| Adamtsl1 | 1,52E-09 | 0,388618026 | 0,326 | 0,07 | 2,42E-05 | 0 | Adamtsl1 |
| Stox2 | 2,12E-09 | 0,289445255 | 0,305 | 0,057 | 3,36E-05 | 0 | Stox2 |
| Igfbp3 | 3,16E-09 | 0,653491481 | 0,979 | 0,697 | 5,01E-05 | 0 | Igfbp3 |
| Id3 | 5,06E-09 | 0,695386536 | 0,758 | 0,478 | 8,02E-05 | 0 | Id3 |
| Arf2 | 5,09E-09 | 0,866073277 | 0,695 | 0,443 | 8,07E-05 | 0 | Arf2 |
| Rab11a | 5,19E-09 | 0,503722226 | 0,895 | 0,833 | 8,22E-05 | 0 | Rab11a |
| Cpe | 5,22E-09 | 0,698095442 | 0,811 | 0,504 | 8,28E-05 | 0 | Cpe |
| mt-Cytb | 5,24E-09 | 0,352698583 | 1 | 1 | 8,31E-05 | 0 | mt-Cytb |
| Apold1 | 8,27E-09 | 0,666182905 | 0,758 | 0,474 | 0,000131115 | 0 | Apold1 |
| Zfp69 | 9,31E-09 | 0,274972036 | 0,432 | 0,136 | 0,000147559 | 0 | Zfp69 |
| Flt1 | 1,15E-08 | 0,49239355 | 0,947 | 0,636 | 0,000182266 | 0 | Flt1 |
| Mmrn2 | 1,26E-08 | 0,589861954 | 0,853 | 0,5 | 0,000200234 | 0 | Mmrn2 |
| Ace | 1,27E-08 | 0,42022655 | 0,432 | 0,149 | 0,000200996 | 0 | Ace |
| Hspb1 | 1,29E-08 | 0,479853097 | 0,505 | 0,202 | 0,000204183 | 0 | Hspb1 |
| Adamts8 | 1,65E-08 | 0,412855384 | 0,253 | 0,044 | 0,000261528 | 0 | Adamts8 |
| Dnm3 | 1,95E-08 | 0,37879572 | 0,379 | 0,118 | 0,000308546 | 0 | Dnm3 |
| Osmr | 2,22E-08 | 0,491255039 | 0,474 | 0,171 | 0,000351763 | 0 | Osmr |
| Atp1b3 | 2,74E-08 | 0,464932713 | 0,979 | 0,89 | 0,000434715 | 0 | Atp1b3 |
| S100a6 | 2,82E-08 | 0,416813727 | 0,958 | 0,768 | 0,000446453 | 0 | S100a6 |
| Gpr182 | 3,53E-08 | 0,516725211 | 0,379 | 0,118 | 0,0005601 | 0 | Gpr182 |
| Ecscr | 4,07E-08 | 0,406862384 | 0,947 | 0,64 | 0,00064519 | 0 | Ecscr |
| Nxpe4 | 4,23E-08 | 0,455781013 | 0,516 | 0,211 | 0,000671276 | 0 | Nxpe4 |
| Adgrg1 | 4,99E-08 | 0,380910274 | 0,726 | 0,355 | 0,000790708 | 0 | Adgrg1 |
| Pcp4l1 | 5,19E-08 | 0,622433446 | 0,305 | 0,079 | 0,000822503 | 0 | Pcp4l1 |
| Palm | 5,81E-08 | 0,500937604 | 0,579 | 0,289 | 0,000921043 | 0 | Palm |
| Itm2a | 6,72E-08 | 0,884598133 | 0,4 | 0,14 | 0,001064539 | 0 | Itm2a |
| Vwf | 7,66E-08 | 0,625687696 | 0,905 | 0,601 | 0,001214071 | 0 | Vwf |
| Lmcd1 | 8,95E-08 | 0,616126156 | 0,421 | 0,162 | 0,001419213 | 0 | Lmcd1 |
| Ceacam1 | 9,96E-08 | 0,531960289 | 0,421 | 0,167 | 0,001579559 | 0 | Ceacam1 |
| Rflnb | 9,99E-08 | 0,456448161 | 0,505 | 0,237 | 0,001583359 | 0 | Rflnb |
| Slc39a8 | 1,05E-07 | 0,668336518 | 0,463 | 0,202 | 0,001658448 | 0 | Slc39a8 |
| Tmem44 | 1,12E-07 | 0,399248912 | 0,347 | 0,105 | 0,001768676 | 0 | Tmem44 |
| Bmpr2 | 1,30E-07 | 0,499589226 | 0,768 | 0,522 | 0,002067562 | 0 | Bmpr2 |
| Selp | 1,71E-07 | 0,568148031 | 0,347 | 0,101 | 0,002712665 | 0 | Selp |
| Shroom4 | 1,74E-07 | 0,502442557 | 0,442 | 0,18 | 0,002750576 | 0 | Shroom4 |
| Pam | 1,75E-07 | 0,266610041 | 0,968 | 0,75 | 0,002778742 | 0 | Pam |
| Zfp423 | 1,90E-07 | 0,408675619 | 0,411 | 0,154 | 0,003004909 | 0 | Zfp423 |
| Arl4a | 2,00E-07 | 0,665638252 | 0,621 | 0,377 | 0,003162551 | 0 | Arl4a |
| Pkp4 | 2,30E-07 | 0,529103435 | 0,526 | 0,246 | 0,003641675 | 0 | Pkp4 |
| Gimap1 | 3,12E-07 | 0,531131344 | 0,632 | 0,36 | 0,004948199 | 0 | Gimap1 |
| Kbtbd11 | 3,15E-07 | 0,357137915 | 0,453 | 0,18 | 0,004994701 | 0 | Kbtbd11 |
| Gimap5 | 3,70E-07 | 0,382157916 | 0,695 | 0,404 | 0,005859218 | 0 | Gimap5 |
| Emcn | 3,83E-07 | 0,487478466 | 0,958 | 0,623 | 0,006071546 | 0 | Emcn |
| Cxx1b | 4,78E-07 | 0,268633959 | 0,874 | 0,596 | 0,007582077 | 0 | Cxx1b |
| Ushbp1 | 5,23E-07 | 0,598490285 | 0,737 | 0,474 | 0,008291494 | 0 | Ushbp1 |
| Lysmd2 | 5,25E-07 | 0,56483513 | 0,558 | 0,298 | 0,008327674 | 0 | Lysmd2 |
| Scgb3a1 | 5,27E-07 | 1,317938787 | 0,442 | 0,197 | 0,008349442 | 0 | Scgb3a1 |
| Tacr1 | 6,91E-07 | 0,629142365 | 0,4 | 0,167 | 0,010947797 | 0 | Tacr1 |
| Ndufa8 | 7,20E-07 | 0,432544009 | 0,884 | 0,746 | 0,011407958 | 0 | Ndufa8 |
| Bvht | 9,48E-07 | 0,369959593 | 0,642 | 0,373 | 0,015032475 | 0 | Bvht |
| Prex2 | 9,74E-07 | 0,445388475 | 0,884 | 0,803 | 0,015440189 | 0 | Prex2 |
| Tns2 | 1,06E-06 | 0,488894367 | 0,558 | 0,307 | 0,016748943 | 0 | Tns2 |
| Pltp | 1,07E-06 | 0,819174384 | 0,526 | 0,294 | 0,017023498 | 0 | Pltp |
| Tmsb10 | 1,19E-06 | 0,33919803 | 1 | 0,965 | 0,018866025 | 0 | Tmsb10 |
| Ehd3 | 1,20E-06 | 0,502661058 | 0,579 | 0,289 | 0,019003373 | 0 | Ehd3 |
| Rassf9 | 1,27E-06 | 0,416063088 | 0,316 | 0,101 | 0,020168711 | 0 | Rassf9 |
| Palmd | 1,36E-06 | 0,585677214 | 0,621 | 0,373 | 0,02151921 | 0 | Palmd |
| Sptbn1 | 1,62E-06 | 0,445990112 | 0,947 | 0,746 | 0,025749919 | 0 | Sptbn1 |
| Ece1 | 1,64E-06 | 0,304211211 | 1 | 0,671 | 0,026022477 | 0 | Ece1 |
| Esam | 1,66E-06 | 0,438487092 | 0,947 | 0,645 | 0,026325541 | 0 | Esam |
| Cldn5 | 1,66E-06 | 0,467605959 | 0,926 | 0,623 | 0,026369331 | 0 | Cldn5 |
| Plvap | 2,22E-06 | 0,433753709 | 0,905 | 0,596 | 0,0351228 | 0 | Plvap |
| Ets1 | 2,46E-06 | 0,404814363 | 0,905 | 0,662 | 0,038951957 | 0 | Ets1 |
| Zfp36l1 | 2,50E-06 | 0,379525952 | 0,958 | 0,965 | 0,039599783 | 0 | Zfp36l1 |
| Kdr | 2,50E-06 | 0,407372253 | 0,895 | 0,566 | 0,039660314 | 0 | Kdr |
| Atox1 | 2,65E-06 | 0,282272393 | 0,947 | 0,982 | 0,041964399 | 0 | Atox1 |
| Foxp1 | 2,67E-06 | 0,553279915 | 0,737 | 0,57 | 0,042372669 | 0 | Foxp1 |
| Ly6e | 2,86E-06 | 0,365011539 | 1 | 0,987 | 0,045318955 | 0 | Ly6e |
| Nxn | 2,93E-06 | 0,399310141 | 0,421 | 0,189 | 0,046439249 | 0 | Nxn |
| Efr3b | 2,97E-06 | 0,514838422 | 0,263 | 0,075 | 0,04708298 | 0 | Efr3b |

| | | | | | | | |
|---|---|---|---|---|---|---|---|
| Dpep2 | 1,42E-45 | 1,043355863 | 0,716 | 0,008 | 2,26E-41 | MAC | Dpep2 |
| Pilra | 2,15E-45 | 1,017843766 | 0,704 | 0,004 | 3,40E-41 | MAC | Pilra |
| Clec4a1 | 2,93E-45 | 1,224646934 | 0,691 | 0 | 4,64E-41 | MAC | Clec4a1 |
| Pid1 | 2,93E-45 | 1,043526642 | 0,691 | 0 | 4,64E-41 | MAC | Pid1 |
| Tifab | 2,93E-45 | 0,893903155 | 0,691 | 0 | 4,64E-41 | MAC | Tifab |
| Pirb | 3,32E-45 | 1,387346463 | 0,741 | 0,021 | 5,26E-41 | MAC | Pirb |
| Lst1 | 4,33E-45 | 0,643469085 | 0,728 | 0,017 | 6,86E-41 | MAC | Lst1 |
| Lgals3 | 8,65E-45 | 2,203308785 | 0,864 | 0,099 | 1,37E-40 | MAC | Lgals3 |
| Pf4 | 1,58E-44 | 1,235653998 | 0,753 | 0,029 | 2,50E-40 | MAC | Pf4 |
| Cd14 | 2,50E-44 | 1,973991269 | 0,938 | 0,145 | 3,97E-40 | MAC | Cd14 |
| Zmynd15 | 1,06E-43 | 1,512281879 | 0,704 | 0,012 | 1,68E-39 | MAC | Zmynd15 |
| Pycard | 1,10E-43 | 1,183325971 | 0,864 | 0,087 | 1,74E-39 | MAC | Pycard |
| AU020206 | 1,51E-43 | 1,118438958 | 0,951 | 0,178 | 2,39E-39 | MAC | AU020206 |
| Efhd2 | 1,74E-43 | 1,550336541 | 0,963 | 0,219 | 2,75E-39 | MAC | Efhd2 |
| Sirpb1b | 1,97E-43 | 0,458000508 | 0,667 | 0 | 3,12E-39 | MAC | Sirpb1b |
| Rgs2 | 2,44E-43 | 1,650651099 | 0,951 | 0,161 | 3,87E-39 | MAC | Rgs2 |
| Tlr2 | 4,10E-43 | 1,300151531 | 0,765 | 0,037 | 6,50E-39 | MAC | Tlr2 |
| Unc93b1 | 5,60E-43 | 1,83824491 | 1 | 0,351 | 8,88E-39 | MAC | Unc93b1 |
| H2-Aa | 6,01E-43 | 2,966786186 | 0,827 | 0,083 | 9,53E-39 | MAC | H2-Aa |
| P2ry6 | 7,69E-43 | 1,356527043 | 0,827 | 0,07 | 1,22E-38 | MAC | P2ry6 |
| Gatm | 2,41E-42 | 1,559059904 | 0,926 | 0,157 | 3,82E-38 | MAC | Gatm |
| Ctsz | 2,54E-42 | 2,09688997 | 1 | 0,413 | 4,02E-38 | MAC | Ctsz |
| Ccl7 | 4,74E-42 | 2,092756852 | 0,679 | 0,012 | 7,52E-38 | MAC | Ccl7 |
| Cd83 | 1,26E-41 | 1,366010777 | 0,642 | 0 | 1,99E-37 | MAC | Cd83 |
| Itgal | 1,26E-41 | 0,788248745 | 0,642 | 0 | 1,99E-37 | MAC | Itgal |
| Ucp2 | 1,35E-41 | 2,064485573 | 1 | 0,405 | 2,14E-37 | MAC | Ucp2 |
| Ntpcr | 3,70E-41 | 1,495518791 | 0,901 | 0,161 | 5,86E-37 | MAC | Ntpcr |
| Cst3 | 4,35E-41 | 1,849523353 | 1 | 0,851 | 6,89E-37 | MAC | Cst3 |
| Zeb2 | 6,25E-41 | 0,941150684 | 0,741 | 0,033 | 9,91E-37 | MAC | Zeb2 |
| Ccl9 | 8,03E-41 | 1,141040972 | 0,642 | 0,004 | 1,27E-36 | MAC | Ccl9 |
| Myo1f | 9,85E-41 | 0,881048734 | 0,63 | 0 | 1,56E-36 | MAC | Myo1f |
| Rbm47 | 9,89E-41 | 0,388600047 | 0,704 | 0,025 | 1,57E-36 | MAC | Rbm47 |
| Hexb | 1,01E-40 | 1,958017343 | 0,988 | 0,298 | 1,60E-36 | MAC | Hexb |
| Grn | 1,61E-40 | 1,850319544 | 1 | 0,773 | 2,55E-36 | MAC | Grn |
| Gm42031 | 2,09E-40 | 0,618574808 | 0,914 | 0,198 | 3,31E-36 | MAC | Gm42031 |
| Gm6377 | 2,52E-40 | 1,042744529 | 0,642 | 0,004 | 3,99E-36 | MAC | Gm6377 |
| Celf2 | 2,62E-40 | 0,88320141 | 0,654 | 0,008 | 4,15E-36 | MAC | Celf2 |
| Ncf2 | 3,99E-40 | 0,968890726 | 0,691 | 0,021 | 6,32E-36 | MAC | Ncf2 |
| Fth1 | 4,75E-40 | 1,914775366 | 1 | 0,988 | 7,54E-36 | MAC | Fth1 |
| Tnfaip8l2 | 7,36E-40 | 0,958776097 | 0,63 | 0,004 | 1,17E-35 | MAC | Tnfaip8l2 |
| Ftl1 | 7,51E-40 | 1,614515569 | 1 | 0,992 | 1,19E-35 | MAC | Ftl1 |
| Spint1 | 7,63E-40 | 0,95144666 | 0,617 | 0 | 1,21E-35 | MAC | Spint1 |
| Il10ra | 7,63E-40 | 0,902622275 | 0,617 | 0 | 1,21E-35 | MAC | Il10ra |
| Lrrc25 | 7,63E-40 | 0,673458635 | 0,617 | 0 | 1,21E-35 | MAC | Lrrc25 |
| Emb | 7,95E-40 | 1,216127566 | 0,642 | 0,008 | 1,26E-35 | MAC | Emb |
| Cd74 | 1,34E-39 | 3,250433361 | 0,951 | 0,24 | 2,12E-35 | MAC | Cd74 |
| Trf | 2,10E-39 | 2,137320354 | 0,963 | 0,326 | 3,32E-35 | MAC | Trf |
| Ppfia4 | 5,01E-39 | 0,953528514 | 0,617 | 0,004 | 7,94E-35 | MAC | Ppfia4 |
| Bhlhe41 | 5,75E-39 | 0,793743502 | 0,667 | 0,017 | 9,12E-35 | MAC | Bhlhe41 |
| Arhgap30 | 5,84E-39 | 0,740906224 | 0,605 | 0 | 9,26E-35 | MAC | Arhgap30 |
| Lat2 | 7,18E-39 | 1,045474674 | 0,728 | 0,045 | 1,14E-34 | MAC | Lat2 |
| Cfp | 8,85E-39 | 0,997574157 | 0,617 | 0,004 | 1,40E-34 | MAC | Cfp |
| Prkcd | 1,40E-38 | 1,198810565 | 0,765 | 0,066 | 2,21E-34 | MAC | Prkcd |
| Lgmn | 1,49E-38 | 1,686533885 | 1 | 0,897 | 2,35E-34 | MAC | Lgmn |
| Rab3il1 | 2,08E-38 | 1,270197707 | 0,802 | 0,083 | 3,30E-34 | MAC | Rab3il1 |
| Cebpb | 2,44E-38 | 1,020953092 | 0,802 | 0,083 | 3,88E-34 | MAC | Cebpb |
| Man2b1 | 2,71E-38 | 1,68834718 | 0,988 | 0,401 | 4,29E-34 | MAC | Man2b1 |
| Dock10 | 3,89E-38 | 0,670253041 | 0,605 | 0,004 | 6,16E-34 | MAC | Dock10 |
| Hk2 | 3,89E-38 | 0,537198907 | 0,605 | 0,004 | 6,16E-34 | MAC | Hk2 |
| Ifi27l2a | 4,05E-38 | 0,863232196 | 0,617 | 0,008 | 6,42E-34 | MAC | Ifi27l2a |
| Olfml3 | 4,07E-38 | 0,944676953 | 0,642 | 0,017 | 6,46E-34 | MAC | Olfml3 |
| Cyba | 6,51E-38 | 1,447496936 | 1 | 0,736 | 1,03E-33 | MAC | Cyba |
| Cebpa | 6,91E-38 | 0,678538118 | 0,63 | 0,012 | 1,10E-33 | MAC | Cebpa |
| Cep85 | 7,38E-38 | 1,125955149 | 0,951 | 0,223 | 1,17E-33 | MAC | Cep85 |
| H2-Eb1 | 7,56E-38 | 2,820339309 | 0,778 | 0,091 | 1,20E-33 | MAC | H2-Eb1 |
| Psap | 9,09E-38 | 1,816068891 | 1 | 0,74 | 1,44E-33 | MAC | Psap |
| Tgfbi | 1,25E-37 | 1,886260686 | 0,975 | 0,285 | 1,98E-33 | MAC | Tgfbi |
| Sdc4 | 1,61E-37 | 1,174851714 | 0,877 | 0,136 | 2,55E-33 | MAC | Sdc4 |
| Ccl12 | 3,29E-37 | 1,327348185 | 0,58 | 0 | 5,22E-33 | MAC | Ccl12 |
| Tlr7 | 3,29E-37 | 0,571550199 | 0,58 | 0 | 5,22E-33 | MAC | Tlr7 |
| Sirpb1a | 3,29E-37 | 0,483691495 | 0,58 | 0 | 5,22E-33 | MAC | Sirpb1a |
| Lpxn | 4,54E-37 | 0,945628138 | 0,593 | 0,004 | 7,20E-33 | MAC | Lpxn |
| Arhgdib | 1,06E-36 | 1,5222498 | 0,951 | 0,322 | 1,69E-32 | MAC | Arhgdib |
| Maf | 1,49E-36 | 0,856992054 | 0,852 | 0,128 | 2,37E-32 | MAC | Maf |
| Adcy7 | 1,70E-36 | 0,839789888 | 0,617 | 0,017 | 2,69E-32 | MAC | Adcy7 |
| Ftl1-ps1 | 1,77E-36 | 0,797047456 | 0,988 | 0,913 | 2,81E-32 | MAC | Ftl1-ps1 |
| Clec7a | 2,25E-36 | 0,963189716 | 0,58 | 0,004 | 3,57E-32 | MAC | Clec7a |
| Rab7b | 2,38E-36 | 0,953264266 | 0,58 | 0,004 | 3,77E-32 | MAC | Rab7b |
| Gsn | 5,04E-36 | 1,154446012 | 0,765 | 0,083 | 7,99E-32 | MAC | Gsn |
| Slc7a8 | 5,85E-36 | 0,671393786 | 0,58 | 0,004 | 9,27E-32 | MAC | Slc7a8 |
| Vsir | 8,31E-36 | 1,17055103 | 0,901 | 0,182 | 1,32E-31 | MAC | Vsir |
| Mafb | 8,90E-36 | 1,590650052 | 0,877 | 0,198 | 1,41E-31 | MAC | Mafb |
| Slc15a3 | 1,27E-35 | 0,760427349 | 0,741 | 0,062 | 2,02E-31 | MAC | Slc15a3 |
| Stxbp2 | 1,54E-35 | 0,791656462 | 0,605 | 0,017 | 2,44E-31 | MAC | Stxbp2 |
| Gm43549 | 1,67E-35 | 0,377947293 | 0,605 | 0,017 | 2,65E-31 | MAC | Gm43549 |
| Themis2 | 1,73E-35 | 0,640242425 | 0,568 | 0,004 | 2,74E-31 | MAC | Themis2 |
| Evi2b | 3,66E-35 | 0,626299369 | 0,593 | 0,012 | 5,81E-31 | MAC | Evi2b |
| Gm43305 | 6,48E-35 | 0,828603696 | 0,988 | 0,368 | 1,03E-30 | MAC | Gm43305 |
| Tmem106a | 1,08E-34 | 1,065465683 | 0,778 | 0,103 | 1,72E-30 | MAC | Tmem106a |
| Adap2 | 1,27E-34 | 0,421789164 | 0,543 | 0 | 2,02E-30 | MAC | Adap2 |
| Parvg | 1,43E-34 | 0,797120824 | 0,556 | 0,004 | 2,26E-30 | MAC | Parvg |
| Dock2 | 1,47E-34 | 0,745931628 | 0,556 | 0,004 | 2,33E-30 | MAC | Dock2 |
| Ctsb | 1,63E-34 | 1,277189877 | 1 | 0,975 | 2,59E-30 | MAC | Ctsb |
| Rab32 | 2,55E-34 | 1,038452288 | 0,704 | 0,066 | 4,04E-30 | MAC | Rab32 |
| Npc2 | 2,66E-34 | 1,352299046 | 1 | 0,636 | 4,21E-30 | MAC | Npc2 |
| Prdx5 | 2,91E-34 | 1,374611256 | 0,988 | 0,607 | 4,62E-30 | MAC | Prdx5 |
| Ptprj | 3,68E-34 | 0,500222645 | 0,568 | 0,008 | 5,84E-30 | MAC | Ptprj |
| Pkib | 9,07E-34 | 0,616131471 | 0,531 | 0 | 1,44E-29 | MAC | Pkib |
| Igsf6 | 9,71E-34 | 0,887651126 | 0,556 | 0,008 | 1,54E-29 | MAC | Igsf6 |
| AB124611 | 1,10E-33 | 0,690231993 | 0,543 | 0,004 | 1,74E-29 | MAC | AB124611 |
| Snx5 | 1,11E-33 | 1,439090225 | 0,963 | 0,401 | 1,75E-29 | MAC | Snx5 |
| Ctsa | 2,05E-33 | 1,413419045 | 1 | 0,682 | 3,25E-29 | MAC | Ctsa |
| Lgals1 | 3,49E-33 | 1,223815921 | 0,988 | 0,426 | 5,54E-29 | MAC | Lgals1 |
| Ctsd | 3,81E-33 | 1,766775887 | 1 | 0,682 | 6,04E-29 | MAC | Ctsd |
| Zfp36 | 5,17E-33 | 1,747150546 | 0,975 | 0,521 | 8,19E-29 | MAC | Zfp36 |
| Mertk | 5,99E-33 | 0,832936549 | 0,642 | 0,041 | 9,50E-29 | MAC | Mertk |
| P2ry12 | 6,52E-33 | 0,808904511 | 0,531 | 0,004 | 1,03E-28 | MAC | P2ry12 |
| Sema4b | 6,64E-33 | 0,288141104 | 0,679 | 0,05 | 1,05E-28 | MAC | Sema4b |
| Ikzf1 | 7,49E-33 | 0,751378546 | 0,531 | 0,004 | 1,19E-28 | MAC | Ikzf1 |
| H2-DMb1 | 8,88E-33 | 1,266057588 | 0,79 | 0,12 | 1,41E-28 | MAC | H2-DMb1 |
| Cd44 | 9,88E-33 | 0,627273201 | 0,531 | 0,004 | 1,57E-28 | MAC | Cd44 |
| Rnase4 | 1,84E-32 | 1,37399573 | 0,914 | 0,248 | 2,92E-28 | MAC | Rnase4 |
| Runx1 | 3,16E-32 | 0,700503396 | 0,58 | 0,025 | 5,01E-28 | MAC | Runx1 |
| Sema4d | 4,44E-32 | 0,58492625 | 0,506 | 0 | 7,03E-28 | MAC | Sema4d |
| Fgl2 | 4,64E-32 | 0,879039726 | 0,519 | 0,004 | 7,36E-28 | MAC | Fgl2 |
| F13a1 | 4,77E-32 | 1,363365067 | 0,519 | 0,004 | 7,57E-28 | MAC | F13a1 |
| Naaa | 4,91E-32 | 0,736130496 | 0,519 | 0,004 | 7,78E-28 | MAC | Naaa |
| Sh2d1b1 | 5,05E-32 | 0,718340252 | 0,519 | 0,004 | 8,00E-28 | MAC | Sh2d1b1 |
| Arl11 | 6,16E-32 | 0,844862586 | 0,593 | 0,029 | 9,76E-28 | MAC | Arl11 |
| Atp13a2 | 8,76E-32 | 0,949076139 | 0,728 | 0,099 | 1,39E-27 | MAC | Atp13a2 |

| | | | | | | | |
|---|---|---|---|---|---|---|---|
| Tm4sf1 | 3,06E-06 | 0,423926489 | 0,958 | 0,632 | 0,048526806 | 0 | Tm4sf1 |
| Sorbs3 | 3,11E-06 | 0,376208232 | 0,832 | 0,627 | 0,049249966 | 0 | Sorbs3 |
| Elk3 | 3,54E-06 | 0,380131109 | 0,937 | 0,772 | 0,056143765 | 0 | Elk3 |
| Timp3 | 4,02E-06 | 0,489366934 | 0,821 | 0,579 | 0,063680617 | 0 | Timp3 |
| Dock6 | 4,09E-06 | 0,415448544 | 0,6 | 0,338 | 0,06476602 | 0 | Dock6 |
| Cmtm8 | 4,19E-06 | 0,319628339 | 0,453 | 0,215 | 0,066437203 | 0 | Cmtm8 |
| Clu | 4,35E-06 | 0,716242929 | 0,421 | 0,189 | 0,068940008 | 0 | Clu |
| Lamb2 | 4,36E-06 | 0,335902195 | 0,358 | 0,136 | 0,069040934 | 0 | Lamb2 |
| Ifitm3 | 4,67E-06 | 0,357121638 | 0,979 | 0,939 | 0,074066168 | 0 | Ifitm3 |
| Mkl2 | 4,68E-06 | 0,585214234 | 0,768 | 0,522 | 0,074257263 | 0 | Mkl2 |
| Cd93 | 4,75E-06 | 0,322084985 | 1 | 0,899 | 0,075270601 | 0 | Cd93 |
| Cmip | 4,87E-06 | 0,496277663 | 0,716 | 0,513 | 0,077214529 | 0 | Cmip |
| Ecm1 | 5,09E-06 | 0,505516163 | 0,884 | 0,759 | 0,080728782 | 0 | Ecm1 |
| Fry | 5,74E-06 | 0,252052617 | 0,632 | 0,373 | 0,091061283 | 0 | Fry |
| Pmp22 | 7,89E-06 | 0,467662952 | 0,4 | 0,189 | 0,125071149 | 0 | Pmp22 |
| Nostrin | 8,02E-06 | 0,414634252 | 0,695 | 0,504 | 0,127139951 | 0 | Nostrin |
| Vamp5 | 8,33E-06 | 0,337434073 | 0,695 | 0,439 | 0,13197018 | 0 | Vamp5 |
| Plec | 1,06E-05 | 0,278941939 | 0,937 | 0,715 | 0,167917629 | 0 | Plec |
| Ppic | 1,10E-05 | 0,378500811 | 0,916 | 0,64 | 0,174491229 | 0 | Ppic |
| Pitpnc1 | 1,13E-05 | 0,412136503 | 0,758 | 0,548 | 0,178600613 | 0 | Pitpnc1 |
| Ephb4 | 1,38E-05 | 0,334243703 | 0,642 | 0,368 | 0,218639944 | 0 | Ephb4 |
| Ripply3 | 1,45E-05 | 0,316997609 | 0,274 | 0,096 | 0,230477342 | 0 | Ripply3 |
| Col4a2 | 1,54E-05 | 0,311179108 | 1 | 0,671 | 0,243552916 | 0 | Col4a2 |
| Tspan12 | 1,55E-05 | 0,582395496 | 0,579 | 0,373 | 0,245228929 | 0 | Tspan12 |
| Oaz2 | 1,62E-05 | 0,565191502 | 0,737 | 0,588 | 0,257295791 | 0 | Oaz2 |
| Armcx2 | 1,76E-05 | 0,534801247 | 0,432 | 0,224 | 0,279456487 | 0 | Armcx2 |
| Efnb2 | 1,77E-05 | 0,510764693 | 0,4 | 0,202 | 0,280791733 | 0 | Efnb2 |
| Tie1 | 1,82E-05 | 0,289525955 | 0,832 | 0,557 | 0,288938053 | 0 | Tie1 |
| Sept4 | 1,87E-05 | 0,404388288 | 0,611 | 0,368 | 0,29564392 | 0 | Septin4 |
| Pkn3 | 1,89E-05 | 0,5971469 | 0,537 | 0,298 | 0,299120303 | 0 | Pkn3 |
| Sdpr | 1,92E-05 | 0,469272005 | 0,442 | 0,215 | 0,30465035 | 0 | Sdpr |
| Synm | 1,98E-05 | 0,330196986 | 0,316 | 0,118 | 0,314502821 | 0 | Synm |
| Capn5 | 2,02E-05 | 0,303420351 | 0,263 | 0,088 | 0,320172242 | 0 | Capn5 |
| Selenom | 2,05E-05 | 0,341010231 | 0,674 | 0,469 | 0,324774099 | 0 | Selenom |
| Car2 | 2,30E-05 | 0,620706211 | 0,358 | 0,171 | 0,364645167 | 0 | Car2 |
| Efna1 | 2,43E-05 | 0,425887766 | 0,832 | 0,57 | 0,384860062 | 0 | Efna1 |
| Cnn3 | 2,43E-05 | 0,35660785 | 0,937 | 0,654 | 0,385334081 | 0 | Cnn3 |
| Npnt | 2,49E-05 | 0,435316653 | 0,295 | 0,11 | 0,395137983 | 0 | Npnt |
| Cers4 | 2,55E-05 | 0,429726399 | 0,379 | 0,171 | 0,404664769 | 0 | Cers4 |
| Adamts9 | 2,63E-05 | 0,397006058 | 0,642 | 0,399 | 0,416642209 | 0 | Adamts9 |
| BC028528 | 2,76E-05 | 0,357808892 | 0,874 | 0,798 | 0,438263344 | 0 | BC028528 |
| Dock9 | 2,89E-05 | 0,359099837 | 0,768 | 0,522 | 0,458352885 | 0 | Dock9 |
| Cd300lg | 2,93E-05 | 0,673112519 | 0,421 | 0,215 | 0,463964358 | 0 | Cd300lg |
| Dok4 | 3,20E-05 | 0,443638775 | 0,411 | 0,202 | 0,506766395 | 0 | Dok4 |
| Mcf2l | 3,28E-05 | 0,546563238 | 0,411 | 0,206 | 0,51917239 | 0 | Mcf2l |
| Adam15 | 3,31E-05 | 0,478913211 | 0,884 | 0,785 | 0,525430334 | 0 | Adam15 |
| Cd59a | 3,42E-05 | 0,447029228 | 0,337 | 0,145 | 0,541896401 | 0 | Cd59a |
| Abcc4 | 3,49E-05 | 0,449078088 | 0,579 | 0,368 | 0,553805579 | 0 | Abcc4 |
| Mxra8 | 3,52E-05 | 0,512989678 | 0,579 | 0,351 | 0,558485899 | 0 | Mxra8 |
| Ppp1r2 | 3,58E-05 | 0,515846889 | 0,747 | 0,623 | 0,567949391 | 0 | Ppp1r2 |
| Nos3 | 3,60E-05 | 0,382745048 | 0,768 | 0,522 | 0,570643157 | 0 | Nos3 |
| Tmem204 | 3,68E-05 | 0,400097825 | 0,705 | 0,487 | 0,584071747 | 0 | Tmem204 |
| Ltbp4 | 3,69E-05 | 0,497571124 | 0,516 | 0,294 | 0,58475958 | 0 | Ltbp4 |
| Npdc1 | 4,12E-05 | 0,425563835 | 0,653 | 0,443 | 0,653723458 | 0 | Npdc1 |
| Stom | 4,14E-05 | 0,452442513 | 0,516 | 0,307 | 0,656450509 | 0 | Stom |
| Exoc3l2 | 4,33E-05 | 0,511188616 | 0,432 | 0,228 | 0,686968627 | 0 | Exoc3l2 |
| Ankrd33b | 4,54E-05 | 0,310264086 | 0,284 | 0,11 | 0,719400005 | 0 | Ankrd33b |
| Sox18 | 4,57E-05 | 0,347022367 | 0,895 | 0,557 | 0,724867487 | 0 | Sox18 |
| AU021092 | 4,66E-05 | 0,304350576 | 0,421 | 0,206 | 0,738909512 | 0 | AU021092 |
| Mgea5 | 4,70E-05 | 0,348332813 | 0,516 | 0,303 | 0,744311623 | 0 | Mgea5 |
| Mprip | 4,87E-05 | 0,597674461 | 0,768 | 0,561 | 0,771295034 | 0 | Mprip |
| Rasip1 | 5,08E-05 | 0,282600384 | 0,821 | 0,504 | 0,804981779 | 0 | Rasip1 |
| Maoa | 5,10E-05 | 0,391178224 | 0,389 | 0,189 | 0,808389576 | 0 | Maoa |
| Adamts1 | 5,39E-05 | 0,448411341 | 0,726 | 0,504 | 0,854072195 | 0 | Adamts1 |
| Hmgn3 | 5,56E-05 | 0,461600665 | 0,484 | 0,294 | 0,88105416 | 0 | Hmgn3 |
| Icam2 | 6,02E-05 | 0,318514296 | 0,821 | 0,575 | 0,953958269 | 0 | Icam2 |
| Slc44a2 | 6,36E-05 | 0,305272002 | 0,579 | 0,329 | 1 | 0 | Slc44a2 |
| Tspan7 | 6,90E-05 | 0,446937778 | 0,495 | 0,268 | 1 | 0 | Tspan7 |
| Tshz2 | 7,06E-05 | 0,357668859 | 0,611 | 0,377 | 1 | 0 | Tshz2 |
| Lama5 | 7,20E-05 | 0,530093272 | 0,558 | 0,355 | 1 | 0 | Lama5 |
| Map1lc3b | 7,28E-05 | 0,367855802 | 0,947 | 0,895 | 1 | 0 | Map1lc3b |
| Gnb1 | 7,53E-05 | 0,298344015 | 1 | 0,996 | 1 | 0 | Gnb1 |
| Gimap4 | 7,92E-05 | 0,394935499 | 0,768 | 0,548 | 1 | 0 | Gimap4 |
| Igfbp4 | 7,98E-05 | 0,406016852 | 0,853 | 0,754 | 1 | 0 | Igfbp4 |
| Arhgef15 | 8,48E-05 | 0,28714017 | 0,726 | 0,526 | 1 | 0 | Arhgef15 |
| Mall | 8,85E-05 | 0,309282259 | 0,495 | 0,298 | 1 | 0 | Mall |
| Itm2b | 8,87E-05 | 0,605536584 | 1 | 0,996 | 1 | 0 | Itm2b |
| Plcb1 | 9,22E-05 | 0,421432759 | 0,358 | 0,167 | 1 | 0 | Plcb1 |
| Cyb5r3 | 9,32E-05 | 0,372176219 | 0,821 | 0,689 | 1 | 0 | Cyb5r3 |
| Pdlim1 | 9,66E-05 | 0,313381306 | 0,884 | 0,632 | 1 | 0 | Pdlim1 |
| S100a13 | 0,00010123 | 0,277970285 | 0,821 | 0,732 | 1 | 0 | S100a13 |
| Limch1 | 0,000109337 | 0,380496399 | 0,442 | 0,241 | 1 | 0 | Limch1 |
| B4galt4 | 0,00011678 | 0,355696205 | 0,347 | 0,162 | 1 | 0 | B4galt4 |
| Rpl35 | 0,000117947 | 0,250603025 | 0,989 | 0,991 | 1 | 0 | Rpl35 |
| Prss23 | 0,000118957 | 0,277632735 | 0,779 | 0,553 | 1 | 0 | Prss23 |
| Ifi44 | 0,000123535 | 0,328348945 | 0,305 | 0,127 | 1 | 0 | Ifi44 |
| Upp1 | 0,00013194 | 0,313553431 | 0,789 | 0,561 | 1 | 0 | Upp1 |
| Col18a1 | 0,000133942 | 0,348489317 | 0,737 | 0,535 | 1 | 0 | Col18a1 |
| Ccnyl1 | 0,000135864 | 0,260073967 | 0,316 | 0,14 | 1 | 0 | Ccnyl1 |
| Aplnr | 0,000138452 | 0,462101053 | 0,705 | 0,504 | 1 | 0 | Aplnr |
| Polr2b | 0,000149036 | 0,303837402 | 0,737 | 0,504 | 1 | 0 | Polr2b |
| S1pr1 | 0,000149574 | 0,267782594 | 0,979 | 0,68 | 1 | 0 | S1pr1 |
| Scarf1 | 0,000155679 | 0,353588691 | 0,558 | 0,342 | 1 | 0 | Scarf1 |
| Cltb | 0,000156107 | 0,350862352 | 0,684 | 0,5 | 1 | 0 | Cltb |
| Adcy4 | 0,000162452 | 0,423113128 | 0,537 | 0,316 | 1 | 0 | Adcy4 |
| Ahnak | 0,000163896 | 0,320056225 | 0,821 | 0,64 | 1 | 0 | Ahnak |
| Lsr | 0,000167269 | 0,454224053 | 0,389 | 0,211 | 1 | 0 | Lsr |
| Tmod3 | 0,000183079 | 0,354483278 | 0,853 | 0,754 | 1 | 0 | Tmod3 |
| Tspan9 | 0,00019122 | 0,28882556 | 0,737 | 0,491 | 1 | 0 | Tspan9 |
| Jup | 0,00020115 | 0,339530259 | 0,884 | 0,671 | 1 | 0 | Jup |
| Rai14 | 0,00021377 | 0,323965392 | 0,674 | 0,465 | 1 | 0 | Rai14 |
| Socs5 | 0,000215422 | 0,360943542 | 0,421 | 0,224 | 1 | 0 | Socs5 |
| Afap1l1 | 0,00022132 | 0,345368358 | 0,653 | 0,452 | 1 | 0 | Afap1l1 |
| Nr2f2 | 0,00023922 | 0,286228094 | 0,526 | 0,316 | 1 | 0 | Nr2f2 |
| Stc1 | 0,00024372 | 0,459974452 | 0,516 | 0,316 | 1 | 0 | Stc1 |
| Ccnd3 | 0,000286157 | 0,331363419 | 0,726 | 0,561 | 1 | 0 | Ccnd3 |
| Mn1 | 0,00028647 | 0,39828985 | 0,295 | 0,136 | 1 | 0 | Mn1 |
| 4930578C19Rik | 0,000290691 | 0,358886431 | 0,453 | 0,259 | 1 | 0 | 4930578C19Rik |
| Gatsl3 | 0,000291191 | 0,359784544 | 0,263 | 0,114 | 1 | 0 | Gatsl3 |
| Ccm2l | 0,000293189 | 0,303050038 | 0,379 | 0,197 | 1 | 0 | Ccm2l |
| Gm37376 | 0,000322314 | 0,450999995 | 0,979 | 0,991 | 1 | 0 | Gm37376 |
| Alad | 0,000339967 | 0,388011005 | 0,6 | 0,434 | 1 | 0 | Alad |
| Acvrl1 | 0,00039222 | 0,448514473 | 0,695 | 0,57 | 1 | 0 | Acvrl1 |
| Fam167b | 0,000400015 | 0,450660557 | 0,663 | 0,509 | 1 | 0 | Fam167b |
| App | 0,000408916 | 0,337804175 | 0,937 | 0,943 | 1 | 0 | App |
| Ick | 0,000416014 | 0,305180611 | 0,379 | 0,219 | 1 | 0 | Ick |
| Anxa3 | 0,000419765 | 0,413171897 | 0,811 | 0,763 | 1 | 0 | Anxa3 |
| Ocln | 0,000434672 | 0,420606076 | 0,274 | 0,123 | 1 | 0 | Ocln |
| Anxa2 | 0,000435631 | 0,263843242 | 0,968 | 0,829 | 1 | 0 | Anxa2 |
| Cald1 | 0,000436311 | 0,292473544 | 0,821 | 0,588 | 1 | 0 | Cald1 |

| | | | | | | | |
|---|---|---|---|---|---|---|---|
| Mapkapk3 | 1,11E-31 | 0,677580928 | 0,531 | 0,008 | 1,76E-27 | MAC | Mapkapk3 |
| Gm22133 | 1,11E-31 | 0,520499036 | 0,963 | 0,781 | 1,76E-27 | MAC | Gm22133 |
| Sh3bp1 | 1,66E-31 | 0,546644652 | 0,543 | 0,012 | 2,63E-27 | MAC | Sh3bp1 |
| Tgfbr1 | 3,01E-31 | 1,042124834 | 0,901 | 0,207 | 4,78E-27 | MAC | Tgfbr1 |
| Lsp1 | 3,05E-31 | 1,015425272 | 0,494 | 0 | 4,83E-27 | MAC | Lsp1 |
| Arhgap9 | 3,05E-31 | 0,534123342 | 0,494 | 0 | 4,83E-27 | MAC | Arhgap9 |
| Tagap | 3,26E-31 | 1,05180134 | 0,519 | 0,008 | 5,17E-27 | MAC | Tagap |
| Milr1 | 3,83E-31 | 0,769847035 | 0,568 | 0,025 | 6,07E-27 | MAC | Milr1 |
| Fyb | 6,18E-31 | 1,187172025 | 0,914 | 0,355 | 9,80E-27 | MAC | Fyb |
| Prkcb | 7,05E-31 | 0,544190178 | 0,506 | 0,004 | 1,12E-26 | MAC | Prkcb |
| Atp6v0c | 1,36E-30 | 1,035339302 | 1 | 0,872 | 2,16E-26 | MAC | Atp6v0c |
| Arhgap15 | 2,07E-30 | 0,587595427 | 0,481 | 0 | 3,28E-26 | MAC | Arhgap15 |
| Runx3 | 2,07E-30 | 0,568885157 | 0,481 | 0 | 3,28E-26 | MAC | Runx3 |
| Hexa | 2,18E-30 | 1,367739242 | 0,914 | 0,322 | 3,45E-26 | MAC | Hexa |
| Itgax | 2,34E-30 | 0,940423925 | 0,494 | 0,004 | 3,70E-26 | MAC | Itgax |
| Tep1 | 2,97E-30 | 0,925350018 | 0,728 | 0,112 | 4,70E-26 | MAC | Tep1 |
| Emilin2 | 3,21E-30 | 0,735592592 | 0,519 | 0,012 | 5,08E-26 | MAC | Emilin2 |
| Cyth4 | 4,45E-30 | 1,136689241 | 0,827 | 0,202 | 7,05E-26 | MAC | Cyth4 |
| Arap1 | 6,23E-30 | 0,662836078 | 0,642 | 0,062 | 9,87E-26 | MAC | Arap1 |
| Arhgap19 | 1,09E-29 | 0,721811133 | 0,556 | 0,029 | 1,73E-25 | MAC | Arhgap19 |
| Abcc3 | 1,19E-29 | 0,69645675 | 0,556 | 0,025 | 1,89E-25 | MAC | Abcc3 |
| Ighm | 1,39E-29 | 1,006104576 | 0,469 | 0 | 2,20E-25 | MAC | Ighm |
| Blnk | 1,39E-29 | 0,9001503 | 0,469 | 0 | 2,20E-25 | MAC | Blnk |
| Col14a1 | 1,39E-29 | 0,803681873 | 0,469 | 0 | 2,20E-25 | MAC | Col14a1 |
| Clec5a | 1,39E-29 | 0,783345542 | 0,469 | 0 | 2,20E-25 | MAC | Clec5a |
| Cxcl14 | 1,56E-29 | 0,679460214 | 0,481 | 0,004 | 2,48E-25 | MAC | Cxcl14 |
| Icosl | 1,77E-29 | 0,311099298 | 0,543 | 0,021 | 2,81E-25 | MAC | Icosl |
| Rps29 | 2,74E-29 | 0,619448213 | 1 | 0,996 | 4,35E-25 | MAC | Rps29 |
| Notch2 | 3,02E-29 | 0,632063456 | 0,901 | 0,252 | 4,79E-25 | MAC | Notch2 |
| Tpd52 | 3,87E-29 | 0,979458038 | 0,938 | 0,273 | 6,14E-25 | MAC | Tpd52 |
| Rassf4 | 4,41E-29 | 0,844686742 | 0,642 | 0,066 | 6,99E-25 | MAC | Rassf4 |
| Fam49b | 4,94E-29 | 1,225835016 | 0,901 | 0,335 | 7,83E-25 | MAC | Fam49b |
| Ptafr | 1,06E-28 | 0,605476117 | 0,469 | 0,004 | 1,68E-24 | MAC | Ptafr |
| Csf3r | 1,12E-28 | 0,796284593 | 0,469 | 0,004 | 1,78E-24 | MAC | Csf3r |
| Nfam1 | 1,13E-28 | 0,766262085 | 0,481 | 0,008 | 1,79E-24 | MAC | Nfam1 |
| Gns | 1,39E-28 | 1,237033357 | 0,938 | 0,339 | 2,21E-24 | MAC | Gns |
| Metrnl | 1,71E-28 | 0,712839286 | 0,531 | 0,025 | 2,71E-24 | MAC | Metrnl |
| Ptk2b | 2,50E-28 | 0,520270734 | 0,481 | 0,008 | 3,96E-24 | MAC | Ptk2b |
| Spp1 | 4,51E-28 | 2,849304487 | 0,568 | 0,045 | 7,14E-24 | MAC | Spp1 |
| Hvcn1 | 6,06E-28 | 0,687798307 | 0,444 | 0 | 9,61E-24 | MAC | Hvcn1 |
| Siglece | 6,06E-28 | 0,62331922 | 0,444 | 0 | 9,61E-24 | MAC | Siglece |
| Stk17b | 7,55E-28 | 0,953648699 | 0,63 | 0,07 | 1,20E-23 | MAC | Stk17b |
| Gmfg | 7,74E-28 | 0,786944147 | 0,889 | 0,264 | 1,23E-23 | MAC | Gmfg |
| B4galnt1 | 1,45E-27 | 0,745542904 | 0,481 | 0,012 | 2,30E-23 | MAC | B4galnt1 |
| Il6ra | 1,51E-27 | 0,578527358 | 0,494 | 0,017 | 2,40E-23 | MAC | Il6ra |
| Glipr1 | 1,56E-27 | 0,594104761 | 0,543 | 0,037 | 2,47E-23 | MAC | Glipr1 |
| Creg1 | 1,86E-27 | 0,938382843 | 0,778 | 0,161 | 2,94E-23 | MAC | Creg1 |
| Fmnl1 | 2,48E-27 | 0,397439907 | 0,568 | 0,041 | 3,92E-23 | MAC | Fmnl1 |
| Gm13166 | 3,24E-27 | 0,267573573 | 0,765 | 0,178 | 5,13E-23 | MAC | Gm13166 |
| Pik3ap1 | 3,93E-27 | 0,46500539 | 0,432 | 0 | 6,24E-23 | MAC | Pik3ap1 |
| Pou2f2 | 3,93E-27 | 0,374063164 | 0,432 | 0 | 6,24E-23 | MAC | Pou2f2 |
| Slc6a6 | 4,03E-27 | 0,792335408 | 0,877 | 0,231 | 6,39E-23 | MAC | Slc6a6 |
| Ms4a4c | 4,73E-27 | 0,992016276 | 0,444 | 0,004 | 7,51E-23 | MAC | Ms4a4c |
| AW112010 | 5,62E-27 | 1,50202659 | 0,827 | 0,236 | 8,91E-23 | MAC | AW112010 |
| H2-Ab1 | 5,67E-27 | 2,549173623 | 0,852 | 0,306 | 8,98E-23 | MAC | H2-Ab1 |
| Camk1d | 6,91E-27 | 0,489802103 | 0,444 | 0,004 | 1,09E-22 | MAC | Camk1d |
| Slfn2 | 7,35E-27 | 0,99484791 | 0,802 | 0,174 | 1,17E-22 | MAC | Slfn2 |
| Fnbp1 | 1,30E-26 | 0,903192124 | 0,778 | 0,174 | 2,05E-22 | MAC | Fnbp1 |
| Selenop | 1,67E-26 | 1,277941566 | 0,988 | 0,818 | 2,64E-22 | MAC | Selenop |
| Tcirg1 | 2,03E-26 | 1,326977134 | 0,889 | 0,376 | 3,22E-22 | MAC | Tcirg1 |
| Fos | 2,48E-26 | 1,538335635 | 0,951 | 0,467 | 3,93E-22 | MAC | Fos |
| Gm14303 | 3,50E-26 | 0,508652242 | 1 | 0,983 | 5,55E-22 | MAC | Gm14303 |
| Rassf5 | 3,65E-26 | 0,824555347 | 0,432 | 0,004 | 5,79E-22 | MAC | Rassf5 |
| Susd3 | 5,46E-26 | 0,617745883 | 0,444 | 0,008 | 8,66E-22 | MAC | Susd3 |
| Abhd12 | 5,56E-26 | 0,777833898 | 0,802 | 0,178 | 8,82E-22 | MAC | Abhd12 |
| Havcr2 | 6,08E-26 | 0,515268894 | 0,432 | 0,004 | 9,63E-22 | MAC | Havcr2 |
| Arhgef6 | 8,94E-26 | 0,368266832 | 0,457 | 0,012 | 1,42E-21 | MAC | Arhgef6 |
| Fabp5 | 1,08E-25 | 0,704699434 | 0,469 | 0,017 | 1,71E-21 | MAC | Fabp5 |
| Plcg2 | 1,25E-25 | 0,660951769 | 0,469 | 0,017 | 1,98E-21 | MAC | Plcg2 |
| Slc43a2 | 1,29E-25 | 0,635968875 | 0,667 | 0,103 | 2,04E-21 | MAC | Slc43a2 |
| Tnfsf13b | 1,60E-25 | 0,737616777 | 0,407 | 0 | 2,54E-21 | MAC | Tnfsf13b |
| Pag1 | 1,60E-25 | 0,347251411 | 0,407 | 0 | 2,54E-21 | MAC | Pag1 |
| Dab2 | 2,06E-25 | 1,346638505 | 0,926 | 0,492 | 3,26E-21 | MAC | Dab2 |
| Syngr1 | 2,13E-25 | 0,476112442 | 0,42 | 0,004 | 3,38E-21 | MAC | Syngr1 |
| Arhgap22 | 2,78E-25 | 0,560932159 | 0,42 | 0,004 | 4,41E-21 | MAC | Arhgap22 |
| Daglb | 3,19E-25 | 0,771501764 | 0,519 | 0,037 | 5,05E-21 | MAC | Daglb |
| Zfp991 | 3,63E-25 | 0,386443859 | 0,593 | 0,062 | 5,75E-21 | MAC | Zfp991 |
| Cd63 | 5,33E-25 | 1,30519191 | 0,951 | 0,595 | 8,45E-21 | MAC | Cd63 |
| Dnase2a | 5,48E-25 | 0,613567353 | 0,519 | 0,037 | 8,69E-21 | MAC | Dnase2a |
| Scpep1 | 5,65E-25 | 0,761035765 | 0,79 | 0,165 | 8,96E-21 | MAC | Scpep1 |
| Snx20 | 6,89E-25 | 0,603895727 | 0,481 | 0,029 | 1,09E-20 | MAC | Snx20 |
| Chd9 | 7,98E-25 | 0,687602891 | 0,951 | 0,43 | 1,27E-20 | MAC | Chd9 |
| Sash3 | 1,01E-24 | 0,586670948 | 0,395 | 0 | 1,60E-20 | MAC | Sash3 |
| Cd300ld | 1,01E-24 | 0,336144078 | 0,395 | 0 | 1,60E-20 | MAC | Cd300ld |
| Pik3cd | 1,30E-24 | 0,684825339 | 0,407 | 0,004 | 2,05E-20 | MAC | Pik3cd |
| Lrmp | 1,33E-24 | 0,516033805 | 0,407 | 0,004 | 2,11E-20 | MAC | Lrmp |
| Kcnn4 | 1,56E-24 | 0,674958892 | 0,407 | 0,004 | 2,48E-20 | MAC | Kcnn4 |
| Cxcl2 | 1,59E-24 | 1,195320277 | 0,42 | 0,008 | 2,53E-20 | MAC | Cxcl2 |
| Cd300a | 1,73E-24 | 0,506783022 | 0,444 | 0,017 | 2,74E-20 | MAC | Cd300a |
| Id2 | 2,40E-24 | 1,296325932 | 0,827 | 0,277 | 3,81E-20 | MAC | Id2 |
| Ppm1h | 5,17E-24 | 0,655674662 | 0,63 | 0,091 | 8,20E-20 | MAC | Ppm1h |
| Slc9a3r1 | 6,01E-24 | 0,666075571 | 0,481 | 0,029 | 9,53E-20 | MAC | Slc9a3r1 |
| Sorl1 | 6,27E-24 | 0,497958317 | 0,383 | 0 | 9,93E-20 | MAC | Sorl1 |
| Gcnt1 | 6,27E-24 | 0,36546244 | 0,383 | 0 | 9,93E-20 | MAC | Gcnt1 |
| Pla2g15 | 7,16E-24 | 0,523222522 | 0,457 | 0,021 | 1,13E-19 | MAC | Pla2g15 |
| Pfkfb4 | 8,91E-24 | 0,475324266 | 0,395 | 0,004 | 1,41E-19 | MAC | Pfkfb4 |
| Gpr34 | 9,40E-24 | 0,701872507 | 0,395 | 0,004 | 1,49E-19 | MAC | Gpr34 |
| Egr2 | 9,43E-24 | 0,940208335 | 0,432 | 0,017 | 1,50E-19 | MAC | Egr2 |
| Tnfaip8 | 1,09E-23 | 0,914377619 | 0,765 | 0,186 | 1,73E-19 | MAC | Tnfaip8 |
| Tnf | 1,11E-23 | 0,938191585 | 0,407 | 0,008 | 1,76E-19 | MAC | Tnf |
| Osm | 3,85E-23 | 0,823072714 | 0,37 | 0 | 6,11E-19 | MAC | Osm |
| Cyp27a1 | 3,85E-23 | 0,713416811 | 0,37 | 0 | 6,11E-19 | MAC | Cyp27a1 |
| Scimp | 3,85E-23 | 0,66422555 | 0,37 | 0 | 6,11E-19 | MAC | Scimp |
| Arhgap4 | 3,85E-23 | 0,49667114 | 0,37 | 0 | 6,11E-19 | MAC | Arhgap4 |
| Slamf7 | 3,85E-23 | 0,433141015 | 0,37 | 0 | 6,11E-19 | MAC | Slamf7 |
| Lgals3bp | 4,49E-23 | 1,092193382 | 0,926 | 0,384 | 7,11E-19 | MAC | Lgals3bp |
| Snx18 | 4,82E-23 | 0,730944186 | 0,543 | 0,066 | 7,63E-19 | MAC | Snx18 |
| Asah1 | 4,94E-23 | 0,999397526 | 0,951 | 0,508 | 7,83E-19 | MAC | Asah1 |
| Esr1 | 5,93E-23 | 0,580748083 | 0,556 | 0,066 | 9,41E-19 | MAC | Esr1 |
| Pim1 | 6,99E-23 | 0,306871403 | 0,815 | 0,248 | 1,11E-18 | MAC | Pim1 |
| Arl4c | 7,36E-23 | 0,627292835 | 0,58 | 0,079 | 1,17E-18 | MAC | Arl4c |
| Cadm1 | 9,60E-23 | 0,62633455 | 0,543 | 0,058 | 1,52E-18 | MAC | Cadm1 |
| Dse | 1,54E-22 | 0,388360609 | 0,407 | 0,012 | 2,44E-18 | MAC | Dse |
| Gpsm3 | 2,14E-22 | 0,40707025 | 0,667 | 0,136 | 3,40E-18 | MAC | Gpsm3 |
| Msrb1 | 2,19E-22 | 0,303364052 | 0,802 | 0,306 | 3,48E-18 | MAC | Msrb1 |
| C4b | 2,34E-22 | 0,710978666 | 0,358 | 0 | 3,72E-18 | MAC | C4b |
| Tlr1 | 2,34E-22 | 0,67315276 | 0,358 | 0 | 3,72E-18 | MAC | Tlr1 |
| Pik3r5 | 2,34E-22 | 0,549391712 | 0,358 | 0 | 3,72E-18 | MAC | Pik3r5 |
| Ptpn22 | 2,34E-22 | 0,434309744 | 0,358 | 0 | 3,72E-18 | MAC | Ptpn22 |
| Wdfy4 | 2,55E-22 | 0,504190759 | 0,444 | 0,025 | 4,04E-18 | MAC | Wdfy4 |

| | | | | | | | |
|---|---|---|---|---|---|---|---|
| Cdr2l | 0,000437714 | 0,365629767 | 0,589 | 0,386 | 1 | 0 | Cdr2l |
| Tspan5 | 0,00044374 | 0,303447469 | 0,358 | 0,189 | 1 | 0 | Tspan5 |
| Sorbs2 | 0,000446037 | 0,322025032 | 0,547 | 0,351 | 1 | 0 | Sorbs2 |
| Eogt | 0,000466652 | 0,55309757 | 0,568 | 0,373 | 1 | 0 | Eogt |
| Fam63a | 0,000496867 | 0,328072977 | 0,484 | 0,303 | 1 | 0 | Fam63a |
| Ybx1 | 0,000510261 | 0,301973322 | 0,937 | 0,952 | 1 | 0 | Ybx1 |
| Tspan13 | 0,000529028 | 0,566830712 | 0,589 | 0,417 | 1 | 0 | Tspan13 |
| Nck1 | 0,000540894 | 0,363930391 | 0,495 | 0,325 | 1 | 0 | Nck1 |
| Fli1 | 0,00056668 | 0,482014207 | 0,905 | 0,825 | 1 | 0 | Fli1 |
| Timp1 | 0,000603247 | 0,25001302 | 0,442 | 0,276 | 1 | 0 | Timp1 |
| Pdia3 | 0,000705182 | 0,293079919 | 0,958 | 0,961 | 1 | 0 | Pdia3 |
| Als2cl | 0,000725488 | 0,275538634 | 0,326 | 0,167 | 1 | 0 | Als2cl |
| Malat1 | 0,000732849 | 0,427893197 | 1 | 0,996 | 1 | 0 | Malat1 |
| Fzd6 | 0,000741442 | 0,300482452 | 0,421 | 0,25 | 1 | 0 | Fzd6 |
| Pros1 | 0,000847969 | 0,466408979 | 0,6 | 0,443 | 1 | 0 | Pros1 |
| Tnfaip1 | 0,000866154 | 0,29941525 | 0,811 | 0,627 | 1 | 0 | Tnfaip1 |
| Gem | 0,000924115 | 0,290103526 | 0,316 | 0,154 | 1 | 0 | Gem |
| Kank3 | 0,000942144 | 0,410702686 | 0,621 | 0,421 | 1 | 0 | Kank3 |
| Luzp1 | 0,000955239 | 0,398114495 | 0,716 | 0,627 | 1 | 0 | Luzp1 |
| Lpcat3 | 0,000982687 | 0,320037629 | 0,379 | 0,215 | 1 | 0 | Lpcat3 |
| Tsc22d3 | 0,000990306 | 0,418887077 | 0,695 | 0,557 | 1 | 0 | Tsc22d3 |
| Gpr146 | 0,001009936 | 0,346527991 | 0,547 | 0,382 | 1 | 0 | Gpr146 |
| Cd81 | 0,001013418 | 0,262863202 | 0,958 | 0,987 | 1 | 0 | Cd81 |
| B3gnt3 | 0,001019346 | 0,30832275 | 0,695 | 0,509 | 1 | 0 | B3gnt3 |
| Fam198b | 0,001033768 | 0,325664713 | 0,263 | 0,118 | 1 | 0 | Fam198b |
| Rnf213 | 0,00104328 | 0,323481113 | 0,579 | 0,404 | 1 | 0 | Rnf213 |
| Laptm4a | 0,001109159 | 0,305544752 | 0,895 | 0,798 | 1 | 0 | Laptm4a |
| Tfpi | 0,001116847 | 0,434091458 | 0,568 | 0,377 | 1 | 0 | Tfpi |
| Gnas | 0,001120349 | 0,288612431 | 0,937 | 0,882 | 1 | 0 | Gnas |
| Cd320 | 0,001195044 | 0,265074555 | 0,284 | 0,14 | 1 | 0 | Cd320 |
| Ptpn14 | 0,001217187 | 0,346765487 | 0,379 | 0,224 | 1 | 0 | Ptpn14 |
| 2010111I01Rik | 0,001264608 | 0,470480942 | 0,705 | 0,522 | 1 | 0 | 2010111I01Rik |
| Stt3b | 0,001275069 | 0,396045955 | 0,632 | 0,496 | 1 | 0 | Stt3b |
| Ehd2 | 0,001363662 | 0,268284944 | 0,674 | 0,478 | 1 | 0 | Ehd2 |
| Nedd4 | 0,001446199 | 0,296919835 | 0,863 | 0,658 | 1 | 0 | Nedd4 |
| Lta4h | 0,001507136 | 0,308033744 | 0,642 | 0,439 | 1 | 0 | Lta4h |
| Notch1 | 0,001530752 | 0,409702913 | 0,768 | 0,64 | 1 | 0 | Notch1 |
| Dusp3 | 0,001556665 | 0,302845777 | 0,937 | 0,798 | 1 | 0 | Dusp3 |
| Pecam1 | 0,00155992 | 0,274808379 | 0,979 | 0,684 | 1 | 0 | Pecam1 |
| Klf2 | 0,001812594 | 0,414434858 | 0,547 | 0,417 | 1 | 0 | Klf2 |
| Jag2 | 0,001890763 | 0,296217663 | 0,411 | 0,254 | 1 | 0 | Jag2 |
| Slc12a7 | 0,00189465 | 0,348685294 | 0,505 | 0,338 | 1 | 0 | Slc12a7 |
| Emp1 | 0,002134327 | 0,312457539 | 0,853 | 0,711 | 1 | 0 | Emp1 |
| Reep3 | 0,002201634 | 0,330127794 | 0,695 | 0,548 | 1 | 0 | Reep3 |
| Arhgap31 | 0,002247071 | 0,316334004 | 0,716 | 0,561 | 1 | 0 | Arhgap31 |
| Ptn | 0,002358197 | 0,589139504 | 0,305 | 0,158 | 1 | 0 | Ptn |
| Cdc42bpb | 0,002633701 | 0,253881473 | 0,611 | 0,452 | 1 | 0 | Cdc42bpb |
| 4931406P16Rik | 0,002635948 | 0,27348609 | 0,611 | 0,469 | 1 | 0 | 4931406P16Rik |
| Lima1 | 0,0026486 | 0,291039443 | 0,421 | 0,268 | 1 | 0 | Lima1 |
| C130074G19Rik | 0,002869102 | 0,307122235 | 0,358 | 0,224 | 1 | 0 | C130074G19Rik |
| Mrpl17 | 0,003275733 | 0,275298199 | 0,779 | 0,693 | 1 | 0 | Mrpl17 |
| Gja1 | 0,003288856 | 0,381821615 | 0,758 | 0,618 | 1 | 0 | Gja1 |
| Cpne8 | 0,003292282 | 0,304366827 | 0,568 | 0,447 | 1 | 0 | Cpne8 |
| Slc25a4 | 0,003325824 | 0,294598278 | 0,895 | 0,833 | 1 | 0 | Slc25a4 |
| Pnrc2 | 0,003601621 | 0,341559742 | 0,695 | 0,57 | 1 | 0 | Pnrc2 |
| Ahr | 0,003604528 | 0,30136991 | 0,305 | 0,171 | 1 | 0 | Ahr |
| Plxna2 | 0,003693453 | 0,314001685 | 0,495 | 0,351 | 1 | 0 | Plxna2 |
| Cttn | 0,003890766 | 0,378094084 | 0,6 | 0,43 | 1 | 0 | Cttn |
| Olfml2a | 0,004411022 | 0,38995935 | 0,442 | 0,303 | 1 | 0 | Olfml2a |
| Cd151 | 0,004649086 | 0,315723406 | 0,579 | 0,434 | 1 | 0 | Cd151 |
| Pls3 | 0,004750182 | 0,349904208 | 0,705 | 0,539 | 1 | 0 | Pls3 |
| Ywhab | 0,004792694 | 0,338421624 | 0,884 | 0,873 | 1 | 0 | Ywhab |
| Dnajc8 | 0,005209188 | 0,287194896 | 0,737 | 0,711 | 1 | 0 | Dnajc8 |
| Zhx1 | 0,005286704 | 0,493134556 | 0,537 | 0,417 | 1 | 0 | Zhx1 |
| Cdipt | 0,005394486 | 0,332928397 | 0,695 | 0,618 | 1 | 0 | Cdipt |
| Spns2 | 0,005482653 | 0,278100945 | 0,379 | 0,241 | 1 | 0 | Spns2 |
| Gsk3b | 0,005486481 | 0,296291168 | 0,884 | 0,781 | 1 | 0 | Gsk3b |
| Fkbp9 | 0,005613439 | 0,333288436 | 0,495 | 0,355 | 1 | 0 | Fkbp9 |
| Lrrc8a | 0,00577684 | 0,252762414 | 0,884 | 0,746 | 1 | 0 | Lrrc8a |
| Cdc42ep2 | 0,006004243 | 0,306080408 | 0,379 | 0,241 | 1 | 0 | Cdc42ep2 |
| Ap2m1 | 0,006209123 | 0,257397884 | 0,895 | 0,895 | 1 | 0 | Ap2m1 |
| Slc39a10 | 0,00669673 | 0,360563947 | 0,4 | 0,272 | 1 | 0 | Slc39a10 |
| Bst1 | 0,006748197 | 0,300239034 | 0,295 | 0,175 | 1 | 0 | Bst1 |
| Uqcc3 | 0,007051579 | 0,319087289 | 0,547 | 0,439 | 1 | 0 | Uqcc3 |
| Myo1b | 0,007133771 | 0,275276827 | 0,695 | 0,513 | 1 | 0 | Myo1b |
| St3gal6 | 0,00728455 | 0,390779936 | 0,484 | 0,373 | 1 | 0 | St3gal6 |
| Procr | 0,007634383 | 0,259257872 | 0,389 | 0,259 | 1 | 0 | Procr |
| Nav1 | 0,008012809 | 0,350536084 | 0,547 | 0,421 | 1 | 0 | Nav1 |
| Rtp4 | 0,008092864 | 0,321528064 | 0,432 | 0,294 | 1 | 0 | Rtp4 |
| Mtmr11 | 0,008220749 | 0,286800135 | 0,347 | 0,206 | 1 | 0 | Mtmr11 |
| Fstl1 | 0,008546979 | 0,2987209 | 0,611 | 0,5 | 1 | 0 | Fstl1 |
| Nrp2 | 0,009025843 | 0,389168182 | 0,579 | 0,474 | 1 | 0 | Nrp2 |
| Hspb8 | 0,009388122 | 0,266084662 | 0,568 | 0,421 | 1 | 0 | Hspb8 |
| Lilrb4a | 1,77E-69 | 2,400385012 | 1 | 0,004 | 2,80E-65 | 1 | Lilrb4a |
| Bcl2a1b | 1,77E-69 | 2,189161117 | 1 | 0,004 | 2,80E-65 | 1 | Bcl2a1b |
| Mpeg1 | 5,11E-69 | 2,010932239 | 0,988 | 0 | 8,10E-65 | 1 | Mpeg1 |
| Tyrobp | 6,88E-69 | 2,918688965 | 1 | 0,008 | 1,09E-64 | 1 | Tyrobp |
| Lilr4b | 2,46E-68 | 2,329919561 | 0,988 | 0,004 | 3,90E-64 | 1 | Lilr4b |
| Fcgr3 | 2,59E-68 | 2,967713703 | 1 | 0,012 | 4,10E-64 | 1 | Fcgr3 |
| Cd52 | 2,59E-68 | 2,278192308 | 1 | 0,012 | 4,10E-64 | 1 | Cd52 |
| Aif1 | 2,59E-68 | 2,103100042 | 1 | 0,012 | 4,10E-64 | 1 | Aif1 |
| Ly86 | 5,88E-68 | 1,946878729 | 0,975 | 0 | 9,32E-64 | 1 | Ly86 |
| Spi1 | 5,88E-68 | 1,790833527 | 0,975 | 0 | 9,32E-64 | 1 | Spi1 |
| Ifi30 | 1,13E-67 | 2,182814707 | 1 | 0,017 | 1,80E-63 | 1 | Ifi30 |
| Apobec1 | 6,67E-67 | 1,931855482 | 0,963 | 0 | 1,06E-62 | 1 | Apobec1 |
| C1qa | 1,13E-66 | 3,788283771 | 1 | 0,025 | 1,80E-62 | 1 | C1qa |
| Fcgr2b | 1,13E-66 | 2,744112754 | 1 | 0,025 | 1,80E-62 | 1 | Fcgr2b |
| Laptm5 | 1,68E-66 | 2,361074799 | 1 | 0,025 | 2,67E-62 | 1 | Laptm5 |
| Csf1r | 3,76E-66 | 2,692010077 | 1 | 0,029 | 5,96E-62 | 1 | Csf1r |
| Pla2g7 | 4,48E-66 | 2,385132456 | 0,988 | 0,021 | 7,09E-62 | 1 | Pla2g7 |
| C1qc | 1,21E-65 | 4,125327152 | 1 | 0,033 | 1,92E-61 | 1 | C1qc |
| Coro1a | 1,37E-65 | 2,157053059 | 1 | 0,029 | 2,18E-61 | 1 | Coro1a |
| Trem2 | 3,33E-65 | 1,787857377 | 0,951 | 0,004 | 5,28E-61 | 1 | Trem2 |
| C3ar1 | 8,20E-65 | 1,837999673 | 0,938 | 0 | 1,30E-60 | 1 | C3ar1 |
| Cd300c2 | 8,20E-65 | 1,436111737 | 0,938 | 0 | 1,30E-60 | 1 | Cd300c2 |
| Pld4 | 1,34E-64 | 2,292815016 | 0,951 | 0,008 | 2,13E-60 | 1 | Pld4 |
| Rgs1 | 3,61E-64 | 3,074974665 | 0,938 | 0,004 | 5,73E-60 | 1 | Rgs1 |
| Itgam | 3,61E-64 | 1,531582192 | 0,938 | 0,004 | 5,73E-60 | 1 | Itgam |
| Cybb | 5,92E-64 | 1,695465599 | 0,951 | 0,012 | 9,39E-60 | 1 | Cybb |
| Ms4a7 | 6,11E-64 | 1,94153161 | 0,951 | 0,012 | 9,68E-60 | 1 | Ms4a7 |
| Cd68 | 1,05E-63 | 2,337275647 | 1 | 0,05 | 1,67E-59 | 1 | Cd68 |
| C1qb | 2,80E-63 | 4,258745352 | 1 | 0,054 | 4,44E-59 | 1 | C1qb |
| Ms4a6c | 6,25E-63 | 1,450614627 | 0,938 | 0,012 | 9,90E-59 | 1 | Ms4a6c |
| Lyz2 | 7,73E-63 | 4,475989832 | 1 | 0,058 | 1,23E-58 | 1 | Lyz2 |
| Ctss | 7,73E-63 | 4,33950261 | 1 | 0,058 | 1,23E-58 | 1 | Ctss |
| Cyp4f18 | 9,52E-63 | 1,741252522 | 0,914 | 0 | 1,51E-58 | 1 | Cyp4f18 |
| Ptprc | 9,52E-63 | 1,496797575 | 0,914 | 0 | 1,51E-58 | 1 | Ptprc |
| Alox5ap | 2,76E-62 | 1,886439132 | 0,963 | 0,025 | 4,38E-58 | 1 | Alox5ap |
| Sirpb1c | 4,34E-62 | 1,470039352 | 0,914 | 0,004 | 6,88E-58 | 1 | Sirpb1c |
| Rgs10 | 1,00E-61 | 1,276072511 | 0,901 | 0 | 1,59E-57 | 1 | Rgs10 |

| | | | | | | | |
|---|---|---|---|---|---|---|---|
| Ang | 3,07E-22 | 0,531061683 | 0,457 | 0,033 | 4,87E-18 | MAC | Ang |
| Was | 3,82E-22 | 0,336419105 | 0,395 | 0,012 | 6,06E-18 | MAC | Was |
| Gpx3 | 5,36E-22 | 0,965896029 | 0,741 | 0,186 | 8,50E-18 | MAC | Gpx3 |
| Rps6ka1 | 5,65E-22 | 0,685936274 | 0,63 | 0,107 | 8,95E-18 | MAC | Rps6ka1 |
| Cstb | 1,09E-21 | 0,962951432 | 0,951 | 0,657 | 1,72E-17 | MAC | Cstb |
| Vcam1 | 1,14E-21 | 1,26914205 | 0,716 | 0,174 | 1,81E-17 | MAC | Vcam1 |
| Ppcdc | 1,37E-21 | 0,843662871 | 0,593 | 0,099 | 2,17E-17 | MAC | Ppcdc |
| Cmtm7 | 1,40E-21 | 0,78567478 | 0,753 | 0,215 | 2,22E-17 | MAC | Cmtm7 |
| Il1b | 1,41E-21 | 1,172471008 | 0,346 | 0 | 2,24E-17 | MAC | Il1b |
| Fam46c | 1,41E-21 | 0,651824784 | 0,346 | 0 | 2,24E-17 | MAC | Fam46c |
| P2ry14 | 1,41E-21 | 0,565459209 | 0,346 | 0 | 2,24E-17 | MAC | P2ry14 |
| Klra17 | 1,41E-21 | 0,522474672 | 0,346 | 0 | 2,24E-17 | MAC | Klra17 |
| Abhd15 | 1,41E-21 | 0,445438485 | 0,346 | 0 | 2,24E-17 | MAC | Abhd15 |
| I830077J02Rik | 1,41E-21 | 0,303579479 | 0,346 | 0 | 2,24E-17 | MAC | I830077J02Rik |
| Tmem86a | 1,73E-21 | 0,710968724 | 0,654 | 0,12 | 2,75E-17 | MAC | Tmem86a |
| Tspan32 | 2,03E-21 | 0,517061799 | 0,358 | 0,004 | 3,21E-17 | MAC | Tspan32 |
| Adssl1 | 2,76E-21 | 0,674850795 | 0,395 | 0,017 | 4,38E-17 | MAC | Adssl1 |
| Ptgs1 | 3,59E-21 | 1,004198622 | 0,679 | 0,153 | 5,69E-17 | MAC | Ptgs1 |
| P2rx4 | 3,64E-21 | 0,958053931 | 0,741 | 0,223 | 5,78E-17 | MAC | P2rx4 |
| Tmsb4x | 3,78E-21 | 0,524215299 | 1 | 1 | 5,99E-17 | MAC | Tmsb4x |
| Blvrb | 4,16E-21 | 0,595817096 | 0,667 | 0,136 | 6,59E-17 | MAC | Blvrb |
| Sowahc | 8,27E-21 | 0,597567389 | 0,444 | 0,033 | 1,31E-16 | MAC | Sowahc |
| Arhgap24 | 8,41E-21 | 0,388451462 | 0,333 | 0 | 1,33E-16 | MAC | Arhgap24 |
| Pstpip1 | 8,41E-21 | 0,362602899 | 0,333 | 0 | 1,33E-16 | MAC | Pstpip1 |
| Clec4d | 8,41E-21 | 0,323312751 | 0,333 | 0 | 1,33E-16 | MAC | Clec4d |
| Lamp1 | 9,83E-21 | 0,800039046 | 1 | 0,893 | 1,56E-16 | MAC | Lamp1 |
| B3gnt8 | 9,99E-21 | 0,568065754 | 0,494 | 0,066 | 1,58E-16 | MAC | B3gnt8 |
| Uap1l1 | 1,07E-20 | 0,807724392 | 0,494 | 0,058 | 1,70E-16 | MAC | Uap1l1 |
| Rnaset2a | 1,09E-20 | 0,481170359 | 0,951 | 0,471 | 1,73E-16 | MAC | Rnaset2a |
| AI662270 | 1,57E-20 | 0,43863666 | 0,432 | 0,033 | 2,48E-16 | MAC | AI662270 |
| Gltp | 1,81E-20 | 0,899557904 | 0,815 | 0,31 | 2,87E-16 | MAC | Gltp |
| Akna | 2,02E-20 | 0,577668582 | 0,494 | 0,054 | 3,21E-16 | MAC | Akna |
| Srgn | 3,25E-20 | 1,041232525 | 0,963 | 0,736 | 5,16E-16 | MAC | Srgn |
| Cysltr1 | 3,48E-20 | 0,562471883 | 0,383 | 0,017 | 5,51E-16 | MAC | Cysltr1 |
| P2ry13 | 4,96E-20 | 0,525826406 | 0,321 | 0 | 7,87E-16 | MAC | P2ry13 |
| Lilra5 | 4,96E-20 | 0,443206512 | 0,321 | 0 | 7,87E-16 | MAC | Lilra5 |
| Acss1 | 4,96E-20 | 0,427991117 | 0,321 | 0 | 7,87E-16 | MAC | Acss1 |
| Cndp2 | 5,18E-20 | 0,959349471 | 0,84 | 0,314 | 8,21E-16 | MAC | Cndp2 |
| Ptpn18 | 7,08E-20 | 0,621629777 | 0,901 | 0,5 | 1,12E-15 | MAC | Ptpn18 |
| Ccdc88b | 8,01E-20 | 0,292537393 | 0,333 | 0,004 | 1,27E-15 | MAC | Ccdc88b |
| Eps8 | 1,04E-19 | 0,311074945 | 0,333 | 0,004 | 1,65E-15 | MAC | Eps8 |
| Ppt1 | 1,50E-19 | 0,966749552 | 0,914 | 0,421 | 2,37E-15 | MAC | Ppt1 |
| Pik3cg | 1,52E-19 | 0,48954774 | 0,481 | 0,054 | 2,41E-15 | MAC | Pik3cg |
| Soat1 | 1,76E-19 | 0,468381438 | 0,778 | 0,215 | 2,79E-15 | MAC | Soat1 |
| Neurl3 | 1,85E-19 | 0,672542852 | 0,593 | 0,107 | 2,93E-15 | MAC | Neurl3 |
| Ccnd2 | 2,14E-19 | 0,591870319 | 0,432 | 0,037 | 3,40E-15 | MAC | Ccnd2 |
| Slc37a2 | 2,75E-19 | 0,455059437 | 0,383 | 0,021 | 4,36E-15 | MAC | Slc37a2 |
| Rasal3 | 2,90E-19 | 0,529944769 | 0,309 | 0 | 4,59E-15 | MAC | Rasal3 |
| Tlr8 | 2,90E-19 | 0,49041397 | 0,309 | 0 | 4,59E-15 | MAC | Tlr8 |
| Btk | 2,90E-19 | 0,398201553 | 0,309 | 0 | 4,59E-15 | MAC | Btk |
| Gm26740 | 2,90E-19 | 0,346443608 | 0,309 | 0 | 4,59E-15 | MAC | Gm26740 |
| Dcxr | 2,92E-19 | 0,427762424 | 0,457 | 0,045 | 4,63E-15 | MAC | Dcxr |
| Smim3 | 3,36E-19 | 0,622848311 | 0,457 | 0,045 | 5,33E-15 | MAC | Smim3 |
| Irs2 | 4,52E-19 | 0,304990634 | 0,321 | 0,004 | 7,17E-15 | MAC | Irs2 |
| Lyl1 | 4,89E-19 | 0,696506323 | 0,506 | 0,079 | 7,75E-15 | MAC | Lyl1 |
| Lacc1 | 5,98E-19 | 0,517177254 | 0,395 | 0,029 | 9,48E-15 | MAC | Lacc1 |
| Dok3 | 6,36E-19 | 0,850796404 | 0,778 | 0,293 | 1,01E-14 | MAC | Dok3 |
| Timp2 | 8,10E-19 | 0,640926194 | 0,642 | 0,149 | 1,28E-14 | MAC | Timp2 |
| Cyfip2 | 8,40E-19 | 0,266014314 | 0,321 | 0,004 | 1,33E-14 | MAC | Cyfip2 |
| Rhoh | 9,49E-19 | 0,542404648 | 0,346 | 0,012 | 1,50E-14 | MAC | Rhoh |
| Hivep3 | 1,13E-18 | 0,42617658 | 0,469 | 0,054 | 1,80E-14 | MAC | Hivep3 |
| Rhog | 1,20E-18 | 0,758091983 | 0,963 | 0,562 | 1,90E-14 | MAC | Rhog |
| Gpr171 | 1,68E-18 | 0,588052816 | 0,296 | 0 | 2,66E-14 | MAC | Gpr171 |
| Myo1g | 1,68E-18 | 0,48580086 | 0,296 | 0 | 2,66E-14 | MAC | Myo1g |
| Galnt6 | 1,68E-18 | 0,477593558 | 0,296 | 0 | 2,66E-14 | MAC | Galnt6 |
| Ikbke | 1,68E-18 | 0,415045879 | 0,296 | 0 | 2,66E-14 | MAC | Ikbke |
| Dennd1c | 1,68E-18 | 0,381452533 | 0,296 | 0 | 2,66E-14 | MAC | Dennd1c |
| Atp1a3 | 1,68E-18 | 0,308285402 | 0,296 | 0 | 2,66E-14 | MAC | Atp1a3 |
| Rplp1 | 1,91E-18 | 0,4786856 | 1 | 0,988 | 3,03E-14 | MAC | Rplp1 |
| Csk | 2,82E-18 | 0,684396563 | 0,728 | 0,219 | 4,47E-14 | MAC | Csk |
| Il4ra | 3,01E-18 | 0,851831436 | 0,753 | 0,223 | 4,78E-14 | MAC | Il4ra |
| Junb | 3,33E-18 | 0,972479719 | 0,963 | 0,686 | 5,28E-14 | MAC | Junb |
| Arg1 | 3,35E-18 | 1,565771763 | 0,321 | 0,008 | 5,31E-14 | MAC | Arg1 |
| Kctd12 | 3,88E-18 | 0,713023622 | 0,704 | 0,202 | 6,14E-14 | MAC | Kctd12 |
| Dtx4 | 3,92E-18 | 0,357952817 | 0,395 | 0,029 | 6,21E-14 | MAC | Dtx4 |
| B3gnt7 | 4,29E-18 | 0,29619881 | 0,321 | 0,008 | 6,80E-14 | MAC | B3gnt7 |
| Lyn | 4,61E-18 | 0,857210201 | 0,938 | 0,537 | 7,31E-14 | MAC | Lyn |
| Tmem37 | 6,50E-18 | 0,795621974 | 0,728 | 0,244 | 1,03E-13 | MAC | Tmem37 |
| Pitpnm1 | 6,74E-18 | 0,666624923 | 0,42 | 0,045 | 1,07E-13 | MAC | Pitpnm1 |
| Hcls1 | 7,00E-18 | 0,681283415 | 0,901 | 0,409 | 1,11E-13 | MAC | Hcls1 |
| Sh3pxd2b | 7,33E-18 | 0,687197637 | 0,617 | 0,161 | 1,16E-13 | MAC | Sh3pxd2b |
| Pnpla7 | 7,50E-18 | 0,334212043 | 0,407 | 0,037 | 1,19E-13 | MAC | Pnpla7 |
| Abr | 7,54E-18 | 0,444957589 | 0,617 | 0,128 | 1,20E-13 | MAC | Abr |
| 5430427O19Rik | 9,59E-18 | 0,596711118 | 0,284 | 0 | 1,52E-13 | MAC | 5430427O19Rik |
| Tnfsf9 | 9,59E-18 | 0,3803071 | 0,284 | 0 | 1,52E-13 | MAC | Tnfsf9 |
| Il21r | 9,59E-18 | 0,360762191 | 0,284 | 0 | 1,52E-13 | MAC | Il21r |
| Shtn1 | 9,59E-18 | 0,295439637 | 0,284 | 0 | 1,52E-13 | MAC | Shtn1 |
| Nlrp3 | 9,59E-18 | 0,26010347 | 0,284 | 0 | 1,52E-13 | MAC | Nlrp3 |
| Gpr183 | 1,01E-17 | 0,492959763 | 0,469 | 0,058 | 1,60E-13 | MAC | Gpr183 |
| Mlxip | 1,05E-17 | 0,604818629 | 0,605 | 0,128 | 1,66E-13 | MAC | Mlxip |
| Ccrl2 | 1,20E-17 | 1,032785251 | 0,568 | 0,124 | 1,90E-13 | MAC | Ccrl2 |
| Ptger4 | 1,51E-17 | 0,535734658 | 0,296 | 0,004 | 2,40E-13 | MAC | Ptger4 |
| Gm9843 | 1,52E-17 | 0,446390455 | 1 | 0,988 | 2,41E-13 | MAC | Gm9843 |
| Gas6 | 2,04E-17 | 0,768216389 | 0,432 | 0,05 | 3,23E-13 | MAC | Gas6 |
| Rasgef1b | 2,19E-17 | 0,456818573 | 0,432 | 0,045 | 3,46E-13 | MAC | Rasgef1b |
| Gmip | 2,21E-17 | 0,50334324 | 0,457 | 0,058 | 3,51E-13 | MAC | Gmip |
| Trex1 | 2,52E-17 | 0,641870459 | 0,494 | 0,083 | 3,99E-13 | MAC | Trex1 |
| Tnfrsf13b | 2,72E-17 | 0,452468117 | 0,296 | 0,004 | 4,32E-13 | MAC | Tnfrsf13b |
| Cryl1 | 3,04E-17 | 0,304046136 | 0,383 | 0,029 | 4,82E-13 | MAC | Cryl1 |
| Egr1 | 3,06E-17 | 1,103825904 | 0,765 | 0,298 | 4,85E-13 | MAC | Egr1 |
| Myo5a | 3,27E-17 | 0,67561715 | 0,605 | 0,14 | 5,18E-13 | MAC | Myo5a |
| Ppp1r21 | 3,74E-17 | 0,481426419 | 0,519 | 0,083 | 5,93E-13 | MAC | Ppp1r21 |
| Rnf130 | 3,78E-17 | 0,601648237 | 0,765 | 0,285 | 6,00E-13 | MAC | Rnf130 |
| Lcp1 | 3,85E-17 | 0,874187342 | 0,975 | 0,719 | 6,10E-13 | MAC | Lcp1 |
| Irf2bp2 | 3,91E-17 | 0,476537671 | 0,704 | 0,219 | 6,20E-13 | MAC | Irf2bp2 |
| Gusb | 4,24E-17 | 0,940811342 | 0,864 | 0,463 | 6,72E-13 | MAC | Gusb |
| Clta | 4,41E-17 | 0,715709448 | 0,963 | 0,694 | 6,99E-13 | MAC | Clta |
| Emp3 | 5,00E-17 | 0,656386728 | 0,852 | 0,343 | 7,93E-13 | MAC | Emp3 |
| Ccl8 | 5,44E-17 | 1,154760223 | 0,272 | 0 | 8,62E-13 | MAC | Ccl8 |
| Ap1b1 | 6,04E-17 | 0,731000105 | 0,704 | 0,223 | 9,58E-13 | MAC | Ap1b1 |
| B2m | 6,16E-17 | 0,505602121 | 1 | 1 | 9,76E-13 | MAC | B2m |
| 1700017B05Rik | 6,80E-17 | 0,775383546 | 0,556 | 0,124 | 1,08E-12 | MAC | 1700017B05Rik |
| Fam134b | 7,82E-17 | 0,381311619 | 0,333 | 0,017 | 1,24E-12 | MAC | Fam134b |
| 1810011H11Rik | 8,74E-17 | 0,384323324 | 0,284 | 0,004 | 1,39E-12 | MAC | 1810011H11Rik |
| Tnfsf13 | 1,05E-16 | 0,437969132 | 0,383 | 0,037 | 1,67E-12 | MAC | Tnfsf13 |
| Paox | 1,09E-16 | 0,395249477 | 0,432 | 0,062 | 1,72E-12 | MAC | Paox |
| Atf3 | 1,28E-16 | 1,028513801 | 0,63 | 0,165 | 2,03E-12 | MAC | Atf3 |
| Phf11b | 1,43E-16 | 0,608505462 | 0,432 | 0,058 | 2,27E-12 | MAC | Phf11b |
| Erp29 | 1,80E-16 | 0,645234386 | 0,901 | 0,504 | 2,85E-12 | MAC | Erp29 |
| Rap2b | 2,14E-16 | 0,662673498 | 0,864 | 0,533 | 3,40E-12 | MAC | Rap2b |

| | | | | | | | |
|---|---|---|---|---|---|---|---|
| Cd53 | 1,57E-61 | 1,85537111 | 0,988 | 0,045 | 2,49E-57 | 1 | Cd53 |
| Ptpn6 | 2,74E-61 | 1,607331114 | 0,914 | 0,008 | 4,35E-57 | 1 | Ptpn6 |
| Clec4a2 | 4,65E-61 | 1,513908845 | 0,901 | 0,004 | 7,37E-57 | 1 | Clec4a2 |
| Lrp1 | 5,38E-61 | 1,668389423 | 0,914 | 0,008 | 8,53E-57 | 1 | Lrp1 |
| Fermt3 | 6,14E-61 | 1,49298526 | 0,901 | 0,004 | 9,73E-57 | 1 | Fermt3 |
| Csf2ra | 7,19E-61 | 1,317595911 | 0,938 | 0,025 | 1,14E-56 | 1 | Csf2ra |
| Ms4a4a | 1,04E-60 | 1,521905739 | 0,889 | 0 | 1,65E-56 | 1 | Ms4a4a |
| Ccl6 | 1,04E-60 | 1,516975938 | 0,889 | 0 | 1,65E-56 | 1 | Ccl6 |
| Fcgr4 | 1,12E-60 | 1,79610778 | 0,914 | 0,008 | 1,77E-56 | 1 | Fcgr4 |
| Fcer1g | 2,14E-60 | 3,44213885 | 1 | 0,083 | 3,40E-56 | 1 | Fcer1g |
| Adgre1 | 4,91E-60 | 1,58175684 | 0,889 | 0,004 | 7,79E-56 | 1 | Adgre1 |
| Wfdc17 | 8,83E-60 | 2,040118947 | 0,901 | 0,012 | 1,40E-55 | 1 | Wfdc17 |
| Clec12a | 1,07E-59 | 1,508464223 | 0,877 | 0 | 1,69E-55 | 1 | Clec12a |
| Rac2 | 2,29E-59 | 1,650161882 | 0,889 | 0,008 | 3,63E-55 | 1 | Rac2 |
| Fam105a | 5,12E-59 | 1,499573611 | 0,877 | 0,004 | 8,11E-55 | 1 | Fam105a |
| Itgb2 | 1,08E-58 | 1,545172129 | 0,864 | 0 | 1,71E-54 | 1 | Itgb2 |
| Lair1 | 1,08E-58 | 1,427215962 | 0,864 | 0 | 1,71E-54 | 1 | Lair1 |
| Cxcl16 | 2,81E-58 | 2,227180165 | 1 | 0,091 | 4,45E-54 | 1 | Cxcl16 |
| C5ar1 | 5,41E-58 | 1,676653978 | 0,864 | 0,004 | 8,58E-54 | 1 | C5ar1 |
| Clec4n | 1,07E-57 | 1,596936596 | 0,852 | 0 | 1,70E-53 | 1 | Clec4n |
| Ctsc | 3,48E-57 | 2,919063102 | 0,988 | 0,103 | 5,51E-53 | 1 | Ctsc |
| Msr1 | 1,05E-56 | 1,516472413 | 0,84 | 0 | 1,67E-52 | 1 | Msr1 |
| Slamf9 | 1,05E-56 | 1,434402142 | 0,84 | 0 | 1,67E-52 | 1 | Slamf9 |
| Arrb2 | 1,37E-56 | 0,863955106 | 0,901 | 0,029 | 2,17E-52 | 1 | Arrb2 |
| Fcgr1 | 5,30E-56 | 1,453640018 | 0,84 | 0,004 | 8,40E-52 | 1 | Fcgr1 |
| Cd300lf | 6,17E-56 | 1,405426153 | 0,84 | 0,004 | 9,78E-52 | 1 | Cd300lf |
| Axl | 8,64E-56 | 1,375333071 | 0,889 | 0,021 | 1,37E-51 | 1 | Axl |
| Acp5 | 1,54E-55 | 1,861950181 | 0,926 | 0,05 | 2,45E-51 | 1 | Acp5 |
| Fcrls | 5,54E-55 | 1,99176459 | 0,827 | 0,004 | 8,78E-51 | 1 | Fcrls |
| Plek | 6,64E-55 | 1,260735668 | 0,827 | 0,004 | 1,05E-50 | 1 | Plek |
| Nckap1l | 7,65E-55 | 1,31726559 | 0,864 | 0,017 | 1,21E-50 | 1 | Nckap1l |
| Cx3cr1 | 9,74E-55 | 1,825864762 | 0,815 | 0 | 1,54E-50 | 1 | Cx3cr1 |
| Sla | 9,74E-55 | 1,434527979 | 0,815 | 0 | 1,54E-50 | 1 | Sla |
| Mrc1 | 1,75E-54 | 1,504749087 | 0,852 | 0,017 | 2,77E-50 | 1 | Mrc1 |
| Cd33 | 3,53E-54 | 1,347542742 | 0,852 | 0,017 | 5,59E-50 | 1 | Cd33 |
| Selplg | 5,30E-54 | 1,472375391 | 0,84 | 0,017 | 8,41E-50 | 1 | Selplg |
| Ccr5 | 2,44E-53 | 1,4336233 | 0,84 | 0,021 | 3,87E-49 | 1 | Ccr5 |
| Tlr13 | 9,56E-53 | 1,074220537 | 0,827 | 0,012 | 1,52E-48 | 1 | Tlr13 |
| Lpcat2 | 3,41E-52 | 1,535071916 | 0,852 | 0,025 | 5,40E-48 | 1 | Lpcat2 |
| Hck | 4,57E-52 | 1,217824949 | 0,79 | 0,004 | 7,24E-48 | 1 | Hck |
| Ccr1 | 5,34E-52 | 1,699117124 | 0,815 | 0,017 | 8,47E-48 | 1 | Ccr1 |
| Evi2a | 8,08E-52 | 1,302358918 | 0,852 | 0,033 | 1,28E-47 | 1 | Evi2a |
| H2-DMa | 1,01E-51 | 1,603167016 | 0,852 | 0,029 | 1,60E-47 | 1 | H2-DMa |
| Ms4a6b | 1,44E-51 | 1,698284467 | 0,864 | 0,037 | 2,29E-47 | 1 | Ms4a6b |
| Mt1 | 2,78E-51 | 1,720681994 | 0,975 | 0,132 | 4,41E-47 | 1 | Mt1 |
| Cd84 | 2,98E-51 | 1,193123543 | 0,827 | 0,025 | 4,73E-47 | 1 | Cd84 |
| Ccl3 | 7,07E-51 | 2,044218471 | 0,765 | 0 | 1,12E-46 | 1 | Ccl3 |
| Slc11a1 | 7,07E-51 | 1,510747551 | 0,765 | 0 | 1,12E-46 | 1 | Slc11a1 |
| Ncf4 | 7,07E-51 | 1,170621712 | 0,765 | 0 | 1,12E-46 | 1 | Ncf4 |
| Gpr65 | 7,07E-51 | 1,131568577 | 0,765 | 0 | 1,12E-46 | 1 | Gpr65 |
| Apoc2 | 7,07E-51 | 1,087753959 | 0,765 | 0 | 1,12E-46 | 1 | Apoc2 |
| Cd86 | 7,07E-51 | 1,022165243 | 0,765 | 0 | 1,12E-46 | 1 | Cd86 |
| Vav1 | 7,07E-51 | 1,008406534 | 0,765 | 0 | 1,12E-46 | 1 | Vav1 |
| Itgb5 | 1,85E-50 | 1,33655926 | 0,79 | 0,012 | 2,94E-46 | 1 | Itgb5 |
| Clec4a3 | 6,31E-50 | 1,132583469 | 0,753 | 0 | 9,99E-46 | 1 | Clec4a3 |
| Bcl2a1d | 6,31E-50 | 0,96999313 | 0,753 | 0 | 9,99E-46 | 1 | Bcl2a1d |
| Hpgds | 9,53E-50 | 0,968983948 | 0,778 | 0,008 | 1,51E-45 | 1 | Hpgds |
| Ncf1 | 1,10E-49 | 1,076307045 | 0,778 | 0,008 | 1,75E-45 | 1 | Ncf1 |
| Gm2a | 1,11E-49 | 1,823513617 | 1 | 0,219 | 1,75E-45 | 1 | Gm2a |
| Ccl2 | 2,04E-49 | 1,96990662 | 0,864 | 0,041 | 3,23E-45 | 1 | Ccl2 |
| Ccl4 | 2,05E-49 | 2,292775707 | 0,765 | 0,008 | 3,25E-45 | 1 | Ccl4 |
| Cd48 | 2,05E-49 | 1,052723344 | 0,765 | 0,008 | 3,25E-45 | 1 | Cd48 |
| Bin2 | 2,83E-49 | 1,287063192 | 0,765 | 0,008 | 4,48E-45 | 1 | Bin2 |
| Cd72 | 4,89E-49 | 1,826717892 | 0,827 | 0,037 | 7,75E-45 | 1 | Cd72 |
| Plbd1 | 5,55E-49 | 1,632109157 | 0,741 | 0 | 8,80E-45 | 1 | Plbd1 |
| Fgd2 | 5,55E-49 | 1,318948008 | 0,741 | 0 | 8,80E-45 | 1 | Fgd2 |
| Aoah | 5,55E-49 | 0,989179868 | 0,741 | 0 | 8,80E-45 | 1 | Aoah |
| Apoe | 1,06E-48 | 5,76904748 | 1 | 0,281 | 1,67E-44 | 1 | Apoe |
| Klra2 | 4,82E-48 | 1,07273413 | 0,728 | 0 | 7,65E-44 | 1 | Klra2 |
| Prune2 | 5,85E-48 | 1,135055499 | 0,84 | 0,041 | 9,27E-44 | 1 | Prune2 |
| Irf5 | 1,08E-47 | 1,501959508 | 0,877 | 0,091 | 1,72E-43 | 1 | Irf5 |
| Ifi207 | 2,25E-47 | 1,035036117 | 0,753 | 0,012 | 3,56E-43 | 1 | Ifi207 |
| Bcl2a1a | 4,14E-47 | 1,005691984 | 0,716 | 0 | 6,56E-43 | 1 | Bcl2a1a |
| Tmem119 | 4,60E-47 | 1,483698869 | 0,728 | 0,004 | 7,29E-43 | 1 | Tmem119 |
| Arhgap45 | 3,50E-46 | 1,004643969 | 0,704 | 0 | 5,55E-42 | 1 | Arhgap45 |
| Tbxas1 | 4,84E-46 | 1,013880169 | 0,728 | 0,008 | 7,67E-42 | 1 | Tbxas1 |
| Sirpa | 6,66E-46 | 1,643392497 | 0,963 | 0,169 | 1,06E-41 | 1 | Sirpa |
| Ctsh | 9,56E-46 | 2,001194247 | 0,988 | 0,252 | 1,51E-41 | 1 | Ctsh |
| Dpep2 | 1,42E-45 | 1,043355863 | 0,716 | 0,008 | 2,26E-41 | 1 | Dpep2 |
| Pilra | 2,15E-45 | 1,017843766 | 0,704 | 0,004 | 3,40E-41 | 1 | Pilra |
| Clec4a1 | 2,93E-45 | 1,224646934 | 0,691 | 0 | 4,64E-41 | 1 | Clec4a1 |
| Pid1 | 2,93E-45 | 1,043526642 | 0,691 | 0 | 4,64E-41 | 1 | Pid1 |
| Tifab | 2,93E-45 | 0,893903155 | 0,691 | 0 | 4,64E-41 | 1 | Tifab |
| Pirb | 3,32E-45 | 1,387346463 | 0,741 | 0,021 | 5,26E-41 | 1 | Pirb |
| Lst1 | 4,33E-45 | 0,643469085 | 0,728 | 0,017 | 6,86E-41 | 1 | Lst1 |
| Lgals3 | 8,65E-45 | 2,203308785 | 0,864 | 0,099 | 1,37E-40 | 1 | Lgals3 |
| Pf4 | 1,58E-44 | 1,235653998 | 0,753 | 0,029 | 2,50E-40 | 1 | Pf4 |
| Cd14 | 2,50E-44 | 1,973991269 | 0,938 | 0,145 | 3,97E-40 | 1 | Cd14 |
| Zmynd15 | 1,06E-43 | 1,512281879 | 0,704 | 0,012 | 1,68E-39 | 1 | Zmynd15 |
| Pycard | 1,10E-43 | 1,183325971 | 0,864 | 0,087 | 1,74E-39 | 1 | Pycard |
| AU020206 | 1,51E-43 | 1,118438958 | 0,951 | 0,178 | 2,39E-39 | 1 | AU020206 |
| Efhd2 | 1,74E-43 | 1,550336541 | 0,963 | 0,219 | 2,75E-39 | 1 | Efhd2 |
| Sirpb1b | 1,97E-43 | 0,458000508 | 0,667 | 0 | 3,12E-39 | 1 | Sirpb1b |
| Rgs2 | 2,44E-43 | 1,650651099 | 0,951 | 0,161 | 3,87E-39 | 1 | Rgs2 |
| Tlr2 | 4,10E-43 | 1,300151531 | 0,765 | 0,037 | 6,50E-39 | 1 | Tlr2 |
| Unc93b1 | 5,60E-43 | 1,83824491 | 1 | 0,351 | 8,88E-39 | 1 | Unc93b1 |
| H2-Aa | 6,01E-43 | 2,966786186 | 0,827 | 0,083 | 9,53E-39 | 1 | H2-Aa |
| P2ry6 | 7,69E-43 | 1,356527043 | 0,827 | 0,07 | 1,22E-38 | 1 | P2ry6 |
| Gatm | 2,41E-42 | 1,559059904 | 0,926 | 0,157 | 3,82E-38 | 1 | Gatm |
| Ctsz | 2,54E-42 | 2,09688997 | 1 | 0,413 | 4,02E-38 | 1 | Ctsz |
| Ccl7 | 4,74E-42 | 2,092756852 | 0,679 | 0,012 | 7,52E-38 | 1 | Ccl7 |
| Cd83 | 1,26E-41 | 1,366010777 | 0,642 | 0 | 1,99E-37 | 1 | Cd83 |
| Itgal | 1,26E-41 | 0,788248745 | 0,642 | 0 | 1,99E-37 | 1 | Itgal |
| Ucp2 | 1,35E-41 | 2,064485573 | 1 | 0,405 | 2,14E-37 | 1 | Ucp2 |
| Ntpcr | 3,70E-41 | 1,495518791 | 0,901 | 0,161 | 5,86E-37 | 1 | Ntpcr |
| Cst3 | 4,35E-41 | 1,849523353 | 1 | 0,851 | 6,89E-37 | 1 | Cst3 |
| Zeb2 | 6,25E-41 | 0,941150684 | 0,741 | 0,033 | 9,91E-37 | 1 | Zeb2 |
| Ccl9 | 8,03E-41 | 1,141040972 | 0,642 | 0,004 | 1,27E-36 | 1 | Ccl9 |
| Myo1f | 9,85E-41 | 0,881048734 | 0,63 | 0 | 1,56E-36 | 1 | Myo1f |
| Rbm47 | 9,89E-41 | 0,388600047 | 0,704 | 0,025 | 1,57E-36 | 1 | Rbm47 |
| Hexb | 1,01E-40 | 1,958017343 | 0,988 | 0,298 | 1,60E-36 | 1 | Hexb |
| Grn | 1,61E-40 | 1,850319544 | 1 | 0,773 | 2,55E-36 | 1 | Grn |
| Gm42031 | 2,09E-40 | 0,618574808 | 0,914 | 0,198 | 3,31E-36 | 1 | Gm42031 |
| Gm6377 | 2,52E-40 | 1,042744529 | 0,642 | 0,004 | 3,99E-36 | 1 | Gm6377 |
| Celf2 | 2,62E-40 | 0,88320141 | 0,654 | 0,008 | 4,15E-36 | 1 | Celf2 |
| Ncf2 | 3,99E-40 | 0,968890726 | 0,691 | 0,021 | 6,32E-36 | 1 | Ncf2 |
| Fth1 | 4,75E-40 | 1,914775366 | 1 | 0,988 | 7,54E-36 | 1 | Fth1 |
| Tnfaip8l2 | 7,36E-40 | 0,958776097 | 0,63 | 0,004 | 1,17E-35 | 1 | Tnfaip8l2 |
| Ftl1 | 7,51E-40 | 1,614515569 | 1 | 0,992 | 1,19E-35 | 1 | Ftl1 |
| Spint1 | 7,63E-40 | 0,95144666 | 0,617 | 0 | 1,21E-35 | 1 | Spint1 |

| | | | | | | | |
|---|---|---|---|---|---|---|---|
| Nampt | 2,41E-16 | 0,727924815 | 0,716 | 0,231 | 3,82E-12 | MAC | Nampt |
| Ckb | 2,65E-16 | 0,919194369 | 0,778 | 0,326 | 4,20E-12 | MAC | Ckb |
| Skap2 | 2,76E-16 | 0,622818388 | 0,765 | 0,277 | 4,38E-12 | MAC | Skap2 |
| Il1rn | 3,05E-16 | 0,711378733 | 0,259 | 0 | 4,84E-12 | MAC | Il1rn |
| Trpm2 | 3,05E-16 | 0,544577763 | 0,259 | 0 | 4,84E-12 | MAC | Trpm2 |
| Cd244 | 3,05E-16 | 0,427697384 | 0,259 | 0 | 4,84E-12 | MAC | Cd244 |
| Psd4 | 3,05E-16 | 0,409417958 | 0,259 | 0 | 4,84E-12 | MAC | Psd4 |
| Vill | 3,05E-16 | 0,379542702 | 0,259 | 0 | 4,84E-12 | MAC | Vill |
| Rnf150 | 3,05E-16 | 0,253934024 | 0,259 | 0 | 4,84E-12 | MAC | Rnf150 |
| Hcst | 4,27E-16 | 0,302186009 | 0,481 | 0,083 | 6,77E-12 | MAC | Hcst |
| Ccr2 | 4,87E-16 | 0,684310477 | 0,272 | 0,004 | 7,72E-12 | MAC | Ccr2 |
| Mknk1 | 5,36E-16 | 0,538016491 | 0,568 | 0,132 | 8,50E-12 | MAC | Mknk1 |
| Sdf2l1 | 5,96E-16 | 0,510732026 | 0,704 | 0,207 | 9,45E-12 | MAC | Sdf2l1 |
| Lrch4 | 6,16E-16 | 0,643674174 | 0,63 | 0,182 | 9,76E-12 | MAC | Lrch4 |
| Atp8a1 | 6,27E-16 | 0,343750138 | 0,556 | 0,112 | 9,94E-12 | MAC | Atp8a1 |
| Rpl10-ps3 | 6,71E-16 | 0,503409129 | 1 | 0,921 | 1,06E-11 | MAC | Rpl10-ps3 |
| Trpv2 | 7,28E-16 | 0,61375328 | 0,407 | 0,058 | 1,15E-11 | MAC | Trpv2 |
| Epsti1 | 7,57E-16 | 0,469595964 | 0,407 | 0,05 | 1,20E-11 | MAC | Epsti1 |
| Cln8 | 7,70E-16 | 0,563221072 | 0,531 | 0,103 | 1,22E-11 | MAC | Cln8 |
| Plxnb2 | 8,64E-16 | 0,8664309 | 0,753 | 0,293 | 1,37E-11 | MAC | Plxnb2 |
| N4bp2l1 | 1,11E-15 | 0,400397317 | 0,284 | 0,008 | 1,76E-11 | MAC | N4bp2l1 |
| Gm29216 | 1,17E-15 | 0,542367253 | 1 | 1 | 1,85E-11 | MAC | Gm29216 |
| Sgpl1 | 1,19E-15 | 0,657117666 | 0,667 | 0,215 | 1,89E-11 | MAC | Sgpl1 |
| Creb5 | 1,21E-15 | 0,382026432 | 0,383 | 0,041 | 1,92E-11 | MAC | Creb5 |
| Inpp5d | 1,23E-15 | 0,741392624 | 0,901 | 0,517 | 1,96E-11 | MAC | Inpp5d |
| Rab20 | 1,38E-15 | 0,685717148 | 0,42 | 0,058 | 2,18E-11 | MAC | Rab20 |
| Pgd | 1,52E-15 | 0,524444206 | 0,802 | 0,306 | 2,41E-11 | MAC | Pgd |
| Ldhb | 1,61E-15 | 0,565261504 | 0,358 | 0,033 | 2,56E-11 | MAC | Ldhb |
| Sp110 | 2,14E-15 | 0,575952333 | 0,654 | 0,182 | 3,39E-11 | MAC | Sp110 |
| Sh3bp2 | 2,29E-15 | 0,313881132 | 0,481 | 0,083 | 3,63E-11 | MAC | Sh3bp2 |
| Gm5150 | 2,75E-15 | 0,333877016 | 0,259 | 0,004 | 4,37E-11 | MAC | Gm5150 |
| Gdf3 | 3,94E-15 | 0,841966805 | 0,296 | 0,017 | 6,24E-11 | MAC | Gdf3 |
| Hk3 | 4,12E-15 | 0,359976259 | 0,259 | 0,004 | 6,53E-11 | MAC | Hk3 |
| Spon1 | 5,08E-15 | 0,369231045 | 0,272 | 0,008 | 8,05E-11 | MAC | Spon1 |
| Iqgap2 | 5,20E-15 | 0,300328243 | 0,272 | 0,008 | 8,24E-11 | MAC | Iqgap2 |
| Fgd4 | 5,36E-15 | 0,253209369 | 0,321 | 0,025 | 8,50E-11 | MAC | Fgd4 |
| Syk | 5,56E-15 | 0,393628744 | 0,506 | 0,103 | 8,81E-11 | MAC | Syk |
| Slc29a3 | 6,02E-15 | 0,563428806 | 0,444 | 0,074 | 9,55E-11 | MAC | Slc29a3 |
| Dhrs3 | 6,22E-15 | 0,742196147 | 0,741 | 0,298 | 9,86E-11 | MAC | Dhrs3 |
| Usp2 | 7,53E-15 | 0,369816958 | 0,296 | 0,017 | 1,19E-10 | MAC | Usp2 |
| Grk2 | 8,91E-15 | 0,644933194 | 0,765 | 0,318 | 1,41E-10 | MAC | Grk2 |
| Fam46a | 9,34E-15 | 0,452056672 | 0,346 | 0,033 | 1,48E-10 | MAC | Fam46a |
| Mt2 | 1,00E-14 | 0,32885324 | 0,432 | 0,066 | 1,58E-10 | MAC | Mt2 |
| Cd37 | 1,22E-14 | 0,256594967 | 0,284 | 0,012 | 1,94E-10 | MAC | Cd37 |
| Rps9 | 1,26E-14 | 0,463912496 | 1 | 0,959 | 2,00E-10 | MAC | Rps9 |
| Sh3bgrl3 | 1,33E-14 | 0,511549595 | 0,988 | 0,955 | 2,10E-10 | MAC | Sh3bgrl3 |
| Rplp2 | 1,59E-14 | 0,43557302 | 1 | 0,971 | 2,51E-10 | MAC | Rplp2 |
| Plk3 | 1,72E-14 | 0,732739522 | 0,481 | 0,103 | 2,73E-10 | MAC | Plk3 |
| Hebp1 | 1,89E-14 | 0,510041106 | 0,679 | 0,236 | 2,99E-10 | MAC | Hebp1 |
| Arpc2 | 2,18E-14 | 0,538401095 | 1 | 0,934 | 3,45E-10 | MAC | Arpc2 |
| Cfb | 2,25E-14 | 0,91965093 | 0,63 | 0,186 | 3,56E-10 | MAC | Cfb |
| Lamtor4 | 2,60E-14 | 0,436108828 | 0,741 | 0,31 | 4,12E-10 | MAC | Lamtor4 |
| Adamtsl4 | 2,73E-14 | 0,494964011 | 0,284 | 0,017 | 4,33E-10 | MAC | Adamtsl4 |
| Nmt1 | 3,00E-14 | 0,511115036 | 0,938 | 0,595 | 4,75E-10 | MAC | Nmt1 |
| Grina | 3,37E-14 | 0,765950615 | 0,79 | 0,372 | 5,34E-10 | MAC | Grina |
| Rbpj | 3,43E-14 | 0,578363677 | 0,704 | 0,264 | 5,43E-10 | MAC | Rbpj |
| Irf8 | 4,52E-14 | 0,626553181 | 0,815 | 0,355 | 7,16E-10 | MAC | Irf8 |
| Limd2 | 4,62E-14 | 0,614563961 | 0,765 | 0,326 | 7,32E-10 | MAC | Limd2 |
| Dusp1 | 5,08E-14 | 1,103805735 | 0,877 | 0,744 | 8,06E-10 | MAC | Dusp1 |
| Il18bp | 5,36E-14 | 0,590276155 | 0,481 | 0,099 | 8,50E-10 | MAC | Il18bp |
| Ier3 | 5,78E-14 | 1,028140617 | 0,778 | 0,397 | 9,16E-10 | MAC | Ier3 |
| Stx7 | 7,18E-14 | 0,695742817 | 0,827 | 0,421 | 1,14E-09 | MAC | Stx7 |
| Cotl1 | 7,39E-14 | 0,565070912 | 0,951 | 0,57 | 1,17E-09 | MAC | Cotl1 |
| Rpl35a | 7,44E-14 | 0,460085251 | 1 | 0,955 | 1,18E-09 | MAC | Rpl35a |
| Adrb2 | 8,23E-14 | 0,690821824 | 0,321 | 0,029 | 1,31E-09 | MAC | Adrb2 |
| Fosb | 8,49E-14 | 0,50847215 | 0,543 | 0,132 | 1,35E-09 | MAC | Fosb |
| Rnf149 | 8,77E-14 | 0,625950326 | 0,469 | 0,103 | 1,39E-09 | MAC | Rnf149 |
| Rasa4 | 1,02E-13 | 0,491052565 | 0,358 | 0,045 | 1,61E-09 | MAC | Rasa4 |
| Aldh2 | 1,13E-13 | 0,528215229 | 0,432 | 0,079 | 1,79E-09 | MAC | Aldh2 |
| Ogfrl1 | 1,30E-13 | 0,402116622 | 0,531 | 0,132 | 2,05E-09 | MAC | Ogfrl1 |
| Stk4 | 1,72E-13 | 0,539395855 | 0,481 | 0,112 | 2,73E-09 | MAC | Stk4 |
| Rel | 1,81E-13 | 0,602543455 | 0,494 | 0,12 | 2,86E-09 | MAC | Rel |
| G530011O06Rik | 1,82E-13 | 0,402739018 | 0,321 | 0,033 | 2,89E-09 | MAC | G530011O06Rik |
| Snx24 | 1,85E-13 | 0,356335862 | 0,407 | 0,066 | 2,94E-09 | MAC | Snx24 |
| Oxct1 | 2,11E-13 | 0,52743885 | 0,519 | 0,132 | 3,35E-09 | MAC | Oxct1 |
| Renbp | 2,74E-13 | 0,593152122 | 0,568 | 0,165 | 4,34E-09 | MAC | Renbp |
| Stab1 | 3,44E-13 | 0,808652468 | 0,938 | 0,897 | 5,46E-09 | MAC | Stab1 |
| Rnaset2b | 3,62E-13 | 0,578768571 | 0,988 | 0,744 | 5,74E-09 | MAC | Rnaset2b |
| Gna15 | 3,73E-13 | 0,443882291 | 0,272 | 0,017 | 5,91E-09 | MAC | Gna15 |
| S100a1 | 3,85E-13 | 0,385697876 | 0,568 | 0,165 | 6,10E-09 | MAC | S100a1 |
| Mgst1 | 4,13E-13 | 0,544946181 | 0,42 | 0,079 | 6,55E-09 | MAC | Mgst1 |
| Nfe2l2 | 4,78E-13 | 0,775013045 | 0,889 | 0,529 | 7,58E-09 | MAC | Nfe2l2 |
| Extl3 | 6,66E-13 | 0,53124816 | 0,691 | 0,256 | 1,06E-08 | MAC | Extl3 |
| Lhfpl2 | 6,77E-13 | 0,627004965 | 0,358 | 0,054 | 1,07E-08 | MAC | Lhfpl2 |
| Dennd4b | 8,28E-13 | 0,473687598 | 0,272 | 0,021 | 1,31E-08 | MAC | Dennd4b |
| Hfe | 8,72E-13 | 0,613687222 | 0,63 | 0,227 | 1,38E-08 | MAC | Hfe |
| Mkrn1 | 9,01E-13 | 0,437200793 | 0,63 | 0,207 | 1,43E-08 | MAC | Mkrn1 |
| Cd302 | 1,12E-12 | 0,557097882 | 0,704 | 0,331 | 1,78E-08 | MAC | Cd302 |
| Zfp36l2 | 1,24E-12 | 0,510718524 | 0,827 | 0,455 | 1,97E-08 | MAC | Zfp36l2 |
| Coro7 | 1,32E-12 | 0,606731924 | 0,716 | 0,302 | 2,09E-08 | MAC | Coro7 |
| Plagl2 | 1,35E-12 | 0,47316562 | 0,432 | 0,091 | 2,15E-08 | MAC | Plagl2 |
| Rps25 | 1,36E-12 | 0,436130949 | 1 | 0,963 | 2,16E-08 | MAC | Rps25 |
| Ezr | 1,43E-12 | 0,444466913 | 0,346 | 0,05 | 2,27E-08 | MAC | Ezr |
| Pfn1 | 1,44E-12 | 0,391110037 | 1 | 0,996 | 2,29E-08 | MAC | Pfn1 |
| Pld3 | 1,46E-12 | 0,651098261 | 0,765 | 0,331 | 2,31E-08 | MAC | Pld3 |
| Ptpn1 | 1,57E-12 | 0,648909454 | 0,827 | 0,45 | 2,49E-08 | MAC | Ptpn1 |
| 1600014C10Rik | 1,94E-12 | 0,435332021 | 0,494 | 0,124 | 3,08E-08 | MAC | 1600014C10Rik |
| Frrs1 | 2,23E-12 | 0,436794436 | 0,358 | 0,054 | 3,53E-08 | MAC | Frrs1 |
| mt-Nd6 | 2,43E-12 | 0,495943928 | 1 | 0,938 | 3,85E-08 | MAC | mt-Nd6 |
| Tmem50b | 2,45E-12 | 0,615273094 | 0,556 | 0,169 | 3,88E-08 | MAC | Tmem50b |
| Mfsd12 | 2,99E-12 | 0,317135005 | 0,395 | 0,074 | 4,74E-08 | MAC | Mfsd12 |
| Dapp1 | 3,27E-12 | 0,461333746 | 0,321 | 0,041 | 5,19E-08 | MAC | Dapp1 |
| Dnmt3a | 3,77E-12 | 0,399229807 | 0,543 | 0,157 | 5,97E-08 | MAC | Dnmt3a |
| Rpl37rt | 3,82E-12 | 0,328749956 | 1 | 0,967 | 6,05E-08 | MAC | Rpl37rt |
| Atp6v1f | 3,93E-12 | 0,52629993 | 0,951 | 0,702 | 6,24E-08 | MAC | Atp6v1f |
| Etv5 | 4,01E-12 | 0,562947656 | 0,469 | 0,12 | 6,36E-08 | MAC | Etv5 |
| Pi4k2a | 4,14E-12 | 0,440058705 | 0,383 | 0,07 | 6,56E-08 | MAC | Pi4k2a |
| Atp6v0b | 4,44E-12 | 0,627465194 | 0,827 | 0,467 | 7,05E-08 | MAC | Atp6v0b |
| Epb41l2 | 4,84E-12 | 0,69486121 | 0,765 | 0,384 | 7,68E-08 | MAC | Epb41l2 |
| Fam213b | 5,37E-12 | 0,335481219 | 0,296 | 0,033 | 8,52E-08 | MAC | Fam213b |
| Vamp8 | 5,43E-12 | 0,60299128 | 0,926 | 0,711 | 8,61E-08 | MAC | Vamp8 |
| Gk | 5,86E-12 | 0,319990012 | 0,284 | 0,029 | 9,28E-08 | MAC | Gk |
| Snx8 | 5,97E-12 | 0,343920183 | 0,346 | 0,054 | 9,47E-08 | MAC | Snx8 |
| Klc4 | 6,54E-12 | 0,485411731 | 0,469 | 0,116 | 1,04E-07 | MAC | Klc4 |
| Fam96a | 8,14E-12 | 0,582269561 | 0,815 | 0,434 | 1,29E-07 | MAC | Fam96a |
| Lrp12 | 8,61E-12 | 0,287319276 | 0,259 | 0,021 | 1,36E-07 | MAC | Lrp12 |
| B3galnt1 | 1,06E-11 | 0,353528189 | 0,284 | 0,029 | 1,68E-07 | MAC | B3galnt1 |
| Mroh1 | 1,16E-11 | 0,370345061 | 0,358 | 0,062 | 1,83E-07 | MAC | Mroh1 |
| Fam234b | 1,23E-11 | 0,347375668 | 0,321 | 0,045 | 1,96E-07 | MAC | Fam234b |
| Samhd1 | 1,32E-11 | 0,577276734 | 0,679 | 0,264 | 2,09E-07 | MAC | Samhd1 |

| | | | | | | | |
|---|---|---|---|---|---|---|---|
| Il10ra | 7,63E-40 | 0,902622275 | 0,617 | 0 | 1,21E-35 | 1 | Il10ra |
| Lrrc25 | 7,63E-40 | 0,673458635 | 0,617 | 0 | 1,21E-35 | 1 | Lrrc25 |
| Emb | 7,95E-40 | 1,216127566 | 0,642 | 0,008 | 1,26E-35 | 1 | Emb |
| Cd74 | 1,34E-39 | 3,250433361 | 0,951 | 0,24 | 2,12E-35 | 1 | Cd74 |
| Trf | 2,10E-39 | 2,137320354 | 0,963 | 0,326 | 3,32E-35 | 1 | Trf |
| Ppfia4 | 5,01E-39 | 0,953528514 | 0,617 | 0,004 | 7,94E-35 | 1 | Ppfia4 |
| Bhlhe41 | 5,75E-39 | 0,793743502 | 0,667 | 0,017 | 9,12E-35 | 1 | Bhlhe41 |
| Arhgap30 | 5,84E-39 | 0,740906224 | 0,605 | 0 | 9,26E-35 | 1 | Arhgap30 |
| Lat2 | 7,18E-39 | 1,045474674 | 0,728 | 0,045 | 1,14E-34 | 1 | Lat2 |
| Cfp | 8,85E-39 | 0,997574157 | 0,617 | 0,004 | 1,40E-34 | 1 | Cfp |
| Prkcd | 1,40E-38 | 1,198810565 | 0,765 | 0,066 | 2,21E-34 | 1 | Prkcd |
| Lgmn | 1,49E-38 | 1,686533885 | 1 | 0,897 | 2,35E-34 | 1 | Lgmn |
| Rab3il1 | 2,08E-38 | 1,270197707 | 0,802 | 0,083 | 3,30E-34 | 1 | Rab3il1 |
| Cebpb | 2,44E-38 | 1,020953092 | 0,802 | 0,083 | 3,88E-34 | 1 | Cebpb |
| Man2b1 | 2,71E-38 | 1,68834718 | 0,988 | 0,401 | 4,29E-34 | 1 | Man2b1 |
| Dock10 | 3,89E-38 | 0,670253041 | 0,605 | 0,004 | 6,16E-34 | 1 | Dock10 |
| Hk2 | 3,89E-38 | 0,537198907 | 0,605 | 0,004 | 6,16E-34 | 1 | Hk2 |
| Ifi27l2a | 4,05E-38 | 0,863232196 | 0,617 | 0,008 | 6,42E-34 | 1 | Ifi27l2a |
| Olfml3 | 4,07E-38 | 0,944676953 | 0,642 | 0,017 | 6,46E-34 | 1 | Olfml3 |
| Cyba | 6,51E-38 | 1,447746936 | 1 | 0,736 | 1,03E-33 | 1 | Cyba |
| Cebpa | 6,91E-38 | 0,678538118 | 0,63 | 0,012 | 1,10E-33 | 1 | Cebpa |
| Cep85 | 7,38E-38 | 1,125955149 | 0,951 | 0,223 | 1,17E-33 | 1 | Cep85 |
| H2-Eb1 | 7,56E-38 | 2,820339309 | 0,778 | 0,091 | 1,20E-33 | 1 | H2-Eb1 |
| Psap | 9,09E-38 | 1,816068891 | 1 | 0,74 | 1,44E-33 | 1 | Psap |
| Tgfbi | 1,25E-37 | 1,886260686 | 0,975 | 0,285 | 1,98E-33 | 1 | Tgfbi |
| Sdc4 | 1,61E-37 | 1,174851714 | 0,877 | 0,136 | 2,55E-33 | 1 | Sdc4 |
| Ccl12 | 3,29E-37 | 1,327348185 | 0,58 | 0 | 5,22E-33 | 1 | Ccl12 |
| Tlr7 | 3,29E-37 | 0,571550199 | 0,58 | 0 | 5,22E-33 | 1 | Tlr7 |
| Sirpb1a | 3,29E-37 | 0,483691495 | 0,58 | 0 | 5,22E-33 | 1 | Sirpb1a |
| Lpxn | 4,54E-37 | 0,945628138 | 0,593 | 0,004 | 7,20E-33 | 1 | Lpxn |
| Arhgdib | 1,06E-36 | 1,5222498 | 0,951 | 0,322 | 1,69E-32 | 1 | Arhgdib |
| Maf | 1,49E-36 | 0,856992054 | 0,852 | 0,128 | 2,37E-32 | 1 | Maf |
| Adcy7 | 1,70E-36 | 0,839789888 | 0,617 | 0,017 | 2,69E-32 | 1 | Adcy7 |
| Ftl1-ps1 | 1,77E-36 | 0,797047456 | 0,988 | 0,913 | 2,81E-32 | 1 | Ftl1-ps1 |
| Clec7a | 2,25E-36 | 0,963189716 | 0,58 | 0,004 | 3,57E-32 | 1 | Clec7a |
| Rab7b | 2,38E-36 | 0,953264266 | 0,58 | 0,004 | 3,77E-32 | 1 | Rab7b |
| Gsn | 5,04E-36 | 1,154446012 | 0,765 | 0,083 | 7,99E-32 | 1 | Gsn |
| Slc7a8 | 5,85E-36 | 0,671393786 | 0,58 | 0,004 | 9,27E-32 | 1 | Slc7a8 |
| Vsir | 8,31E-36 | 1,17055103 | 0,901 | 0,182 | 1,32E-31 | 1 | Vsir |
| Mafb | 8,90E-36 | 1,590650052 | 0,877 | 0,198 | 1,41E-31 | 1 | Mafb |
| Slc15a3 | 1,27E-35 | 0,760427349 | 0,741 | 0,062 | 2,02E-31 | 1 | Slc15a3 |
| Stxbp2 | 1,54E-35 | 0,791656462 | 0,605 | 0,017 | 2,44E-31 | 1 | Stxbp2 |
| Gm43549 | 1,67E-35 | 0,377947293 | 0,605 | 0,017 | 2,65E-31 | 1 | Gm43549 |
| Themis2 | 1,73E-35 | 0,640242425 | 0,568 | 0,004 | 2,74E-31 | 1 | Themis2 |
| Evi2b | 3,66E-35 | 0,626299369 | 0,593 | 0,012 | 5,81E-31 | 1 | Evi2b |
| Gm43305 | 6,48E-35 | 0,828603696 | 0,988 | 0,368 | 1,03E-30 | 1 | Gm43305 |
| Tmem106a | 1,08E-34 | 1,065465683 | 0,778 | 0,103 | 1,72E-30 | 1 | Tmem106a |
| March1 | 1,27E-34 | 0,693771947 | 0,543 | 0 | 2,02E-30 | 1 | March1 |
| Adap2 | 1,27E-34 | 0,421789164 | 0,543 | 0 | 2,02E-30 | 1 | Adap2 |
| Parvg | 1,43E-34 | 0,797120824 | 0,556 | 0,004 | 2,26E-30 | 1 | Parvg |
| Dock2 | 1,47E-34 | 0,745931628 | 0,556 | 0,004 | 2,33E-30 | 1 | Dock2 |
| Ctsb | 1,63E-34 | 1,277189877 | 1 | 0,975 | 2,59E-30 | 1 | Ctsb |
| Rab32 | 2,55E-34 | 1,038452288 | 0,704 | 0,066 | 4,04E-30 | 1 | Rab32 |
| Npc2 | 2,66E-34 | 1,352299046 | 1 | 0,636 | 4,21E-30 | 1 | Npc2 |
| Prdx5 | 2,91E-34 | 1,374611256 | 0,988 | 0,607 | 4,62E-30 | 1 | Prdx5 |
| Ptprj | 3,68E-34 | 0,500222645 | 0,568 | 0,008 | 5,84E-30 | 1 | Ptprj |
| Pkib | 9,07E-34 | 0,616131471 | 0,531 | 0 | 1,44E-29 | 1 | Pkib |
| Igsf6 | 9,71E-34 | 0,887651126 | 0,556 | 0,008 | 1,54E-29 | 1 | Igsf6 |
| AB124611 | 1,10E-33 | 0,690231993 | 0,543 | 0,004 | 1,74E-29 | 1 | AB124611 |
| Snx5 | 1,11E-33 | 1,439090225 | 0,963 | 0,401 | 1,75E-29 | 1 | Snx5 |
| Ctsa | 2,05E-33 | 1,413419045 | 1 | 0,682 | 3,25E-29 | 1 | Ctsa |
| Lgals1 | 3,49E-33 | 1,223815921 | 0,988 | 0,426 | 5,54E-29 | 1 | Lgals1 |
| Ctsd | 3,81E-33 | 1,766775887 | 1 | 0,682 | 6,04E-29 | 1 | Ctsd |
| Zfp36 | 5,17E-33 | 1,747150546 | 0,975 | 0,521 | 8,19E-29 | 1 | Zfp36 |
| Mertk | 5,99E-33 | 0,832936549 | 0,642 | 0,041 | 9,50E-29 | 1 | Mertk |
| P2ry12 | 6,52E-33 | 0,808904511 | 0,531 | 0,004 | 1,03E-28 | 1 | P2ry12 |
| Sema4b | 6,64E-33 | 0,288141104 | 0,679 | 0,05 | 1,05E-28 | 1 | Sema4b |
| Ikzf1 | 7,49E-33 | 0,751378546 | 0,531 | 0,004 | 1,19E-28 | 1 | Ikzf1 |
| H2-DMb1 | 8,88E-33 | 1,266057588 | 0,79 | 0,12 | 1,41E-28 | 1 | H2-DMb1 |
| Cd44 | 9,88E-33 | 0,627273201 | 0,531 | 0,004 | 1,57E-28 | 1 | Cd44 |
| Rnase4 | 1,84E-32 | 1,37399573 | 0,914 | 0,248 | 2,92E-28 | 1 | Rnase4 |
| Runx1 | 3,16E-32 | 0,700503396 | 0,58 | 0,025 | 5,01E-28 | 1 | Runx1 |
| Sema4d | 4,44E-32 | 0,58492625 | 0,506 | 0 | 7,03E-28 | 1 | Sema4d |
| Fgl2 | 4,64E-32 | 0,879039726 | 0,519 | 0,004 | 7,36E-28 | 1 | Fgl2 |
| F13a1 | 4,77E-32 | 1,363365067 | 0,519 | 0,004 | 7,57E-28 | 1 | F13a1 |
| Naaa | 4,91E-32 | 0,736130496 | 0,519 | 0,004 | 7,78E-28 | 1 | Naaa |
| Sh2d1b1 | 5,05E-32 | 0,718340252 | 0,519 | 0,004 | 8,00E-28 | 1 | Sh2d1b1 |
| Arl11 | 6,16E-32 | 0,844862586 | 0,593 | 0,029 | 9,76E-28 | 1 | Arl11 |
| Atp13a2 | 8,76E-32 | 0,949076139 | 0,728 | 0,099 | 1,39E-27 | 1 | Atp13a2 |
| Mapkapk3 | 1,11E-31 | 0,677580928 | 0,531 | 0,008 | 1,76E-27 | 1 | Mapkapk3 |
| Gm22133 | 1,11E-31 | 0,520499036 | 0,963 | 0,781 | 1,76E-27 | 1 | Gm22133 |
| Sh3bp1 | 1,66E-31 | 0,546644652 | 0,543 | 0,012 | 2,63E-27 | 1 | Sh3bp1 |
| Tgfbr1 | 3,01E-31 | 1,042124834 | 0,901 | 0,207 | 4,78E-27 | 1 | Tgfbr1 |
| Lsp1 | 3,05E-31 | 1,015425272 | 0,494 | 0 | 4,83E-27 | 1 | Lsp1 |
| Arhgap9 | 3,05E-31 | 0,534213342 | 0,494 | 0 | 4,83E-27 | 1 | Arhgap9 |
| Tagap | 3,26E-31 | 1,05180134 | 0,519 | 0,008 | 5,17E-27 | 1 | Tagap |
| Milr1 | 3,83E-31 | 0,769847035 | 0,568 | 0,025 | 6,07E-27 | 1 | Milr1 |
| Fyb | 6,18E-31 | 1,187172025 | 0,914 | 0,355 | 9,80E-27 | 1 | Fyb |
| Prkcb | 7,05E-31 | 0,544190178 | 0,506 | 0,004 | 1,12E-26 | 1 | Prkcb |
| Atp6v0c | 1,36E-30 | 1,035339302 | 1 | 0,872 | 2,16E-26 | 1 | Atp6v0c |
| Arhgap15 | 2,07E-30 | 0,587595427 | 0,481 | 0 | 3,28E-26 | 1 | Arhgap15 |
| Runx3 | 2,07E-30 | 0,568885157 | 0,481 | 0 | 3,28E-26 | 1 | Runx3 |
| Hexa | 2,18E-30 | 1,367739242 | 0,914 | 0,322 | 3,45E-26 | 1 | Hexa |
| Itgax | 2,34E-30 | 0,940423925 | 0,494 | 0,004 | 3,70E-26 | 1 | Itgax |
| Tep1 | 2,97E-30 | 0,925350018 | 0,728 | 0,112 | 4,70E-26 | 1 | Tep1 |
| Emilin2 | 3,21E-30 | 0,735592592 | 0,519 | 0,012 | 5,08E-26 | 1 | Emilin2 |
| Cyth4 | 4,45E-30 | 1,136689241 | 0,827 | 0,202 | 7,05E-26 | 1 | Cyth4 |
| Arap1 | 6,23E-30 | 0,662836078 | 0,642 | 0,062 | 9,87E-26 | 1 | Arap1 |
| Arhgap19 | 1,09E-29 | 0,721811133 | 0,556 | 0,029 | 1,73E-25 | 1 | Arhgap19 |
| Abcc3 | 1,19E-29 | 0,69645675 | 0,556 | 0,025 | 1,89E-25 | 1 | Abcc3 |
| Ighm | 1,39E-29 | 1,006104576 | 0,469 | 0 | 2,20E-25 | 1 | Ighm |
| Blnk | 1,39E-29 | 0,9001503 | 0,469 | 0 | 2,20E-25 | 1 | Blnk |
| Col14a1 | 1,39E-29 | 0,803681873 | 0,469 | 0 | 2,20E-25 | 1 | Col14a1 |
| Clec5a | 1,39E-29 | 0,783345542 | 0,469 | 0 | 2,20E-25 | 1 | Clec5a |
| Cxcl14 | 1,56E-29 | 0,679460214 | 0,481 | 0,004 | 2,48E-25 | 1 | Cxcl14 |
| Icosl | 1,77E-29 | 0,311099298 | 0,543 | 0,021 | 2,81E-25 | 1 | Icosl |
| Rps29 | 2,74E-29 | 0,619448213 | 1 | 0,996 | 4,35E-25 | 1 | Rps29 |
| Notch2 | 3,02E-29 | 0,632063456 | 0,901 | 0,252 | 4,79E-25 | 1 | Notch2 |
| Tpd52 | 3,87E-29 | 0,979458038 | 0,938 | 0,273 | 6,14E-25 | 1 | Tpd52 |
| Rassf4 | 4,41E-29 | 0,844686742 | 0,642 | 0,066 | 6,99E-25 | 1 | Rassf4 |
| Fam49b | 4,94E-29 | 1,225835016 | 0,901 | 0,335 | 7,83E-25 | 1 | Fam49b |
| Ptafr | 1,06E-28 | 0,605476117 | 0,469 | 0,004 | 1,68E-24 | 1 | Ptafr |
| Csf3r | 1,12E-28 | 0,796284593 | 0,469 | 0,004 | 1,78E-24 | 1 | Csf3r |
| Nfam1 | 1,13E-28 | 0,766262085 | 0,481 | 0,008 | 1,79E-24 | 1 | Nfam1 |
| Gns | 1,39E-28 | 1,237033357 | 0,938 | 0,339 | 2,21E-24 | 1 | Gns |
| Metrnl | 1,71E-28 | 0,712839286 | 0,531 | 0,025 | 2,71E-24 | 1 | Metrnl |
| Ptk2b | 2,50E-28 | 0,520270734 | 0,481 | 0,008 | 3,96E-24 | 1 | Ptk2b |
| Spp1 | 4,51E-28 | 2,849304487 | 0,568 | 0,045 | 7,14E-24 | 1 | Spp1 |
| Hvcn1 | 6,06E-28 | 0,687798307 | 0,444 | 0 | 9,61E-24 | 1 | Hvcn1 |
| Siglece | 6,06E-28 | 0,62331922 | 0,444 | 0 | 9,61E-24 | 1 | Siglece |

| | | | | | | | |
|---|---|---|---|---|---|---|---|
| Psmb8 | 1,79E-11 | 0,62020329 | 0,951 | 0,74 | 2,84E-07 | MAC | Psmb8 |
| Ubash3b | 1,83E-11 | 0,284984232 | 0,321 | 0,045 | 2,90E-07 | MAC | Ubash3b |
| Socs3 | 1,89E-11 | 0,785208164 | 0,765 | 0,388 | 3,00E-07 | MAC | Socs3 |
| H3f3a | 2,23E-11 | 0,515651002 | 0,975 | 0,843 | 3,53E-07 | MAC | H3f3a |
| Psen2 | 2,25E-11 | 0,478803646 | 0,556 | 0,19 | 3,57E-07 | MAC | Psen2 |
| Aldh3b1 | 2,53E-11 | 0,388904171 | 0,296 | 0,037 | 4,01E-07 | MAC | Aldh3b1 |
| Fau | 2,68E-11 | 0,314117353 | 1 | 0,988 | 4,25E-07 | MAC | Fau |
| Coro1b | 3,74E-11 | 0,701413397 | 0,889 | 0,599 | 5,92E-07 | MAC | Coro1b |
| Hmox1 | 3,80E-11 | 0,595803795 | 0,593 | 0,223 | 6,03E-07 | MAC | Hmox1 |
| Cyp4v3 | 4,96E-11 | 0,348145221 | 0,284 | 0,033 | 7,86E-07 | MAC | Cyp4v3 |
| Lcp2 | 6,21E-11 | 0,505509481 | 0,827 | 0,364 | 9,84E-07 | MAC | Lcp2 |
| Olfm1 | 6,45E-11 | 0,28378505 | 0,333 | 0,054 | 1,02E-06 | MAC | Olfm1 |
| Tiparp | 6,76E-11 | 0,56007753 | 0,395 | 0,091 | 1,07E-06 | MAC | Tiparp |
| Ifi204 | 7,26E-11 | 0,710974407 | 0,531 | 0,19 | 1,15E-06 | MAC | Ifi204 |
| Gla | 7,43E-11 | 0,259021277 | 0,321 | 0,05 | 1,18E-06 | MAC | Gla |
| Bscl2 | 7,47E-11 | 0,440721991 | 0,543 | 0,186 | 1,18E-06 | MAC | Bscl2 |
| Rtcb | 8,23E-11 | 0,505700893 | 0,827 | 0,409 | 1,30E-06 | MAC | Rtcb |
| Casp1 | 9,47E-11 | 0,447189032 | 0,395 | 0,091 | 1,50E-06 | MAC | Casp1 |
| Rps24 | 9,66E-11 | 0,366724221 | 1 | 0,988 | 1,53E-06 | MAC | Rps24 |
| Znrf2 | 9,77E-11 | 0,276292022 | 0,506 | 0,157 | 1,55E-06 | MAC | Znrf2 |
| Naip5 | 1,05E-10 | 0,283946835 | 0,272 | 0,033 | 1,66E-06 | MAC | Naip5 |
| B4galt1 | 1,05E-10 | 0,52061707 | 0,889 | 0,657 | 1,66E-06 | MAC | B4galt1 |
| Rpl37 | 1,05E-10 | 0,309653305 | 1 | 0,975 | 1,66E-06 | MAC | Rpl37 |
| Man2b2 | 1,18E-10 | 0,506227063 | 0,506 | 0,157 | 1,88E-06 | MAC | Man2b2 |
| Tifa | 1,21E-10 | 0,502810017 | 0,642 | 0,244 | 1,91E-06 | MAC | Tifa |
| Rin2 | 1,45E-10 | 0,398051064 | 0,568 | 0,202 | 2,30E-06 | MAC | Rin2 |
| Tyk2 | 1,70E-10 | 0,271148252 | 0,284 | 0,037 | 2,69E-06 | MAC | Tyk2 |
| Slc8b1 | 1,75E-10 | 0,341325667 | 0,333 | 0,062 | 2,77E-06 | MAC | Slc8b1 |
| Fam217b | 1,80E-10 | 0,342401797 | 0,333 | 0,062 | 2,86E-06 | MAC | Fam217b |
| Zbtb7b | 1,88E-10 | 0,270810632 | 0,358 | 0,074 | 2,97E-06 | MAC | Zbtb7b |
| Rps28 | 1,94E-10 | 0,320508052 | 1 | 0,992 | 3,07E-06 | MAC | Rps28 |
| Arpc1b | 1,96E-10 | 0,462137086 | 1 | 0,905 | 3,10E-06 | MAC | Arpc1b |
| Sfxn5 | 2,05E-10 | 0,284802255 | 0,358 | 0,074 | 3,25E-06 | MAC | Sfxn5 |
| Ptpn7 | 2,29E-10 | 0,339500348 | 0,346 | 0,066 | 3,62E-06 | MAC | Ptpn7 |
| Gpx1 | 2,33E-10 | 0,630117103 | 1 | 0,921 | 3,69E-06 | MAC | Gpx1 |
| Rab24 | 2,41E-10 | 0,510135775 | 0,617 | 0,269 | 3,83E-06 | MAC | Rab24 |
| Rogdi | 2,58E-10 | 0,497205869 | 0,58 | 0,223 | 4,10E-06 | MAC | Rogdi |
| Fnip1 | 2,94E-10 | 0,387543954 | 0,444 | 0,136 | 4,66E-06 | MAC | Fnip1 |
| Pold4 | 3,16E-10 | 0,331141164 | 0,568 | 0,215 | 5,01E-06 | MAC | Pold4 |
| Ypel3 | 3,32E-10 | 0,523306983 | 0,556 | 0,211 | 5,26E-06 | MAC | Ypel3 |
| Rpl10a | 3,35E-10 | 0,359997518 | 1 | 0,921 | 5,31E-06 | MAC | Rpl10a |
| Taldo1 | 3,49E-10 | 0,504992913 | 0,864 | 0,603 | 5,53E-06 | MAC | Taldo1 |
| Rpl26 | 3,70E-10 | 0,332330144 | 1 | 0,963 | 5,87E-06 | MAC | Rpl26 |
| Clcn7 | 4,20E-10 | 0,574743996 | 0,531 | 0,194 | 6,66E-06 | MAC | Clcn7 |
| Dnase1l1 | 4,32E-10 | 0,386661687 | 0,42 | 0,107 | 6,84E-06 | MAC | Dnase1l1 |
| Fkbp2 | 4,40E-10 | 0,409685297 | 0,716 | 0,393 | 6,98E-06 | MAC | Fkbp2 |
| AI413582 | 4,44E-10 | 0,435651638 | 0,58 | 0,26 | 7,03E-06 | MAC | AI413582 |
| Oas1g | 4,72E-10 | 0,308331101 | 0,284 | 0,041 | 7,48E-06 | MAC | Oas1g |
| Slc35f6 | 4,75E-10 | 0,266989958 | 0,407 | 0,095 | 7,53E-06 | MAC | Slc35f6 |
| Cited2 | 6,22E-10 | 0,261250454 | 0,333 | 0,066 | 9,85E-06 | MAC | Cited2 |
| Khk | 6,63E-10 | 0,268554416 | 0,259 | 0,033 | 1,05E-05 | MAC | Khk |
| Ppp1r18 | 7,91E-10 | 0,466495029 | 0,815 | 0,554 | 1,25E-05 | MAC | Ppp1r18 |
| Rassf2 | 8,37E-10 | 0,265859129 | 0,667 | 0,273 | 1,33E-05 | MAC | Rassf2 |
| Plekha2 | 9,11E-10 | 0,32913824 | 0,37 | 0,083 | 1,44E-05 | MAC | Plekha2 |
| Gabarap | 9,52E-10 | 0,465031578 | 0,975 | 0,806 | 1,51E-05 | MAC | Gabarap |
| Pisd-ps1 | 9,61E-10 | 0,415847684 | 0,568 | 0,252 | 1,52E-05 | MAC | Pisd-ps1 |
| Nfkbiz | 1,11E-09 | 0,803398477 | 0,63 | 0,298 | 1,76E-05 | MAC | Nfkbiz |
| Trim8 | 1,14E-09 | 0,316832618 | 0,58 | 0,236 | 1,81E-05 | MAC | Trim8 |
| Pacs2 | 1,15E-09 | 0,326344429 | 0,444 | 0,136 | 1,82E-05 | MAC | Pacs2 |
| Gga1 | 1,17E-09 | 0,287208463 | 0,531 | 0,19 | 1,86E-05 | MAC | Gga1 |
| Kdm7a | 1,18E-09 | 0,27440017 | 0,481 | 0,153 | 1,87E-05 | MAC | Kdm7a |
| Atp6v1b2 | 1,28E-09 | 0,636540955 | 0,642 | 0,31 | 2,04E-05 | MAC | Atp6v1b2 |
| Acox3 | 1,32E-09 | 0,447666567 | 0,432 | 0,124 | 2,10E-05 | MAC | Acox3 |
| Grb2 | 1,41E-09 | 0,41818104 | 0,679 | 0,339 | 2,24E-05 | MAC | Grb2 |
| Rab43 | 1,43E-09 | 0,444088343 | 0,531 | 0,207 | 2,26E-05 | MAC | Rab43 |
| Lpin2 | 1,53E-09 | 0,378320467 | 0,333 | 0,07 | 2,43E-05 | MAC | Lpin2 |
| Plgrkt | 1,56E-09 | 0,368980329 | 0,556 | 0,211 | 2,47E-05 | MAC | Plgrkt |
| Plekhm2 | 1,59E-09 | 0,410650711 | 0,296 | 0,054 | 2,51E-05 | MAC | Plekhm2 |
| Plin2 | 1,59E-09 | 0,542380816 | 0,691 | 0,31 | 2,53E-05 | MAC | Plin2 |
| Tmem104 | 1,75E-09 | 0,421002829 | 0,42 | 0,12 | 2,77E-05 | MAC | Tmem104 |
| Snx2 | 1,76E-09 | 0,596221547 | 0,827 | 0,483 | 2,79E-05 | MAC | Snx2 |
| Rilpl2 | 1,78E-09 | 0,530644209 | 0,444 | 0,145 | 2,82E-05 | MAC | Rilpl2 |
| Naip2 | 1,89E-09 | 0,308571636 | 0,407 | 0,107 | 3,00E-05 | MAC | Naip2 |
| Map3k8 | 2,12E-09 | 0,32260867 | 0,469 | 0,153 | 3,37E-05 | MAC | Map3k8 |
| Gm28437 | 2,31E-09 | 0,300053937 | 1 | 0,992 | 3,66E-05 | MAC | Gm28437 |
| Bmp2k | 2,51E-09 | 0,376162685 | 0,988 | 0,86 | 3,99E-05 | MAC | Bmp2k |
| Taok3 | 2,70E-09 | 0,356951209 | 0,457 | 0,149 | 4,28E-05 | MAC | Taok3 |
| Igsf8 | 2,87E-09 | 0,462153522 | 0,358 | 0,095 | 4,55E-05 | MAC | Igsf8 |
| 5031439G07Rik | 2,93E-09 | 0,332936568 | 0,506 | 0,178 | 4,64E-05 | MAC | 5031439G07Rik |
| Pqlc3 | 2,95E-09 | 0,351605941 | 0,296 | 0,058 | 4,68E-05 | MAC | Pqlc3 |
| Snx1 | 3,05E-09 | 0,556844509 | 0,704 | 0,368 | 4,83E-05 | MAC | Snx1 |
| Morc3 | 3,05E-09 | 0,385446898 | 0,481 | 0,157 | 4,84E-05 | MAC | Morc3 |
| Nfkbid | 3,23E-09 | 0,54807823 | 0,333 | 0,074 | 5,12E-05 | MAC | Nfkbid |
| Tmem176b | 3,96E-09 | 0,603594458 | 0,914 | 0,715 | 6,28E-05 | MAC | Tmem176b |
| Letmd1 | 4,13E-09 | 0,301215122 | 0,296 | 0,054 | 6,55E-05 | MAC | Letmd1 |
| Tmem219 | 4,71E-09 | 0,352341813 | 0,506 | 0,178 | 7,46E-05 | MAC | Tmem219 |
| Mfsd1 | 4,75E-09 | 0,562724908 | 0,815 | 0,508 | 7,54E-05 | MAC | Mfsd1 |
| Tmem268 | 4,91E-09 | 0,360747318 | 0,272 | 0,045 | 7,78E-05 | MAC | Tmem268 |
| Btg2 | 5,44E-09 | 0,763811883 | 0,716 | 0,438 | 8,62E-05 | MAC | Btg2 |
| Rps8 | 6,31E-09 | 0,414751109 | 1 | 0,975 | 0,000100056 | MAC | Rps8 |
| Man2a2 | 7,43E-09 | 0,305383657 | 0,321 | 0,07 | 0,000117767 | MAC | Man2a2 |
| Vps18 | 7,89E-09 | 0,482439892 | 0,358 | 0,099 | 0,000125044 | MAC | Vps18 |
| Rpl32 | 7,94E-09 | 0,289426801 | 1 | 0,988 | 0,00012583 | MAC | Rpl32 |
| Ap1g2 | 8,39E-09 | 0,535053461 | 0,259 | 0,045 | 0,000132948 | MAC | Ap1g2 |
| Arpc4 | 9,24E-09 | 0,379032921 | 1 | 0,884 | 0,000146448 | MAC | Arpc4 |
| Crlf3 | 1,03E-08 | 0,410666623 | 0,407 | 0,116 | 0,000163151 | MAC | Crlf3 |
| Arf6 | 1,07E-08 | 0,471831406 | 0,753 | 0,442 | 0,000169072 | MAC | Arf6 |
| Nupr1 | 1,10E-08 | 0,261345505 | 0,272 | 0,05 | 0,000174104 | MAC | Nupr1 |
| Aup1 | 1,23E-08 | 0,541470424 | 0,79 | 0,525 | 0,000194308 | MAC | Aup1 |
| Tsc22d4 | 1,27E-08 | 0,362072565 | 0,716 | 0,397 | 0,000201491 | MAC | Tsc22d4 |
| Mapkapk2 | 1,58E-08 | 0,31206058 | 0,642 | 0,281 | 0,000250506 | MAC | Mapkapk2 |
| Phf11a | 1,59E-08 | 0,261339435 | 0,259 | 0,045 | 0,000251722 | MAC | Phf11a |
| Pip4k2a | 1,65E-08 | 0,358965477 | 0,432 | 0,14 | 0,000261708 | MAC | Pip4k2a |
| Aftph | 2,43E-08 | 0,280698347 | 0,383 | 0,107 | 0,000384763 | MAC | Aftph |
| Rsrp1 | 2,47E-08 | 0,595035526 | 0,926 | 0,719 | 0,000391913 | MAC | Rsrp1 |
| Pak1 | 2,68E-08 | 0,266462141 | 0,333 | 0,079 | 0,000425143 | MAC | Pak1 |
| Ppp1r15a | 3,46E-08 | 0,574772393 | 0,654 | 0,318 | 0,000548282 | MAC | Ppp1r15a |
| Galc | 3,47E-08 | 0,255565691 | 0,346 | 0,095 | 0,000549302 | MAC | Galc |
| Fuca2 | 3,69E-08 | 0,434636214 | 0,506 | 0,194 | 0,000584657 | MAC | Fuca2 |
| Mthfs | 4,48E-08 | 0,259588896 | 0,469 | 0,169 | 0,000710056 | MAC | Mthfs |
| Serp1 | 4,57E-08 | 0,512257959 | 0,864 | 0,574 | 0,000724582 | MAC | Serp1 |
| Rragc | 4,73E-08 | 0,287752796 | 0,519 | 0,202 | 0,000750369 | MAC | Rragc |
| Il17ra | 4,94E-08 | 0,339720252 | 0,37 | 0,103 | 0,000782408 | MAC | Il17ra |
| Atp6ap2 | 5,13E-08 | 0,503285279 | 0,679 | 0,355 | 0,000813986 | MAC | Atp6ap2 |
| Cd274 | 5,17E-08 | 0,373104432 | 0,284 | 0,062 | 0,000819388 | MAC | Cd274 |
| Pdlim4 | 5,55E-08 | 0,338972994 | 0,309 | 0,074 | 0,000879132 | MAC | Pdlim4 |
| Cpne2 | 5,69E-08 | 0,331404293 | 0,593 | 0,277 | 0,000901295 | MAC | Cpne2 |
| Sfi1 | 5,70E-08 | 0,288851182 | 0,494 | 0,198 | 0,000903212 | MAC | Sfi1 |
| Dna2 | 5,92E-08 | 0,254922514 | 0,272 | 0,054 | 0,000938223 | MAC | Dna2 |
| Acsl1 | 6,01E-08 | 0,37854247 | 0,321 | 0,083 | 0,00095261 | MAC | Acsl1 |

| | | | | | | | |
|---|---|---|---|---|---|---|---|
| Stk17b | 7,55E-28 | 0,953648699 | 0,63 | 0,07 | 1,20E-23 | 1 | Stk17b |
| Gmfg | 7,74E-28 | 0,786944147 | 0,889 | 0,264 | 1,23E-23 | 1 | Gmfg |
| B4galnt1 | 1,45E-27 | 0,745542904 | 0,481 | 0,012 | 2,30E-23 | 1 | B4galnt1 |
| Il6ra | 1,51E-27 | 0,578527358 | 0,494 | 0,017 | 2,40E-23 | 1 | Il6ra |
| Glipr1 | 1,56E-27 | 0,594104761 | 0,543 | 0,037 | 2,47E-23 | 1 | Glipr1 |
| Creg1 | 1,86E-27 | 0,938382843 | 0,778 | 0,161 | 2,94E-23 | 1 | Creg1 |
| Fmnl1 | 2,48E-27 | 0,397439907 | 0,568 | 0,041 | 3,92E-23 | 1 | Fmnl1 |
| Gm13166 | 3,24E-27 | 0,267573573 | 0,765 | 0,178 | 5,13E-23 | 1 | Gm13166 |
| Pik3ap1 | 3,93E-27 | 0,46500539 | 0,432 | 0 | 6,24E-23 | 1 | Pik3ap1 |
| Pou2f2 | 3,93E-27 | 0,374063164 | 0,432 | 0 | 6,24E-23 | 1 | Pou2f2 |
| Slc6a6 | 4,03E-27 | 0,792335408 | 0,877 | 0,231 | 6,39E-23 | 1 | Slc6a6 |
| Ms4a4c | 4,73E-27 | 0,992016276 | 0,444 | 0,004 | 7,51E-23 | 1 | Ms4a4c |
| AW112010 | 5,62E-27 | 1,50202659 | 0,827 | 0,236 | 8,91E-23 | 1 | AW112010 |
| H2-Ab1 | 5,67E-27 | 2,549173623 | 0,852 | 0,306 | 8,98E-23 | 1 | H2-Ab1 |
| Camk1d | 6,91E-27 | 0,489802103 | 0,444 | 0,004 | 1,09E-22 | 1 | Camk1d |
| Slfn2 | 7,35E-27 | 0,99484791 | 0,802 | 0,174 | 1,17E-22 | 1 | Slfn2 |
| Fnbp1 | 1,30E-26 | 0,903192124 | 0,778 | 0,174 | 2,05E-22 | 1 | Fnbp1 |
| Selenop | 1,67E-26 | 1,277941566 | 0,988 | 0,818 | 2,64E-22 | 1 | Selenop |
| Tcirg1 | 2,03E-26 | 1,326977134 | 0,889 | 0,376 | 3,22E-22 | 1 | Tcirg1 |
| Fos | 2,48E-26 | 1,538335635 | 0,951 | 0,467 | 3,93E-22 | 1 | Fos |
| Gm14303 | 3,50E-26 | 0,508652242 | 1 | 0,983 | 5,55E-22 | 1 | Gm14303 |
| Rassf5 | 3,65E-26 | 0,824555347 | 0,432 | 0,004 | 5,79E-22 | 1 | Rassf5 |
| Susd3 | 5,46E-26 | 0,617745883 | 0,444 | 0,008 | 8,66E-22 | 1 | Susd3 |
| Abhd12 | 5,56E-26 | 0,777833898 | 0,802 | 0,178 | 8,82E-22 | 1 | Abhd12 |
| Havcr2 | 6,08E-26 | 0,515268894 | 0,432 | 0,004 | 9,63E-22 | 1 | Havcr2 |
| Arhgef6 | 8,94E-26 | 0,368266832 | 0,457 | 0,012 | 1,42E-21 | 1 | Arhgef6 |
| Fabp5 | 1,08E-25 | 0,704699434 | 0,469 | 0,017 | 1,71E-21 | 1 | Fabp5 |
| Plcg2 | 1,25E-25 | 0,660951769 | 0,469 | 0,017 | 1,98E-21 | 1 | Plcg2 |
| Slc43a2 | 1,29E-25 | 0,635968875 | 0,667 | 0,103 | 2,04E-21 | 1 | Slc43a2 |
| Tnfsf13b | 1,60E-25 | 0,737616777 | 0,407 | 0 | 2,54E-21 | 1 | Tnfsf13b |
| Pag1 | 1,60E-25 | 0,347251411 | 0,407 | 0 | 2,54E-21 | 1 | Pag1 |
| Dab2 | 2,06E-25 | 1,346638505 | 0,926 | 0,492 | 3,26E-21 | 1 | Dab2 |
| Syngr1 | 2,13E-25 | 0,476112442 | 0,42 | 0,004 | 3,38E-21 | 1 | Syngr1 |
| Arhgap22 | 2,78E-25 | 0,560932159 | 0,42 | 0,004 | 4,41E-21 | 1 | Arhgap22 |
| Daglb | 3,19E-25 | 0,771501764 | 0,519 | 0,037 | 5,05E-21 | 1 | Daglb |
| Zfp991 | 3,63E-25 | 0,386443859 | 0,593 | 0,062 | 5,75E-21 | 1 | Zfp991 |
| Cd63 | 5,33E-25 | 1,30519191 | 0,951 | 0,595 | 8,45E-21 | 1 | Cd63 |
| Dnase2a | 5,48E-25 | 0,613567353 | 0,519 | 0,037 | 8,69E-21 | 1 | Dnase2a |
| Scpep1 | 5,65E-25 | 0,761035765 | 0,79 | 0,165 | 8,96E-21 | 1 | Scpep1 |
| Snx20 | 6,89E-25 | 0,603895727 | 0,481 | 0,029 | 1,09E-20 | 1 | Snx20 |
| Chd9 | 7,98E-25 | 0,687602891 | 0,951 | 0,43 | 1,27E-20 | 1 | Chd9 |
| Sash3 | 1,01E-24 | 0,586670948 | 0,395 | 0 | 1,60E-20 | 1 | Sash3 |
| Cd300ld | 1,01E-24 | 0,336144078 | 0,395 | 0 | 1,60E-20 | 1 | Cd300ld |
| Pik3cd | 1,30E-24 | 0,684825339 | 0,407 | 0,004 | 2,05E-20 | 1 | Pik3cd |
| Lrmp | 1,33E-24 | 0,516033805 | 0,407 | 0,004 | 2,11E-20 | 1 | Lrmp |
| Kcnn4 | 1,56E-24 | 0,674958892 | 0,407 | 0,004 | 2,48E-20 | 1 | Kcnn4 |
| Cxcl2 | 1,59E-24 | 1,195320277 | 0,42 | 0,008 | 2,53E-20 | 1 | Cxcl2 |
| Cd300a | 1,73E-24 | 0,506783022 | 0,444 | 0,017 | 2,74E-20 | 1 | Cd300a |
| Id2 | 2,40E-24 | 1,296325932 | 0,827 | 0,277 | 3,81E-20 | 1 | Id2 |
| Ppm1h | 5,17E-24 | 0,655674662 | 0,63 | 0,091 | 8,20E-20 | 1 | Ppm1h |
| Slc9a3r1 | 6,01E-24 | 0,666075571 | 0,481 | 0,029 | 9,53E-20 | 1 | Slc9a3r1 |
| Sorl1 | 6,27E-24 | 0,497958317 | 0,383 | 0 | 9,93E-20 | 1 | Sorl1 |
| Gcnt1 | 6,27E-24 | 0,36546244 | 0,383 | 0 | 9,93E-20 | 1 | Gcnt1 |
| Pla2g15 | 7,16E-24 | 0,523222522 | 0,457 | 0,021 | 1,13E-19 | 1 | Pla2g15 |
| Pfkfb4 | 8,91E-24 | 0,475324266 | 0,395 | 0,004 | 1,41E-19 | 1 | Pfkfb4 |
| Gpr34 | 9,40E-24 | 0,701872507 | 0,395 | 0,004 | 1,49E-19 | 1 | Gpr34 |
| Egr2 | 9,43E-24 | 0,940208335 | 0,432 | 0,017 | 1,50E-19 | 1 | Egr2 |
| Tnfaip8 | 1,09E-23 | 0,914377619 | 0,765 | 0,186 | 1,73E-19 | 1 | Tnfaip8 |
| Tnf | 1,11E-23 | 0,938191585 | 0,407 | 0,008 | 1,76E-19 | 1 | Tnf |
| Osm | 3,85E-23 | 0,823072714 | 0,37 | 0 | 6,11E-19 | 1 | Osm |
| Cyp27a1 | 3,85E-23 | 0,713416811 | 0,37 | 0 | 6,11E-19 | 1 | Cyp27a1 |
| Scimp | 3,85E-23 | 0,66422555 | 0,37 | 0 | 6,11E-19 | 1 | Scimp |
| Arhgap4 | 3,85E-23 | 0,49667114 | 0,37 | 0 | 6,11E-19 | 1 | Arhgap4 |
| Slamf7 | 3,85E-23 | 0,433141015 | 0,37 | 0 | 6,11E-19 | 1 | Slamf7 |
| Lgals3bp | 4,49E-23 | 1,092193382 | 0,926 | 0,384 | 7,11E-19 | 1 | Lgals3bp |
| Snx18 | 4,82E-23 | 0,730944186 | 0,543 | 0,066 | 7,63E-19 | 1 | Snx18 |
| Asah1 | 4,94E-23 | 0,999397526 | 0,951 | 0,508 | 7,83E-19 | 1 | Asah1 |
| Esr1 | 5,93E-23 | 0,580748083 | 0,556 | 0,066 | 9,41E-19 | 1 | Esr1 |
| Pim1 | 6,99E-23 | 0,306871403 | 0,815 | 0,248 | 1,11E-18 | 1 | Pim1 |
| Arl4c | 7,36E-23 | 0,627292835 | 0,58 | 0,079 | 1,17E-18 | 1 | Arl4c |
| Cadm1 | 9,60E-23 | 0,62633455 | 0,543 | 0,058 | 1,52E-18 | 1 | Cadm1 |
| Dse | 1,54E-22 | 0,388360609 | 0,407 | 0,012 | 2,44E-18 | 1 | Dse |
| Gpsm3 | 2,14E-22 | 0,40707025 | 0,667 | 0,136 | 3,40E-18 | 1 | Gpsm3 |
| Msrb1 | 2,19E-22 | 0,303364052 | 0,802 | 0,306 | 3,48E-18 | 1 | Msrb1 |
| C4b | 2,34E-22 | 0,710978666 | 0,358 | 0 | 3,72E-18 | 1 | C4b |
| Tlr1 | 2,34E-22 | 0,67315276 | 0,358 | 0 | 3,72E-18 | 1 | Tlr1 |
| Pik3r5 | 2,34E-22 | 0,549391712 | 0,358 | 0 | 3,72E-18 | 1 | Pik3r5 |
| Ptpn22 | 2,34E-22 | 0,434309744 | 0,358 | 0 | 3,72E-18 | 1 | Ptpn22 |
| Wdfy4 | 2,55E-22 | 0,504190759 | 0,444 | 0,025 | 4,04E-18 | 1 | Wdfy4 |
| Ang | 3,07E-22 | 0,531061683 | 0,457 | 0,033 | 4,87E-18 | 1 | Ang |
| Was | 3,82E-22 | 0,336419105 | 0,395 | 0,012 | 6,06E-18 | 1 | Was |
| Gpx3 | 5,36E-22 | 0,965896029 | 0,741 | 0,186 | 8,50E-18 | 1 | Gpx3 |
| Rps6ka1 | 5,65E-22 | 0,685936274 | 0,63 | 0,107 | 8,95E-18 | 1 | Rps6ka1 |
| Cstb | 1,09E-21 | 0,962951432 | 0,951 | 0,657 | 1,72E-17 | 1 | Cstb |
| Vcam1 | 1,14E-21 | 1,26914205 | 0,716 | 0,174 | 1,81E-17 | 1 | Vcam1 |
| Ppcdc | 1,37E-21 | 0,843662871 | 0,593 | 0,099 | 2,17E-17 | 1 | Ppcdc |
| Cmtm7 | 1,40E-21 | 0,78567478 | 0,753 | 0,215 | 2,22E-17 | 1 | Cmtm7 |
| Il1b | 1,41E-21 | 1,172471008 | 0,346 | 0 | 2,24E-17 | 1 | Il1b |
| Fam46c | 1,41E-21 | 0,651824784 | 0,346 | 0 | 2,24E-17 | 1 | Fam46c |
| P2ry14 | 1,41E-21 | 0,565459209 | 0,346 | 0 | 2,24E-17 | 1 | P2ry14 |
| Klra17 | 1,41E-21 | 0,522474672 | 0,346 | 0 | 2,24E-17 | 1 | Klra17 |
| Abhd15 | 1,41E-21 | 0,445438485 | 0,346 | 0 | 2,24E-17 | 1 | Abhd15 |
| I830077J02Rik | 1,41E-21 | 0,303579479 | 0,346 | 0 | 2,24E-17 | 1 | I830077J02Rik |
| Tmem86a | 1,73E-21 | 0,710968724 | 0,654 | 0,12 | 2,75E-17 | 1 | Tmem86a |
| Tspan32 | 2,03E-21 | 0,517061799 | 0,358 | 0,004 | 3,21E-17 | 1 | Tspan32 |
| Adssl1 | 2,76E-21 | 0,674850795 | 0,395 | 0,017 | 4,38E-17 | 1 | Adssl1 |
| Ptgs1 | 3,59E-21 | 1,004198622 | 0,679 | 0,153 | 5,69E-17 | 1 | Ptgs1 |
| P2rx4 | 3,64E-21 | 0,958053931 | 0,741 | 0,223 | 5,78E-17 | 1 | P2rx4 |
| Tmsb4x | 3,78E-21 | 0,524215299 | 1 | 1 | 5,99E-17 | 1 | Tmsb4x |
| Blvrb | 4,16E-21 | 0,595817096 | 0,667 | 0,136 | 6,59E-17 | 1 | Blvrb |
| Sowahc | 8,27E-21 | 0,597567389 | 0,444 | 0,033 | 1,31E-16 | 1 | Sowahc |
| Arhgap24 | 8,41E-21 | 0,388451462 | 0,333 | 0 | 1,33E-16 | 1 | Arhgap24 |
| Pstpip1 | 8,41E-21 | 0,362602899 | 0,333 | 0 | 1,33E-16 | 1 | Pstpip1 |
| Clec4d | 8,41E-21 | 0,323312751 | 0,333 | 0 | 1,33E-16 | 1 | Clec4d |
| Lamp1 | 9,83E-21 | 0,800039046 | 1 | 0,893 | 1,56E-16 | 1 | Lamp1 |
| B3gnt8 | 9,99E-21 | 0,568065754 | 0,494 | 0,066 | 1,58E-16 | 1 | B3gnt8 |
| Uap1l1 | 1,07E-20 | 0,807724392 | 0,494 | 0,058 | 1,70E-16 | 1 | Uap1l1 |
| Rnaset2a | 1,09E-20 | 0,481170359 | 0,951 | 0,471 | 1,73E-16 | 1 | Rnaset2a |
| AI662270 | 1,57E-20 | 0,43863666 | 0,432 | 0,033 | 2,48E-16 | 1 | AI662270 |
| Gltp | 1,81E-20 | 0,899557904 | 0,815 | 0,31 | 2,87E-16 | 1 | Gltp |
| Akna | 2,02E-20 | 0,577668582 | 0,494 | 0,054 | 3,21E-16 | 1 | Akna |
| Srgn | 3,25E-20 | 1,041232525 | 0,963 | 0,736 | 5,16E-16 | 1 | Srgn |
| Cysltr1 | 3,48E-20 | 0,562471883 | 0,383 | 0,017 | 5,51E-16 | 1 | Cysltr1 |
| P2ry13 | 4,96E-20 | 0,525826406 | 0,321 | 0 | 7,87E-16 | 1 | P2ry13 |
| Lilra5 | 4,96E-20 | 0,443206512 | 0,321 | 0 | 7,87E-16 | 1 | Lilra5 |
| Acss1 | 4,96E-20 | 0,427991117 | 0,321 | 0 | 7,87E-16 | 1 | Acss1 |
| Cndp2 | 5,18E-20 | 0,959349471 | 0,84 | 0,314 | 8,21E-16 | 1 | Cndp2 |
| Ptpn18 | 7,08E-20 | 0,621629777 | 0,901 | 0,5 | 1,12E-15 | 1 | Ptpn18 |
| Ccdc88b | 8,01E-20 | 0,292537393 | 0,333 | 0,004 | 1,27E-15 | 1 | Ccdc88b |
| Eps8 | 1,04E-19 | 0,311074945 | 0,333 | 0,004 | 1,65E-15 | 1 | Eps8 |

| | | | | | | | |
|---|---|---|---|---|---|---|---|
| Tnfrsf1b | 6,71E-08 | 0,304839792 | 0,617 | 0,281 | 0,001064223 | MAC | Tnfrsf1b |
| Akr1b3 | 6,91E-08 | 0,337364873 | 0,716 | 0,364 | 0,001094718 | MAC | Akr1b3 |
| Ncoa3 | 7,04E-08 | 0,404264804 | 0,741 | 0,421 | 0,001115937 | MAC | Ncoa3 |
| Ifngr1 | 7,53E-08 | 0,574604344 | 0,889 | 0,657 | 0,001194152 | MAC | Ifngr1 |
| Gm42418 | 7,91E-08 | 0,324815373 | 0,753 | 0,496 | 0,001253297 | MAC | Gm42418 |
| Psmb9 | 8,54E-08 | 0,411952036 | 0,778 | 0,492 | 0,001353094 | MAC | Psmb9 |
| Parp1 | 8,68E-08 | 0,462397937 | 0,556 | 0,248 | 0,001375698 | MAC | Parp1 |
| Klf6 | 8,75E-08 | 0,559287297 | 0,827 | 0,554 | 0,00138672 | MAC | Klf6 |
| Tmem189 | 8,82E-08 | 0,460042083 | 0,531 | 0,244 | 0,001398114 | MAC | Tmem189 |
| Ndufa13 | 9,80E-08 | 0,39710895 | 0,864 | 0,707 | 0,001553524 | MAC | Ndufa13 |
| Cyth1 | 9,95E-08 | 0,385011682 | 0,568 | 0,256 | 0,001577463 | MAC | Cyth1 |
| Cdk9 | 1,03E-07 | 0,379559798 | 0,494 | 0,19 | 0,001625808 | MAC | Cdk9 |
| Rack1 | 1,04E-07 | 0,404986594 | 1 | 0,926 | 0,001650619 | MAC | Rack1 |
| Mapk14 | 1,27E-07 | 0,422128891 | 0,519 | 0,219 | 0,002005811 | MAC | Mapk14 |
| Gm7536 | 1,27E-07 | 0,341241622 | 0,988 | 0,938 | 0,002016701 | MAC | Gm7536 |
| Flcn | 1,40E-07 | 0,510609678 | 0,358 | 0,107 | 0,002213389 | MAC | Flcn |
| Rap2a | 1,40E-07 | 0,390838553 | 0,457 | 0,169 | 0,002222496 | MAC | Rap2a |
| Vps26a | 1,43E-07 | 0,368504367 | 0,58 | 0,277 | 0,002262571 | MAC | Vps26a |
| Slc25a45 | 1,52E-07 | 0,387565813 | 0,284 | 0,066 | 0,002408527 | MAC | Slc25a45 |
| Fes | 1,74E-07 | 0,400772623 | 0,691 | 0,335 | 0,002761886 | MAC | Fes |
| Dgkz | 1,78E-07 | 0,511542067 | 0,741 | 0,467 | 0,002822216 | MAC | Dgkz |
| Hspa1a | 1,80E-07 | 1,174612449 | 0,519 | 0,269 | 0,002846349 | MAC | Hspa1a |
| Tmem192 | 1,84E-07 | 0,479401824 | 0,407 | 0,153 | 0,002909046 | MAC | Tmem192 |
| Wdr81 | 1,87E-07 | 0,269206671 | 0,259 | 0,054 | 0,002968104 | MAC | Wdr81 |
| Dennd1a | 2,00E-07 | 0,251730345 | 0,432 | 0,161 | 0,0031665 | MAC | Dennd1a |
| Plekhb2 | 2,06E-07 | 0,297982996 | 0,469 | 0,178 | 0,00327252 | MAC | Plekhb2 |
| 2210016F16Rik | 2,19E-07 | 0,283348809 | 0,432 | 0,153 | 0,003478031 | MAC | 2210016F16Rik |
| Nceh1 | 2,29E-07 | 0,342073045 | 0,333 | 0,095 | 0,003625298 | MAC | Nceh1 |
| Gaa | 2,29E-07 | 0,32016142 | 0,444 | 0,161 | 0,003626059 | MAC | Gaa |
| Adam9 | 2,35E-07 | 0,360520503 | 0,79 | 0,467 | 0,003719169 | MAC | Adam9 |
| Rpl37a | 2,46E-07 | 0,253571596 | 1 | 0,996 | 0,003897212 | MAC | Rpl37a |
| Myo9b | 2,53E-07 | 0,399475533 | 0,58 | 0,285 | 0,004006585 | MAC | Myo9b |
| Mdfic | 2,71E-07 | 0,363253621 | 0,506 | 0,219 | 0,004294311 | MAC | Mdfic |
| Slc35c2 | 2,72E-07 | 0,33979996 | 0,432 | 0,165 | 0,004313519 | MAC | Slc35c2 |
| Rps13 | 2,80E-07 | 0,34570896 | 0,963 | 0,913 | 0,004438004 | MAC | Rps13 |
| Gm10275 | 2,81E-07 | 0,321386147 | 0,988 | 0,888 | 0,004455872 | MAC | Gm10275 |
| Tnfrsf21 | 2,84E-07 | 0,315534155 | 0,284 | 0,066 | 0,004505844 | MAC | Tnfrsf21 |
| Rps26 | 2,95E-07 | 0,256091399 | 1 | 0,992 | 0,004679862 | MAC | Rps26 |
| Rgs19 | 3,32E-07 | 0,343106929 | 0,568 | 0,269 | 0,00525813 | MAC | Rgs19 |
| Camk1 | 3,47E-07 | 0,562994961 | 0,716 | 0,417 | 0,005507566 | MAC | Camk1 |
| Akr1a1 | 3,52E-07 | 0,40842193 | 0,951 | 0,913 | 0,005578477 | MAC | Akr1a1 |
| Nptn | 3,60E-07 | 0,474010964 | 0,716 | 0,475 | 0,005704295 | MAC | Nptn |
| Dpp7 | 3,78E-07 | 0,264869139 | 0,284 | 0,07 | 0,005995632 | MAC | Dpp7 |
| Tpp1 | 4,17E-07 | 0,450689817 | 0,79 | 0,517 | 0,006611116 | MAC | Tpp1 |
| Dleu2 | 4,20E-07 | 0,322452128 | 0,605 | 0,306 | 0,006653505 | MAC | Dleu2 |
| Pigs | 4,45E-07 | 0,456908012 | 0,519 | 0,227 | 0,007060433 | MAC | Pigs |
| Cdt1 | 4,85E-07 | 0,251512871 | 0,346 | 0,103 | 0,007682716 | MAC | Cdt1 |
| Taf6l | 4,86E-07 | 0,363202758 | 0,593 | 0,293 | 0,007709968 | MAC | Taf6l |
| Cox14 | 5,11E-07 | 0,342245265 | 0,691 | 0,45 | 0,008108268 | MAC | Cox14 |
| Blvra | 5,15E-07 | 0,309984188 | 0,704 | 0,376 | 0,008163604 | MAC | Blvra |
| Gsto1 | 5,27E-07 | 0,388805125 | 0,728 | 0,446 | 0,008359742 | MAC | Gsto1 |
| Ssh2 | 5,30E-07 | 0,566563372 | 0,753 | 0,492 | 0,008408263 | MAC | Ssh2 |
| Stat6 | 5,37E-07 | 0,46673398 | 0,543 | 0,244 | 0,008511975 | MAC | Stat6 |
| Crip1 | 5,43E-07 | 0,577245067 | 0,765 | 0,554 | 0,008613079 | MAC | Crip1 |
| Btg1 | 5,49E-07 | 0,45796331 | 0,938 | 0,789 | 0,008706478 | MAC | Btg1 |
| Scand1 | 5,67E-07 | 0,281490816 | 0,741 | 0,521 | 0,008985107 | MAC | Scand1 |
| Vrk2 | 6,35E-07 | 0,279880631 | 0,259 | 0,062 | 0,010059752 | MAC | Vrk2 |
| Npc1 | 7,38E-07 | 0,275798249 | 0,395 | 0,14 | 0,01169185 | MAC | Npc1 |
| Atp6v1a | 8,23E-07 | 0,420850531 | 0,691 | 0,388 | 0,013040315 | MAC | Atp6v1a |
| Rpl21 | 9,14E-07 | 0,302270835 | 1 | 0,963 | 0,014486009 | MAC | Rpl21 |
| Gm26917 | 9,87E-07 | 0,334005648 | 0,988 | 0,855 | 0,015653415 | MAC | Gm26917 |
| Htatip2 | 1,00E-06 | 0,316806292 | 0,407 | 0,149 | 0,015872115 | MAC | Htatip2 |
| Pmepa1 | 1,03E-06 | 0,375002635 | 0,79 | 0,521 | 0,016399906 | MAC | Pmepa1 |
| Cnp | 1,04E-06 | 0,291586412 | 0,407 | 0,145 | 0,016449048 | MAC | Cnp |
| mt-Atp6 | 1,10E-06 | 0,306721723 | 1 | 0,983 | 0,017496445 | MAC | mt-Atp6 |
| G6pdx | 1,14E-06 | 0,423892542 | 0,395 | 0,149 | 0,018137153 | MAC | G6pdx |
| Atp5e | 1,20E-06 | 0,314681132 | 0,975 | 0,864 | 0,018985818 | MAC | Atp5e |
| AC121965.1 | 1,26E-06 | 0,29458315 | 0,901 | 0,764 | 0,020048374 | MAC | AC121965.1 |
| Cox8a | 1,38E-06 | 0,332429412 | 0,951 | 0,88 | 0,021860291 | MAC | Cox8a |
| Cap1 | 1,39E-06 | 0,335142778 | 0,938 | 0,76 | 0,021984693 | MAC | Cap1 |
| Tex10 | 1,46E-06 | 0,309308564 | 0,432 | 0,178 | 0,023171982 | MAC | Tex10 |
| Edem1 | 1,59E-06 | 0,306618845 | 0,654 | 0,355 | 0,025200255 | MAC | Edem1 |
| Actb | 1,60E-06 | 0,275931915 | 1 | 1 | 0,025410614 | MAC | Actb |
| Per1 | 1,67E-06 | 0,274415015 | 0,617 | 0,331 | 0,026488303 | MAC | Per1 |
| Slc36a1 | 1,74E-06 | 0,320110234 | 0,259 | 0,066 | 0,027607457 | MAC | Slc36a1 |
| Gnpda1 | 1,75E-06 | 0,525451447 | 0,383 | 0,149 | 0,027700075 | MAC | Gnpda1 |
| 2810474O19Rik | 1,83E-06 | 0,340457258 | 0,679 | 0,393 | 0,028980775 | MAC | 2810474O19Rik |
| Txnip | 1,88E-06 | 0,250458743 | 0,741 | 0,409 | 0,029872936 | MAC | Txnip |
| Gba | 1,90E-06 | 0,421069872 | 0,593 | 0,302 | 0,030151438 | MAC | Gba |
| Dtnbp1 | 1,99E-06 | 0,264801413 | 0,358 | 0,124 | 0,031552819 | MAC | Dtnbp1 |
| Naglu | 2,11E-06 | 0,336616854 | 0,395 | 0,153 | 0,03342043 | MAC | Naglu |
| Cox6b1 | 2,28E-06 | 0,31456173 | 0,988 | 0,926 | 0,036129408 | MAC | Cox6b1 |
| Prex1 | 2,29E-06 | 0,477377259 | 0,679 | 0,409 | 0,036277186 | MAC | Prex1 |
| Fam50a | 2,29E-06 | 0,271526568 | 0,37 | 0,136 | 0,036346359 | MAC | Fam50a |
| Hn1 | 2,33E-06 | 0,448412304 | 0,827 | 0,55 | 0,036906254 | MAC | Hn1 |
| Tgif1 | 2,42E-06 | 0,377933745 | 0,605 | 0,302 | 0,038291948 | MAC | Tgif1 |
| Gapdh | 2,46E-06 | 0,278842381 | 1 | 0,988 | 0,039008427 | MAC | Gapdh |
| M6pr | 2,50E-06 | 0,432072169 | 0,84 | 0,558 | 0,039681663 | MAC | M6pr |
| Anxa4 | 2,56E-06 | 0,331962501 | 0,593 | 0,31 | 0,040586703 | MAC | Anxa4 |
| Mrps36 | 2,58E-06 | 0,268668285 | 0,506 | 0,273 | 0,040968991 | MAC | Mrps36 |
| Rnf19b | 2,59E-06 | 0,340871007 | 0,395 | 0,153 | 0,041126594 | MAC | Rnf19b |
| Atp6ap1 | 2,62E-06 | 0,355720406 | 0,827 | 0,512 | 0,041529857 | MAC | Atp6ap1 |
| Tmem176a | 2,67E-06 | 0,525873627 | 0,778 | 0,607 | 0,042316403 | MAC | Tmem176a |
| Rcbtb2 | 2,70E-06 | 0,374847106 | 0,506 | 0,236 | 0,042797029 | MAC | Rcbtb2 |
| Bcl2l1 | 2,88E-06 | 0,50225105 | 0,58 | 0,322 | 0,045626493 | MAC | Bcl2l1 |
| Trafd1 | 3,02E-06 | 0,514206768 | 0,605 | 0,331 | 0,047899631 | MAC | Trafd1 |
| Nhlrc3 | 3,26E-06 | 0,349961382 | 0,432 | 0,182 | 0,051735818 | MAC | Nhlrc3 |
| Wipf1 | 3,68E-06 | 0,36239454 | 0,556 | 0,322 | 0,058387708 | MAC | Wipf1 |
| Cnpy3 | 4,09E-06 | 0,382239611 | 0,543 | 0,273 | 0,064770194 | MAC | Cnpy3 |
| Elf4 | 4,16E-06 | 0,25896207 | 0,346 | 0,12 | 0,06601765 | MAC | Elf4 |
| Actr2 | 4,63E-06 | 0,25243645 | 1 | 0,938 | 0,073358429 | MAC | Actr2 |
| Rps27a | 4,72E-06 | 0,266554083 | 1 | 0,975 | 0,074895431 | MAC | Rps27a |
| Csf2rb | 4,83E-06 | 0,414080554 | 0,593 | 0,322 | 0,076625401 | MAC | Csf2rb |
| Ddt | 4,91E-06 | 0,255558656 | 0,407 | 0,169 | 0,077902543 | MAC | Ddt |
| Rnh1 | 5,07E-06 | 0,37633876 | 0,815 | 0,496 | 0,080439727 | MAC | Rnh1 |
| Zfp622 | 5,08E-06 | 0,276062419 | 0,407 | 0,161 | 0,080573376 | MAC | Zfp622 |
| Tor1a | 5,15E-06 | 0,377415083 | 0,531 | 0,285 | 0,081602432 | MAC | Tor1a |
| Smpdl3a | 5,91E-06 | 0,27364814 | 0,457 | 0,186 | 0,093753494 | MAC | Smpdl3a |
| Pacsin2 | 6,49E-06 | 0,301107499 | 0,642 | 0,335 | 0,102865868 | MAC | Pacsin2 |
| Cyb5r4 | 6,78E-06 | 0,31605335 | 0,457 | 0,202 | 0,107544474 | MAC | Cyb5r4 |
| Ubl3 | 6,98E-06 | 0,42328339 | 0,778 | 0,525 | 0,110574542 | MAC | Ubl3 |
| Tmem55b | 7,18E-06 | 0,369626709 | 0,457 | 0,223 | 0,113858887 | MAC | Tmem55b |
| Pip5k1c | 8,09E-06 | 0,449937828 | 0,593 | 0,36 | 0,128269372 | MAC | Pip5k1c |
| Lifr | 8,29E-06 | 0,374037249 | 0,321 | 0,116 | 0,131471297 | MAC | Lifr |
| Mfsd11 | 8,59E-06 | 0,310880442 | 0,333 | 0,12 | 0,136236867 | MAC | Mfsd11 |
| Irak4 | 8,73E-06 | 0,261272298 | 0,358 | 0,14 | 0,138343019 | MAC | Irak4 |
| Ccdc115 | 8,80E-06 | 0,328650839 | 0,481 | 0,231 | 0,139516881 | MAC | Ccdc115 |
| Glrx | 8,93E-06 | 0,394557061 | 0,617 | 0,335 | 0,141488708 | MAC | Glrx |
| Ddx28 | 9,28E-06 | 0,337459919 | 0,259 | 0,079 | 0,147046663 | MAC | Ddx28 |
| Ndufa2 | 9,67E-06 | 0,348987899 | 0,951 | 0,868 | 0,153322264 | MAC | Ndufa2 |

| | | | | | | | |
|---|---|---|---|---|---|---|---|
| Ppt1 | 1,50E-19 | 0,966749552 | 0,914 | 0,421 | 2,37E-15 | 1 | Ppt1 |
| Pik3cg | 1,52E-19 | 0,48954774 | 0,481 | 0,054 | 2,41E-15 | 1 | Pik3cg |
| Soat1 | 1,76E-19 | 0,468381438 | 0,778 | 0,215 | 2,79E-15 | 1 | Soat1 |
| Neurl3 | 1,85E-19 | 0,672542852 | 0,593 | 0,107 | 2,93E-15 | 1 | Neurl3 |
| Ccnd2 | 2,14E-19 | 0,591870319 | 0,432 | 0,037 | 3,40E-15 | 1 | Ccnd2 |
| Slc37a2 | 2,75E-19 | 0,455059437 | 0,383 | 0,021 | 4,36E-15 | 1 | Slc37a2 |
| Rasal3 | 2,90E-19 | 0,529944769 | 0,309 | 0 | 4,59E-15 | 1 | Rasal3 |
| Tlr8 | 2,90E-19 | 0,49041397 | 0,309 | 0 | 4,59E-15 | 1 | Tlr8 |
| Btk | 2,90E-19 | 0,398201553 | 0,309 | 0 | 4,59E-15 | 1 | Btk |
| Gm26740 | 2,90E-19 | 0,346443608 | 0,309 | 0 | 4,59E-15 | 1 | Gm26740 |
| Dcxr | 2,92E-19 | 0,427762424 | 0,457 | 0,045 | 4,63E-15 | 1 | Dcxr |
| Smim3 | 3,36E-19 | 0,622848311 | 0,457 | 0,045 | 5,33E-15 | 1 | Smim3 |
| Irs2 | 4,52E-19 | 0,304990634 | 0,321 | 0,004 | 7,17E-15 | 1 | Irs2 |
| Lyl1 | 4,89E-19 | 0,696506323 | 0,506 | 0,079 | 7,75E-15 | 1 | Lyl1 |
| Lacc1 | 5,98E-19 | 0,517177254 | 0,395 | 0,029 | 9,48E-15 | 1 | Lacc1 |
| Dok3 | 6,36E-19 | 0,850796404 | 0,778 | 0,293 | 1,01E-14 | 1 | Dok3 |
| Timp2 | 8,10E-19 | 0,640926194 | 0,642 | 0,149 | 1,28E-14 | 1 | Timp2 |
| Cyfip2 | 8,40E-19 | 0,266014314 | 0,321 | 0,004 | 1,33E-14 | 1 | Cyfip2 |
| Rhoh | 9,49E-19 | 0,542404648 | 0,346 | 0,012 | 1,50E-14 | 1 | Rhoh |
| Hivep3 | 1,13E-18 | 0,42617658 | 0,469 | 0,054 | 1,80E-14 | 1 | Hivep3 |
| Rhog | 1,20E-18 | 0,758091983 | 0,963 | 0,562 | 1,90E-14 | 1 | Rhog |
| Gpr171 | 1,68E-18 | 0,588052816 | 0,296 | 0 | 2,66E-14 | 1 | Gpr171 |
| Myo1g | 1,68E-18 | 0,48580086 | 0,296 | 0 | 2,66E-14 | 1 | Myo1g |
| Galnt6 | 1,68E-18 | 0,477593558 | 0,296 | 0 | 2,66E-14 | 1 | Galnt6 |
| Ikbke | 1,68E-18 | 0,415045879 | 0,296 | 0 | 2,66E-14 | 1 | Ikbke |
| Dennd1c | 1,68E-18 | 0,381452533 | 0,296 | 0 | 2,66E-14 | 1 | Dennd1c |
| Atp1a3 | 1,68E-18 | 0,308285402 | 0,296 | 0 | 2,66E-14 | 1 | Atp1a3 |
| Rplp1 | 1,91E-18 | 0,4786856 | 1 | 0,988 | 3,03E-14 | 1 | Rplp1 |
| Csk | 2,82E-18 | 0,684396563 | 0,728 | 0,219 | 4,47E-14 | 1 | Csk |
| Il4ra | 3,01E-18 | 0,851831436 | 0,753 | 0,223 | 4,78E-14 | 1 | Il4ra |
| Junb | 3,33E-18 | 0,972479719 | 0,963 | 0,686 | 5,28E-14 | 1 | Junb |
| Arg1 | 3,35E-18 | 1,565771763 | 0,321 | 0,008 | 5,31E-14 | 1 | Arg1 |
| Kctd12 | 3,88E-18 | 0,713023622 | 0,704 | 0,202 | 6,14E-14 | 1 | Kctd12 |
| Dtx4 | 3,92E-18 | 0,357952817 | 0,395 | 0,029 | 6,21E-14 | 1 | Dtx4 |
| B3gnt7 | 4,29E-18 | 0,29619881 | 0,321 | 0,008 | 6,80E-14 | 1 | B3gnt7 |
| Lyn | 4,61E-18 | 0,857210201 | 0,938 | 0,537 | 7,31E-14 | 1 | Lyn |
| Tmem37 | 6,50E-18 | 0,795621974 | 0,728 | 0,244 | 1,03E-13 | 1 | Tmem37 |
| Pitpnm1 | 6,74E-18 | 0,666624923 | 0,42 | 0,045 | 1,07E-13 | 1 | Pitpnm1 |
| Hcls1 | 7,00E-18 | 0,681283415 | 0,901 | 0,409 | 1,11E-13 | 1 | Hcls1 |
| Sh3pxd2b | 7,33E-18 | 0,687197637 | 0,617 | 0,161 | 1,16E-13 | 1 | Sh3pxd2b |
| Pnpla7 | 7,50E-18 | 0,334212043 | 0,407 | 0,037 | 1,19E-13 | 1 | Pnpla7 |
| Abr | 7,54E-18 | 0,444957589 | 0,617 | 0,128 | 1,20E-13 | 1 | Abr |
| 5430427O19Rik | 9,59E-18 | 0,596711118 | 0,284 | 0 | 1,52E-13 | 1 | 5430427O19Rik |
| Tnfsf9 | 9,59E-18 | 0,3803071 | 0,284 | 0 | 1,52E-13 | 1 | Tnfsf9 |
| Il21r | 9,59E-18 | 0,360762191 | 0,284 | 0 | 1,52E-13 | 1 | Il21r |
| Shtn1 | 9,59E-18 | 0,295439637 | 0,284 | 0 | 1,52E-13 | 1 | Shtn1 |
| Nlrp3 | 9,59E-18 | 0,26010347 | 0,284 | 0 | 1,52E-13 | 1 | Nlrp3 |
| Gpr183 | 1,01E-17 | 0,492959763 | 0,469 | 0,058 | 1,60E-13 | 1 | Gpr183 |
| Mlxip | 1,05E-17 | 0,604818629 | 0,605 | 0,128 | 1,66E-13 | 1 | Mlxip |
| Ccrl2 | 1,20E-17 | 1,032785251 | 0,568 | 0,124 | 1,90E-13 | 1 | Ccrl2 |
| Ptger4 | 1,51E-17 | 0,535734658 | 0,296 | 0,004 | 2,40E-13 | 1 | Ptger4 |
| Gm9843 | 1,52E-17 | 0,446390455 | 1 | 0,988 | 2,41E-13 | 1 | Gm9843 |
| Gas6 | 2,04E-17 | 0,768216389 | 0,432 | 0,05 | 3,23E-13 | 1 | Gas6 |
| Rasgef1b | 2,19E-17 | 0,456818573 | 0,432 | 0,045 | 3,46E-13 | 1 | Rasgef1b |
| Gmip | 2,21E-17 | 0,50334324 | 0,457 | 0,058 | 3,51E-13 | 1 | Gmip |
| Trex1 | 2,52E-17 | 0,641870459 | 0,494 | 0,083 | 3,99E-13 | 1 | Trex1 |
| Tnfrsf13b | 2,72E-17 | 0,452468117 | 0,296 | 0,004 | 4,32E-13 | 1 | Tnfrsf13b |
| Cryl1 | 3,04E-17 | 0,304046136 | 0,383 | 0,029 | 4,82E-13 | 1 | Cryl1 |
| Egr1 | 3,06E-17 | 1,103825904 | 0,765 | 0,298 | 4,85E-13 | 1 | Egr1 |
| Myo5a | 3,27E-17 | 0,67561715 | 0,605 | 0,14 | 5,18E-13 | 1 | Myo5a |
| Ppp1r21 | 3,74E-17 | 0,481426419 | 0,519 | 0,083 | 5,93E-13 | 1 | Ppp1r21 |
| Rnf130 | 3,78E-17 | 0,601648237 | 0,765 | 0,285 | 6,00E-13 | 1 | Rnf130 |
| Lcp1 | 3,85E-17 | 0,874187342 | 0,975 | 0,719 | 6,10E-13 | 1 | Lcp1 |
| Irf2bp2 | 3,91E-17 | 0,476537671 | 0,704 | 0,219 | 6,20E-13 | 1 | Irf2bp2 |
| Gusb | 4,24E-17 | 0,940811342 | 0,864 | 0,463 | 6,72E-13 | 1 | Gusb |
| Clta | 4,41E-17 | 0,715709448 | 0,963 | 0,694 | 6,99E-13 | 1 | Clta |
| Emp3 | 5,00E-17 | 0,656386728 | 0,852 | 0,343 | 7,93E-13 | 1 | Emp3 |
| Ccl8 | 5,44E-17 | 1,154760223 | 0,272 | 0 | 8,62E-13 | 1 | Ccl8 |
| Ap1b1 | 6,04E-17 | 0,731000105 | 0,704 | 0,223 | 9,58E-13 | 1 | Ap1b1 |
| B2m | 6,16E-17 | 0,505602121 | 1 | 1 | 9,76E-13 | 1 | B2m |
| 1700017B05Rik | 6,80E-17 | 0,775383546 | 0,556 | 0,124 | 1,08E-12 | 1 | 1700017B05Rik |
| Fam134b | 7,82E-17 | 0,381311619 | 0,333 | 0,017 | 1,24E-12 | 1 | Fam134b |
| 1810011H11Rik | 8,74E-17 | 0,384323324 | 0,284 | 0,004 | 1,39E-12 | 1 | 1810011H11Rik |
| Tnfsf13 | 1,05E-16 | 0,437969132 | 0,383 | 0,037 | 1,67E-12 | 1 | Tnfsf13 |
| Paox | 1,09E-16 | 0,395249477 | 0,432 | 0,062 | 1,72E-12 | 1 | Paox |
| Atf3 | 1,28E-16 | 1,028513801 | 0,63 | 0,165 | 2,03E-12 | 1 | Atf3 |
| Phf11b | 1,43E-16 | 0,608505462 | 0,432 | 0,058 | 2,27E-12 | 1 | Phf11b |
| Erp29 | 1,80E-16 | 0,645234386 | 0,901 | 0,504 | 2,85E-12 | 1 | Erp29 |
| Rap2b | 2,14E-16 | 0,662673498 | 0,864 | 0,533 | 3,40E-12 | 1 | Rap2b |
| Nampt | 2,41E-16 | 0,727924815 | 0,716 | 0,231 | 3,82E-12 | 1 | Nampt |
| Ckb | 2,65E-16 | 0,919194369 | 0,778 | 0,326 | 4,20E-12 | 1 | Ckb |
| Skap2 | 2,76E-16 | 0,622818388 | 0,765 | 0,277 | 4,38E-12 | 1 | Skap2 |
| Il1rn | 3,05E-16 | 0,711378733 | 0,259 | 0 | 4,84E-12 | 1 | Il1rn |
| Trpm2 | 3,05E-16 | 0,544577763 | 0,259 | 0 | 4,84E-12 | 1 | Trpm2 |
| Cd244 | 3,05E-16 | 0,427697384 | 0,259 | 0 | 4,84E-12 | 1 | Cd244 |
| Psd4 | 3,05E-16 | 0,409417958 | 0,259 | 0 | 4,84E-12 | 1 | Psd4 |
| Vill | 3,05E-16 | 0,379542702 | 0,259 | 0 | 4,84E-12 | 1 | Vill |
| Rnf150 | 3,05E-16 | 0,253934024 | 0,259 | 0 | 4,84E-12 | 1 | Rnf150 |
| Hcst | 4,27E-16 | 0,302186009 | 0,481 | 0,083 | 6,77E-12 | 1 | Hcst |
| Ccr2 | 4,87E-16 | 0,684310477 | 0,272 | 0,004 | 7,72E-12 | 1 | Ccr2 |
| Mknk1 | 5,36E-16 | 0,538016491 | 0,568 | 0,132 | 8,50E-12 | 1 | Mknk1 |
| Sdf2l1 | 5,96E-16 | 0,510732026 | 0,704 | 0,207 | 9,45E-12 | 1 | Sdf2l1 |
| Lrch4 | 6,16E-16 | 0,643674174 | 0,63 | 0,182 | 9,76E-12 | 1 | Lrch4 |
| Atp8a1 | 6,27E-16 | 0,343750138 | 0,556 | 0,112 | 9,94E-12 | 1 | Atp8a1 |
| Rpl10-ps3 | 6,71E-16 | 0,503409129 | 1 | 0,921 | 1,06E-11 | 1 | Rpl10-ps3 |
| Trpv2 | 7,28E-16 | 0,61375328 | 0,407 | 0,058 | 1,15E-11 | 1 | Trpv2 |
| Epsti1 | 7,57E-16 | 0,469595964 | 0,407 | 0,05 | 1,20E-11 | 1 | Epsti1 |
| Cln8 | 7,70E-16 | 0,563221072 | 0,531 | 0,103 | 1,22E-11 | 1 | Cln8 |
| Plxnb2 | 8,64E-16 | 0,8664309 | 0,753 | 0,293 | 1,37E-11 | 1 | Plxnb2 |
| N4bp2l1 | 1,11E-15 | 0,400397317 | 0,284 | 0,008 | 1,76E-11 | 1 | N4bp2l1 |
| Gm29216 | 1,17E-15 | 0,542367253 | 1 | 1 | 1,85E-11 | 1 | Gm29216 |
| Sgpl1 | 1,19E-15 | 0,657117666 | 0,667 | 0,215 | 1,89E-11 | 1 | Sgpl1 |
| Creb5 | 1,21E-15 | 0,382026432 | 0,383 | 0,041 | 1,92E-11 | 1 | Creb5 |
| Inpp5d | 1,23E-15 | 0,741392624 | 0,901 | 0,517 | 1,96E-11 | 1 | Inpp5d |
| Rab20 | 1,38E-15 | 0,685717148 | 0,42 | 0,058 | 2,18E-11 | 1 | Rab20 |
| Pgd | 1,52E-15 | 0,524444206 | 0,802 | 0,306 | 2,41E-11 | 1 | Pgd |
| Ldhb | 1,61E-15 | 0,565261504 | 0,358 | 0,033 | 2,56E-11 | 1 | Ldhb |
| Sp110 | 2,14E-15 | 0,575952333 | 0,654 | 0,182 | 3,39E-11 | 1 | Sp110 |
| Sh3bp2 | 2,29E-15 | 0,313881132 | 0,481 | 0,083 | 3,63E-11 | 1 | Sh3bp2 |
| Gm5150 | 2,75E-15 | 0,333877016 | 0,259 | 0,004 | 4,37E-11 | 1 | Gm5150 |
| Gdf3 | 3,94E-15 | 0,841966805 | 0,296 | 0,017 | 6,24E-11 | 1 | Gdf3 |
| Hk3 | 4,12E-15 | 0,359976259 | 0,259 | 0,004 | 6,53E-11 | 1 | Hk3 |
| Spon1 | 5,08E-15 | 0,369231045 | 0,272 | 0,008 | 8,05E-11 | 1 | Spon1 |
| Iqgap2 | 5,20E-15 | 0,300328243 | 0,272 | 0,008 | 8,24E-11 | 1 | Iqgap2 |
| Fgd4 | 5,36E-15 | 0,253209369 | 0,321 | 0,025 | 8,50E-11 | 1 | Fgd4 |
| Syk | 5,56E-15 | 0,393628744 | 0,506 | 0,103 | 8,81E-11 | 1 | Syk |
| Slc29a3 | 6,02E-15 | 0,563428806 | 0,444 | 0,074 | 9,55E-11 | 1 | Slc29a3 |
| Dhrs3 | 6,22E-15 | 0,742196147 | 0,741 | 0,298 | 9,86E-11 | 1 | Dhrs3 |
| Usp2 | 7,53E-15 | 0,369816958 | 0,296 | 0,017 | 1,19E-10 | 1 | Usp2 |
| Grk2 | 8,91E-15 | 0,644933194 | 0,765 | 0,318 | 1,41E-10 | 1 | Grk2 |

| | | | | | | | |
|---|---|---|---|---|---|---|---|
| Ncapg2 | 9,76E-06 | 0,415761684 | 0,346 | 0,136 | 0,15469208 | MAC | Ncapg2 |
| Kxd1 | 9,89E-06 | 0,287555683 | 0,494 | 0,236 | 0,156810509 | MAC | Kxd1 |
| Ms4a6d | 9,95E-06 | 0,449845956 | 0,914 | 0,777 | 0,157661587 | MAC | Ms4a6d |
| Psmb10 | 1,01E-05 | 0,351103358 | 0,716 | 0,463 | 0,160047223 | MAC | Psmb10 |
| Ccnl1 | 1,04E-05 | 0,399996713 | 0,654 | 0,409 | 0,164085963 | MAC | Ccnl1 |
| Cuta | 1,14E-05 | 0,257625009 | 0,679 | 0,446 | 0,1807053 | MAC | Cuta |
| Siva1 | 1,17E-05 | 0,369090619 | 0,395 | 0,169 | 0,185836272 | MAC | Siva1 |
| Arrdc1 | 1,24E-05 | 0,35977545 | 0,568 | 0,306 | 0,196907732 | MAC | Arrdc1 |
| Nfil3 | 1,32E-05 | 0,374162748 | 0,272 | 0,087 | 0,209915239 | MAC | Nfil3 |
| Adam17 | 1,39E-05 | 0,375295267 | 0,58 | 0,343 | 0,220950103 | MAC | Adam17 |
| Acer3 | 1,43E-05 | 0,432093874 | 0,481 | 0,24 | 0,226900967 | MAC | Acer3 |
| Smap2 | 1,58E-05 | 0,283444208 | 0,543 | 0,281 | 0,250145372 | MAC | Smap2 |
| BC005537 | 1,59E-05 | 0,463756994 | 0,691 | 0,442 | 0,251665269 | MAC | BC005537 |
| Rps12l1 | 1,59E-05 | 0,313469105 | 0,963 | 0,855 | 0,252374209 | MAC | Rps12l1 |
| Mif | 1,65E-05 | 0,350488226 | 0,963 | 0,814 | 0,261009391 | MAC | Mif |
| Bmyc | 1,74E-05 | 0,312660213 | 0,432 | 0,207 | 0,276256042 | MAC | Bmyc |
| Acp2 | 1,80E-05 | 0,38597389 | 0,432 | 0,194 | 0,285111216 | MAC | Acp2 |
| Pabpc1 | 1,89E-05 | 0,250776888 | 0,988 | 0,942 | 0,300015297 | MAC | Pabpc1 |
| Rhbdf2 | 2,17E-05 | 0,36100564 | 0,296 | 0,107 | 0,344556075 | MAC | Rhbdf2 |
| Heatr5a | 2,27E-05 | 0,404769656 | 0,37 | 0,161 | 0,359961722 | MAC | Heatr5a |
| Rnf13 | 2,45E-05 | 0,315453115 | 0,654 | 0,368 | 0,388255311 | MAC | Rnf13 |
| Tmem14c | 2,46E-05 | 0,311756394 | 0,765 | 0,537 | 0,390646008 | MAC | Tmem14c |
| Wdr26 | 2,56E-05 | 0,353666909 | 0,691 | 0,438 | 0,405755681 | MAC | Wdr26 |
| Mon2 | 2,74E-05 | 0,256787941 | 0,333 | 0,128 | 0,435091688 | MAC | Mon2 |
| Ifi211 | 3,16E-05 | 0,555267337 | 0,358 | 0,157 | 0,500438229 | MAC | Ifi211 |
| Spg21 | 3,16E-05 | 0,253956438 | 0,63 | 0,347 | 0,501644693 | MAC | Spg21 |
| Got1 | 3,17E-05 | 0,315694764 | 0,333 | 0,128 | 0,502705264 | MAC | Got1 |
| Fkbp5 | 3,30E-05 | 0,326733948 | 0,704 | 0,488 | 0,522835258 | MAC | Fkbp5 |
| Rpl17 | 3,31E-05 | 0,284752912 | 0,988 | 0,905 | 0,525460259 | MAC | Rpl17 |
| Tab2 | 3,32E-05 | 0,297073291 | 0,494 | 0,26 | 0,526972407 | MAC | Tab2 |
| Rbfa | 3,38E-05 | 0,293478347 | 0,42 | 0,19 | 0,536569295 | MAC | Rbfa |
| Acaa1a | 4,03E-05 | 0,288458877 | 0,543 | 0,289 | 0,638575488 | MAC | Acaa1a |
| Rab8b | 4,27E-05 | 0,353639868 | 0,617 | 0,38 | 0,677268308 | MAC | Rab8b |
| Zmiz2 | 4,58E-05 | 0,373698644 | 0,481 | 0,264 | 0,726072889 | MAC | Zmiz2 |
| Il10rb | 4,78E-05 | 0,392888027 | 0,667 | 0,426 | 0,757823196 | MAC | Il10rb |
| Stk24 | 4,80E-05 | 0,323531773 | 0,37 | 0,169 | 0,761298812 | MAC | Stk24 |
| Mcl1 | 4,82E-05 | 0,321860004 | 0,84 | 0,653 | 0,763520793 | MAC | Mcl1 |
| Haus8 | 4,93E-05 | 0,280554427 | 0,272 | 0,099 | 0,780957369 | MAC | Haus8 |
| Gna13 | 4,97E-05 | 0,347415731 | 0,691 | 0,467 | 0,787658785 | MAC | Gna13 |
| Rpl7a | 5,03E-05 | 0,271818892 | 1 | 0,955 | 0,796827427 | MAC | Rpl7a |
| Srsf9 | 5,06E-05 | 0,278987998 | 0,605 | 0,397 | 0,801722727 | MAC | Srsf9 |
| Tbc1d14 | 5,53E-05 | 0,292106893 | 0,407 | 0,19 | 0,876494101 | MAC | Tbc1d14 |
| Calr | 5,71E-05 | 0,276697 | 0,963 | 0,872 | 0,905681229 | MAC | Calr |
| Psme1 | 5,76E-05 | 0,35124452 | 0,852 | 0,707 | 0,912454285 | MAC | Psme1 |
| Lamp2 | 5,91E-05 | 0,362412043 | 0,901 | 0,707 | 0,936613516 | MAC | Lamp2 |
| Neu1 | 6,10E-05 | 0,407221473 | 0,457 | 0,244 | 0,96739424 | MAC | Neu1 |
| Atp2b1 | 6,12E-05 | 0,253957571 | 0,84 | 0,533 | 0,97041529 | MAC | Atp2b1 |
| Cdc42se1 | 6,27E-05 | 0,374070549 | 0,741 | 0,554 | 0,993821394 | MAC | Cdc42se1 |
| Cox5a | 6,81E-05 | 0,329080342 | 0,914 | 0,752 | 1 | MAC | Cox5a |
| Naga | 6,85E-05 | 0,327309289 | 0,556 | 0,314 | 1 | MAC | Naga |
| Lamtor1 | 6,90E-05 | 0,309293295 | 0,728 | 0,504 | 1 | MAC | Lamtor1 |
| Rpsa | 7,55E-05 | 0,259320969 | 1 | 0,934 | 1 | MAC | Rpsa |
| Fam174a | 7,74E-05 | 0,335653986 | 0,531 | 0,306 | 1 | MAC | Fam174a |
| Rasa3 | 7,81E-05 | 0,281353895 | 0,383 | 0,174 | 1 | MAC | Rasa3 |
| Tln1 | 7,98E-05 | 0,297268989 | 0,926 | 0,831 | 1 | MAC | Tln1 |
| Gab2 | 9,01E-05 | 0,349502881 | 0,494 | 0,277 | 1 | MAC | Gab2 |
| Gna12 | 9,38E-05 | 0,418271329 | 0,691 | 0,471 | 1 | MAC | Gna12 |
| Tkt | 0,000104102 | 0,254276801 | 0,765 | 0,537 | 1 | MAC | Tkt |
| Pnrc1 | 0,000110131 | 0,337376004 | 0,58 | 0,372 | 1 | MAC | Pnrc1 |
| Pgam1 | 0,000121458 | 0,481171014 | 0,877 | 0,674 | 1 | MAC | Pgam1 |
| Agps | 0,000134388 | 0,252685423 | 0,469 | 0,24 | 1 | MAC | Agps |
| Slc48a1 | 0,000135331 | 0,263373231 | 0,42 | 0,207 | 1 | MAC | Slc48a1 |
| Scamp2 | 0,000140012 | 0,272735841 | 0,691 | 0,455 | 1 | MAC | Scamp2 |
| Fam234a | 0,0001411 | 0,280142013 | 0,358 | 0,161 | 1 | MAC | Fam234a |
| Ski | 0,000141441 | 0,331579784 | 0,654 | 0,45 | 1 | MAC | Ski |
| Xylt2 | 0,000143447 | 0,257927367 | 0,272 | 0,107 | 1 | MAC | Xylt2 |
| Psme2 | 0,000143635 | 0,275848433 | 0,889 | 0,764 | 1 | MAC | Psme2 |
| Ap2a2 | 0,000147513 | 0,304147864 | 0,728 | 0,554 | 1 | MAC | Ap2a2 |
| Oaz1 | 0,000154596 | 0,264464644 | 1 | 0,959 | 1 | MAC | Oaz1 |
| Herc4 | 0,000154734 | 0,359136314 | 0,358 | 0,174 | 1 | MAC | Herc4 |
| Vdac2 | 0,000160588 | 0,303634252 | 0,877 | 0,649 | 1 | MAC | Vdac2 |
| Frmd4b | 0,00016283 | 0,475232633 | 0,531 | 0,331 | 1 | MAC | Frmd4b |
| Ahsa1 | 0,000173995 | 0,355986811 | 0,543 | 0,331 | 1 | MAC | Ahsa1 |
| Hpf1 | 0,000174231 | 0,275094629 | 0,481 | 0,277 | 1 | MAC | Hpf1 |
| Fkbp15 | 0,000187463 | 0,283872936 | 0,531 | 0,31 | 1 | MAC | Fkbp15 |
| Pmvk | 0,000200092 | 0,287175187 | 0,432 | 0,219 | 1 | MAC | Pmvk |
| Narfl | 0,000215593 | 0,272199991 | 0,259 | 0,099 | 1 | MAC | Narfl |
| Rpl36al | 0,000217303 | 0,291709092 | 0,938 | 0,798 | 1 | MAC | Rpl36al |
| Capza2 | 0,000217745 | 0,255354865 | 0,951 | 0,818 | 1 | MAC | Capza2 |
| Tob2 | 0,000231997 | 0,271365236 | 0,37 | 0,194 | 1 | MAC | Tob2 |
| Mdh1 | 0,000238528 | 0,282455577 | 0,654 | 0,434 | 1 | MAC | Mdh1 |
| Ubc | 0,000245708 | 0,302117554 | 1 | 0,971 | 1 | MAC | Ubc |
| Ccdc86 | 0,000248756 | 0,299585161 | 0,333 | 0,149 | 1 | MAC | Ccdc86 |
| mt-Nd5 | 0,000254438 | 0,258544722 | 0,988 | 0,979 | 1 | MAC | mt-Nd5 |
| Sptssa | 0,000280441 | 0,314468222 | 0,691 | 0,521 | 1 | MAC | Sptssa |
| Sec11c | 0,000281978 | 0,343426263 | 0,679 | 0,496 | 1 | MAC | Sec11c |
| Colgalt1 | 0,000285128 | 0,444913719 | 0,605 | 0,442 | 1 | MAC | Colgalt1 |
| Atp6v0e | 0,000294512 | 0,277202448 | 0,864 | 0,665 | 1 | MAC | Atp6v0e |
| Jun | 0,000296263 | 0,439974447 | 0,654 | 0,459 | 1 | MAC | Jun |
| Syngr2 | 0,000328371 | 0,358140639 | 0,815 | 0,599 | 1 | MAC | Syngr2 |
| Incenp | 0,000388183 | 0,312358927 | 0,284 | 0,128 | 1 | MAC | Incenp |
| Tpt1 | 0,000393596 | 0,256080445 | 1 | 0,983 | 1 | MAC | Tpt1 |
| Gstp1 | 0,000410133 | 0,283807883 | 0,63 | 0,475 | 1 | MAC | Gstp1 |
| Ivns1abp | 0,000421008 | 0,339652005 | 0,827 | 0,616 | 1 | MAC | Ivns1abp |
| Lrpap1 | 0,000434266 | 0,294307354 | 0,605 | 0,384 | 1 | MAC | Lrpap1 |
| Map3k1 | 0,000443908 | 0,327620612 | 0,617 | 0,455 | 1 | MAC | Map3k1 |
| Man1c1 | 0,000459086 | 0,28352606 | 0,432 | 0,252 | 1 | MAC | Man1c1 |
| Cept1 | 0,000579451 | 0,257571054 | 0,42 | 0,231 | 1 | MAC | Cept1 |
| Plbd2 | 0,000580448 | 0,292592948 | 0,519 | 0,314 | 1 | MAC | Plbd2 |
| Cltc | 0,000586272 | 0,326264032 | 0,938 | 0,851 | 1 | MAC | Cltc |
| mt-Nd2 | 0,000606054 | 0,27418639 | 0,975 | 0,979 | 1 | MAC | mt-Nd2 |
| Tnfaip3 | 0,000608246 | 0,51500519 | 0,259 | 0,116 | 1 | MAC | Tnfaip3 |
| Rpl10 | 0,000622519 | 0,272833737 | 0,988 | 0,93 | 1 | MAC | Rpl10 |
| 2310035C23Rik | 0,000688 | 0,275192349 | 0,469 | 0,289 | 1 | MAC | 2310035C23Rik |
| Tm6sf1 | 0,000732851 | 0,319738078 | 0,432 | 0,236 | 1 | MAC | Tm6sf1 |
| Mrps26 | 0,000850556 | 0,253855037 | 0,481 | 0,293 | 1 | MAC | Mrps26 |
| Tsc22d3 | 0,001002564 | 0,269443601 | 0,741 | 0,55 | 1 | MAC | Tsc22d3 |
| Rfc2 | 0,001011926 | 0,295254248 | 0,556 | 0,368 | 1 | MAC | Rfc2 |
| Dram2 | 0,001028189 | 0,279026799 | 0,716 | 0,558 | 1 | MAC | Dram2 |
| Tpi1 | 0,001106805 | 0,291732728 | 0,926 | 0,711 | 1 | MAC | Tpi1 |
| Echs1 | 0,001148989 | 0,355575642 | 0,481 | 0,322 | 1 | MAC | Echs1 |
| Arhgap25 | 0,001207998 | 0,317985339 | 0,568 | 0,368 | 1 | MAC | Arhgap25 |
| Zfyve19 | 0,001300237 | 0,327754084 | 0,272 | 0,132 | 1 | MAC | Zfyve19 |
| Ensa | 0,00130699 | 0,3004799 | 0,605 | 0,409 | 1 | MAC | Ensa |
| Cdca3 | 0,001342141 | 0,41526518 | 0,259 | 0,12 | 1 | MAC | Cdca3 |
| Mif4gd | 0,001384247 | 0,258736045 | 0,395 | 0,219 | 1 | MAC | Mif4gd |
| Gpr137b-ps | 0,001428522 | 0,265826649 | 0,432 | 0,26 | 1 | MAC | Gpr137b-ps |
| Dnm2 | 0,001570228 | 0,256464864 | 0,741 | 0,579 | 1 | MAC | Dnm2 |
| Marcks | 0,001595358 | 0,270161179 | 1 | 0,959 | 1 | MAC | Marcks |
| Atp6v1c1 | 0,00165266 | 0,302802144 | 0,506 | 0,331 | 1 | MAC | Atp6v1c1 |

| | | | | | | | |
|---|---|---|---|---|---|---|---|
| Fam46a | 9,34E-15 | 0,452056672 | 0,346 | 0,033 | 1,48E-10 | 1 | Fam46a |
| Mt2 | 1,00E-14 | 0,32885324 | 0,432 | 0,066 | 1,58E-10 | 1 | Mt2 |
| Cd37 | 1,22E-14 | 0,256594967 | 0,284 | 0,012 | 1,94E-10 | 1 | Cd37 |
| Rps9 | 1,26E-14 | 0,463912496 | 1 | 0,959 | 2,00E-10 | 1 | Rps9 |
| Sh3bgrl3 | 1,33E-14 | 0,511549595 | 0,988 | 0,955 | 2,10E-10 | 1 | Sh3bgrl3 |
| Rplp2 | 1,59E-14 | 0,43557302 | 1 | 0,971 | 2,51E-10 | 1 | Rplp2 |
| Plk3 | 1,72E-14 | 0,732739522 | 0,481 | 0,103 | 2,73E-10 | 1 | Plk3 |
| Hebp1 | 1,89E-14 | 0,510041106 | 0,679 | 0,236 | 2,99E-10 | 1 | Hebp1 |
| Arpc2 | 2,18E-14 | 0,538401095 | 1 | 0,934 | 3,45E-10 | 1 | Arpc2 |
| Cfb | 2,25E-14 | 0,91965093 | 0,63 | 0,186 | 3,56E-10 | 1 | Cfb |
| Lamtor4 | 2,60E-14 | 0,436108828 | 0,741 | 0,31 | 4,12E-10 | 1 | Lamtor4 |
| Adamtsl4 | 2,73E-14 | 0,494964011 | 0,284 | 0,017 | 4,33E-10 | 1 | Adamtsl4 |
| Nmt1 | 3,00E-14 | 0,511115036 | 0,938 | 0,595 | 4,75E-10 | 1 | Nmt1 |
| Grina | 3,37E-14 | 0,765950615 | 0,79 | 0,372 | 5,34E-10 | 1 | Grina |
| Rbpj | 3,43E-14 | 0,578363677 | 0,704 | 0,264 | 5,43E-10 | 1 | Rbpj |
| Irf8 | 4,52E-14 | 0,626553181 | 0,815 | 0,355 | 7,16E-10 | 1 | Irf8 |
| Limd2 | 4,62E-14 | 0,614563961 | 0,765 | 0,326 | 7,32E-10 | 1 | Limd2 |
| Dusp1 | 5,08E-14 | 1,103805735 | 0,877 | 0,744 | 8,06E-10 | 1 | Dusp1 |
| Il18bp | 5,36E-14 | 0,590276155 | 0,481 | 0,099 | 8,50E-10 | 1 | Il18bp |
| Ier3 | 5,78E-14 | 1,028140617 | 0,778 | 0,397 | 9,16E-10 | 1 | Ier3 |
| Stx7 | 7,18E-14 | 0,695742817 | 0,827 | 0,421 | 1,14E-09 | 1 | Stx7 |
| Cotl1 | 7,39E-14 | 0,565070912 | 0,951 | 0,57 | 1,17E-09 | 1 | Cotl1 |
| Rpl35a | 7,44E-14 | 0,460085251 | 1 | 0,955 | 1,18E-09 | 1 | Rpl35a |
| Adrb2 | 8,23E-14 | 0,690821824 | 0,321 | 0,029 | 1,31E-09 | 1 | Adrb2 |
| Fosb | 8,49E-14 | 0,50847215 | 0,543 | 0,132 | 1,35E-09 | 1 | Fosb |
| Rnf149 | 8,77E-14 | 0,625950326 | 0,469 | 0,103 | 1,39E-09 | 1 | Rnf149 |
| Rasa4 | 1,02E-13 | 0,491052565 | 0,358 | 0,045 | 1,61E-09 | 1 | Rasa4 |
| Aldh2 | 1,13E-13 | 0,528215229 | 0,432 | 0,079 | 1,79E-09 | 1 | Aldh2 |
| Ogfrl1 | 1,30E-13 | 0,402116622 | 0,531 | 0,132 | 2,05E-09 | 1 | Ogfrl1 |
| Stk4 | 1,72E-13 | 0,539395855 | 0,481 | 0,112 | 2,73E-09 | 1 | Stk4 |
| Rel | 1,81E-13 | 0,602543455 | 0,494 | 0,12 | 2,86E-09 | 1 | Rel |
| G530011O06Rik | 1,82E-13 | 0,402739018 | 0,321 | 0,033 | 2,89E-09 | 1 | G530011O06Rik |
| Snx24 | 1,85E-13 | 0,356335862 | 0,407 | 0,066 | 2,94E-09 | 1 | Snx24 |
| Oxct1 | 2,11E-13 | 0,52743885 | 0,519 | 0,132 | 3,35E-09 | 1 | Oxct1 |
| Renbp | 2,74E-13 | 0,593152122 | 0,568 | 0,165 | 4,34E-09 | 1 | Renbp |
| Stab1 | 3,44E-13 | 0,808652468 | 0,938 | 0,897 | 5,46E-09 | 1 | Stab1 |
| Rnaset2b | 3,62E-13 | 0,578768571 | 0,988 | 0,744 | 5,74E-09 | 1 | Rnaset2b |
| Gna15 | 3,73E-13 | 0,443882291 | 0,272 | 0,017 | 5,91E-09 | 1 | Gna15 |
| S100a1 | 3,85E-13 | 0,385697876 | 0,568 | 0,165 | 6,10E-09 | 1 | S100a1 |
| Mgst1 | 4,13E-13 | 0,544946181 | 0,42 | 0,079 | 6,55E-09 | 1 | Mgst1 |
| Nfe2l2 | 4,78E-13 | 0,775013045 | 0,889 | 0,529 | 7,58E-09 | 1 | Nfe2l2 |
| Extl3 | 6,66E-13 | 0,53124816 | 0,691 | 0,256 | 1,06E-08 | 1 | Extl3 |
| Lhfpl2 | 6,77E-13 | 0,627004965 | 0,358 | 0,054 | 1,07E-08 | 1 | Lhfpl2 |
| Dennd4b | 8,28E-13 | 0,473687598 | 0,272 | 0,021 | 1,31E-08 | 1 | Dennd4b |
| Hfe | 8,72E-13 | 0,613687222 | 0,63 | 0,227 | 1,38E-08 | 1 | Hfe |
| Mkrn1 | 9,01E-13 | 0,437200793 | 0,63 | 0,207 | 1,43E-08 | 1 | Mkrn1 |
| Cd302 | 1,12E-12 | 0,557097882 | 0,704 | 0,331 | 1,78E-08 | 1 | Cd302 |
| Zfp36l2 | 1,24E-12 | 0,510718524 | 0,827 | 0,455 | 1,97E-08 | 1 | Zfp36l2 |
| Coro7 | 1,32E-12 | 0,606731924 | 0,716 | 0,302 | 2,09E-08 | 1 | Coro7 |
| Plagl2 | 1,35E-12 | 0,47316562 | 0,432 | 0,091 | 2,15E-08 | 1 | Plagl2 |
| Rps25 | 1,36E-12 | 0,436130949 | 1 | 0,963 | 2,16E-08 | 1 | Rps25 |
| Ezr | 1,43E-12 | 0,444466913 | 0,346 | 0,05 | 2,27E-08 | 1 | Ezr |
| Pfn1 | 1,44E-12 | 0,391110037 | 1 | 0,996 | 2,29E-08 | 1 | Pfn1 |
| Pld3 | 1,46E-12 | 0,651098261 | 0,765 | 0,331 | 2,31E-08 | 1 | Pld3 |
| Ptpn1 | 1,57E-12 | 0,648909454 | 0,827 | 0,45 | 2,49E-08 | 1 | Ptpn1 |
| 1600014C10Rik | 1,94E-12 | 0,435332021 | 0,494 | 0,124 | 3,08E-08 | 1 | 1600014C10Rik |
| Frrs1 | 2,23E-12 | 0,436794436 | 0,358 | 0,054 | 3,53E-08 | 1 | Frrs1 |
| mt-Nd6 | 2,43E-12 | 0,495943928 | 1 | 0,938 | 3,85E-08 | 1 | mt-Nd6 |
| Tmem50b | 2,45E-12 | 0,615273094 | 0,556 | 0,169 | 3,88E-08 | 1 | Tmem50b |
| Mfsd12 | 2,99E-12 | 0,317135005 | 0,395 | 0,074 | 4,74E-08 | 1 | Mfsd12 |
| Dapp1 | 3,27E-12 | 0,461333746 | 0,321 | 0,041 | 5,19E-08 | 1 | Dapp1 |
| Dnmt3a | 3,77E-12 | 0,399229807 | 0,543 | 0,157 | 5,97E-08 | 1 | Dnmt3a |
| Rpl37rt | 3,82E-12 | 0,328749956 | 1 | 0,967 | 6,05E-08 | 1 | Rpl37rt |
| Atp6v1f | 3,93E-12 | 0,52629993 | 0,951 | 0,702 | 6,24E-08 | 1 | Atp6v1f |
| Etv5 | 4,01E-12 | 0,562947656 | 0,469 | 0,12 | 6,36E-08 | 1 | Etv5 |
| Pi4k2a | 4,14E-12 | 0,440058705 | 0,383 | 0,07 | 6,56E-08 | 1 | Pi4k2a |
| Atp6v0b | 4,44E-12 | 0,627465194 | 0,827 | 0,467 | 7,05E-08 | 1 | Atp6v0b |
| Epb41l2 | 4,84E-12 | 0,69486121 | 0,765 | 0,384 | 7,68E-08 | 1 | Epb41l2 |
| Fam213b | 5,37E-12 | 0,335481219 | 0,296 | 0,033 | 8,52E-08 | 1 | Fam213b |
| Vamp8 | 5,43E-12 | 0,60299128 | 0,926 | 0,711 | 8,61E-08 | 1 | Vamp8 |
| Gk | 5,86E-12 | 0,319990012 | 0,284 | 0,029 | 9,28E-08 | 1 | Gk |
| Snx8 | 5,97E-12 | 0,343920183 | 0,346 | 0,054 | 9,47E-08 | 1 | Snx8 |
| Klc4 | 6,54E-12 | 0,485411731 | 0,469 | 0,116 | 1,04E-07 | 1 | Klc4 |
| Fam96a | 8,14E-12 | 0,582269561 | 0,815 | 0,434 | 1,29E-07 | 1 | Fam96a |
| Lrp12 | 8,61E-12 | 0,287319276 | 0,259 | 0,021 | 1,36E-07 | 1 | Lrp12 |
| B3galnt1 | 1,06E-11 | 0,353528189 | 0,284 | 0,029 | 1,68E-07 | 1 | B3galnt1 |
| Mroh1 | 1,16E-11 | 0,370345061 | 0,358 | 0,062 | 1,83E-07 | 1 | Mroh1 |
| Fam234b | 1,23E-11 | 0,347375668 | 0,321 | 0,045 | 1,96E-07 | 1 | Fam234b |
| Samhd1 | 1,32E-11 | 0,577276734 | 0,679 | 0,264 | 2,09E-07 | 1 | Samhd1 |
| Psmb8 | 1,79E-11 | 0,62020329 | 0,951 | 0,74 | 2,84E-07 | 1 | Psmb8 |
| Ubash3b | 1,83E-11 | 0,284984232 | 0,321 | 0,045 | 2,90E-07 | 1 | Ubash3b |
| Socs3 | 1,89E-11 | 0,785208164 | 0,765 | 0,388 | 3,00E-07 | 1 | Socs3 |
| H3f3a | 2,23E-11 | 0,515651002 | 0,975 | 0,843 | 3,53E-07 | 1 | H3f3a |
| Psen2 | 2,25E-11 | 0,478803646 | 0,556 | 0,19 | 3,57E-07 | 1 | Psen2 |
| Aldh3b1 | 2,53E-11 | 0,388904171 | 0,296 | 0,037 | 4,01E-07 | 1 | Aldh3b1 |
| Fau | 2,68E-11 | 0,314117353 | 1 | 0,988 | 4,25E-07 | 1 | Fau |
| Coro1b | 3,74E-11 | 0,701413397 | 0,889 | 0,599 | 5,92E-07 | 1 | Coro1b |
| Hmox1 | 3,80E-11 | 0,595803795 | 0,593 | 0,223 | 6,03E-07 | 1 | Hmox1 |
| Cyp4v3 | 4,96E-11 | 0,348145221 | 0,284 | 0,033 | 7,86E-07 | 1 | Cyp4v3 |
| Lcp2 | 6,21E-11 | 0,505509481 | 0,827 | 0,364 | 9,84E-07 | 1 | Lcp2 |
| Olfm1 | 6,45E-11 | 0,28378505 | 0,333 | 0,054 | 1,02E-06 | 1 | Olfm1 |
| Tiparp | 6,76E-11 | 0,56007753 | 0,395 | 0,091 | 1,07E-06 | 1 | Tiparp |
| Ifi204 | 7,26E-11 | 0,710974407 | 0,531 | 0,19 | 1,15E-06 | 1 | Ifi204 |
| Gla | 7,43E-11 | 0,259021277 | 0,321 | 0,05 | 1,18E-06 | 1 | Gla |
| Bscl2 | 7,47E-11 | 0,440721991 | 0,543 | 0,186 | 1,18E-06 | 1 | Bscl2 |
| Rtcb | 8,23E-11 | 0,505700893 | 0,827 | 0,409 | 1,30E-06 | 1 | Rtcb |
| Casp1 | 9,47E-11 | 0,447189032 | 0,395 | 0,091 | 1,50E-06 | 1 | Casp1 |
| Rps24 | 9,66E-11 | 0,366724221 | 1 | 0,988 | 1,53E-06 | 1 | Rps24 |
| Znrf2 | 9,77E-11 | 0,276292022 | 0,506 | 0,157 | 1,55E-06 | 1 | Znrf2 |
| Naip5 | 1,05E-10 | 0,283946835 | 0,272 | 0,033 | 1,66E-06 | 1 | Naip5 |
| B4galt1 | 1,05E-10 | 0,52061707 | 0,889 | 0,657 | 1,66E-06 | 1 | B4galt1 |
| Rpl37 | 1,05E-10 | 0,309653305 | 1 | 0,975 | 1,66E-06 | 1 | Rpl37 |
| Man2b2 | 1,18E-10 | 0,506227063 | 0,506 | 0,157 | 1,88E-06 | 1 | Man2b2 |
| Tifa | 1,21E-10 | 0,502810017 | 0,642 | 0,244 | 1,91E-06 | 1 | Tifa |
| Rin2 | 1,45E-10 | 0,398051064 | 0,568 | 0,202 | 2,30E-06 | 1 | Rin2 |
| Tyk2 | 1,70E-10 | 0,271148252 | 0,284 | 0,037 | 2,69E-06 | 1 | Tyk2 |
| Slc8b1 | 1,75E-10 | 0,341325667 | 0,333 | 0,062 | 2,77E-06 | 1 | Slc8b1 |
| Fam217b | 1,80E-10 | 0,342401797 | 0,333 | 0,062 | 2,86E-06 | 1 | Fam217b |
| Zbtb7b | 1,88E-10 | 0,270810632 | 0,358 | 0,074 | 2,97E-06 | 1 | Zbtb7b |
| Rps28 | 1,94E-10 | 0,320508052 | 1 | 0,992 | 3,07E-06 | 1 | Rps28 |
| Arpc1b | 1,96E-10 | 0,462137086 | 1 | 0,905 | 3,10E-06 | 1 | Arpc1b |
| Sfxn5 | 2,05E-10 | 0,284802255 | 0,358 | 0,074 | 3,25E-06 | 1 | Sfxn5 |
| Ptpn7 | 2,29E-10 | 0,339500348 | 0,346 | 0,066 | 3,62E-06 | 1 | Ptpn7 |
| Gpx1 | 2,33E-10 | 0,630117103 | 1 | 0,921 | 3,69E-06 | 1 | Gpx1 |
| Rab24 | 2,41E-10 | 0,510135775 | 0,617 | 0,269 | 3,83E-06 | 1 | Rab24 |
| Rogdi | 2,58E-10 | 0,497205869 | 0,58 | 0,223 | 4,10E-06 | 1 | Rogdi |
| Fnip1 | 2,94E-10 | 0,387543954 | 0,444 | 0,136 | 4,66E-06 | 1 | Fnip1 |
| Pold4 | 3,16E-10 | 0,331141164 | 0,568 | 0,215 | 5,01E-06 | 1 | Pold4 |
| Ypel3 | 3,32E-10 | 0,523306983 | 0,556 | 0,211 | 5,26E-06 | 1 | Ypel3 |
| Rpl10a | 3,35E-10 | 0,359997518 | 1 | 0,921 | 5,31E-06 | 1 | Rpl10a |

| | | | | | | | |
|---|---|---|---|---|---|---|---|
| Cerk | 0,001671041 | 0,255840249 | 0,543 | 0,347 | 1 | MAC | Cerk |
| Nubp1 | 0,001682947 | 0,352596296 | 0,444 | 0,289 | 1 | MAC | Nubp1 |
| Tuba1c | 0,001700921 | 0,299674033 | 0,79 | 0,603 | 1 | MAC | Tuba1c |
| Rrbp1 | 0,001727621 | 0,288315791 | 0,938 | 0,835 | 1 | MAC | Rrbp1 |
| Brk1 | 0,001770722 | 0,287077428 | 0,815 | 0,711 | 1 | MAC | Brk1 |
| Ifnar2 | 0,001816522 | 0,305014514 | 0,667 | 0,517 | 1 | MAC | Ifnar2 |
| Sirt2 | 0,001851002 | 0,299959535 | 0,531 | 0,355 | 1 | MAC | Sirt2 |
| Cisd2 | 0,001872841 | 0,258851665 | 0,506 | 0,326 | 1 | MAC | Cisd2 |
| Washc2 | 0,001874363 | 0,325287528 | 0,407 | 0,244 | 1 | MAC | Washc2 |
| Ndufb10 | 0,001968151 | 0,274319719 | 0,765 | 0,719 | 1 | MAC | Ndufb10 |
| Camta2 | 0,002125487 | 0,325067527 | 0,407 | 0,256 | 1 | MAC | Camta2 |
| Htra2 | 0,002204692 | 0,318091171 | 0,309 | 0,165 | 1 | MAC | Htra2 |
| Slc25a5 | 0,002585475 | 0,291702116 | 0,938 | 0,888 | 1 | MAC | Slc25a5 |
| Commd9 | 0,002827814 | 0,276233384 | 0,333 | 0,186 | 1 | MAC | Commd9 |
| Ndufs7 | 0,003125321 | 0,258165094 | 0,691 | 0,521 | 1 | MAC | Ndufs7 |
| Rab11fip5 | 0,003313717 | 0,259085413 | 0,457 | 0,302 | 1 | MAC | Rab11fip5 |
| Bnip3 | 0,003384699 | 0,287352749 | 0,346 | 0,174 | 1 | MAC | Bnip3 |
| Cyp4f16 | 0,003393531 | 0,274379499 | 0,259 | 0,132 | 1 | MAC | Cyp4f16 |
| Herpud1 | 0,00362017 | 0,32477929 | 0,568 | 0,393 | 1 | MAC | Herpud1 |
| Phip | 0,00378353 | 0,274333164 | 0,37 | 0,231 | 1 | MAC | Phip |
| Ier5 | 0,004077352 | 0,302594807 | 0,444 | 0,298 | 1 | MAC | Ier5 |
| Zfp810 | 0,004144817 | 0,273331743 | 0,259 | 0,128 | 1 | MAC | Zfp810 |
| Rpl22l1 | 0,004217647 | 0,283652359 | 0,864 | 0,76 | 1 | MAC | Rpl22l1 |
| Preb | 0,004552737 | 0,323296902 | 0,444 | 0,31 | 1 | MAC | Preb |
| Oas1a | 0,004839028 | 0,278052257 | 0,309 | 0,174 | 1 | MAC | Oas1a |
| Rabggta | 0,004937431 | 0,371226076 | 0,284 | 0,157 | 1 | MAC | Rabggta |
| Nr4a1 | 0,004988915 | 0,608428637 | 0,494 | 0,351 | 1 | MAC | Nr4a1 |
| Lamtor3 | 0,0053705 | 0,32271594 | 0,519 | 0,413 | 1 | MAC | Lamtor3 |
| Leng8 | 0,00618549 | 0,277088141 | 0,58 | 0,417 | 1 | MAC | Leng8 |
| Ifnar1 | 0,006287453 | 0,313146503 | 0,63 | 0,467 | 1 | MAC | Ifnar1 |
| Dnajc13 | 0,006490648 | 0,263354753 | 0,346 | 0,215 | 1 | MAC | Dnajc13 |
| Itpripl1 | 0,007411859 | 0,25218796 | 0,407 | 0,256 | 1 | MAC | Itpripl1 |
| Rbck1 | 0,008204591 | 0,273263975 | 0,481 | 0,343 | 1 | MAC | Rbck1 |
| Hat1 | 0,008668322 | 0,321692578 | 0,358 | 0,223 | 1 | MAC | Hat1 |
| Atp6v0d1 | 0,009870302 | 0,266712217 | 0,654 | 0,574 | 1 | MAC | Atp6v0d1 |
| Tmem252 | 3,15E-17 | 0,823016189 | 0,976 | 0,506 | 4,99E-13 | T_EC | Tmem252 |
| Tnfrsf9 | 2,25E-16 | 0,891224275 | 0,585 | 0,145 | 3,56E-12 | T_EC | Tnfrsf9 |
| Esam | 9,01E-16 | 0,762601895 | 0,988 | 0,647 | 1,43E-11 | T_EC | Esam |
| Cpe | 2,13E-15 | 0,87507239 | 0,939 | 0,477 | 3,38E-11 | T_EC | Cpe |
| Igfbp7 | 2,82E-15 | 0,747603266 | 0,988 | 0,693 | 4,47E-11 | T_EC | Igfbp7 |
| Tbx1 | 3,61E-15 | 0,482108457 | 0,402 | 0,054 | 5,72E-11 | T_EC | Tbx1 |
| Pdlim1 | 8,07E-15 | 0,721956251 | 0,976 | 0,614 | 1,28E-10 | T_EC | Pdlim1 |
| S100a11 | 2,54E-14 | 0,596465584 | 0,988 | 0,809 | 4,02E-10 | T_EC | S100a11 |
| Eef1a1 | 2,93E-14 | 0,359834991 | 1 | 1 | 4,65E-10 | T_EC | Eef1a1 |
| Cldn5 | 3,68E-14 | 0,960102031 | 0,963 | 0,627 | 5,84E-10 | T_EC | Cldn5 |
| Vwa1 | 6,38E-14 | 0,783973987 | 0,951 | 0,627 | 1,01E-09 | T_EC | Vwa1 |
| Igfbp3 | 1,41E-13 | 0,824192353 | 1 | 0,705 | 2,23E-09 | T_EC | Igfbp3 |
| Adcy4 | 4,13E-13 | 0,691609538 | 0,707 | 0,27 | 6,55E-09 | T_EC | Adcy4 |
| Tm4sf1 | 6,01E-13 | 0,714597809 | 0,988 | 0,639 | 9,53E-09 | T_EC | Tm4sf1 |
| Ramp2 | 6,13E-13 | 0,583302166 | 0,988 | 0,606 | 9,72E-09 | T_EC | Ramp2 |
| Fkbp1a | 6,85E-13 | 0,550680874 | 1 | 0,905 | 1,09E-08 | T_EC | Fkbp1a |
| Itm2a | 7,84E-13 | 0,96928811 | 0,488 | 0,124 | 1,24E-08 | T_EC | Itm2a |
| Sox17 | 1,38E-12 | 0,647802502 | 0,598 | 0,199 | 2,19E-08 | T_EC | Sox17 |
| Egfl7 | 1,43E-12 | 0,617074937 | 0,988 | 0,676 | 2,27E-08 | T_EC | Egfl7 |
| Car2 | 2,03E-12 | 0,577860939 | 0,5 | 0,133 | 3,22E-08 | T_EC | Car2 |
| Sox18 | 2,31E-12 | 0,735765542 | 0,902 | 0,573 | 3,67E-08 | T_EC | Sox18 |
| Dusp2 | 5,28E-12 | 0,618663687 | 0,744 | 0,328 | 8,37E-08 | T_EC | Dusp2 |
| Cttn | 8,56E-12 | 0,61601156 | 0,793 | 0,373 | 1,36E-07 | T_EC | Cttn |
| Prnp | 1,32E-11 | 0,648312265 | 0,927 | 0,552 | 2,10E-07 | T_EC | Prnp |
| Clec14a | 4,27E-11 | 0,727857027 | 0,854 | 0,452 | 6,78E-07 | T_EC | Clec14a |
| Tmem88 | 5,23E-11 | 0,52050525 | 0,951 | 0,61 | 8,29E-07 | T_EC | Tmem88 |
| Tagln2 | 5,27E-11 | 0,579656078 | 0,939 | 0,838 | 8,36E-07 | T_EC | Tagln2 |
| Gata2 | 7,44E-11 | 0,520428961 | 0,537 | 0,174 | 1,18E-06 | T_EC | Gata2 |
| Cd34 | 9,91E-11 | 0,546632245 | 0,988 | 0,664 | 1,57E-06 | T_EC | Cd34 |
| Thsd1 | 1,39E-10 | 0,569434592 | 0,756 | 0,365 | 2,20E-06 | T_EC | Thsd1 |
| Cxx1a | 3,95E-10 | 0,408081382 | 0,951 | 0,618 | 6,25E-06 | T_EC | Cxx1a |
| Impdh1 | 4,29E-10 | 0,506388904 | 0,744 | 0,373 | 6,80E-06 | T_EC | Impdh1 |
| Lysmd2 | 4,38E-10 | 0,405117354 | 0,683 | 0,27 | 6,94E-06 | T_EC | Lysmd2 |
| Grcc10 | 5,28E-10 | 0,37500351 | 0,976 | 0,685 | 8,37E-06 | T_EC | Grcc10 |
| Maoa | 6,21E-10 | 0,481987592 | 0,5 | 0,162 | 9,84E-06 | T_EC | Maoa |
| Armcx2 | 6,66E-10 | 0,417877115 | 0,549 | 0,195 | 1,06E-05 | T_EC | Armcx2 |
| Copb2 | 7,08E-10 | 0,620490804 | 0,72 | 0,407 | 1,12E-05 | T_EC | Copb2 |
| Adgrl4 | 7,36E-10 | 0,572549531 | 0,976 | 0,618 | 1,17E-05 | T_EC | Adgrl4 |
| Npdc1 | 1,18E-09 | 0,457827796 | 0,817 | 0,398 | 1,86E-05 | T_EC | Npdc1 |
| Abcg2 | 1,31E-09 | 0,556793794 | 0,854 | 0,506 | 2,08E-05 | T_EC | Abcg2 |
| Rhoc | 1,47E-09 | 0,532017079 | 0,963 | 0,651 | 2,32E-05 | T_EC | Rhoc |
| Csrp2 | 1,54E-09 | 0,329873245 | 0,622 | 0,274 | 2,44E-05 | T_EC | Csrp2 |
| Mpzl1 | 1,56E-09 | 0,439114944 | 0,878 | 0,49 | 2,48E-05 | T_EC | Mpzl1 |
| Scgb3a1 | 1,57E-09 | 1,06888038 | 0,524 | 0,183 | 2,49E-05 | T_EC | Scgb3a1 |
| Grap | 1,72E-09 | 0,508688325 | 0,756 | 0,382 | 2,73E-05 | T_EC | Grap |
| Rplp0 | 1,83E-09 | 0,360316676 | 1 | 0,979 | 2,91E-05 | T_EC | Rplp0 |
| Mxra8 | 1,98E-09 | 0,466805308 | 0,707 | 0,32 | 3,14E-05 | T_EC | Mxra8 |
| Stx6 | 2,07E-09 | 0,659899003 | 0,732 | 0,432 | 3,29E-05 | T_EC | Stx6 |
| Ccnd3 | 2,13E-09 | 0,524065264 | 0,841 | 0,531 | 3,37E-05 | T_EC | Ccnd3 |
| Slc2a1 | 2,15E-09 | 0,873112021 | 0,89 | 0,593 | 3,40E-05 | T_EC | Slc2a1 |
| Cxx1b | 2,41E-09 | 0,293357495 | 0,951 | 0,585 | 3,82E-05 | T_EC | Cxx1b |
| Rhoj | 2,69E-09 | 0,446409557 | 0,817 | 0,465 | 4,26E-05 | T_EC | Rhoj |
| Tnfaip1 | 2,89E-09 | 0,440492444 | 0,927 | 0,598 | 4,58E-05 | T_EC | Tnfaip1 |
| Vim | 3,14E-09 | 0,444108182 | 1 | 0,971 | 4,97E-05 | T_EC | Vim |
| Cebpd | 3,17E-09 | 0,634194324 | 0,695 | 0,394 | 5,02E-05 | T_EC | Cebpd |
| Pcdhgc3 | 3,84E-09 | 0,349896135 | 0,378 | 0,095 | 6,09E-05 | T_EC | Pcdhgc3 |
| Ifitm3 | 3,85E-09 | 0,412280928 | 1 | 0,934 | 6,10E-05 | T_EC | Ifitm3 |
| Actn4 | 3,94E-09 | 0,435539892 | 0,988 | 0,784 | 6,25E-05 | T_EC | Actn4 |
| Hspb8 | 4,86E-09 | 0,648008937 | 0,72 | 0,378 | 7,70E-05 | T_EC | Hspb8 |
| Itgb1 | 5,22E-09 | 0,554490641 | 1 | 0,921 | 8,27E-05 | T_EC | Itgb1 |
| Robo4 | 5,58E-09 | 0,416599959 | 0,89 | 0,49 | 8,84E-05 | T_EC | Robo4 |
| Rpl13 | 6,22E-09 | 0,296782766 | 1 | 0,988 | 9,86E-05 | T_EC | Rpl13 |
| Prkcdbp | 7,00E-09 | 0,475400162 | 0,988 | 0,635 | 0,00011103 | T_EC | Prkcdbp |
| St3gal6 | 7,04E-09 | 0,452014279 | 0,671 | 0,315 | 0,000111647 | T_EC | St3gal6 |
| Fkbp10 | 7,54E-09 | 0,552074387 | 0,744 | 0,402 | 0,000119564 | T_EC | Fkbp10 |
| Ctla2a | 9,64E-09 | 0,481874353 | 0,988 | 0,668 | 0,000152746 | T_EC | Ctla2a |
| Ly6a | 1,11E-08 | 0,518687329 | 0,963 | 0,718 | 0,000176545 | T_EC | Ly6a |
| Icam2 | 1,17E-08 | 0,424463218 | 0,902 | 0,56 | 0,000186182 | T_EC | Icam2 |
| Arhgef7 | 1,22E-08 | 0,419585112 | 0,768 | 0,444 | 0,000192617 | T_EC | Arhgef7 |
| Tpm1 | 1,29E-08 | 0,479282892 | 0,878 | 0,598 | 0,000204005 | T_EC | Tpm1 |
| Ly6e | 1,37E-08 | 0,425740322 | 0,988 | 0,992 | 0,000217867 | T_EC | Ly6e |
| H2-Q6 | 1,55E-08 | 0,52548706 | 0,695 | 0,365 | 0,000245518 | T_EC | H2-Q6 |
| Grrp1 | 1,64E-08 | 0,397779347 | 0,707 | 0,332 | 0,000260564 | T_EC | Grrp1 |
| Ldha | 1,86E-08 | 0,438139412 | 0,951 | 0,921 | 0,000295483 | T_EC | Ldha |
| Lsr | 1,98E-08 | 0,502316235 | 0,5 | 0,183 | 0,000314452 | T_EC | Lsr |
| Arhgef15 | 2,54E-08 | 0,308224159 | 0,841 | 0,498 | 0,000402936 | T_EC | Arhgef15 |
| Cd200 | 2,55E-08 | 0,464570118 | 0,976 | 0,676 | 0,000404286 | T_EC | Cd200 |
| Tbrg1 | 2,74E-08 | 0,464351755 | 0,768 | 0,432 | 0,000433976 | T_EC | Tbrg1 |
| Cnn3 | 2,80E-08 | 0,376790191 | 1 | 0,647 | 0,000443824 | T_EC | Cnn3 |
| S100a16 | 2,96E-08 | 0,423755033 | 0,927 | 0,593 | 0,000468723 | T_EC | S100a16 |
| Acer2 | 3,18E-08 | 0,51885655 | 0,622 | 0,274 | 0,000504194 | T_EC | Acer2 |
| Rbpms | 4,78E-08 | 0,3703281 | 0,927 | 0,573 | 0,000757378 | T_EC | Rbpms |
| Acvrl1 | 4,98E-08 | 0,527987203 | 0,841 | 0,527 | 0,000789819 | T_EC | Acvrl1 |
| Ly6c1 | 5,31E-08 | 0,491027113 | 0,927 | 0,548 | 0,000841357 | T_EC | Ly6c1 |
| Nectin2 | 6,19E-08 | 0,516664032 | 0,683 | 0,357 | 0,000980758 | T_EC | Nectin2 |

| | | | | | | | |
|---|---|---|---|---|---|---|---|
| Taldo1 | 3,49E-10 | 0,504992913 | 0,864 | 0,603 | 5,53E-06 | 1 | Taldo1 |
| Rpl26 | 3,70E-10 | 0,332330144 | 1 | 0,963 | 5,87E-06 | 1 | Rpl26 |
| Clcn7 | 4,20E-10 | 0,574743996 | 0,531 | 0,194 | 6,66E-06 | 1 | Clcn7 |
| Dnase1l1 | 4,32E-10 | 0,386661687 | 0,42 | 0,107 | 6,84E-06 | 1 | Dnase1l1 |
| Fkbp2 | 4,40E-10 | 0,409685297 | 0,716 | 0,393 | 6,98E-06 | 1 | Fkbp2 |
| AI413582 | 4,44E-10 | 0,435651638 | 0,58 | 0,26 | 7,03E-06 | 1 | AI413582 |
| Oas1g | 4,72E-10 | 0,308331101 | 0,284 | 0,041 | 7,48E-06 | 1 | Oas1g |
| Slc35f6 | 4,75E-10 | 0,266989958 | 0,407 | 0,095 | 7,53E-06 | 1 | Slc35f6 |
| Cited2 | 6,22E-10 | 0,261250454 | 0,333 | 0,066 | 9,85E-06 | 1 | Cited2 |
| Khk | 6,63E-10 | 0,268554416 | 0,259 | 0,033 | 1,05E-05 | 1 | Khk |
| Ppp1r18 | 7,91E-10 | 0,466495029 | 0,815 | 0,554 | 1,25E-05 | 1 | Ppp1r18 |
| Rassf2 | 8,37E-10 | 0,265859129 | 0,667 | 0,273 | 1,33E-05 | 1 | Rassf2 |
| Plekha2 | 9,11E-10 | 0,32913824 | 0,37 | 0,083 | 1,44E-05 | 1 | Plekha2 |
| Gabarap | 9,52E-10 | 0,465031578 | 0,975 | 0,806 | 1,51E-05 | 1 | Gabarap |
| Pisd-ps1 | 9,61E-10 | 0,415847684 | 0,568 | 0,252 | 1,52E-05 | 1 | Pisd-ps1 |
| Nfkbiz | 1,11E-09 | 0,803398477 | 0,63 | 0,298 | 1,76E-05 | 1 | Nfkbiz |
| Trim8 | 1,14E-09 | 0,316832618 | 0,58 | 0,236 | 1,81E-05 | 1 | Trim8 |
| Pacs2 | 1,15E-09 | 0,326344429 | 0,444 | 0,136 | 1,82E-05 | 1 | Pacs2 |
| Gga1 | 1,17E-09 | 0,287208463 | 0,531 | 0,19 | 1,86E-05 | 1 | Gga1 |
| Kdm7a | 1,18E-09 | 0,27440017 | 0,481 | 0,153 | 1,87E-05 | 1 | Kdm7a |
| Atp6v1b2 | 1,28E-09 | 0,636540955 | 0,642 | 0,31 | 2,04E-05 | 1 | Atp6v1b2 |
| Acox3 | 1,32E-09 | 0,447666567 | 0,432 | 0,124 | 2,10E-05 | 1 | Acox3 |
| Grb2 | 1,41E-09 | 0,41818104 | 0,679 | 0,339 | 2,24E-05 | 1 | Grb2 |
| Rab43 | 1,43E-09 | 0,444088343 | 0,531 | 0,207 | 2,26E-05 | 1 | Rab43 |
| Lpin2 | 1,53E-09 | 0,378320467 | 0,333 | 0,07 | 2,43E-05 | 1 | Lpin2 |
| Plgrkt | 1,56E-09 | 0,368980329 | 0,556 | 0,211 | 2,47E-05 | 1 | Plgrkt |
| Plekhm2 | 1,59E-09 | 0,410650711 | 0,296 | 0,054 | 2,51E-05 | 1 | Plekhm2 |
| Plin2 | 1,59E-09 | 0,542380816 | 0,691 | 0,31 | 2,53E-05 | 1 | Plin2 |
| Tmem104 | 1,75E-09 | 0,421002829 | 0,42 | 0,12 | 2,77E-05 | 1 | Tmem104 |
| Snx2 | 1,76E-09 | 0,596221547 | 0,827 | 0,483 | 2,79E-05 | 1 | Snx2 |
| Rilpl2 | 1,78E-09 | 0,530644209 | 0,444 | 0,145 | 2,82E-05 | 1 | Rilpl2 |
| Naip2 | 1,89E-09 | 0,308571636 | 0,407 | 0,107 | 3,00E-05 | 1 | Naip2 |
| Map3k8 | 2,12E-09 | 0,32260867 | 0,469 | 0,153 | 3,37E-05 | 1 | Map3k8 |
| Gm28437 | 2,31E-09 | 0,300053937 | 1 | 0,992 | 3,66E-05 | 1 | Gm28437 |
| Bmp2k | 2,51E-09 | 0,376162685 | 0,988 | 0,86 | 3,99E-05 | 1 | Bmp2k |
| Taok3 | 2,70E-09 | 0,356951209 | 0,457 | 0,149 | 4,28E-05 | 1 | Taok3 |
| Igsf8 | 2,87E-09 | 0,462153522 | 0,358 | 0,095 | 4,55E-05 | 1 | Igsf8 |
| 5031439G07Rik | 2,93E-09 | 0,332936568 | 0,506 | 0,178 | 4,64E-05 | 1 | 5031439G07Rik |
| Pqlc3 | 2,95E-09 | 0,351605941 | 0,296 | 0,058 | 4,68E-05 | 1 | Pqlc3 |
| Snx1 | 3,05E-09 | 0,556844509 | 0,704 | 0,368 | 4,83E-05 | 1 | Snx1 |
| Morc3 | 3,05E-09 | 0,385446898 | 0,481 | 0,157 | 4,84E-05 | 1 | Morc3 |
| Nfkbid | 3,23E-09 | 0,54807823 | 0,333 | 0,074 | 5,12E-05 | 1 | Nfkbid |
| Tmem176b | 3,96E-09 | 0,603594458 | 0,914 | 0,715 | 6,28E-05 | 1 | Tmem176b |
| Letmd1 | 4,13E-09 | 0,301215122 | 0,296 | 0,054 | 6,55E-05 | 1 | Letmd1 |
| Tmem219 | 4,71E-09 | 0,352341913 | 0,506 | 0,178 | 7,46E-05 | 1 | Tmem219 |
| Mfsd1 | 4,75E-09 | 0,562724908 | 0,815 | 0,508 | 7,54E-05 | 1 | Mfsd1 |
| Tmem268 | 4,91E-09 | 0,360747318 | 0,272 | 0,045 | 7,78E-05 | 1 | Tmem268 |
| Btg2 | 5,44E-09 | 0,763811883 | 0,716 | 0,438 | 8,62E-05 | 1 | Btg2 |
| Rps8 | 6,31E-09 | 0,414751109 | 1 | 0,975 | 0,000100056 | 1 | Rps8 |
| Man2a2 | 7,43E-09 | 0,305383657 | 0,321 | 0,07 | 0,000117767 | 1 | Man2a2 |
| Vps18 | 7,89E-09 | 0,482439892 | 0,358 | 0,099 | 0,000125044 | 1 | Vps18 |
| Rpl32 | 7,94E-09 | 0,289426801 | 1 | 0,988 | 0,00012583 | 1 | Rpl32 |
| Ap1g2 | 8,39E-09 | 0,535053461 | 0,259 | 0,045 | 0,000132948 | 1 | Ap1g2 |
| Arpc4 | 9,24E-09 | 0,379032921 | 1 | 0,884 | 0,000146448 | 1 | Arpc4 |
| Crlf3 | 1,03E-08 | 0,410666623 | 0,407 | 0,116 | 0,000163151 | 1 | Crlf3 |
| Arf6 | 1,07E-08 | 0,471831406 | 0,753 | 0,442 | 0,000169072 | 1 | Arf6 |
| Nupr1 | 1,10E-08 | 0,261345505 | 0,272 | 0,05 | 0,000174104 | 1 | Nupr1 |
| Aup1 | 1,23E-08 | 0,541470424 | 0,79 | 0,525 | 0,000194308 | 1 | Aup1 |
| Tsc22d4 | 1,27E-08 | 0,362072565 | 0,716 | 0,397 | 0,000201491 | 1 | Tsc22d4 |
| Mapkapk2 | 1,58E-08 | 0,31206058 | 0,642 | 0,281 | 0,000250506 | 1 | Mapkapk2 |
| Phf11a | 1,59E-08 | 0,261339435 | 0,259 | 0,045 | 0,000251722 | 1 | Phf11a |
| Pip4k2a | 1,65E-08 | 0,358965477 | 0,432 | 0,14 | 0,000261708 | 1 | Pip4k2a |
| Aftph | 2,43E-08 | 0,280698347 | 0,383 | 0,107 | 0,000384763 | 1 | Aftph |
| Rsrp1 | 2,47E-08 | 0,595035526 | 0,926 | 0,719 | 0,000391913 | 1 | Rsrp1 |
| Pak1 | 2,68E-08 | 0,266462141 | 0,333 | 0,079 | 0,000425143 | 1 | Pak1 |
| Ppp1r15a | 3,46E-08 | 0,574772393 | 0,654 | 0,318 | 0,000548282 | 1 | Ppp1r15a |
| Galc | 3,47E-08 | 0,255565691 | 0,346 | 0,095 | 0,000549302 | 1 | Galc |
| Fuca2 | 3,69E-08 | 0,434636214 | 0,506 | 0,194 | 0,000584657 | 1 | Fuca2 |
| Mthfs | 4,48E-08 | 0,259588896 | 0,469 | 0,169 | 0,000710056 | 1 | Mthfs |
| Serp1 | 4,57E-08 | 0,512257959 | 0,864 | 0,574 | 0,000724582 | 1 | Serp1 |
| Rragc | 4,73E-08 | 0,287752796 | 0,519 | 0,202 | 0,000750369 | 1 | Rragc |
| Il17ra | 4,94E-08 | 0,339720252 | 0,37 | 0,103 | 0,000782408 | 1 | Il17ra |
| Atp6ap2 | 5,13E-08 | 0,503285279 | 0,679 | 0,355 | 0,000813986 | 1 | Atp6ap2 |
| Cd274 | 5,17E-08 | 0,373104432 | 0,284 | 0,062 | 0,000819388 | 1 | Cd274 |
| Pdlim4 | 5,55E-08 | 0,338972994 | 0,309 | 0,074 | 0,000879132 | 1 | Pdlim4 |
| Cpne2 | 5,69E-08 | 0,331404293 | 0,593 | 0,277 | 0,000901295 | 1 | Cpne2 |
| Sfi1 | 5,70E-08 | 0,288851182 | 0,494 | 0,198 | 0,000903212 | 1 | Sfi1 |
| Dna2 | 5,92E-08 | 0,254922514 | 0,272 | 0,054 | 0,000938223 | 1 | Dna2 |
| Acsl1 | 6,01E-08 | 0,37854247 | 0,321 | 0,083 | 0,00095261 | 1 | Acsl1 |
| Tnfrsf1b | 6,71E-08 | 0,304839792 | 0,617 | 0,281 | 0,001064223 | 1 | Tnfrsf1b |
| Akr1b3 | 6,91E-08 | 0,337364873 | 0,716 | 0,364 | 0,001094718 | 1 | Akr1b3 |
| Ncoa3 | 7,04E-08 | 0,404264804 | 0,741 | 0,421 | 0,001115937 | 1 | Ncoa3 |
| Ifngr1 | 7,53E-08 | 0,574604344 | 0,889 | 0,657 | 0,001194152 | 1 | Ifngr1 |
| Gm42418 | 7,91E-08 | 0,324815373 | 0,753 | 0,496 | 0,001253297 | 1 | Gm42418 |
| Psmb9 | 8,54E-08 | 0,411952036 | 0,778 | 0,492 | 0,001353094 | 1 | Psmb9 |
| Parp1 | 8,68E-08 | 0,462397937 | 0,556 | 0,248 | 0,001375698 | 1 | Parp1 |
| Klf6 | 8,75E-08 | 0,559287297 | 0,827 | 0,554 | 0,00138672 | 1 | Klf6 |
| Tmem189 | 8,82E-08 | 0,460042083 | 0,531 | 0,244 | 0,001398114 | 1 | Tmem189 |
| Ndufa13 | 9,80E-08 | 0,39710895 | 0,864 | 0,707 | 0,001553524 | 1 | Ndufa13 |
| Cyth1 | 9,95E-08 | 0,385011682 | 0,568 | 0,256 | 0,001577463 | 1 | Cyth1 |
| Cdk9 | 1,03E-07 | 0,379559798 | 0,494 | 0,19 | 0,001625808 | 1 | Cdk9 |
| Rack1 | 1,04E-07 | 0,404986594 | 1 | 0,926 | 0,001650619 | 1 | Rack1 |
| Mapk14 | 1,27E-07 | 0,422128891 | 0,519 | 0,219 | 0,002005811 | 1 | Mapk14 |
| Gm7536 | 1,27E-07 | 0,341241622 | 0,988 | 0,938 | 0,002016701 | 1 | Gm7536 |
| Flcn | 1,40E-07 | 0,510609678 | 0,358 | 0,107 | 0,002213389 | 1 | Flcn |
| Rap2a | 1,40E-07 | 0,390838553 | 0,457 | 0,169 | 0,002222496 | 1 | Rap2a |
| Vps26a | 1,43E-07 | 0,368504367 | 0,58 | 0,277 | 0,002262571 | 1 | Vps26a |
| Slc25a45 | 1,52E-07 | 0,387565813 | 0,284 | 0,066 | 0,002408527 | 1 | Slc25a45 |
| Fes | 1,74E-07 | 0,400772623 | 0,691 | 0,335 | 0,002761886 | 1 | Fes |
| Dgkz | 1,78E-07 | 0,511542067 | 0,741 | 0,467 | 0,002822216 | 1 | Dgkz |
| Hspa1a | 1,80E-07 | 1,174612449 | 0,519 | 0,269 | 0,002846349 | 1 | Hspa1a |
| Tmem192 | 1,84E-07 | 0,479401824 | 0,407 | 0,153 | 0,002909046 | 1 | Tmem192 |
| Wdr81 | 1,87E-07 | 0,269206671 | 0,259 | 0,054 | 0,002968104 | 1 | Wdr81 |
| Dennd1a | 2,00E-07 | 0,251730345 | 0,432 | 0,161 | 0,0031665 | 1 | Dennd1a |
| Plekhb2 | 2,06E-07 | 0,297982996 | 0,469 | 0,178 | 0,00327252 | 1 | Plekhb2 |
| 2210016F16Rik | 2,19E-07 | 0,283348809 | 0,432 | 0,153 | 0,003478031 | 1 | 2210016F16Rik |
| Nceh1 | 2,29E-07 | 0,342073045 | 0,333 | 0,095 | 0,003625298 | 1 | Nceh1 |
| Gaa | 2,29E-07 | 0,32016142 | 0,444 | 0,161 | 0,003626059 | 1 | Gaa |
| Adam9 | 2,35E-07 | 0,360520503 | 0,79 | 0,467 | 0,003719169 | 1 | Adam9 |
| Rpl37a | 2,46E-07 | 0,253571596 | 1 | 0,996 | 0,003897212 | 1 | Rpl37a |
| Myo9b | 2,53E-07 | 0,399475533 | 0,58 | 0,285 | 0,004006585 | 1 | Myo9b |
| Mdfic | 2,71E-07 | 0,363253621 | 0,506 | 0,219 | 0,004294311 | 1 | Mdfic |
| Slc35c2 | 2,72E-07 | 0,33979996 | 0,432 | 0,165 | 0,004313519 | 1 | Slc35c2 |
| Rps13 | 2,80E-07 | 0,34570896 | 0,963 | 0,913 | 0,004438004 | 1 | Rps13 |
| Gm10275 | 2,81E-07 | 0,321386147 | 0,988 | 0,888 | 0,004455872 | 1 | Gm10275 |
| Tnfrsf21 | 2,84E-07 | 0,315534155 | 0,284 | 0,066 | 0,004505844 | 1 | Tnfrsf21 |
| Rps26 | 2,95E-07 | 0,256091399 | 1 | 0,992 | 0,004679862 | 1 | Rps26 |
| Rgs19 | 3,32E-07 | 0,343106929 | 0,568 | 0,269 | 0,00525813 | 1 | Rgs19 |
| Camk1 | 3,47E-07 | 0,562994961 | 0,716 | 0,417 | 0,005507566 | 1 | Camk1 |
| Akr1a1 | 3,52E-07 | 0,40842193 | 0,951 | 0,913 | 0,005578477 | 1 | Akr1a1 |

| | | | | | | | |
|---|---|---|---|---|---|---|---|
| Wfdc1 | 6,31E-08 | 0,432724312 | 0,256 | 0,05 | 0,000999998 | T_EC | Wfdc1 |
| Ecscr | 6,48E-08 | 0,374573642 | 0,976 | 0,647 | 0,001026418 | T_EC | Ecscr |
| Ushbp1 | 6,58E-08 | 0,399820977 | 0,78 | 0,473 | 0,001042486 | T_EC | Ushbp1 |
| Tspan3 | 7,71E-08 | 0,467716208 | 0,671 | 0,349 | 0,001222256 | T_EC | Tspan3 |
| 9430020K01Rik | 8,34E-08 | 0,303883539 | 0,915 | 0,481 | 0,001322416 | T_EC | 9430020K01Rik |
| Nedd4 | 8,37E-08 | 0,377398486 | 0,951 | 0,639 | 0,001326877 | T_EC | Nedd4 |
| Tsc22d1 | 8,45E-08 | 0,613130367 | 0,841 | 0,51 | 0,001339329 | T_EC | Tsc22d1 |
| Ptrf | 8,89E-08 | 0,406101603 | 0,963 | 0,614 | 0,001408555 | T_EC | Ptrf |
| Tmem43 | 9,59E-08 | 0,569684015 | 0,549 | 0,253 | 0,001520776 | T_EC | Tmem43 |
| Ptprk | 9,93E-08 | 0,315851266 | 0,732 | 0,373 | 0,001573465 | T_EC | Ptprk |
| H2-Q7 | 1,02E-07 | 0,529178343 | 0,854 | 0,589 | 0,001618428 | T_EC | H2-Q7 |
| Atp5b | 1,13E-07 | 0,441417165 | 0,951 | 0,909 | 0,001794007 | T_EC | Atp5b |
| Dstn | 1,19E-07 | 0,472931624 | 0,915 | 0,763 | 0,001882732 | T_EC | Dstn |
| Upp1 | 1,22E-07 | 0,436551436 | 0,902 | 0,535 | 0,001928108 | T_EC | Upp1 |
| Map4k5 | 1,23E-07 | 0,384573362 | 0,744 | 0,398 | 0,001944014 | T_EC | Map4k5 |
| Arf4 | 1,25E-07 | 0,469907911 | 0,927 | 0,751 | 0,001985047 | T_EC | Arf4 |
| Smtn | 1,39E-07 | 0,361939302 | 0,854 | 0,494 | 0,002197384 | T_EC | Smtn |
| Meox1 | 1,44E-07 | 0,471030642 | 0,695 | 0,386 | 0,002288262 | T_EC | Meox1 |
| Eif4g2 | 1,47E-07 | 0,407585471 | 0,976 | 0,88 | 0,002335678 | T_EC | Eif4g2 |
| Bcam | 1,49E-07 | 0,33290461 | 0,402 | 0,137 | 0,00236304 | T_EC | Bcam |
| Gm10123 | 1,51E-07 | 0,283411564 | 0,988 | 0,988 | 0,002391179 | T_EC | Gm10123 |
| Tax1bp3 | 1,58E-07 | 0,385392714 | 0,915 | 0,685 | 0,002498471 | T_EC | Tax1bp3 |
| Adgrg1 | 1,77E-07 | 0,4055812 | 0,707 | 0,382 | 0,002799458 | T_EC | Adgrg1 |
| Eif5a | 1,80E-07 | 0,423060832 | 0,988 | 0,929 | 0,002850584 | T_EC | Eif5a |
| Lef1 | 1,84E-07 | 0,322660703 | 0,341 | 0,104 | 0,002920113 | T_EC | Lef1 |
| Rmnd5b | 2,00E-07 | 0,33783044 | 0,561 | 0,245 | 0,003174285 | T_EC | Rmnd5b |
| Slc7a5 | 2,08E-07 | 0,330476085 | 0,5 | 0,191 | 0,0032964 | T_EC | Slc7a5 |
| Mcam | 2,14E-07 | 0,441792822 | 1 | 0,639 | 0,003392892 | T_EC | Mcam |
| Slc39a8 | 2,28E-07 | 0,475907959 | 0,488 | 0,207 | 0,003615648 | T_EC | Slc39a8 |
| Vwf | 2,36E-07 | 0,410254989 | 0,939 | 0,606 | 0,003739535 | T_EC | Vwf |
| Slco2b1 | 2,44E-07 | 0,600308719 | 0,646 | 0,386 | 0,003872263 | T_EC | Slco2b1 |
| Gimap5 | 2,50E-07 | 0,294991962 | 0,756 | 0,398 | 0,003960502 | T_EC | Gimap5 |
| Arf1 | 2,52E-07 | 0,365444011 | 0,988 | 0,921 | 0,00399371 | T_EC | Arf1 |
| Bok | 2,60E-07 | 0,449362284 | 0,634 | 0,315 | 0,004123365 | T_EC | Bok |
| Ankrd37 | 2,61E-07 | 0,538580249 | 0,561 | 0,278 | 0,00413013 | T_EC | Ankrd37 |
| Rela | 2,89E-07 | 0,426504701 | 0,72 | 0,448 | 0,004588992 | T_EC | Rela |
| Mall | 2,99E-07 | 0,262824575 | 0,561 | 0,286 | 0,004737591 | T_EC | Mall |
| Id3 | 3,00E-07 | 0,456418034 | 0,841 | 0,465 | 0,004754232 | T_EC | Id3 |
| Phb | 3,21E-07 | 0,291132383 | 0,78 | 0,502 | 0,005093723 | T_EC | Phb |
| Ercc1 | 3,30E-07 | 0,301939076 | 0,634 | 0,311 | 0,005226835 | T_EC | Ercc1 |
| Snrk | 3,33E-07 | 0,499739374 | 0,902 | 0,614 | 0,005284882 | T_EC | Snrk |
| Rasip1 | 3,36E-07 | 0,338623152 | 0,854 | 0,51 | 0,005319068 | T_EC | Rasip1 |
| Pdlim7 | 3,43E-07 | 0,39077543 | 0,915 | 0,664 | 0,005443408 | T_EC | Pdlim7 |
| Sparc | 3,48E-07 | 0,396511425 | 1 | 0,722 | 0,005510587 | T_EC | Sparc |
| Trp53i11 | 4,07E-07 | 0,62858834 | 0,841 | 0,61 | 0,006446555 | T_EC | Trp53i11 |
| Ocln | 4,08E-07 | 0,417400576 | 0,341 | 0,108 | 0,006474005 | T_EC | Ocln |
| Vcl | 4,18E-07 | 0,37650416 | 0,683 | 0,361 | 0,006626308 | T_EC | Vcl |
| Elk3 | 4,57E-07 | 0,306941257 | 1 | 0,759 | 0,007240884 | T_EC | Elk3 |
| Lta4h | 4,69E-07 | 0,330471934 | 0,732 | 0,419 | 0,0074344 | T_EC | Lta4h |
| Cdc42ep4 | 4,70E-07 | 0,372057946 | 0,524 | 0,237 | 0,007445985 | T_EC | Cdc42ep4 |
| Fam167b | 4,80E-07 | 0,372640579 | 0,805 | 0,469 | 0,007612037 | T_EC | Fam167b |
| Bcl6b | 5,28E-07 | 0,461664642 | 0,573 | 0,274 | 0,008375254 | T_EC | Bcl6b |
| Lypla1 | 5,81E-07 | 0,356400336 | 0,768 | 0,477 | 0,009202994 | T_EC | Lypla1 |
| Ufd1 | 5,88E-07 | 0,320563102 | 0,622 | 0,32 | 0,009314338 | T_EC | Ufd1 |
| Gcsh | 5,95E-07 | 0,4074231 | 0,561 | 0,299 | 0,009434681 | T_EC | Gcsh |
| Fam129a | 6,12E-07 | 0,308999188 | 0,427 | 0,158 | 0,009701833 | T_EC | Fam129a |
| Nostrin | 6,30E-07 | 0,294241202 | 0,817 | 0,473 | 0,009985131 | T_EC | Nostrin |
| Gimap4 | 6,80E-07 | 0,391484104 | 0,854 | 0,531 | 0,010778424 | T_EC | Gimap4 |
| Kank3 | 6,92E-07 | 0,51805767 | 0,671 | 0,415 | 0,010967245 | T_EC | Kank3 |
| Rab11a | 7,13E-07 | 0,413571596 | 0,976 | 0,809 | 0,011304786 | T_EC | Rab11a |
| Lpcat4 | 8,01E-07 | 0,265379089 | 0,39 | 0,141 | 0,012705171 | T_EC | Lpcat4 |
| Eif2s2 | 8,36E-07 | 0,301281907 | 0,951 | 0,813 | 0,013255135 | T_EC | Eif2s2 |
| Calu | 9,35E-07 | 0,436883317 | 0,927 | 0,697 | 0,014821579 | T_EC | Calu |
| Mgp | 9,41E-07 | 0,443514164 | 0,427 | 0,154 | 0,014913658 | T_EC | Mgp |
| Arrdc3 | 9,73E-07 | 0,562602754 | 0,573 | 0,286 | 0,015417812 | T_EC | Arrdc3 |
| Tspan13 | 9,80E-07 | 0,501905729 | 0,695 | 0,39 | 0,015538214 | T_EC | Tspan13 |
| Tuba1a | 1,02E-06 | 0,463288595 | 0,927 | 0,651 | 0,016166862 | T_EC | Tuba1a |
| Polr2b | 1,03E-06 | 0,282689312 | 0,78 | 0,502 | 0,016258881 | T_EC | Polr2b |
| Lgals9 | 1,04E-06 | 0,40672392 | 0,976 | 0,896 | 0,016482774 | T_EC | Lgals9 |
| Lpin3 | 1,06E-06 | 0,367443228 | 0,366 | 0,129 | 0,016761079 | T_EC | Lpin3 |
| Lhfp | 1,07E-06 | 0,33839898 | 0,268 | 0,071 | 0,016977114 | T_EC | Lhfp |
| Cfl2 | 1,12E-06 | 0,398700393 | 0,561 | 0,282 | 0,017691043 | T_EC | Cfl2 |
| Capg | 1,13E-06 | 0,402011624 | 0,951 | 0,718 | 0,017950629 | T_EC | Capg |
| Cnbp | 1,13E-06 | 0,367091774 | 0,976 | 0,88 | 0,017971806 | T_EC | Cnbp |
| Sept4 | 1,15E-06 | 0,454493291 | 0,659 | 0,365 | 0,018202075 | T_EC | Sept4 |
| Ick | 1,26E-06 | 0,333726464 | 0,463 | 0,199 | 0,019926274 | T_EC | Ick |
| Scarf1 | 1,26E-06 | 0,36600928 | 0,646 | 0,324 | 0,019999257 | T_EC | Scarf1 |
| Chchd2 | 1,28E-06 | 0,263087323 | 0,951 | 0,909 | 0,020302137 | T_EC | Chchd2 |
| Csrp1 | 1,45E-06 | 0,416660052 | 0,744 | 0,506 | 0,022992559 | T_EC | Csrp1 |
| Rrp1 | 1,57E-06 | 0,408661767 | 0,744 | 0,485 | 0,024857986 | T_EC | Rrp1 |
| Lypla2 | 1,59E-06 | 0,266428752 | 0,707 | 0,419 | 0,025189228 | T_EC | Lypla2 |
| Nid2 | 1,62E-06 | 0,476300785 | 0,89 | 0,535 | 0,025743542 | T_EC | Nid2 |
| Actg1 | 1,64E-06 | 0,38529985 | 1 | 0,992 | 0,025974992 | T_EC | Actg1 |
| Arhgap29 | 1,72E-06 | 0,376996216 | 0,817 | 0,469 | 0,027234926 | T_EC | Arhgap29 |
| Eogt | 1,81E-06 | 0,449450132 | 0,659 | 0,353 | 0,028617829 | T_EC | Eogt |
| Tubb6 | 1,88E-06 | 0,427891917 | 0,927 | 0,676 | 0,029815818 | T_EC | Tubb6 |
| Fermt2 | 1,94E-06 | 0,315703894 | 0,829 | 0,527 | 0,030811613 | T_EC | Fermt2 |
| Pros1 | 1,99E-06 | 0,334148384 | 0,72 | 0,411 | 0,031473423 | T_EC | Pros1 |
| Trib2 | 2,08E-06 | 0,359739278 | 0,451 | 0,191 | 0,032996176 | T_EC | Trib2 |
| Prss23 | 2,11E-06 | 0,445629851 | 0,878 | 0,531 | 0,033396001 | T_EC | Prss23 |
| Rpl6 | 2,39E-06 | 0,250494547 | 1 | 0,992 | 0,037963497 | T_EC | Rpl6 |
| Rnf14 | 2,42E-06 | 0,353093283 | 0,561 | 0,286 | 0,038352753 | T_EC | Rnf14 |
| Crtap | 2,44E-06 | 0,309281676 | 0,707 | 0,402 | 0,038634598 | T_EC | Crtap |
| Cdh2 | 2,46E-06 | 0,259995317 | 0,378 | 0,137 | 0,038928064 | T_EC | Cdh2 |
| Fam43a | 2,52E-06 | 0,382195557 | 0,659 | 0,361 | 0,039878095 | T_EC | Fam43a |
| Eef1g | 2,68E-06 | 0,373874095 | 0,963 | 0,809 | 0,042554735 | T_EC | Eef1g |
| Anxa7 | 2,82E-06 | 0,402971845 | 0,78 | 0,535 | 0,044773648 | T_EC | Anxa7 |
| Afap1l1 | 2,86E-06 | 0,390573585 | 0,732 | 0,436 | 0,045402542 | T_EC | Afap1l1 |
| Tmem44 | 3,08E-06 | 0,380382786 | 0,341 | 0,12 | 0,048754327 | T_EC | Tmem44 |
| Sh2d3c | 3,17E-06 | 0,357489871 | 0,805 | 0,548 | 0,050241955 | T_EC | Sh2d3c |
| Eif4a-ps4 | 3,60E-06 | 0,324428056 | 0,732 | 0,535 | 0,057004889 | T_EC | Eif4a-ps4 |
| Clic4 | 3,80E-06 | 0,392606752 | 0,951 | 0,817 | 0,060306894 | T_EC | Clic4 |
| Trp53 | 3,92E-06 | 0,34729513 | 0,732 | 0,481 | 0,062213779 | T_EC | Trp53 |
| Tnfaip8l1 | 4,16E-06 | 0,296409325 | 0,561 | 0,274 | 0,065999638 | T_EC | Tnfaip8l1 |
| Gabpb1 | 4,21E-06 | 0,27264265 | 0,5 | 0,228 | 0,066720788 | T_EC | Gabpb1 |
| Myo1b | 4,34E-06 | 0,365802447 | 0,78 | 0,494 | 0,068796564 | T_EC | Myo1b |
| Rps6-ps4 | 4,35E-06 | 0,255016402 | 0,988 | 0,959 | 0,069008103 | T_EC | Rps6-ps4 |
| Fyttd1 | 4,44E-06 | 0,262487762 | 0,646 | 0,357 | 0,070314408 | T_EC | Fyttd1 |
| Hmgn3 | 4,72E-06 | 0,250565336 | 0,573 | 0,274 | 0,074826717 | T_EC | Hmgn3 |
| Osbpl9 | 4,90E-06 | 0,385795094 | 0,756 | 0,548 | 0,077647563 | T_EC | Osbpl9 |
| Rpl15-ps3 | 5,23E-06 | 0,270308594 | 0,927 | 0,909 | 0,082863051 | T_EC | Rpl15-ps3 |
| Ica1 | 5,23E-06 | 0,302138386 | 0,61 | 0,336 | 0,082897974 | T_EC | Ica1 |
| Rps2 | 5,42E-06 | 0,289742078 | 0,988 | 0,988 | 0,085927405 | T_EC | Rps2 |
| Id1 | 5,43E-06 | 0,492030735 | 0,72 | 0,448 | 0,086100989 | T_EC | Id1 |
| Cav1 | 5,62E-06 | 0,54487878 | 0,72 | 0,477 | 0,089069239 | T_EC | Cav1 |
| AU021092 | 5,63E-06 | 0,257500161 | 0,476 | 0,199 | 0,089170177 | T_EC | AU021092 |
| Tek | 5,78E-06 | 0,252081806 | 0,768 | 0,432 | 0,091612331 | T_EC | Tek |
| Trim47 | 6,03E-06 | 0,346110491 | 0,476 | 0,212 | 0,09560695 | T_EC | Trim47 |
| Fbxw9 | 6,06E-06 | 0,258548741 | 0,317 | 0,108 | 0,096109653 | T_EC | Fbxw9 |
| Bambi | 6,07E-06 | 0,355276422 | 0,537 | 0,257 | 0,096158928 | T_EC | Bambi |

| | | | | | | | |
|---|---|---|---|---|---|---|---|
| Nptn | 3,60E-07 | 0,474010964 | 0,716 | 0,475 | 0,005704295 | 1 | Nptn |
| Dpp7 | 3,78E-07 | 0,264869139 | 0,284 | 0,07 | 0,005995632 | 1 | Dpp7 |
| Tpp1 | 4,17E-07 | 0,450689817 | 0,79 | 0,517 | 0,006611116 | 1 | Tpp1 |
| Dleu2 | 4,20E-07 | 0,322452128 | 0,605 | 0,306 | 0,006653505 | 1 | Dleu2 |
| Pigs | 4,45E-07 | 0,456908012 | 0,519 | 0,227 | 0,007060433 | 1 | Pigs |
| Cdt1 | 4,85E-07 | 0,251512871 | 0,346 | 0,103 | 0,007682716 | 1 | Cdt1 |
| Taf6l | 4,86E-07 | 0,363202758 | 0,593 | 0,293 | 0,007709968 | 1 | Taf6l |
| Cox14 | 5,11E-07 | 0,342245265 | 0,691 | 0,45 | 0,008108268 | 1 | Cox14 |
| Blvra | 5,15E-07 | 0,309984188 | 0,704 | 0,376 | 0,008163604 | 1 | Blvra |
| Gsto1 | 5,27E-07 | 0,388805125 | 0,728 | 0,446 | 0,008359742 | 1 | Gsto1 |
| Ssh2 | 5,30E-07 | 0,566563372 | 0,753 | 0,492 | 0,008408263 | 1 | Ssh2 |
| Stat6 | 5,37E-07 | 0,46673398 | 0,543 | 0,244 | 0,008511975 | 1 | Stat6 |
| Crip1 | 5,43E-07 | 0,577245067 | 0,765 | 0,554 | 0,008613079 | 1 | Crip1 |
| Btg1 | 5,49E-07 | 0,45796331 | 0,938 | 0,789 | 0,008706478 | 1 | Btg1 |
| Scand1 | 5,67E-07 | 0,281490816 | 0,741 | 0,521 | 0,008985107 | 1 | Scand1 |
| Vrk2 | 6,35E-07 | 0,279880631 | 0,259 | 0,062 | 0,010059752 | 1 | Vrk2 |
| Npc1 | 7,38E-07 | 0,275798249 | 0,395 | 0,14 | 0,01169185 | 1 | Npc1 |
| Atp6v1a | 8,23E-07 | 0,420850531 | 0,691 | 0,388 | 0,013040315 | 1 | Atp6v1a |
| Rpl21 | 9,14E-07 | 0,302270835 | 1 | 0,963 | 0,014486009 | 1 | Rpl21 |
| Gm26917 | 9,87E-07 | 0,334005648 | 0,988 | 0,855 | 0,015653415 | 1 | Gm26917 |
| Htatip2 | 1,00E-06 | 0,316806292 | 0,407 | 0,149 | 0,015872115 | 1 | Htatip2 |
| Pmepa1 | 1,03E-06 | 0,375002635 | 0,79 | 0,521 | 0,016399906 | 1 | Pmepa1 |
| Cnp | 1,04E-06 | 0,291586412 | 0,407 | 0,145 | 0,016449048 | 1 | Cnp |
| mt-Atp6 | 1,10E-06 | 0,306721723 | 1 | 0,983 | 0,017496445 | 1 | mt-Atp6 |
| G6pdx | 1,14E-06 | 0,423892542 | 0,395 | 0,149 | 0,018137153 | 1 | G6pdx |
| Atp5e | 1,20E-06 | 0,314681132 | 0,975 | 0,864 | 0,018985818 | 1 | Atp5e |
| AC121965.1 | 1,26E-06 | 0,29458315 | 0,901 | 0,764 | 0,020048374 | 1 | AC121965.1 |
| Cox8a | 1,38E-06 | 0,332429412 | 0,951 | 0,88 | 0,021860291 | 1 | Cox8a |
| Cap1 | 1,39E-06 | 0,335142778 | 0,938 | 0,76 | 0,021984693 | 1 | Cap1 |
| Tex10 | 1,46E-06 | 0,309308564 | 0,432 | 0,178 | 0,023171982 | 1 | Tex10 |
| Edem1 | 1,59E-06 | 0,306618845 | 0,654 | 0,355 | 0,025200255 | 1 | Edem1 |
| Actb | 1,60E-06 | 0,275931915 | 1 | 1 | 0,025410614 | 1 | Actb |
| Per1 | 1,67E-06 | 0,274415015 | 0,617 | 0,331 | 0,026488303 | 1 | Per1 |
| Slc36a1 | 1,74E-06 | 0,320110234 | 0,259 | 0,066 | 0,027607457 | 1 | Slc36a1 |
| Gnpda1 | 1,75E-06 | 0,525451447 | 0,383 | 0,149 | 0,027700075 | 1 | Gnpda1 |
| 2810474O19Rik | 1,83E-06 | 0,340457258 | 0,679 | 0,393 | 0,028980775 | 1 | 2810474O19Rik |
| Txnip | 1,88E-06 | 0,250458743 | 0,741 | 0,409 | 0,029872936 | 1 | Txnip |
| Gba | 1,90E-06 | 0,421069872 | 0,593 | 0,302 | 0,030151438 | 1 | Gba |
| Dtnbp1 | 1,99E-06 | 0,264801413 | 0,358 | 0,124 | 0,031552819 | 1 | Dtnbp1 |
| Naglu | 2,11E-06 | 0,336616854 | 0,395 | 0,153 | 0,03342043 | 1 | Naglu |
| Cox6b1 | 2,28E-06 | 0,31456173 | 0,988 | 0,926 | 0,036129408 | 1 | Cox6b1 |
| Prex1 | 2,29E-06 | 0,477377259 | 0,679 | 0,409 | 0,036277186 | 1 | Prex1 |
| Fam50a | 2,29E-06 | 0,271526568 | 0,37 | 0,136 | 0,036346359 | 1 | Fam50a |
| Hn1 | 2,33E-06 | 0,448412304 | 0,827 | 0,55 | 0,036906254 | 1 | Hn1 |
| Tgif1 | 2,42E-06 | 0,377933745 | 0,605 | 0,302 | 0,038291948 | 1 | Tgif1 |
| Gapdh | 2,46E-06 | 0,278842381 | 1 | 0,988 | 0,039008427 | 1 | Gapdh |
| M6pr | 2,50E-06 | 0,432072169 | 0,84 | 0,558 | 0,039681663 | 1 | M6pr |
| Anxa4 | 2,56E-06 | 0,331962501 | 0,593 | 0,31 | 0,040586703 | 1 | Anxa4 |
| Mrps36 | 2,58E-06 | 0,268668285 | 0,506 | 0,273 | 0,040968991 | 1 | Mrps36 |
| Rnf19b | 2,59E-06 | 0,340871007 | 0,395 | 0,153 | 0,041126594 | 1 | Rnf19b |
| Atp6ap1 | 2,62E-06 | 0,355720406 | 0,827 | 0,512 | 0,041529857 | 1 | Atp6ap1 |
| Tmem176a | 2,67E-06 | 0,525873627 | 0,778 | 0,607 | 0,042316403 | 1 | Tmem176a |
| Rcbtb2 | 2,70E-06 | 0,374847106 | 0,506 | 0,236 | 0,042797029 | 1 | Rcbtb2 |
| Bcl2l1 | 2,88E-06 | 0,50225105 | 0,58 | 0,322 | 0,045626493 | 1 | Bcl2l1 |
| Trafd1 | 3,02E-06 | 0,514206768 | 0,605 | 0,331 | 0,047899631 | 1 | Trafd1 |
| Nhlrc3 | 3,26E-06 | 0,349961382 | 0,432 | 0,182 | 0,051735818 | 1 | Nhlrc3 |
| Wipf1 | 3,68E-06 | 0,36239454 | 0,556 | 0,322 | 0,058387708 | 1 | Wipf1 |
| Cnpy3 | 4,09E-06 | 0,382239611 | 0,543 | 0,273 | 0,064770194 | 1 | Cnpy3 |
| Elf4 | 4,16E-06 | 0,25896207 | 0,346 | 0,12 | 0,06601765 | 1 | Elf4 |
| Actr2 | 4,63E-06 | 0,25243645 | 1 | 0,938 | 0,073358429 | 1 | Actr2 |
| Rps27a | 4,72E-06 | 0,266554083 | 1 | 0,975 | 0,074895431 | 1 | Rps27a |
| Csf2rb | 4,83E-06 | 0,414080554 | 0,593 | 0,322 | 0,076625401 | 1 | Csf2rb |
| Ddt | 4,91E-06 | 0,255558656 | 0,407 | 0,169 | 0,077902543 | 1 | Ddt |
| Rnh1 | 5,07E-06 | 0,37633876 | 0,815 | 0,496 | 0,080439727 | 1 | Rnh1 |
| Zfp622 | 5,08E-06 | 0,276062419 | 0,407 | 0,161 | 0,080573376 | 1 | Zfp622 |
| Tor1a | 5,15E-06 | 0,377415083 | 0,531 | 0,285 | 0,081602432 | 1 | Tor1a |
| Smpdl3a | 5,91E-06 | 0,27364814 | 0,457 | 0,186 | 0,093753494 | 1 | Smpdl3a |
| Pacsin2 | 6,49E-06 | 0,301107499 | 0,642 | 0,335 | 0,102865868 | 1 | Pacsin2 |
| Cyb5r4 | 6,78E-06 | 0,31605335 | 0,457 | 0,202 | 0,107544474 | 1 | Cyb5r4 |
| Ubl3 | 6,98E-06 | 0,42328339 | 0,778 | 0,525 | 0,110574542 | 1 | Ubl3 |
| Tmem55b | 7,18E-06 | 0,369626709 | 0,457 | 0,223 | 0,113858887 | 1 | Tmem55b |
| Pip5k1c | 8,09E-06 | 0,449937828 | 0,593 | 0,36 | 0,128269372 | 1 | Pip5k1c |
| Lifr | 8,29E-06 | 0,374037249 | 0,321 | 0,116 | 0,131471297 | 1 | Lifr |
| Mfsd11 | 8,59E-06 | 0,310880442 | 0,333 | 0,12 | 0,136236867 | 1 | Mfsd11 |
| Irak4 | 8,73E-06 | 0,261272298 | 0,358 | 0,14 | 0,138343019 | 1 | Irak4 |
| Ccdc115 | 8,80E-06 | 0,328650839 | 0,481 | 0,231 | 0,139516881 | 1 | Ccdc115 |
| Glrx | 8,93E-06 | 0,394557061 | 0,617 | 0,335 | 0,141488708 | 1 | Glrx |
| Ddx28 | 9,28E-06 | 0,337459919 | 0,259 | 0,079 | 0,147046663 | 1 | Ddx28 |
| Ndufa2 | 9,67E-06 | 0,348987899 | 0,951 | 0,868 | 0,153322264 | 1 | Ndufa2 |
| Ncapg2 | 9,76E-06 | 0,415761684 | 0,346 | 0,136 | 0,15469208 | 1 | Ncapg2 |
| Kxd1 | 9,89E-06 | 0,287555683 | 0,494 | 0,236 | 0,156810509 | 1 | Kxd1 |
| Ms4a6d | 9,95E-06 | 0,449845956 | 0,914 | 0,777 | 0,157661587 | 1 | Ms4a6d |
| Psmb10 | 1,01E-05 | 0,351103358 | 0,716 | 0,463 | 0,160047223 | 1 | Psmb10 |
| Ccnl1 | 1,04E-05 | 0,399996713 | 0,654 | 0,409 | 0,164085963 | 1 | Ccnl1 |
| Cuta | 1,14E-05 | 0,257625009 | 0,679 | 0,446 | 0,1807053 | 1 | Cuta |
| Siva1 | 1,17E-05 | 0,369090619 | 0,395 | 0,169 | 0,185836272 | 1 | Siva1 |
| Arrdc1 | 1,24E-05 | 0,35977545 | 0,568 | 0,306 | 0,196907732 | 1 | Arrdc1 |
| Nfil3 | 1,32E-05 | 0,374162748 | 0,272 | 0,087 | 0,209915239 | 1 | Nfil3 |
| Adam17 | 1,39E-05 | 0,375295267 | 0,58 | 0,343 | 0,220950103 | 1 | Adam17 |
| Acer3 | 1,43E-05 | 0,432093874 | 0,481 | 0,24 | 0,226900967 | 1 | Acer3 |
| Smap2 | 1,58E-05 | 0,283444208 | 0,543 | 0,281 | 0,250145372 | 1 | Smap2 |
| BC005537 | 1,59E-05 | 0,463756994 | 0,691 | 0,442 | 0,251665269 | 1 | BC005537 |
| Rps12l1 | 1,59E-05 | 0,313469105 | 0,963 | 0,855 | 0,252374209 | 1 | Rps12l1 |
| Mif | 1,65E-05 | 0,350488226 | 0,963 | 0,814 | 0,261009391 | 1 | Mif |
| Bmyc | 1,74E-05 | 0,312660213 | 0,432 | 0,207 | 0,276256042 | 1 | Bmyc |
| Acp2 | 1,80E-05 | 0,38597389 | 0,432 | 0,194 | 0,285111216 | 1 | Acp2 |
| Pabpc1 | 1,89E-05 | 0,250776888 | 0,988 | 0,942 | 0,300015297 | 1 | Pabpc1 |
| Rhbdf2 | 2,17E-05 | 0,36100564 | 0,296 | 0,107 | 0,344556075 | 1 | Rhbdf2 |
| Heatr5a | 2,27E-05 | 0,404769656 | 0,37 | 0,161 | 0,359961722 | 1 | Heatr5a |
| Rnf13 | 2,45E-05 | 0,315453115 | 0,654 | 0,368 | 0,388255311 | 1 | Rnf13 |
| Tmem14c | 2,46E-05 | 0,311756394 | 0,765 | 0,537 | 0,390646008 | 1 | Tmem14c |
| Wdr26 | 2,56E-05 | 0,353666909 | 0,691 | 0,438 | 0,405755681 | 1 | Wdr26 |
| Mon2 | 2,74E-05 | 0,256787941 | 0,333 | 0,128 | 0,435091688 | 1 | Mon2 |
| Ifi211 | 3,16E-05 | 0,555267337 | 0,358 | 0,157 | 0,500438229 | 1 | Ifi211 |
| Spg21 | 3,16E-05 | 0,253956438 | 0,63 | 0,347 | 0,501644693 | 1 | Spg21 |
| Got1 | 3,17E-05 | 0,315694764 | 0,333 | 0,128 | 0,502705264 | 1 | Got1 |
| Fkbp5 | 3,30E-05 | 0,326733948 | 0,704 | 0,488 | 0,522835258 | 1 | Fkbp5 |
| Rpl17 | 3,31E-05 | 0,284752912 | 0,988 | 0,905 | 0,525460259 | 1 | Rpl17 |
| Tab2 | 3,32E-05 | 0,297073291 | 0,494 | 0,26 | 0,526972407 | 1 | Tab2 |
| Rbfa | 3,38E-05 | 0,293478347 | 0,42 | 0,19 | 0,536569295 | 1 | Rbfa |
| Acaa1a | 4,03E-05 | 0,288458877 | 0,543 | 0,289 | 0,638575488 | 1 | Acaa1a |
| Rab8b | 4,27E-05 | 0,353639868 | 0,617 | 0,38 | 0,677268308 | 1 | Rab8b |
| Zmiz2 | 4,58E-05 | 0,373698644 | 0,481 | 0,264 | 0,726072889 | 1 | Zmiz2 |
| Il10rb | 4,78E-05 | 0,392888027 | 0,667 | 0,426 | 0,757823196 | 1 | Il10rb |
| Stk24 | 4,80E-05 | 0,323531773 | 0,37 | 0,169 | 0,761298812 | 1 | Stk24 |
| Mcl1 | 4,82E-05 | 0,321860004 | 0,84 | 0,653 | 0,763520793 | 1 | Mcl1 |
| Haus8 | 4,93E-05 | 0,280554427 | 0,272 | 0,099 | 0,780957369 | 1 | Haus8 |
| Gna13 | 4,97E-05 | 0,347415731 | 0,691 | 0,467 | 0,787658785 | 1 | Gna13 |
| Rpl7a | 5,03E-05 | 0,271818892 | 1 | 0,955 | 0,796827427 | 1 | Rpl7a |
| Srsf9 | 5,06E-05 | 0,278987998 | 0,605 | 0,397 | 0,801722727 | 1 | Srsf9 |

| | | | | | | | |
|---|---|---|---|---|---|---|---|
| Sema6d | 6,28E-06 | 0,250383969 | 0,89 | 0,535 | 0,099492562 | T_EC | Sema6d |
| Cyp2j6 | 6,30E-06 | 0,291832023 | 0,305 | 0,1 | 0,099811296 | T_EC | Cyp2j6 |
| Tm9sf2 | 6,50E-06 | 0,346984137 | 0,915 | 0,71 | 0,1030946 | T_EC | Tm9sf2 |
| Bhlhe40 | 6,60E-06 | 0,52660806 | 0,78 | 0,564 | 0,104577512 | T_EC | Bhlhe40 |
| Alas1 | 6,99E-06 | 0,4465916 | 0,622 | 0,344 | 0,110805631 | T_EC | Alas1 |
| Plk2 | 7,42E-06 | 0,390617344 | 0,927 | 0,676 | 0,117610059 | T_EC | Plk2 |
| Slc25a3 | 7,54E-06 | 0,348393432 | 0,939 | 0,867 | 0,119475447 | T_EC | Slc25a3 |
| Ppic | 7,65E-06 | 0,353644682 | 0,939 | 0,647 | 0,121338104 | T_EC | Ppic |
| Lrrc8c | 7,79E-06 | 0,380456012 | 0,793 | 0,556 | 0,123547509 | T_EC | Lrrc8c |
| Rars | 7,85E-06 | 0,308588328 | 0,549 | 0,278 | 0,124426846 | T_EC | Rars |
| Tie1 | 7,88E-06 | 0,301167906 | 0,866 | 0,56 | 0,124915876 | T_EC | Tie1 |
| Ackr2 | 8,54E-06 | 0,41685338 | 0,512 | 0,257 | 0,13539986 | T_EC | Ackr2 |
| Igfbp4 | 9,01E-06 | 0,416875378 | 0,927 | 0,734 | 0,142789163 | T_EC | Igfbp4 |
| Ddah2 | 9,24E-06 | 0,299551222 | 0,683 | 0,378 | 0,14645499 | T_EC | Ddah2 |
| Flt1 | 9,25E-06 | 0,398640498 | 0,963 | 0,647 | 0,146588291 | T_EC | Flt1 |
| Arl1 | 9,29E-06 | 0,308389944 | 0,756 | 0,502 | 0,147318107 | T_EC | Arl1 |
| Cd320 | 9,32E-06 | 0,299545717 | 0,341 | 0,129 | 0,147747552 | T_EC | Cd320 |
| Ubl7 | 9,37E-06 | 0,296953334 | 0,61 | 0,361 | 0,148473301 | T_EC | Ubl7 |
| Prkd2 | 9,49E-06 | 0,282254578 | 0,671 | 0,361 | 0,150403522 | T_EC | Prkd2 |
| Spata6 | 9,69E-06 | 0,262951903 | 0,598 | 0,307 | 0,153623506 | T_EC | Spata6 |
| Atp5c1 | 9,81E-06 | 0,376896725 | 0,915 | 0,697 | 0,155486894 | T_EC | Atp5c1 |
| Psmc3 | 1,02E-05 | 0,348102301 | 0,854 | 0,739 | 0,161390505 | T_EC | Psmc3 |
| Flnb | 1,04E-05 | 0,262901826 | 0,78 | 0,465 | 0,164142374 | T_EC | Flnb |
| Mrpl17 | 1,08E-05 | 0,305747555 | 0,902 | 0,656 | 0,170937975 | T_EC | Mrpl17 |
| Adamts1 | 1,14E-05 | 0,510500445 | 0,78 | 0,498 | 0,180625804 | T_EC | Adamts1 |
| Cops5 | 1,18E-05 | 0,425545605 | 0,622 | 0,39 | 0,187418596 | T_EC | Cops5 |
| Fzd6 | 1,23E-05 | 0,250378561 | 0,488 | 0,237 | 0,195120714 | T_EC | Fzd6 |
| Tmsb10 | 1,24E-05 | 0,305035649 | 1 | 0,967 | 0,19645351 | T_EC | Tmsb10 |
| Slc9a3r2 | 1,25E-05 | 0,325462082 | 0,841 | 0,552 | 0,19769553 | T_EC | Slc9a3r2 |
| Tsen34 | 1,26E-05 | 0,349061774 | 0,683 | 0,436 | 0,200244423 | T_EC | Tsen34 |
| B3gnt3 | 1,29E-05 | 0,306303045 | 0,793 | 0,485 | 0,204800323 | T_EC | B3gnt3 |
| Slc25a4 | 1,32E-05 | 0,32803961 | 0,939 | 0,822 | 0,209539462 | T_EC | Slc25a4 |
| Myl12a | 1,34E-05 | 0,251440614 | 1 | 0,954 | 0,213053751 | T_EC | Myl12a |
| Lxn | 1,36E-05 | 0,299701885 | 0,854 | 0,564 | 0,215230097 | T_EC | Lxn |
| Palmd | 1,44E-05 | 0,404722203 | 0,634 | 0,382 | 0,227967763 | T_EC | Palmd |
| Serinc3 | 1,48E-05 | 0,265767965 | 0,988 | 0,983 | 0,234017429 | T_EC | Serinc3 |
| Slc30a4 | 1,48E-05 | 0,377173272 | 0,354 | 0,149 | 0,234028836 | T_EC | Slc30a4 |
| Tspan15 | 1,49E-05 | 0,266362726 | 0,573 | 0,299 | 0,235735297 | T_EC | Tspan15 |
| Gipc1 | 1,54E-05 | 0,309300611 | 0,72 | 0,448 | 0,243355619 | T_EC | Gipc1 |
| Ndufs2 | 1,60E-05 | 0,319495291 | 0,768 | 0,523 | 0,253550572 | T_EC | Ndufs2 |
| Rps5 | 1,64E-05 | 0,280738757 | 1 | 0,946 | 0,259894041 | T_EC | Rps5 |
| S100a6 | 1,66E-05 | 0,314000382 | 0,988 | 0,768 | 0,263864275 | T_EC | S100a6 |
| Cd82 | 1,75E-05 | 0,45639662 | 0,598 | 0,357 | 0,277039428 | T_EC | Cd82 |
| Rab6a | 1,75E-05 | 0,273247476 | 0,768 | 0,519 | 0,277164796 | T_EC | Rab6a |
| Tpm3 | 1,75E-05 | 0,268065349 | 0,976 | 0,946 | 0,27808094 | T_EC | Tpm3 |
| Itm2c | 1,90E-05 | 0,381559995 | 0,939 | 0,851 | 0,301187803 | T_EC | Itm2c |
| Hnrnpk | 1,93E-05 | 0,354661262 | 1 | 0,959 | 0,305189796 | T_EC | Hnrnpk |
| Pglyrp1 | 2,03E-05 | 0,279769801 | 0,561 | 0,299 | 0,322213695 | T_EC | Pglyrp1 |
| Anxa2 | 2,08E-05 | 0,346345768 | 1 | 0,826 | 0,329758393 | T_EC | Anxa2 |
| Cyb5r3 | 2,18E-05 | 0,302959945 | 0,915 | 0,664 | 0,345019658 | T_EC | Cyb5r3 |
| She | 2,18E-05 | 0,295663731 | 0,585 | 0,315 | 0,34562321 | T_EC | She |
| Rer1 | 2,18E-05 | 0,294891687 | 0,854 | 0,656 | 0,345684761 | T_EC | Rer1 |
| Cdipt | 2,18E-05 | 0,36597326 | 0,829 | 0,577 | 0,345752526 | T_EC | Cdipt |
| Eif3m | 2,23E-05 | 0,300118193 | 0,829 | 0,585 | 0,353910452 | T_EC | Eif3m |
| Nrbp1 | 2,29E-05 | 0,387510919 | 0,756 | 0,544 | 0,363369354 | T_EC | Nrbp1 |
| Ece1 | 2,29E-05 | 0,28633203 | 1 | 0,689 | 0,363596515 | T_EC | Ece1 |
| Hes1 | 2,30E-05 | 0,3186896 | 0,768 | 0,515 | 0,364165972 | T_EC | Hes1 |
| Puf60 | 2,33E-05 | 0,323807521 | 0,768 | 0,531 | 0,369397924 | T_EC | Puf60 |
| Mtmr11 | 2,41E-05 | 0,322817228 | 0,415 | 0,191 | 0,382105717 | T_EC | Mtmr11 |
| Cct5 | 2,52E-05 | 0,3568722 | 0,866 | 0,693 | 0,398903049 | T_EC | Cct5 |
| Pex14 | 2,62E-05 | 0,253813202 | 0,415 | 0,195 | 0,415879061 | T_EC | Pex14 |
| Ppp1r2 | 2,67E-05 | 0,399797259 | 0,829 | 0,602 | 0,423751933 | T_EC | Ppp1r2 |
| Gimap6 | 2,70E-05 | 0,260830815 | 0,951 | 0,631 | 0,427292113 | T_EC | Gimap6 |
| Aplnr | 2,70E-05 | 0,361572297 | 0,78 | 0,49 | 0,428675839 | T_EC | Aplnr |
| Gnb4 | 2,70E-05 | 0,304445457 | 0,659 | 0,378 | 0,428677019 | T_EC | Gnb4 |
| Serpinb6b | 2,73E-05 | 0,329604835 | 0,28 | 0,1 | 0,433507493 | T_EC | Serpinb6b |
| Ssr2 | 2,86E-05 | 0,288179427 | 0,793 | 0,589 | 0,453015202 | T_EC | Ssr2 |
| Adm | 2,90E-05 | 0,396018488 | 0,524 | 0,261 | 0,459457511 | T_EC | Adm |
| Kitl | 2,92E-05 | 0,294326439 | 0,524 | 0,261 | 0,46278349 | T_EC | Kitl |
| Tfg | 2,96E-05 | 0,263484991 | 0,646 | 0,369 | 0,468908876 | T_EC | Tfg |
| Ptp4a3 | 3,12E-05 | 0,251318797 | 0,78 | 0,556 | 0,494760917 | T_EC | Ptp4a3 |
| Prelid1 | 3,25E-05 | 0,280176091 | 0,963 | 0,867 | 0,514527627 | T_EC | Prelid1 |
| Cav2 | 3,25E-05 | 0,337565739 | 0,768 | 0,498 | 0,515255689 | T_EC | Cav2 |
| Morf4l2 | 3,34E-05 | 0,36999902 | 0,902 | 0,78 | 0,529230546 | T_EC | Morf4l2 |
| Nmi | 3,35E-05 | 0,282326238 | 0,524 | 0,295 | 0,530472241 | T_EC | Nmi |
| Tpm4 | 3,38E-05 | 0,329377638 | 0,988 | 0,905 | 0,535920435 | T_EC | Tpm4 |
| Surf4 | 3,41E-05 | 0,394739082 | 0,854 | 0,693 | 0,540481511 | T_EC | Surf4 |
| Mtx2 | 3,64E-05 | 0,277375784 | 0,537 | 0,29 | 0,57686998 | T_EC | Mtx2 |
| Sorbs3 | 3,80E-05 | 0,284122937 | 0,902 | 0,614 | 0,602918398 | T_EC | Sorbs3 |
| Adam15 | 3,81E-05 | 0,3700465 | 0,915 | 0,78 | 0,603614624 | T_EC | Adam15 |
| Sox7 | 3,81E-05 | 0,339353008 | 0,768 | 0,51 | 0,603969486 | T_EC | Sox7 |
| Lpcat3 | 3,85E-05 | 0,305663784 | 0,427 | 0,207 | 0,61006984 | T_EC | Lpcat3 |
| F2r | 3,85E-05 | 0,501262933 | 0,537 | 0,303 | 0,610625159 | T_EC | F2r |
| Efna1 | 4,00E-05 | 0,256977581 | 0,878 | 0,568 | 0,63470916 | T_EC | Efna1 |
| Eng | 4,02E-05 | 0,289926412 | 0,976 | 0,747 | 0,636748246 | T_EC | Eng |
| Tmem204 | 4,07E-05 | 0,273164435 | 0,78 | 0,473 | 0,645706202 | T_EC | Tmem204 |
| 1110004F10Rik | 4,10E-05 | 0,331191603 | 0,817 | 0,568 | 0,649198949 | T_EC | 1110004F10Rik |
| Abcb1a | 4,13E-05 | 0,409092243 | 0,549 | 0,336 | 0,654207213 | T_EC | Abcb1a |
| Bzw1 | 4,15E-05 | 0,31837779 | 0,89 | 0,755 | 0,658398215 | T_EC | Bzw1 |
| Epas1 | 4,23E-05 | 0,333777684 | 0,878 | 0,568 | 0,670234671 | T_EC | Epas1 |
| Suclg2 | 4,48E-05 | 0,333911754 | 0,451 | 0,232 | 0,710037658 | T_EC | Suclg2 |
| Oaz2 | 4,68E-05 | 0,401967018 | 0,817 | 0,568 | 0,741640702 | T_EC | Oaz2 |
| Ndufv1 | 4,71E-05 | 0,268075251 | 0,622 | 0,378 | 0,746539857 | T_EC | Ndufv1 |
| Ubtd1 | 4,84E-05 | 0,354440828 | 0,573 | 0,353 | 0,766882032 | T_EC | Ubtd1 |
| Xbp1 | 5,03E-05 | 0,296771666 | 0,927 | 0,78 | 0,796991003 | T_EC | Xbp1 |
| Alad | 5,05E-05 | 0,294515774 | 0,659 | 0,423 | 0,800905429 | T_EC | Alad |
| Tmem214 | 5,08E-05 | 0,39158971 | 0,39 | 0,187 | 0,804573443 | T_EC | Tmem214 |
| Mgat1 | 5,11E-05 | 0,334953512 | 0,659 | 0,427 | 0,810709642 | T_EC | Mgat1 |
| 6430562O15Rik | 5,30E-05 | 0,339620545 | 0,293 | 0,108 | 0,84008813 | T_EC | 6430562O15Rik |
| Dll4 | 5,34E-05 | 0,285420407 | 0,683 | 0,402 | 0,847181089 | T_EC | Dll4 |
| Trim16 | 5,37E-05 | 0,322074482 | 0,415 | 0,191 | 0,851846866 | T_EC | Trim16 |
| Dynll1 | 5,38E-05 | 0,294400978 | 0,939 | 0,822 | 0,853373128 | T_EC | Dynll1 |
| Ddx5 | 5,85E-05 | 0,293823835 | 1 | 0,979 | 0,927663134 | T_EC | Ddx5 |
| Hmgcl | 5,92E-05 | 0,282738172 | 0,622 | 0,361 | 0,939071675 | T_EC | Hmgcl |
| Traf7 | 6,09E-05 | 0,276006318 | 0,585 | 0,34 | 0,965595747 | T_EC | Traf7 |
| Pcp4l1 | 6,18E-05 | 0,581360826 | 0,268 | 0,104 | 0,979480306 | T_EC | Pcp4l1 |
| Rab18 | 6,54E-05 | 0,286485197 | 0,768 | 0,498 | 1 | T_EC | Rab18 |
| Slc52a3 | 6,58E-05 | 0,302487391 | 0,341 | 0,145 | 1 | T_EC | Slc52a3 |
| Ext2 | 6,62E-05 | 0,281135896 | 0,634 | 0,398 | 1 | T_EC | Ext2 |
| Msn | 6,64E-05 | 0,281736628 | 1 | 0,954 | 1 | T_EC | Msn |
| Pebp1 | 7,09E-05 | 0,320878477 | 0,744 | 0,548 | 1 | T_EC | Pebp1 |
| Rpl8 | 7,11E-05 | 0,259056769 | 0,976 | 0,938 | 1 | T_EC | Rpl8 |
| Frmd8 | 7,24E-05 | 0,328009845 | 0,451 | 0,232 | 1 | T_EC | Frmd8 |
| Fkbp9 | 7,41E-05 | 0,342265849 | 0,573 | 0,336 | 1 | T_EC | Fkbp9 |
| Sh3bp5 | 7,51E-05 | 0,489482809 | 0,585 | 0,39 | 1 | T_EC | Sh3bp5 |
| Capn2 | 7,98E-05 | 0,304617173 | 0,671 | 0,423 | 1 | T_EC | Capn2 |
| Ehd3 | 8,31E-05 | 0,330045418 | 0,561 | 0,311 | 1 | T_EC | Ehd3 |
| Dnajc10 | 8,48E-05 | 0,343679385 | 0,61 | 0,378 | 1 | T_EC | Dnajc10 |
| Ctnnb1 | 8,85E-05 | 0,268110839 | 0,963 | 0,784 | 1 | T_EC | Ctnnb1 |
| Pgm1 | 9,13E-05 | 0,268208715 | 0,646 | 0,378 | 1 | T_EC | Pgm1 |

| | | | | | | | |
|---|---|---|---|---|---|---|---|
| Tbc1d14 | 5,53E-05 | 0,292106893 | 0,407 | 0,19 | 0,876494101 | 1 | Tbc1d14 |
| Calr | 5,71E-05 | 0,276697 | 0,963 | 0,872 | 0,905681229 | 1 | Calr |
| Psme1 | 5,76E-05 | 0,35124452 | 0,852 | 0,707 | 0,912454285 | 1 | Psme1 |
| Lamp2 | 5,91E-05 | 0,362412043 | 0,901 | 0,707 | 0,936613516 | 1 | Lamp2 |
| Neu1 | 6,10E-05 | 0,407221473 | 0,457 | 0,244 | 0,96739424 | 1 | Neu1 |
| Atp2b1 | 6,12E-05 | 0,253957571 | 0,84 | 0,533 | 0,97041529 | 1 | Atp2b1 |
| Cdc42se1 | 6,27E-05 | 0,374070549 | 0,741 | 0,554 | 0,993821394 | 1 | Cdc42se1 |
| Cox5a | 6,81E-05 | 0,329080342 | 0,914 | 0,752 | 1 | 1 | Cox5a |
| Naga | 6,85E-05 | 0,327309289 | 0,556 | 0,314 | 1 | 1 | Naga |
| Lamtor1 | 6,90E-05 | 0,309293295 | 0,728 | 0,504 | 1 | 1 | Lamtor1 |
| Rpsa | 7,55E-05 | 0,259320969 | 1 | 0,934 | 1 | 1 | Rpsa |
| Fam174a | 7,74E-05 | 0,335653986 | 0,531 | 0,306 | 1 | 1 | Fam174a |
| Rasa3 | 7,81E-05 | 0,281353895 | 0,383 | 0,174 | 1 | 1 | Rasa3 |
| Tln1 | 7,98E-05 | 0,297268989 | 0,926 | 0,831 | 1 | 1 | Tln1 |
| Gab2 | 9,01E-05 | 0,349502881 | 0,494 | 0,277 | 1 | 1 | Gab2 |
| Gna12 | 9,38E-05 | 0,418271329 | 0,691 | 0,471 | 1 | 1 | Gna12 |
| Tkt | 0,000104102 | 0,254276801 | 0,765 | 0,537 | 1 | 1 | Tkt |
| Pnrc1 | 0,000110131 | 0,337376004 | 0,58 | 0,372 | 1 | 1 | Pnrc1 |
| Pgam1 | 0,000121458 | 0,481171014 | 0,877 | 0,674 | 1 | 1 | Pgam1 |
| Agps | 0,000134388 | 0,252685423 | 0,469 | 0,24 | 1 | 1 | Agps |
| Slc48a1 | 0,000135331 | 0,263373231 | 0,42 | 0,207 | 1 | 1 | Slc48a1 |
| Scamp2 | 0,000140012 | 0,272735841 | 0,691 | 0,455 | 1 | 1 | Scamp2 |
| Fam234a | 0,0001411 | 0,280142013 | 0,358 | 0,161 | 1 | 1 | Fam234a |
| Ski | 0,000141441 | 0,331579784 | 0,654 | 0,45 | 1 | 1 | Ski |
| Xylt2 | 0,000143447 | 0,257927367 | 0,272 | 0,107 | 1 | 1 | Xylt2 |
| Psme2 | 0,000143635 | 0,275848433 | 0,889 | 0,764 | 1 | 1 | Psme2 |
| Ap2a2 | 0,000147513 | 0,304147864 | 0,728 | 0,554 | 1 | 1 | Ap2a2 |
| Oaz1 | 0,000154596 | 0,264464644 | 1 | 0,959 | 1 | 1 | Oaz1 |
| Herc4 | 0,000154734 | 0,359136314 | 0,358 | 0,174 | 1 | 1 | Herc4 |
| Vdac2 | 0,000160588 | 0,303634252 | 0,877 | 0,649 | 1 | 1 | Vdac2 |
| Frmd4b | 0,00016283 | 0,475232633 | 0,531 | 0,331 | 1 | 1 | Frmd4b |
| Ahsa1 | 0,000173995 | 0,355986811 | 0,543 | 0,331 | 1 | 1 | Ahsa1 |
| Hpf1 | 0,000174231 | 0,275094629 | 0,481 | 0,277 | 1 | 1 | Hpf1 |
| Fkbp15 | 0,000187463 | 0,283872936 | 0,531 | 0,31 | 1 | 1 | Fkbp15 |
| Pmvk | 0,000200092 | 0,287175187 | 0,432 | 0,219 | 1 | 1 | Pmvk |
| Narfl | 0,000215593 | 0,272199991 | 0,259 | 0,099 | 1 | 1 | Narfl |
| Rpl36al | 0,000217303 | 0,291709092 | 0,938 | 0,798 | 1 | 1 | Rpl36al |
| Capza2 | 0,000217745 | 0,255354865 | 0,951 | 0,818 | 1 | 1 | Capza2 |
| Tob2 | 0,000231997 | 0,271365236 | 0,37 | 0,194 | 1 | 1 | Tob2 |
| Mdh1 | 0,000238528 | 0,282455577 | 0,654 | 0,434 | 1 | 1 | Mdh1 |
| Ubc | 0,000245708 | 0,302117554 | 1 | 0,971 | 1 | 1 | Ubc |
| Ccdc86 | 0,000248756 | 0,299585161 | 0,333 | 0,149 | 1 | 1 | Ccdc86 |
| mt-Nd5 | 0,000254438 | 0,258544722 | 0,988 | 0,979 | 1 | 1 | mt-Nd5 |
| Sptssa | 0,000280441 | 0,314468222 | 0,691 | 0,521 | 1 | 1 | Sptssa |
| Sec11c | 0,000281978 | 0,343426263 | 0,679 | 0,496 | 1 | 1 | Sec11c |
| Colgalt1 | 0,000285128 | 0,444913719 | 0,605 | 0,442 | 1 | 1 | Colgalt1 |
| Atp6v0e | 0,000294512 | 0,277202448 | 0,864 | 0,665 | 1 | 1 | Atp6v0e |
| Jun | 0,000296263 | 0,439974447 | 0,654 | 0,459 | 1 | 1 | Jun |
| Syngr2 | 0,000328371 | 0,358140639 | 0,815 | 0,599 | 1 | 1 | Syngr2 |
| Incenp | 0,000388183 | 0,312358927 | 0,284 | 0,128 | 1 | 1 | Incenp |
| Tpt1 | 0,000393596 | 0,256080445 | 1 | 0,983 | 1 | 1 | Tpt1 |
| Gstp1 | 0,000410133 | 0,283807883 | 0,63 | 0,475 | 1 | 1 | Gstp1 |
| Ivns1abp | 0,000421008 | 0,339652005 | 0,827 | 0,616 | 1 | 1 | Ivns1abp |
| Lrpap1 | 0,000434266 | 0,294307354 | 0,605 | 0,384 | 1 | 1 | Lrpap1 |
| Map3k1 | 0,000443908 | 0,327620612 | 0,617 | 0,455 | 1 | 1 | Map3k1 |
| Man1c1 | 0,000459086 | 0,28352606 | 0,432 | 0,252 | 1 | 1 | Man1c1 |
| Cept1 | 0,000579451 | 0,257571054 | 0,42 | 0,231 | 1 | 1 | Cept1 |
| Plbd2 | 0,000580448 | 0,292592948 | 0,519 | 0,314 | 1 | 1 | Plbd2 |
| Cltc | 0,000586272 | 0,326264032 | 0,938 | 0,851 | 1 | 1 | Cltc |
| mt-Nd2 | 0,000606054 | 0,27418639 | 0,975 | 0,979 | 1 | 1 | mt-Nd2 |
| Tnfaip3 | 0,000608246 | 0,51500519 | 0,259 | 0,116 | 1 | 1 | Tnfaip3 |
| Rpl10 | 0,000622519 | 0,272833737 | 0,988 | 0,93 | 1 | 1 | Rpl10 |
| 2310035C23Rik | 0,000688 | 0,275192349 | 0,469 | 0,289 | 1 | 1 | 2310035C23Rik |
| Tm6sf1 | 0,000732851 | 0,319738078 | 0,432 | 0,236 | 1 | 1 | Tm6sf1 |
| Mrps26 | 0,000850556 | 0,253855037 | 0,481 | 0,293 | 1 | 1 | Mrps26 |
| Tsc22d3 | 0,001002564 | 0,269443601 | 0,741 | 0,55 | 1 | 1 | Tsc22d3 |
| Rfc2 | 0,001011926 | 0,295254248 | 0,556 | 0,368 | 1 | 1 | Rfc2 |
| Dram2 | 0,001028189 | 0,279026799 | 0,716 | 0,558 | 1 | 1 | Dram2 |
| Tpi1 | 0,001106805 | 0,291732728 | 0,926 | 0,711 | 1 | 1 | Tpi1 |
| Echs1 | 0,001148989 | 0,355575642 | 0,481 | 0,322 | 1 | 1 | Echs1 |
| Arhgap25 | 0,001207998 | 0,317985339 | 0,568 | 0,368 | 1 | 1 | Arhgap25 |
| Zfyve19 | 0,001300237 | 0,327754084 | 0,272 | 0,132 | 1 | 1 | Zfyve19 |
| Ensa | 0,00130699 | 0,3004799 | 0,605 | 0,409 | 1 | 1 | Ensa |
| Cdca3 | 0,001342141 | 0,41526518 | 0,259 | 0,12 | 1 | 1 | Cdca3 |
| Mif4gd | 0,001384247 | 0,258736045 | 0,395 | 0,219 | 1 | 1 | Mif4gd |
| Gpr137b-ps | 0,001428522 | 0,265826649 | 0,432 | 0,26 | 1 | 1 | Gpr137b-ps |
| Dnm2 | 0,001570228 | 0,256464864 | 0,741 | 0,579 | 1 | 1 | Dnm2 |
| Marcks | 0,001595358 | 0,270161179 | 1 | 0,959 | 1 | 1 | Marcks |
| Atp6v1c1 | 0,00165266 | 0,302802144 | 0,506 | 0,331 | 1 | 1 | Atp6v1c1 |
| Cerk | 0,001671041 | 0,255840249 | 0,543 | 0,347 | 1 | 1 | Cerk |
| Nubp1 | 0,001682947 | 0,352596296 | 0,444 | 0,289 | 1 | 1 | Nubp1 |
| Tuba1c | 0,001700921 | 0,299674033 | 0,79 | 0,603 | 1 | 1 | Tuba1c |
| Rrbp1 | 0,001727621 | 0,288315791 | 0,938 | 0,835 | 1 | 1 | Rrbp1 |
| Brk1 | 0,001770722 | 0,287077428 | 0,815 | 0,711 | 1 | 1 | Brk1 |
| Ifnar2 | 0,001816522 | 0,305014514 | 0,667 | 0,517 | 1 | 1 | Ifnar2 |
| Sirt2 | 0,001851002 | 0,299959535 | 0,531 | 0,355 | 1 | 1 | Sirt2 |
| Cisd2 | 0,001872841 | 0,258851665 | 0,506 | 0,326 | 1 | 1 | Cisd2 |
| Washc2 | 0,001874363 | 0,325287528 | 0,407 | 0,244 | 1 | 1 | Washc2 |
| Ndufb10 | 0,001968151 | 0,274319719 | 0,765 | 0,719 | 1 | 1 | Ndufb10 |
| Camta2 | 0,002125487 | 0,325067527 | 0,407 | 0,256 | 1 | 1 | Camta2 |
| Htra2 | 0,002204692 | 0,318091171 | 0,309 | 0,165 | 1 | 1 | Htra2 |
| Slc25a5 | 0,002585475 | 0,291702116 | 0,938 | 0,888 | 1 | 1 | Slc25a5 |
| Commd9 | 0,002827814 | 0,276233384 | 0,333 | 0,186 | 1 | 1 | Commd9 |
| Ndufs7 | 0,003125321 | 0,258165094 | 0,691 | 0,521 | 1 | 1 | Ndufs7 |
| Rab11fip5 | 0,003313717 | 0,259085413 | 0,457 | 0,302 | 1 | 1 | Rab11fip5 |
| Bnip3 | 0,003384699 | 0,287352749 | 0,346 | 0,174 | 1 | 1 | Bnip3 |
| Cyp4f16 | 0,003393531 | 0,274379499 | 0,259 | 0,132 | 1 | 1 | Cyp4f16 |
| Herpud1 | 0,00362017 | 0,32477929 | 0,568 | 0,393 | 1 | 1 | Herpud1 |
| Phip | 0,00378353 | 0,274333164 | 0,37 | 0,231 | 1 | 1 | Phip |
| Ier5 | 0,004077352 | 0,302594807 | 0,444 | 0,298 | 1 | 1 | Ier5 |
| Zfp810 | 0,004144817 | 0,273331743 | 0,259 | 0,128 | 1 | 1 | Zfp810 |
| Rpl22l1 | 0,004217647 | 0,283652359 | 0,864 | 0,76 | 1 | 1 | Rpl22l1 |
| Preb | 0,004552737 | 0,323296902 | 0,444 | 0,31 | 1 | 1 | Preb |
| Oas1a | 0,004839028 | 0,278052257 | 0,309 | 0,174 | 1 | 1 | Oas1a |
| Rabggta | 0,004937431 | 0,371226076 | 0,284 | 0,157 | 1 | 1 | Rabggta |
| Nr4a1 | 0,004988915 | 0,608428637 | 0,494 | 0,351 | 1 | 1 | Nr4a1 |
| Lamtor3 | 0,0053705 | 0,32271594 | 0,519 | 0,413 | 1 | 1 | Lamtor3 |
| Leng8 | 0,00618549 | 0,277088141 | 0,58 | 0,417 | 1 | 1 | Leng8 |
| Ifnar1 | 0,006287453 | 0,313146503 | 0,63 | 0,467 | 1 | 1 | Ifnar1 |
| Dnajc13 | 0,006490648 | 0,263354753 | 0,346 | 0,215 | 1 | 1 | Dnajc13 |
| Itpripl1 | 0,007411859 | 0,25218796 | 0,407 | 0,256 | 1 | 1 | Itpripl1 |
| Rbck1 | 0,008204591 | 0,273263975 | 0,481 | 0,343 | 1 | 1 | Rbck1 |
| Hat1 | 0,008668322 | 0,321692578 | 0,358 | 0,223 | 1 | 1 | Hat1 |
| Atp6v0d1 | 0,009870302 | 0,266712217 | 0,654 | 0,574 | 1 | 1 | Atp6v0d1 |
| Tnfrsf9 | 2,15E-20 | 1,277038647 | 0,683 | 0,154 | 3,41E-16 | 2 | Tnfrsf9 |
| Rhoc | 1,41E-19 | 0,957044276 | 0,984 | 0,669 | 2,23E-15 | 2 | Rhoc |
| Prnd | 8,13E-18 | 1,296642228 | 0,635 | 0,142 | 1,29E-13 | 2 | Prnd |
| Vwa1 | 8,08E-16 | 0,887936298 | 0,984 | 0,642 | 1,28E-11 | 2 | Vwa1 |
| Prnp | 2,40E-15 | 0,845804343 | 0,984 | 0,565 | 3,81E-11 | 2 | Prnp |
| Tnfaip8l1 | 1,08E-14 | 0,726570586 | 0,746 | 0,25 | 1,71E-10 | 2 | Tnfaip8l1 |

| | | | | | | | |
|---|---|---|---|---|---|---|---|
| Wdr75 | 9,14E-05 | 0,364094618 | 0,317 | 0,137 | 1 | T_EC | Wdr75 |
| Ctnnbip1 | 9,65E-05 | 0,28745625 | 0,707 | 0,494 | 1 | T_EC | Ctnnbip1 |
| Gdpd5 | 9,90E-05 | 0,384517449 | 0,329 | 0,149 | 1 | T_EC | Gdpd5 |
| Tmem184b | 0,000106455 | 0,27783176 | 0,732 | 0,494 | 1 | T_EC | Tmem184b |
| Crip2 | 0,000108011 | 0,266330263 | 0,976 | 0,643 | 1 | T_EC | Crip2 |
| Txnrd1 | 0,000109612 | 0,28595238 | 0,561 | 0,332 | 1 | T_EC | Txnrd1 |
| Trim27 | 0,000118074 | 0,294048058 | 0,5 | 0,278 | 1 | T_EC | Trim27 |
| Psma5 | 0,000118142 | 0,286883142 | 0,878 | 0,689 | 1 | T_EC | Psma5 |
| Stxbp1 | 0,000120522 | 0,335510268 | 0,585 | 0,349 | 1 | T_EC | Stxbp1 |
| Litaf | 0,000120976 | 0,293293686 | 0,988 | 0,867 | 1 | T_EC | Litaf |
| Ubqln1 | 0,000124769 | 0,333968118 | 0,585 | 0,373 | 1 | T_EC | Ubqln1 |
| Cd109 | 0,000127439 | 0,283954725 | 0,622 | 0,373 | 1 | T_EC | Cd109 |
| Stt3b | 0,000128806 | 0,261561712 | 0,72 | 0,473 | 1 | T_EC | Stt3b |
| Polr2e | 0,000128939 | 0,285420311 | 0,646 | 0,436 | 1 | T_EC | Polr2e |
| Elavl1 | 0,000130511 | 0,299205128 | 0,598 | 0,394 | 1 | T_EC | Elavl1 |
| 1810011O10Rik | 0,000133277 | 0,265946557 | 0,476 | 0,245 | 1 | T_EC | 1810011O10Rik |
| Pja1 | 0,000136348 | 0,329175304 | 0,317 | 0,137 | 1 | T_EC | Pja1 |
| Plpp3 | 0,000136515 | 0,332923696 | 0,732 | 0,485 | 1 | T_EC | Plpp3 |
| Nck1 | 0,000138415 | 0,282859286 | 0,561 | 0,311 | 1 | T_EC | Nck1 |
| Fzd4 | 0,000146206 | 0,36299121 | 0,671 | 0,44 | 1 | T_EC | Fzd4 |
| Rflnb | 0,000147564 | 0,33612519 | 0,476 | 0,261 | 1 | T_EC | Rflnb |
| Ppa1 | 0,000153707 | 0,275690519 | 0,366 | 0,17 | 1 | T_EC | Ppa1 |
| Lrg1 | 0,00015389 | 0,257885189 | 0,927 | 0,647 | 1 | T_EC | Lrg1 |
| Rwdd4a | 0,000154522 | 0,274540932 | 0,341 | 0,162 | 1 | T_EC | Rwdd4a |
| Sparcl1 | 0,000154762 | 0,313107526 | 0,963 | 0,643 | 1 | T_EC | Sparcl1 |
| Cdc42ep3 | 0,000158675 | 0,301959481 | 0,646 | 0,39 | 1 | T_EC | Cdc42ep3 |
| Aprt | 0,000159936 | 0,251426567 | 0,817 | 0,635 | 1 | T_EC | Aprt |
| Rbm18 | 0,000161618 | 0,302331426 | 0,378 | 0,178 | 1 | T_EC | Rbm18 |
| Dbn1 | 0,000172428 | 0,252854594 | 0,72 | 0,444 | 1 | T_EC | Dbn1 |
| Agpat4 | 0,000173199 | 0,261453063 | 0,378 | 0,187 | 1 | T_EC | Agpat4 |
| Ubxn6 | 0,000179976 | 0,251300328 | 0,451 | 0,232 | 1 | T_EC | Ubxn6 |
| Snai1 | 0,000181607 | 0,280060107 | 0,415 | 0,22 | 1 | T_EC | Snai1 |
| Metap2 | 0,000187279 | 0,352695527 | 0,829 | 0,656 | 1 | T_EC | Metap2 |
| Tmco1 | 0,000188264 | 0,301987541 | 0,756 | 0,552 | 1 | T_EC | Tmco1 |
| Itfg1 | 0,000190034 | 0,376308649 | 0,561 | 0,34 | 1 | T_EC | Itfg1 |
| Nsmce1 | 0,00019551 | 0,254058078 | 0,622 | 0,398 | 1 | T_EC | Nsmce1 |
| Hspa5 | 0,000211376 | 0,41896865 | 0,951 | 0,867 | 1 | T_EC | Hspa5 |
| Pfkp | 0,00021559 | 0,362174141 | 0,61 | 0,407 | 1 | T_EC | Pfkp |
| Nxpe4 | 0,000219882 | 0,276959055 | 0,463 | 0,245 | 1 | T_EC | Nxpe4 |
| Mapre1 | 0,000226821 | 0,271457116 | 0,829 | 0,71 | 1 | T_EC | Mapre1 |
| Abcc4 | 0,000227011 | 0,308988813 | 0,61 | 0,369 | 1 | T_EC | Abcc4 |
| Ran | 0,000230078 | 0,291680195 | 0,915 | 0,78 | 1 | T_EC | Ran |
| Srsf6 | 0,000238514 | 0,278581779 | 0,744 | 0,519 | 1 | T_EC | Srsf6 |
| Gpi1 | 0,000242156 | 0,271360275 | 0,951 | 0,801 | 1 | T_EC | Gpi1 |
| Itga3 | 0,000244882 | 0,26163826 | 0,537 | 0,328 | 1 | T_EC | Itga3 |
| Upf3a | 0,000248734 | 0,251975961 | 0,427 | 0,216 | 1 | T_EC | Upf3a |
| Gnas | 0,000250287 | 0,262889823 | 0,951 | 0,88 | 1 | T_EC | Gnas |
| Yipf1 | 0,000261579 | 0,30254566 | 0,451 | 0,257 | 1 | T_EC | Yipf1 |
| Rfk | 0,000262451 | 0,332706681 | 0,622 | 0,461 | 1 | T_EC | Rfk |
| Ifi44 | 0,000265546 | 0,278489864 | 0,317 | 0,133 | 1 | T_EC | Ifi44 |
| H13 | 0,00026574 | 0,257080832 | 0,854 | 0,651 | 1 | T_EC | H13 |
| Sec62 | 0,00027191 | 0,266170569 | 0,915 | 0,768 | 1 | T_EC | Sec62 |
| Cdc16 | 0,000271916 | 0,29327058 | 0,488 | 0,278 | 1 | T_EC | Cdc16 |
| Anapc4 | 0,000275335 | 0,321270525 | 0,488 | 0,299 | 1 | T_EC | Anapc4 |
| Crem | 0,000277191 | 0,330736499 | 0,524 | 0,315 | 1 | T_EC | Crem |
| Asns | 0,000318266 | 0,316292573 | 0,28 | 0,116 | 1 | T_EC | Asns |
| Plp2 | 0,000320133 | 0,36607576 | 0,793 | 0,568 | 1 | T_EC | Plp2 |
| Pttg1ip | 0,00034162 | 0,259971161 | 0,817 | 0,589 | 1 | T_EC | Pttg1ip |
| Skp1a | 0,000347643 | 0,257470084 | 0,915 | 0,797 | 1 | T_EC | Skp1a |
| Pkm | 0,000355565 | 0,280888298 | 1 | 0,975 | 1 | T_EC | Pkm |
| Myzap | 0,000374347 | 0,281879046 | 0,549 | 0,34 | 1 | T_EC | Myzap |
| Sec61a1 | 0,000383226 | 0,279587783 | 0,902 | 0,726 | 1 | T_EC | Sec61a1 |
| Prpsap1 | 0,000383298 | 0,326186094 | 0,549 | 0,34 | 1 | T_EC | Prpsap1 |
| Ccny | 0,000384656 | 0,344966544 | 0,488 | 0,303 | 1 | T_EC | Ccny |
| Tubb4b | 0,000390117 | 0,392969098 | 0,695 | 0,523 | 1 | T_EC | Tubb4b |
| Snrpb | 0,00039313 | 0,299802129 | 0,915 | 0,776 | 1 | T_EC | Snrpb |
| Csnk1d | 0,000402223 | 0,387275212 | 0,622 | 0,44 | 1 | T_EC | Csnk1d |
| Rcn1 | 0,000412995 | 0,26468262 | 0,402 | 0,22 | 1 | T_EC | Rcn1 |
| Scarb1 | 0,000435139 | 0,280044498 | 0,659 | 0,456 | 1 | T_EC | Scarb1 |
| Amotl2 | 0,000477685 | 0,281904587 | 0,268 | 0,112 | 1 | T_EC | Amotl2 |
| Gja1 | 0,000482923 | 0,327842672 | 0,829 | 0,602 | 1 | T_EC | Gja1 |
| Prkar1a | 0,000504374 | 0,283985033 | 0,963 | 0,876 | 1 | T_EC | Prkar1a |
| Gorasp2 | 0,000509206 | 0,307097865 | 0,78 | 0,61 | 1 | T_EC | Gorasp2 |
| Tanc1 | 0,000520361 | 0,255253764 | 0,427 | 0,232 | 1 | T_EC | Tanc1 |
| Mbtps1 | 0,000536695 | 0,322892439 | 0,415 | 0,232 | 1 | T_EC | Mbtps1 |
| Arhgap27 | 0,000537935 | 0,272723519 | 0,549 | 0,332 | 1 | T_EC | Arhgap27 |
| Rnf19a | 0,000544444 | 0,272169822 | 0,415 | 0,228 | 1 | T_EC | Rnf19a |
| Cx3cl1 | 0,000551739 | 0,291572525 | 0,329 | 0,166 | 1 | T_EC | Cx3cl1 |
| Smad1 | 0,000566753 | 0,37827003 | 0,72 | 0,552 | 1 | T_EC | Smad1 |
| Fbxo8 | 0,00060431 | 0,324480226 | 0,329 | 0,17 | 1 | T_EC | Fbxo8 |
| Tmed4 | 0,00060565 | 0,253820347 | 0,598 | 0,365 | 1 | T_EC | Tmed4 |
| Marcksl1 | 0,0006243 | 0,25436132 | 0,902 | 0,772 | 1 | T_EC | Marcksl1 |
| Sft2d2 | 0,00062513 | 0,325920731 | 0,5 | 0,324 | 1 | T_EC | Sft2d2 |
| Cers5 | 0,000640119 | 0,327542311 | 0,537 | 0,361 | 1 | T_EC | Cers5 |
| Gpr4 | 0,00064122 | 0,315017128 | 0,768 | 0,552 | 1 | T_EC | Gpr4 |
| Lmna | 0,000646946 | 0,262804051 | 0,878 | 0,672 | 1 | T_EC | Lmna |
| Abhd4 | 0,000687063 | 0,290878932 | 0,549 | 0,361 | 1 | T_EC | Abhd4 |
| Sdha | 0,000709541 | 0,26106231 | 0,659 | 0,49 | 1 | T_EC | Sdha |
| Rbbp7 | 0,000738909 | 0,252146651 | 0,756 | 0,531 | 1 | T_EC | Rbbp7 |
| Cnn2 | 0,00075129 | 0,268671119 | 0,768 | 0,589 | 1 | T_EC | Cnn2 |
| Dusp6 | 0,00077969 | 0,279404078 | 0,951 | 0,822 | 1 | T_EC | Dusp6 |
| Lmo2 | 0,000800987 | 0,302769133 | 0,512 | 0,336 | 1 | T_EC | Lmo2 |
| Mprip | 0,000819301 | 0,271484987 | 0,78 | 0,568 | 1 | T_EC | Mprip |
| Timp3 | 0,000861046 | 0,34738411 | 0,878 | 0,573 | 1 | T_EC | Timp3 |
| Tubb2a | 0,000876079 | 0,306949942 | 0,61 | 0,427 | 1 | T_EC | Tubb2a |
| Tns2 | 0,000916991 | 0,258150195 | 0,537 | 0,328 | 1 | T_EC | Tns2 |
| Kit | 0,000923471 | 0,2683537 | 0,561 | 0,336 | 1 | T_EC | Kit |
| Serbp1 | 0,000925125 | 0,254165999 | 0,902 | 0,817 | 1 | T_EC | Serbp1 |
| Cltb | 0,000936438 | 0,256532091 | 0,732 | 0,494 | 1 | T_EC | Cltb |
| Galnt18 | 0,001044895 | 0,438748906 | 0,366 | 0,207 | 1 | T_EC | Galnt18 |
| Tcaf1 | 0,001074184 | 0,269095187 | 0,305 | 0,145 | 1 | T_EC | Tcaf1 |
| Gpcpd1 | 0,001083271 | 0,33520236 | 0,512 | 0,315 | 1 | T_EC | Gpcpd1 |
| Ncl | 0,001088085 | 0,368584325 | 0,902 | 0,822 | 1 | T_EC | Ncl |
| Sh3bp4 | 0,001111836 | 0,382226531 | 0,39 | 0,224 | 1 | T_EC | Sh3bp4 |
| Stk25 | 0,001190218 | 0,321567653 | 0,72 | 0,568 | 1 | T_EC | Stk25 |
| Eif1ax | 0,001200785 | 0,290249951 | 0,61 | 0,427 | 1 | T_EC | Eif1ax |
| Pfkl | 0,001225118 | 0,401022308 | 0,634 | 0,456 | 1 | T_EC | Pfkl |
| Fkbp8 | 0,001243782 | 0,288484486 | 0,732 | 0,544 | 1 | T_EC | Fkbp8 |
| Cct7 | 0,001247137 | 0,327248803 | 0,78 | 0,614 | 1 | T_EC | Cct7 |
| Apold1 | 0,001302701 | 0,292502658 | 0,732 | 0,498 | 1 | T_EC | Apold1 |
| Nifk | 0,001303013 | 0,278329846 | 0,451 | 0,286 | 1 | T_EC | Nifk |
| Cct8 | 0,001335425 | 0,281821189 | 0,793 | 0,71 | 1 | T_EC | Cct8 |
| Irgm1 | 0,001378798 | 0,262057005 | 0,524 | 0,324 | 1 | T_EC | Irgm1 |
| Prnd | 0,001391351 | 0,400770037 | 0,366 | 0,195 | 1 | T_EC | Prnd |
| Lrrc41 | 0,00146201 | 0,262249409 | 0,329 | 0,174 | 1 | T_EC | Lrrc41 |
| Armc1 | 0,001725458 | 0,295229791 | 0,28 | 0,141 | 1 | T_EC | Armc1 |
| Txnl1 | 0,001738172 | 0,285222951 | 0,573 | 0,427 | 1 | T_EC | Txnl1 |
| Pltp | 0,001772555 | 0,376540638 | 0,5 | 0,315 | 1 | T_EC | Pltp |
| Gde1 | 0,001798469 | 0,26293907 | 0,561 | 0,39 | 1 | T_EC | Gde1 |
| Cdk4 | 0,001935729 | 0,256885205 | 0,793 | 0,631 | 1 | T_EC | Cdk4 |
| Myadm | 0,002015117 | 0,334060649 | 0,817 | 0,751 | 1 | T_EC | Myadm |

| | | | | | | | |
|---|---|---|---|---|---|---|---|
| Trp53i11 | 2,17E-13 | 0,866035928 | 0,968 | 0,596 | 3,43E-09 | 2 | Trp53i11 |
| Mcam | 2,28E-13 | 0,760743123 | 1 | 0,665 | 3,62E-09 | 2 | Mcam |
| Fscn1 | 2,60E-13 | 0,612866322 | 1 | 0,642 | 4,12E-09 | 2 | Fscn1 |
| Pcdhgc3 | 2,43E-12 | 0,462330778 | 0,46 | 0,096 | 3,86E-08 | 2 | Pcdhgc3 |
| Actg1 | 2,63E-12 | 0,535528158 | 1 | 0,992 | 4,18E-08 | 2 | Actg1 |
| Cd82 | 2,90E-12 | 0,789469018 | 0,746 | 0,338 | 4,60E-08 | 2 | Cd82 |
| Fkbp1a | 3,20E-12 | 0,611331738 | 1 | 0,912 | 5,07E-08 | 2 | Fkbp1a |
| Csrp1 | 3,47E-12 | 0,638176967 | 0,873 | 0,492 | 5,49E-08 | 2 | Csrp1 |
| Slc2a1 | 6,73E-12 | 0,657894423 | 0,968 | 0,596 | 1,07E-07 | 2 | Slc2a1 |
| Sh2d5 | 1,00E-11 | 0,3571215 | 0,492 | 0,112 | 1,59E-07 | 2 | Sh2d5 |
| Prkd2 | 1,08E-11 | 0,541415727 | 0,778 | 0,358 | 1,71E-07 | 2 | Prkd2 |
| Rhoj | 1,08E-11 | 0,503120506 | 0,921 | 0,465 | 1,72E-07 | 2 | Rhoj |
| Meox1 | 1,48E-11 | 0,594067289 | 0,825 | 0,377 | 2,35E-07 | 2 | Meox1 |
| Calu | 1,72E-11 | 0,623906294 | 0,984 | 0,7 | 2,73E-07 | 2 | Calu |
| Myadm | 1,77E-11 | 0,762800997 | 0,937 | 0,727 | 2,80E-07 | 2 | Myadm |
| Pdlim11 | 1,88E-11 | 0,656638195 | 0,984 | 0,638 | 2,98E-07 | 2 | Pdlim1 |
| Adm | 2,00E-11 | 0,745137838 | 0,683 | 0,242 | 3,18E-07 | 2 | Adm |
| Kit | 2,11E-11 | 0,55788509 | 0,778 | 0,3 | 3,34E-07 | 2 | Kit |
| Ndfip1 | 2,83E-11 | 0,613567865 | 0,968 | 0,704 | 4,48E-07 | 2 | Ndfip1 |
| Itgb1 | 3,50E-11 | 0,624454568 | 1 | 0,927 | 5,55E-07 | 2 | Itgb1 |
| Nid2 | 3,62E-11 | 0,752179391 | 0,952 | 0,546 | 5,74E-07 | 2 | Nid2 |
| Cldn51 | 3,79E-11 | 0,754219163 | 1 | 0,642 | 6,00E-07 | 2 | Cldn5 |
| 6430562O15Rik | 4,27E-11 | 0,284861862 | 0,429 | 0,088 | 6,76E-07 | 2 | 6430562O15Rik |
| Pde4b | 1,64E-10 | 0,623284663 | 0,921 | 0,538 | 2,60E-06 | 2 | Pde4b |
| Clic6 | 1,74E-10 | 0,356201711 | 0,302 | 0,042 | 2,75E-06 | 2 | Clic6 |
| Tpm1 | 2,13E-10 | 0,58623003 | 0,921 | 0,608 | 3,38E-06 | 2 | Tpm1 |
| Smtn | 2,37E-10 | 0,568237005 | 0,889 | 0,512 | 3,75E-06 | 2 | Smtn |
| Stx6 | 3,08E-10 | 0,749753274 | 0,762 | 0,446 | 4,88E-06 | 2 | Stx6 |
| Cd276 | 3,18E-10 | 0,474707169 | 0,698 | 0,273 | 5,04E-06 | 2 | Cd276 |
| Adgrl4 | 3,46E-10 | 0,655871967 | 1 | 0,638 | 5,48E-06 | 2 | Adgrl4 |
| Grap | 3,97E-10 | 0,585246261 | 0,81 | 0,396 | 6,29E-06 | 2 | Grap |
| Fermt2 | 7,72E-10 | 0,585619576 | 0,889 | 0,535 | 1,22E-05 | 2 | Fermt2 |
| Tm4sf11 | 8,00E-10 | 0,597822042 | 0,984 | 0,665 | 1,27E-05 | 2 | Tm4sf1 |
| Vim | 8,31E-10 | 0,537923402 | 1 | 0,973 | 1,32E-05 | 2 | Vim |
| Bcl6b | 8,36E-10 | 0,575216825 | 0,667 | 0,273 | 1,33E-05 | 2 | Bcl6b |
| Bok | 1,09E-09 | 0,512182608 | 0,714 | 0,319 | 1,72E-05 | 2 | Bok |
| Tspan2 | 1,14E-09 | 0,437934267 | 0,381 | 0,088 | 1,81E-05 | 2 | Tspan2 |
| Sox7 | 1,22E-09 | 0,508263607 | 0,905 | 0,496 | 1,93E-05 | 2 | Sox7 |
| Nectin2 | 1,36E-09 | 0,608272068 | 0,746 | 0,365 | 2,15E-05 | 2 | Nectin2 |
| Ppm1j | 2,05E-09 | 0,340177538 | 0,429 | 0,115 | 3,25E-05 | 2 | Ppm1j |
| Cmtm3 | 2,07E-09 | 0,503232027 | 0,968 | 0,681 | 3,28E-05 | 2 | Cmtm3 |
| Itm2c | 2,43E-09 | 0,499409163 | 0,984 | 0,846 | 3,85E-05 | 2 | Itm2c |
| Tubb6 | 2,48E-09 | 0,621619429 | 0,968 | 0,685 | 3,93E-05 | 2 | Tubb6 |
| Odc1 | 2,76E-09 | 0,630428217 | 0,778 | 0,412 | 4,37E-05 | 2 | Odc1 |
| Rnf125 | 3,13E-09 | 0,431587706 | 0,476 | 0,142 | 4,96E-05 | 2 | Rnf125 |
| S100a16 | 3,20E-09 | 0,478709978 | 0,968 | 0,608 | 5,07E-05 | 2 | S100a16 |
| Upp11 | 4,21E-09 | 0,482057726 | 0,984 | 0,542 | 6,67E-05 | 2 | Upp1 |
| Clic4 | 5,36E-09 | 0,553916591 | 0,968 | 0,823 | 8,49E-05 | 2 | Clic4 |
| Tbrg1 | 6,59E-09 | 0,473956787 | 0,825 | 0,442 | 0,000104521 | 2 | Tbrg1 |
| F2r | 7,66E-09 | 0,722355203 | 0,651 | 0,292 | 0,000121432 | 2 | F2r |
| Ankrd37 | 7,74E-09 | 0,577780219 | 0,635 | 0,281 | 0,00012263 | 2 | Ankrd37 |
| Sparc | 7,75E-09 | 0,475758419 | 1 | 0,742 | 0,000122924 | 2 | Sparc |
| Sox181 | 8,61E-09 | 0,558470715 | 0,937 | 0,588 | 0,00013643 | 2 | Sox18 |
| Ecscr1 | 8,68E-09 | 0,435594463 | 1 | 0,665 | 0,000137533 | 2 | Ecscr |
| Tmem2521 | 1,31E-08 | 0,40183287 | 0,984 | 0,538 | 0,000208231 | 2 | Tmem252 |
| Plk2 | 1,35E-08 | 0,51934366 | 0,984 | 0,681 | 0,000213494 | 2 | Plk2 |
| Myl12a | 1,42E-08 | 0,408567202 | 1 | 0,958 | 0,000224631 | 2 | Myl12a |
| Cnn31 | 1,48E-08 | 0,495210294 | 1 | 0,673 | 0,000233873 | 2 | Cnn3 |
| Impdh1 | 1,64E-08 | 0,512920417 | 0,778 | 0,392 | 0,000260743 | 2 | Impdh1 |
| Prkcdbp | 1,67E-08 | 0,521429039 | 0,984 | 0,662 | 0,000264408 | 2 | Prkcdbp |
| Mast4 | 2,33E-08 | 0,312528019 | 0,841 | 0,469 | 0,000368811 | 2 | Mast4 |
| Tagln2 | 2,53E-08 | 0,497750681 | 0,968 | 0,838 | 0,000400387 | 2 | Tagln2 |
| Sh2d3c | 2,66E-08 | 0,511727819 | 0,889 | 0,546 | 0,000421239 | 2 | Sh2d3c |
| Mpzl1 | 2,81E-08 | 0,522357627 | 0,889 | 0,515 | 0,000445758 | 2 | Mpzl1 |
| Nox4 | 2,91E-08 | 0,388116588 | 0,429 | 0,123 | 0,000460862 | 2 | Nox4 |
| Esam1 | 4,12E-08 | 0,50210432 | 1 | 0,669 | 0,000653099 | 2 | Esam |
| Pxdn | 4,17E-08 | 0,482932196 | 0,825 | 0,462 | 0,000660247 | 2 | Pxdn |
| Pkm | 4,82E-08 | 0,479019032 | 1 | 0,977 | 0,000764626 | 2 | Pkm |
| Slc38a3 | 4,83E-08 | 0,274386603 | 0,286 | 0,054 | 0,000765274 | 2 | Slc38a3 |
| Serpinh1 | 5,26E-08 | 0,335014224 | 0,984 | 0,654 | 0,000834343 | 2 | Serpinh1 |
| Marcksl1 | 5,46E-08 | 0,433352142 | 0,984 | 0,762 | 0,000864866 | 2 | Marcksl1 |
| Akr1b8 | 5,69E-08 | 0,33367551 | 0,683 | 0,308 | 0,000901668 | 2 | Akr1b8 |
| Mmp14 | 6,70E-08 | 0,545920466 | 0,984 | 0,835 | 0,001061747 | 2 | Mmp14 |
| Arf4 | 7,27E-08 | 0,526581046 | 0,937 | 0,762 | 0,001151879 | 2 | Arf4 |
| Creb3l2 | 7,31E-08 | 0,406748848 | 0,873 | 0,496 | 0,001158524 | 2 | Creb3l2 |
| Bhlhe40 | 7,33E-08 | 0,4979892 | 0,873 | 0,558 | 0,001162199 | 2 | Bhlhe40 |
| Tuba1a | 7,35E-08 | 0,645328015 | 0,937 | 0,669 | 0,001165239 | 2 | Tuba1a |
| Hspb81 | 7,38E-08 | 0,660217284 | 0,73 | 0,4 | 0,001170346 | 2 | Hspb8 |
| Dbn1 | 7,63E-08 | 0,399256333 | 0,841 | 0,435 | 0,001208808 | 2 | Dbn1 |
| Lama4 | 8,24E-08 | 0,455072423 | 0,921 | 0,562 | 0,001305527 | 2 | Lama4 |
| Anxa1 | 8,28E-08 | 0,450509834 | 0,921 | 0,65 | 0,001313035 | 2 | Anxa1 |
| Tpm3 | 9,03E-08 | 0,384462235 | 1 | 0,942 | 0,001431964 | 2 | Tpm3 |
| Ethe1 | 9,47E-08 | 0,320372121 | 0,476 | 0,169 | 0,001500786 | 2 | Ethe1 |
| Calm1 | 9,51E-08 | 0,335028681 | 1 | 0,992 | 0,001507248 | 2 | Calm1 |
| Copb2 | 9,89E-08 | 0,504641018 | 0,746 | 0,423 | 0,001567312 | 2 | Copb2 |
| Piezo2 | 1,12E-07 | 0,375815664 | 0,841 | 0,446 | 0,001767583 | 2 | Piezo2 |
| Ercc1 | 1,25E-07 | 0,398972003 | 0,651 | 0,331 | 0,001987523 | 2 | Ercc1 |
| Tcf4 | 1,33E-07 | 0,371398623 | 1 | 0,854 | 0,002115851 | 2 | Tcf4 |
| Ddah2 | 1,38E-07 | 0,435175112 | 0,746 | 0,385 | 0,002185895 | 2 | Ddah2 |
| Tpm4 | 1,49E-07 | 0,433753109 | 1 | 0,908 | 0,002356045 | 2 | Tpm4 |
| Cd341 | 1,50E-07 | 0,527304063 | 1 | 0,685 | 0,002375835 | 2 | Cd34 |
| Mmp15 | 1,55E-07 | 0,274167272 | 0,651 | 0,277 | 0,002450296 | 2 | Mmp15 |
| Cx3cl1 | 1,82E-07 | 0,574283387 | 0,429 | 0,154 | 0,0028827 | 2 | Cx3cl1 |
| Dstn | 1,83E-07 | 0,464154755 | 0,952 | 0,765 | 0,002904384 | 2 | Dstn |
| Smad1 | 1,88E-07 | 0,579380837 | 0,825 | 0,538 | 0,002986634 | 2 | Smad1 |
| Cpe1 | 2,03E-07 | 0,578959355 | 0,889 | 0,523 | 0,003220583 | 2 | Cpe |
| Itga5 | 2,23E-07 | 0,454197275 | 0,841 | 0,577 | 0,003542347 | 2 | Itga5 |
| Pald1 | 2,33E-07 | 0,497618925 | 0,794 | 0,481 | 0,003699368 | 2 | Pald1 |
| Chst7 | 2,39E-07 | 0,379934305 | 0,413 | 0,127 | 0,00378255 | 2 | Chst7 |
| Clec1a | 2,40E-07 | 0,292412824 | 0,698 | 0,331 | 0,00381231 | 2 | Clec1a |
| Rbp1 | 2,70E-07 | 0,399058734 | 0,921 | 0,554 | 0,004272744 | 2 | Rbp1 |
| Tnfaip11 | 2,77E-07 | 0,48420966 | 0,905 | 0,627 | 0,004385804 | 2 | Tnfaip1 |
| Eef1a1 | 2,81E-07 | 0,262297406 | 1 | 1 | 0,004451501 | 2 | Eef1a1 |
| Kitl | 2,85E-07 | 0,418158994 | 0,603 | 0,262 | 0,00451677 | 2 | Kitl |
| Gimap41 | 2,89E-07 | 0,470178406 | 0,889 | 0,546 | 0,004583496 | 2 | Gimap4 |
| Arf1 | 2,94E-07 | 0,386830352 | 1 | 0,923 | 0,004655078 | 2 | Arf1 |
| Atp9a | 2,98E-07 | 0,420857282 | 0,54 | 0,219 | 0,004718035 | 2 | Atp9a |
| Plxnd1 | 3,26E-07 | 0,391023654 | 1 | 0,788 | 0,005163712 | 2 | Plxnd1 |
| Ralb | 3,41E-07 | 0,347607365 | 0,984 | 0,804 | 0,005407961 | 2 | Ralb |
| Tmsb101 | 4,28E-07 | 0,416217081 | 1 | 0,969 | 0,006778766 | 2 | Tmsb10 |
| Lcp21 | 4,41E-07 | 0,522378495 | 0,714 | 0,423 | 0,006983359 | 2 | Lcp2 |
| Dusp2 | 4,50E-07 | 0,452973621 | 0,714 | 0,365 | 0,00713855 | 2 | Dusp2 |
| Thsd1 | 4,70E-07 | 0,44503263 | 0,746 | 0,396 | 0,007443583 | 2 | Thsd1 |
| Kcne3 | 5,15E-07 | 0,402642037 | 0,825 | 0,396 | 0,008167865 | 2 | Kcne3 |
| Tgfb1i1 | 5,20E-07 | 0,384586798 | 0,651 | 0,296 | 0,008247187 | 2 | Tgfb1i1 |
| Arl1 | 5,57E-07 | 0,336785379 | 0,841 | 0,5 | 0,008824275 | 2 | Arl1 |
| Msn | 6,62E-07 | 0,421694843 | 1 | 0,958 | 0,010500046 | 2 | Msn |
| Gpr4 | 7,35E-07 | 0,326513363 | 0,873 | 0,542 | 0,011649234 | 2 | Gpr4 |
| Kank31 | 7,39E-07 | 0,323669044 | 0,762 | 0,412 | 0,011716653 | 2 | Kank3 |
| Lamb1 | 7,87E-07 | 0,339038816 | 0,968 | 0,588 | 0,012480814 | 2 | Lamb1 |
| Spata6 | 8,05E-07 | 0,362581831 | 0,651 | 0,315 | 0,012760906 | 2 | Spata6 |

| | | | | | | | |
|---|---|---|---|---|---|---|---|
| Ranbp3 | 0,002057836 | 0,266840045 | 0,415 | 0,253 | 1 | T_EC | Ranbp3 |
| Hadhb | 0,002066665 | 0,261336463 | 0,585 | 0,415 | 1 | T_EC | Hadhb |
| Ddx58 | 0,002177872 | 0,332001328 | 0,415 | 0,266 | 1 | T_EC | Ddx58 |
| Smu1 | 0,002295399 | 0,30591275 | 0,549 | 0,378 | 1 | T_EC | Smu1 |
| Mars | 0,002371589 | 0,30893894 | 0,341 | 0,195 | 1 | T_EC | Mars |
| Apba3 | 0,002381846 | 0,260663855 | 0,317 | 0,17 | 1 | T_EC | Apba3 |
| Bsg | 0,002438888 | 0,294328032 | 0,963 | 0,917 | 1 | T_EC | Bsg |
| Anapc5 | 0,002486652 | 0,260290973 | 0,768 | 0,651 | 1 | T_EC | Anapc5 |
| Pald1 | 0,002494374 | 0,330978552 | 0,646 | 0,506 | 1 | T_EC | Pald1 |
| Zc3h15 | 0,002524294 | 0,371452442 | 0,537 | 0,386 | 1 | T_EC | Zc3h15 |
| Glrx3 | 0,002812966 | 0,272334563 | 0,695 | 0,519 | 1 | T_EC | Glrx3 |
| Eif1a | 0,002841848 | 0,279361232 | 0,585 | 0,423 | 1 | T_EC | Eif1a |
| Chtop | 0,003064956 | 0,257093086 | 0,561 | 0,398 | 1 | T_EC | Chtop |
| Rhot1 | 0,003066525 | 0,27545181 | 0,354 | 0,199 | 1 | T_EC | Rhot1 |
| Il11ra1 | 0,003100966 | 0,25708298 | 0,366 | 0,203 | 1 | T_EC | Il11ra1 |
| Myl12b | 0,003122161 | 0,252869909 | 0,976 | 0,917 | 1 | T_EC | Myl12b |
| Angptl4 | 0,003336935 | 0,287971047 | 0,341 | 0,178 | 1 | T_EC | Angptl4 |
| Ddx39 | 0,003383477 | 0,265711517 | 0,683 | 0,535 | 1 | T_EC | Ddx39 |
| Eif3l | 0,003538695 | 0,251386695 | 0,695 | 0,548 | 1 | T_EC | Eif3l |
| Ssb | 0,00382179 | 0,294950634 | 0,793 | 0,685 | 1 | T_EC | Ssb |
| Maged2 | 0,003845653 | 0,296932141 | 0,39 | 0,249 | 1 | T_EC | Maged2 |
| Dgcr2 | 0,00422467 | 0,304362548 | 0,524 | 0,361 | 1 | T_EC | Dgcr2 |
| Smarcb1 | 0,004267479 | 0,289042761 | 0,488 | 0,361 | 1 | T_EC | Smarcb1 |
| Rcn2 | 0,004622575 | 0,311327985 | 0,634 | 0,502 | 1 | T_EC | Rcn2 |
| Ssr3 | 0,004625689 | 0,280728694 | 0,89 | 0,755 | 1 | T_EC | Ssr3 |
| Vcp | 0,004629361 | 0,282905943 | 0,805 | 0,755 | 1 | T_EC | Vcp |
| Zfp68 | 0,00472084 | 0,266665457 | 0,305 | 0,162 | 1 | T_EC | Zfp68 |
| Wdr77 | 0,004817554 | 0,323065319 | 0,341 | 0,203 | 1 | T_EC | Wdr77 |
| Mat2a | 0,004897053 | 0,27039299 | 0,646 | 0,494 | 1 | T_EC | Mat2a |
| Psmd2 | 0,004990883 | 0,270898678 | 0,695 | 0,589 | 1 | T_EC | Psmd2 |
| Ube3c | 0,005234808 | 0,269117021 | 0,366 | 0,216 | 1 | T_EC | Ube3c |
| Api5 | 0,00588477 | 0,270498292 | 0,634 | 0,506 | 1 | T_EC | Api5 |
| Prcp | 0,006288544 | 0,332971686 | 0,756 | 0,685 | 1 | T_EC | Prcp |
| Usp4 | 0,006851996 | 0,295390248 | 0,512 | 0,369 | 1 | T_EC | Usp4 |
| Fbxl5 | 0,007021882 | 0,354376836 | 0,463 | 0,344 | 1 | T_EC | Fbxl5 |
| Ctso | 0,007105881 | 0,253758368 | 0,329 | 0,191 | 1 | T_EC | Ctso |
| Ctgf | 0,007373529 | 0,519423125 | 0,415 | 0,257 | 1 | T_EC | Ctgf |
| Slc46a3 | 0,007738888 | 0,289482029 | 0,305 | 0,178 | 1 | T_EC | Slc46a3 |
| Ehd4 | 0,008048354 | 0,272906161 | 0,89 | 0,739 | 1 | T_EC | Ehd4 |
| Mmp14 | 0,008123814 | 0,280902257 | 0,915 | 0,846 | 1 | T_EC | Mmp14 |
| Odc1 | 0,008316062 | 0,300607112 | 0,585 | 0,448 | 1 | T_EC | Odc1 |
| Rnf213 | 0,008896741 | 0,327192935 | 0,561 | 0,419 | 1 | T_EC | Rnf213 |
| Azin1 | 0,009195984 | 0,468449122 | 0,439 | 0,282 | 1 | T_EC | Azin1 |
| mt-Rnr1 | 2,29E-26 | 0,81308994 | 1 | 1 | 3,63E-22 | T_CLEC | mt-Rnr1 |
| S1pr1 | 3,95E-24 | 0,933278024 | 0,988 | 0,552 | 6,26E-20 | T_CLEC | S1pr1 |
| Gng11 | 5,71E-24 | 0,963026124 | 0,975 | 0,528 | 9,06E-20 | T_CLEC | Gng11 |
| Adgrf5 | 8,71E-23 | 1,057222851 | 0,925 | 0,442 | 1,38E-18 | T_CLEC | Adgrf5 |
| Plvap | 1,54E-22 | 1,093123712 | 0,944 | 0,436 | 2,45E-18 | T_CLEC | Plvap |
| Malat1 | 5,93E-22 | 1,047396874 | 0,994 | 1 | 9,41E-18 | T_CLEC | Malat1 |
| Col4a2 | 1,25E-21 | 0,870300769 | 1 | 0,54 | 1,99E-17 | T_CLEC | Col4a2 |
| Cdh13 | 2,97E-21 | 1,03353919 | 0,906 | 0,393 | 4,70E-17 | T_CLEC | Cdh13 |
| Col4a1 | 1,32E-19 | 0,812560223 | 0,994 | 0,564 | 2,10E-15 | T_CLEC | Col4a1 |
| Gm37376 | 1,54E-19 | 0,999218131 | 0,988 | 0,988 | 2,44E-15 | T_CLEC | Gm37376 |
| Flt4 | 2,71E-19 | 0,859119683 | 0,844 | 0,454 | 4,29E-15 | T_CLEC | Flt4 |
| Hspg2 | 4,22E-19 | 0,781529612 | 0,956 | 0,46 | 6,69E-15 | T_CLEC | Hspg2 |
| Cdh5 | 1,15E-18 | 0,683474182 | 0,975 | 0,497 | 1,83E-14 | T_CLEC | Cdh5 |
| Sema3f | 2,55E-18 | 0,655028031 | 0,862 | 0,38 | 4,04E-14 | T_CLEC | Sema3f |
| Sptbn1 | 3,65E-18 | 0,810687416 | 0,981 | 0,632 | 5,78E-14 | T_CLEC | Sptbn1 |
| Ets1 | 5,77E-18 | 0,80075104 | 0,944 | 0,528 | 9,14E-14 | T_CLEC | Ets1 |
| Gpihbp1 | 1,04E-17 | 1,153867816 | 0,756 | 0,307 | 1,65E-13 | T_CLEC | Gpihbp1 |
| Lrrc8a | 4,74E-17 | 0,845816575 | 0,912 | 0,663 | 7,51E-13 | T_CLEC | Lrrc8a |
| Pecam1 | 7,84E-17 | 0,725161956 | 0,981 | 0,564 | 1,24E-12 | T_CLEC | Pecam1 |
| AY036118 | 2,83E-16 | 0,472079981 | 1 | 1 | 4,49E-12 | T_CLEC | AY036118 |
| Ahnak | 8,17E-16 | 0,728874009 | 0,869 | 0,521 | 1,29E-11 | T_CLEC | Ahnak |
| Heg1 | 8,58E-16 | 0,725091534 | 0,912 | 0,472 | 1,36E-11 | T_CLEC | Heg1 |
| Ece11 | 1,26E-15 | 0,597328385 | 1 | 0,54 | 2,00E-11 | T_CLEC | Ece1 |
| Sparcl11 | 2,90E-15 | 0,710229414 | 0,944 | 0,509 | 4,60E-11 | T_CLEC | Sparcl1 |
| Lama4 | 3,56E-15 | 0,710526894 | 0,856 | 0,411 | 5,64E-11 | T_CLEC | Lama4 |
| Cd93 | 3,80E-15 | 0,6694057 | 0,994 | 0,865 | 6,03E-11 | T_CLEC | Cd93 |
| Lrg11 | 3,92E-15 | 0,776991566 | 0,956 | 0,485 | 6,21E-11 | T_CLEC | Lrg1 |
| Insr | 6,46E-15 | 0,73003272 | 0,806 | 0,423 | 1,02E-10 | T_CLEC | Insr |
| Kdr | 6,67E-15 | 0,696572223 | 0,894 | 0,436 | 1,06E-10 | T_CLEC | Kdr |
| Emcn | 9,74E-15 | 0,749146708 | 0,938 | 0,509 | 1,54E-10 | T_CLEC | Emcn |
| Tcf4 | 1,07E-14 | 0,648588086 | 0,988 | 0,779 | 1,70E-10 | T_CLEC | Tcf4 |
| Crip21 | 2,00E-14 | 0,648529373 | 0,925 | 0,534 | 3,18E-10 | T_CLEC | Crip2 |
| Tpm41 | 7,17E-14 | 0,574379727 | 0,988 | 0,865 | 1,14E-09 | T_CLEC | Tpm4 |
| Col18a1 | 9,33E-14 | 0,749285745 | 0,794 | 0,399 | 1,48E-09 | T_CLEC | Col18a1 |
| Sept2 | 1,07E-13 | 0,686184369 | 0,969 | 0,951 | 1,69E-09 | T_CLEC | Sept2 |
| Lamc1 | 1,11E-13 | 0,686985287 | 0,888 | 0,485 | 1,76E-09 | T_CLEC | Lamc1 |
| Flt11 | 1,11E-13 | 0,594394715 | 0,956 | 0,503 | 1,76E-09 | T_CLEC | Flt1 |
| Cyyr1 | 1,12E-13 | 0,724119727 | 0,825 | 0,442 | 1,78E-09 | T_CLEC | Cyyr1 |
| Rbp1 | 1,29E-13 | 0,621627668 | 0,85 | 0,405 | 2,04E-09 | T_CLEC | Rbp1 |
| Eng1 | 1,38E-13 | 0,624551948 | 0,981 | 0,632 | 2,18E-09 | T_CLEC | Eng |
| Tmsb101 | 1,44E-13 | 0,509184651 | 1 | 0,951 | 2,28E-09 | T_CLEC | Tmsb10 |
| Timp31 | 1,52E-13 | 0,559223638 | 0,856 | 0,448 | 2,40E-09 | T_CLEC | Timp3 |
| Podxl | 1,60E-13 | 0,504959531 | 0,819 | 0,374 | 2,54E-09 | T_CLEC | Podxl |
| Lamb1 | 1,92E-13 | 0,701250924 | 0,869 | 0,46 | 3,04E-09 | T_CLEC | Lamb1 |
| Sparc1 | 3,00E-13 | 0,666475156 | 1 | 0,589 | 4,76E-09 | T_CLEC | Sparc |
| Aqp1 | 3,93E-13 | 0,818001907 | 0,344 | 0,031 | 6,23E-09 | T_CLEC | Aqp1 |
| Serpinh1 | 4,80E-13 | 0,461652542 | 0,938 | 0,503 | 7,61E-09 | T_CLEC | Serpinh1 |
| Gimap61 | 5,70E-13 | 0,642261641 | 0,919 | 0,509 | 9,03E-09 | T_CLEC | Gimap6 |
| Nid1 | 7,80E-13 | 0,621257242 | 0,888 | 0,491 | 1,24E-08 | T_CLEC | Nid1 |
| Fabp4 | 8,58E-13 | 0,88242415 | 0,344 | 0,037 | 1,36E-08 | T_CLEC | Fabp4 |
| Pam | 1,31E-12 | 0,379525636 | 0,95 | 0,681 | 2,07E-08 | T_CLEC | Pam |
| Abcc9 | 1,77E-12 | 0,864876323 | 0,606 | 0,227 | 2,80E-08 | T_CLEC | Abcc9 |
| Ppic1 | 1,94E-12 | 0,561824013 | 0,938 | 0,509 | 3,07E-08 | T_CLEC | Ppic |
| Ednrb | 1,96E-12 | 0,883897344 | 0,644 | 0,264 | 3,11E-08 | T_CLEC | Ednrb |
| Gja11 | 2,17E-12 | 0,580685533 | 0,869 | 0,454 | 3,43E-08 | T_CLEC | Gja1 |
| Cald1 | 2,25E-12 | 0,550439359 | 0,844 | 0,472 | 3,56E-08 | T_CLEC | Cald1 |
| Olfml2a | 2,55E-12 | 0,715108592 | 0,519 | 0,172 | 4,05E-08 | T_CLEC | Olfml2a |
| Ywhaz | 2,64E-12 | 0,487502115 | 1 | 0,988 | 4,18E-08 | T_CLEC | Ywhaz |
| Fscn1 | 2,96E-12 | 0,606873093 | 0,912 | 0,515 | 4,70E-08 | T_CLEC | Fscn1 |
| Jup | 3,82E-12 | 0,66745007 | 0,9 | 0,571 | 6,06E-08 | T_CLEC | Jup |
| Plpp1 | 3,99E-12 | 0,744118187 | 0,812 | 0,466 | 6,33E-08 | T_CLEC | Plpp1 |
| Grb10 | 4,09E-12 | 0,660519572 | 0,625 | 0,27 | 6,48E-08 | T_CLEC | Grb10 |
| Ptgr1 | 4,24E-12 | 1,063582916 | 0,7 | 0,393 | 6,72E-08 | T_CLEC | Ptgr1 |
| Fstl1 | 5,38E-12 | 0,621109317 | 0,725 | 0,344 | 8,53E-08 | T_CLEC | Fstl1 |
| Ramp3 | 6,18E-12 | 0,791127318 | 0,494 | 0,141 | 9,80E-08 | T_CLEC | Ramp3 |
| Zeb1 | 6,23E-12 | 0,658743109 | 0,662 | 0,344 | 9,88E-08 | T_CLEC | Zeb1 |
| Ldb2 | 8,08E-12 | 0,528778092 | 0,819 | 0,436 | 1,28E-07 | T_CLEC | Ldb2 |
| Cd2001 | 1,01E-11 | 0,503412191 | 1 | 0,509 | 1,61E-07 | T_CLEC | Cd200 |
| Mcam1 | 1,04E-11 | 0,492840524 | 0,944 | 0,521 | 1,65E-07 | T_CLEC | Mcam |
| Ptprb | 1,08E-11 | 0,614548448 | 0,856 | 0,466 | 1,71E-07 | T_CLEC | Ptprb |
| Slco2a1 | 1,34E-11 | 0,529877842 | 0,744 | 0,356 | 2,12E-07 | T_CLEC | Slco2a1 |
| Ppfibp1 | 2,03E-11 | 0,557509781 | 0,825 | 0,509 | 3,21E-07 | T_CLEC | Ppfibp1 |
| Mmrn2 | 2,41E-11 | 0,64425187 | 0,794 | 0,417 | 3,83E-07 | T_CLEC | Mmrn2 |
| Anxa21 | 2,72E-11 | 0,508225134 | 0,975 | 0,767 | 4,31E-07 | T_CLEC | Anxa2 |
| Ecscr1 | 3,26E-11 | 0,546572318 | 0,956 | 0,509 | 5,17E-07 | T_CLEC | Ecscr |
| Nfib | 3,29E-11 | 0,588688742 | 0,794 | 0,454 | 5,21E-07 | T_CLEC | Nfib |
| Nos3 | 3,48E-11 | 0,465241915 | 0,8 | 0,393 | 5,51E-07 | T_CLEC | Nos3 |
| Plec | 3,83E-11 | 0,542649557 | 0,919 | 0,644 | 6,08E-07 | T_CLEC | Plec |

| | | | | | | | |
|---|---|---|---|---|---|---|---|
| Npdc11 | 8,18E-07 | 0,327937011 | 0,81 | 0,431 | 0,012968633 | 2 | Npdc1 |
| Fam129a | 8,18E-07 | 0,317329168 | 0,46 | 0,169 | 0,012973754 | 2 | Fam129a |
| H2-Q7 | 9,10E-07 | 0,528451652 | 0,873 | 0,604 | 0,0144191 | 2 | H2-Q7 |
| Fam43a | 9,45E-07 | 0,472280008 | 0,683 | 0,377 | 0,014975638 | 2 | Fam43a |
| Rela | 1,02E-06 | 0,377519284 | 0,746 | 0,462 | 0,016223463 | 2 | Rela |
| Pls31 | 1,15E-06 | 0,385414762 | 0,841 | 0,527 | 0,018269637 | 2 | Pls3 |
| Pdlim7 | 1,16E-06 | 0,337906354 | 0,968 | 0,669 | 0,018421085 | 2 | Pdlim7 |
| Tspan18 | 1,17E-06 | 0,305436177 | 0,857 | 0,485 | 0,018478324 | 2 | Tspan18 |
| Cdc42ep4 | 1,18E-06 | 0,36475443 | 0,556 | 0,25 | 0,018754019 | 2 | Cdc42ep4 |
| Cdc42ep1 | 1,21E-06 | 0,371350763 | 0,73 | 0,392 | 0,019139329 | 2 | Cdc42ep1 |
| Ackr3 | 1,24E-06 | 0,399265824 | 0,714 | 0,35 | 0,01958391 | 2 | Ackr3 |
| Chst1 | 1,35E-06 | 0,365323548 | 0,524 | 0,231 | 0,021448848 | 2 | Chst1 |
| Angpt2 | 1,37E-06 | 0,626271416 | 0,508 | 0,212 | 0,021769401 | 2 | Angpt2 |
| Aldoa | 1,57E-06 | 0,379426 | 0,984 | 0,95 | 0,024962939 | 2 | Aldoa |
| Lpin3 | 1,74E-06 | 0,393801801 | 0,397 | 0,138 | 0,027547412 | 2 | Lpin3 |
| Abhd4 | 1,82E-06 | 0,529650577 | 0,635 | 0,354 | 0,028899458 | 2 | Abhd4 |
| Eif4a-ps4 | 1,89E-06 | 0,263216019 | 0,841 | 0,523 | 0,029967378 | 2 | Eif4a-ps4 |
| Tspan15 | 1,92E-06 | 0,333712066 | 0,635 | 0,304 | 0,030398667 | 2 | Tspan15 |
| Exoc3l | 1,93E-06 | 0,359582539 | 0,603 | 0,277 | 0,03065834 | 2 | Exoc3l |
| Aplnr1 | 1,94E-06 | 0,34804054 | 0,889 | 0,485 | 0,030810128 | 2 | Aplnr |
| Fzd4 | 2,00E-06 | 0,397519786 | 0,762 | 0,435 | 0,031710255 | 2 | Fzd4 |
| Cttn1 | 2,07E-06 | 0,297706935 | 0,73 | 0,419 | 0,032824628 | 2 | Cttn |
| Mtx2 | 2,10E-06 | 0,286498736 | 0,619 | 0,288 | 0,033318419 | 2 | Mtx2 |
| Lxn | 2,36E-06 | 0,441332581 | 0,889 | 0,577 | 0,037365582 | 2 | Lxn |
| Myl6 | 2,37E-06 | 0,274186435 | 1 | 0,992 | 0,037503431 | 2 | Myl6 |
| Ppp1r16b | 2,46E-06 | 0,257327348 | 0,794 | 0,504 | 0,039042673 | 2 | Ppp1r16b |
| Mfge8 | 2,47E-06 | 0,390592576 | 0,873 | 0,527 | 0,039116994 | 2 | Mfge8 |
| Tmem254c | 2,48E-06 | 0,265779706 | 0,762 | 0,404 | 0,03925437 | 2 | Tmem254c |
| Tmem123 | 2,50E-06 | 0,379246464 | 0,841 | 0,581 | 0,039703202 | 2 | Tmem123 |
| Kctd10 | 2,51E-06 | 0,39490959 | 0,698 | 0,346 | 0,039754907 | 2 | Kctd10 |
| Plat | 2,72E-06 | 0,36611283 | 0,698 | 0,338 | 0,043083162 | 2 | Plat |
| Sec22b | 2,78E-06 | 0,297017516 | 0,794 | 0,454 | 0,044023483 | 2 | Sec22b |
| Lrrc8c | 2,86E-06 | 0,408629759 | 0,841 | 0,562 | 0,045303937 | 2 | Lrrc8c |
| Trp53 | 2,88E-06 | 0,442234924 | 0,794 | 0,485 | 0,0456542 | 2 | Trp53 |
| Sparcl1 | 2,89E-06 | 0,37406883 | 1 | 0,658 | 0,04584764 | 2 | Sparcl1 |
| Nedd41 | 3,00E-06 | 0,334812558 | 0,984 | 0,654 | 0,047483703 | 2 | Nedd4 |
| Adamts7 | 3,02E-06 | 0,328187509 | 0,413 | 0,154 | 0,047923775 | 2 | Adamts7 |
| Map4k5 | 3,30E-06 | 0,428952895 | 0,714 | 0,431 | 0,05234809 | 2 | Map4k5 |
| Rsu1 | 3,42E-06 | 0,411523022 | 0,921 | 0,696 | 0,054212554 | 2 | Rsu1 |
| Lypla2 | 3,48E-06 | 0,260968414 | 0,73 | 0,435 | 0,055098353 | 2 | Lypla2 |
| B3gnt31 | 3,59E-06 | 0,353005512 | 0,841 | 0,496 | 0,056975295 | 2 | B3gnt3 |
| Schip1 | 3,59E-06 | 0,251500616 | 0,841 | 0,496 | 0,056975295 | 2 | Schip1 |
| Surf4 | 3,65E-06 | 0,507699659 | 0,857 | 0,704 | 0,057913634 | 2 | Surf4 |
| Pfkp | 3,69E-06 | 0,450320751 | 0,683 | 0,404 | 0,058488157 | 2 | Pfkp |
| Hes1 | 3,70E-06 | 0,333943021 | 0,841 | 0,515 | 0,058593303 | 2 | Hes1 |
| Ptrf | 3,82E-06 | 0,287609987 | 0,968 | 0,638 | 0,060529479 | 2 | Ptrf |
| Dad1 | 3,98E-06 | 0,294029394 | 0,968 | 0,827 | 0,063162596 | 2 | Dad1 |
| Lpcat4 | 4,16E-06 | 0,262911647 | 0,413 | 0,154 | 0,06599397 | 2 | Lpcat4 |
| Lhfp | 4,18E-06 | 0,338497084 | 0,286 | 0,081 | 0,066283578 | 2 | Lhfp |
| Nes | 4,73E-06 | 0,378845789 | 0,587 | 0,304 | 0,074908873 | 2 | Nes |
| Kdelr3 | 4,87E-06 | 0,315991854 | 0,429 | 0,165 | 0,077227638 | 2 | Kdelr3 |
| Ccdc85b | 4,98E-06 | 0,327141379 | 0,778 | 0,504 | 0,078967705 | 2 | Ccdc85b |
| Cfl2 | 5,02E-06 | 0,337261529 | 0,587 | 0,296 | 0,079652772 | 2 | Cfl2 |
| Atp5b | 5,16E-06 | 0,387504771 | 1 | 0,9 | 0,081806253 | 2 | Atp5b |
| Capg | 5,20E-06 | 0,352675289 | 0,968 | 0,731 | 0,082372375 | 2 | Capg |
| Synpo | 5,21E-06 | 0,277133531 | 0,762 | 0,446 | 0,082598099 | 2 | Synpo |
| Smg6 | 5,24E-06 | 0,39993007 | 0,571 | 0,273 | 0,083126485 | 2 | Smg6 |
| Cd2001 | 5,34E-06 | 0,357092173 | 1 | 0,692 | 0,084610735 | 2 | Cd200 |
| Nostrin1 | 5,59E-06 | 0,283099704 | 0,857 | 0,488 | 0,088583518 | 2 | Nostrin |
| Lgals9 | 5,62E-06 | 0,418757755 | 1 | 0,896 | 0,089071704 | 2 | Lgals9 |
| Actn4 | 5,63E-06 | 0,350200975 | 0,968 | 0,804 | 0,089202912 | 2 | Actn4 |
| Oaf | 5,64E-06 | 0,322708162 | 0,698 | 0,369 | 0,089467984 | 2 | Oaf |
| Map1b | 5,86E-06 | 0,296314417 | 0,73 | 0,388 | 0,092878508 | 2 | Map1b |
| Pja1 | 5,89E-06 | 0,381382151 | 0,381 | 0,135 | 0,093331239 | 2 | Pja1 |
| F11r | 6,01E-06 | 0,388723575 | 1 | 0,854 | 0,095259633 | 2 | F11r |
| Sept41 | 6,05E-06 | 0,347656891 | 0,714 | 0,373 | 0,095879655 | 2 | Sept4 |
| Sh3bgrl | 6,23E-06 | 0,42149577 | 0,921 | 0,769 | 0,098834125 | 2 | Sh3bgrl |
| Ptprk | 6,37E-06 | 0,278588801 | 0,746 | 0,396 | 0,101000016 | 2 | Ptprk |
| Eif5a | 6,39E-06 | 0,357519028 | 0,984 | 0,935 | 0,101324743 | 2 | Eif5a |
| Snrk | 6,42E-06 | 0,443148812 | 0,889 | 0,638 | 0,101709505 | 2 | Snrk |
| Yipf1 | 6,43E-06 | 0,372254106 | 0,524 | 0,254 | 0,101907584 | 2 | Yipf1 |
| Stap2 | 6,84E-06 | 0,321452232 | 0,667 | 0,338 | 0,108393191 | 2 | Stap2 |
| Mapk6 | 7,15E-06 | 0,31338828 | 0,683 | 0,388 | 0,113367502 | 2 | Mapk6 |
| Pcdh1 | 7,17E-06 | 0,285974951 | 0,683 | 0,385 | 0,113627921 | 2 | Pcdh1 |
| Arhgap18 | 7,56E-06 | 0,449052888 | 0,698 | 0,396 | 0,11987973 | 2 | Arhgap18 |
| Prkar1a | 7,64E-06 | 0,428450698 | 0,968 | 0,881 | 0,121047856 | 2 | Prkar1a |
| Rcn1 | 7,94E-06 | 0,295857084 | 0,476 | 0,215 | 0,125832157 | 2 | Rcn1 |
| Hyal2 | 8,06E-06 | 0,295659136 | 0,698 | 0,358 | 0,127758403 | 2 | Hyal2 |
| F2rl3 | 8,40E-06 | 0,254440367 | 0,349 | 0,115 | 0,133197362 | 2 | F2rl3 |
| Eml1 | 8,55E-06 | 0,290413855 | 0,73 | 0,381 | 0,135497952 | 2 | Eml1 |
| Eif6 | 8,90E-06 | 0,331973913 | 0,81 | 0,569 | 0,141045678 | 2 | Eif6 |
| Morf4l2 | 9,87E-06 | 0,410183765 | 0,921 | 0,785 | 0,156463581 | 2 | Morf4l2 |
| Nfkbie | 9,88E-06 | 0,261794827 | 0,333 | 0,108 | 0,15668882 | 2 | Nfkbie |
| Ppic1 | 1,00E-05 | 0,36054847 | 0,968 | 0,662 | 0,158623001 | 2 | Ppic |
| Cdc42ep3 | 1,04E-05 | 0,371898845 | 0,73 | 0,388 | 0,165207585 | 2 | Cdc42ep3 |
| Ifitm31 | 1,06E-05 | 0,375684645 | 1 | 0,938 | 0,168579094 | 2 | Ifitm3 |
| 1810011O10Rik | 1,11E-05 | 0,382570746 | 0,524 | 0,25 | 0,176427927 | 2 | 1810011O10Rik |
| Pqbp1 | 1,16E-05 | 0,268842147 | 0,667 | 0,35 | 0,183152418 | 2 | Pqbp1 |
| 9430020K01Rik | 1,19E-05 | 0,296906512 | 0,873 | 0,523 | 0,188347281 | 2 | 9430020K01Rik |
| Cct5 | 1,27E-05 | 0,34906159 | 0,921 | 0,692 | 0,201194404 | 2 | Cct5 |
| Tmem881 | 1,30E-05 | 0,289089723 | 0,984 | 0,627 | 0,205801933 | 2 | Tmem88 |
| Sec13 | 1,31E-05 | 0,319717692 | 0,81 | 0,558 | 0,206926299 | 2 | Sec13 |
| 4930523C07Rik | 1,38E-05 | 0,276423126 | 0,413 | 0,162 | 0,218435326 | 2 | 4930523C07Rik |
| Gnb4 | 1,39E-05 | 0,302869799 | 0,714 | 0,385 | 0,221085393 | 2 | Gnb4 |
| Bin1 | 1,41E-05 | 0,288607044 | 0,524 | 0,238 | 0,222747217 | 2 | Bin1 |
| Fzr1 | 1,46E-05 | 0,255321369 | 0,444 | 0,185 | 0,231028972 | 2 | Fzr1 |
| Galnt1 | 1,48E-05 | 0,339795944 | 0,889 | 0,638 | 0,234255702 | 2 | Galnt1 |
| Itgb3 | 1,53E-05 | 0,341216174 | 0,587 | 0,319 | 0,242392545 | 2 | Itgb3 |
| Ctnna1 | 1,58E-05 | 0,356030103 | 0,968 | 0,762 | 0,251121105 | 2 | Ctnna1 |
| N4bp3 | 1,60E-05 | 0,291532519 | 0,667 | 0,354 | 0,254143027 | 2 | N4bp3 |
| Ece11 | 1,62E-05 | 0,311733119 | 1 | 0,712 | 0,256618739 | 2 | Ece1 |
| Laptm4b | 1,62E-05 | 0,27565182 | 0,508 | 0,235 | 0,256865563 | 2 | Laptm4b |
| Ubtd1 | 1,62E-05 | 0,319024551 | 0,619 | 0,358 | 0,257402706 | 2 | Ubtd1 |
| Serpinb6b | 1,67E-05 | 0,256945043 | 0,317 | 0,104 | 0,263954398 | 2 | Serpinb6b |
| Sox17 | 1,82E-05 | 0,321910329 | 0,508 | 0,25 | 0,288700392 | 2 | Sox17 |
| Zbtb20 | 1,82E-05 | 0,309782766 | 0,778 | 0,523 | 0,2892054 | 2 | Zbtb20 |
| Pde2a | 1,85E-05 | 0,365538896 | 0,635 | 0,35 | 0,293582308 | 2 | Pde2a |
| Rmnd5b | 1,87E-05 | 0,273387714 | 0,556 | 0,269 | 0,296771042 | 2 | Rmnd5b |
| Lpar6 | 1,95E-05 | 0,388013264 | 0,746 | 0,431 | 0,309422745 | 2 | Lpar6 |
| Tmem120b | 2,01E-05 | 0,32840121 | 0,349 | 0,127 | 0,317846147 | 2 | Tmem120b |
| Tes | 2,04E-05 | 0,283577552 | 0,937 | 0,669 | 0,323407019 | 2 | Tes |
| Rbpms | 2,12E-05 | 0,286753512 | 0,921 | 0,6 | 0,33661471 | 2 | Rbpms |
| Nrep | 2,13E-05 | 0,283820777 | 0,762 | 0,45 | 0,33841234 | 2 | Nrep |
| Cyp2j6 | 2,21E-05 | 0,371895166 | 0,317 | 0,112 | 0,350228658 | 2 | Cyp2j6 |
| Rab1a | 2,27E-05 | 0,2958709 | 0,968 | 0,804 | 0,359059151 | 2 | Rab1a |
| Ptp4a3 | 2,33E-05 | 0,324327157 | 0,841 | 0,558 | 0,370008189 | 2 | Ptp4a3 |
| Dcaf13 | 2,40E-05 | 0,304178037 | 0,556 | 0,285 | 0,380736648 | 2 | Dcaf13 |
| Cdk4 | 2,46E-05 | 0,321816654 | 0,873 | 0,623 | 0,389574952 | 2 | Cdk4 |
| Sh3bp4 | 2,60E-05 | 0,461029998 | 0,476 | 0,215 | 0,412300589 | 2 | Sh3bp4 |
| Hid1 | 2,61E-05 | 0,260626079 | 0,317 | 0,108 | 0,412964777 | 2 | Hid1 |
| S100a11 | 2,62E-05 | 0,401060196 | 0,984 | 0,823 | 0,415597091 | 2 | S100a11 |

| | | | | | | | |
|---|---|---|---|---|---|---|---|
| Plxnd1 | 4,39E-11 | 0,473421831 | 0,956 | 0,706 | 6,96E-07 | T_CLEC | Plxnd1 |
| Efna11 | 5,81E-11 | 0,567177011 | 0,844 | 0,454 | 9,21E-07 | T_CLEC | Efna1 |
| Epas11 | 8,38E-11 | 0,488914665 | 0,844 | 0,454 | 1,33E-06 | T_CLEC | Epas1 |
| F11r | 8,62E-11 | 0,494761613 | 0,944 | 0,822 | 1,37E-06 | T_CLEC | F11r |
| Tspan9 | 8,88E-11 | 0,52308318 | 0,744 | 0,387 | 1,41E-06 | T_CLEC | Tspan9 |
| Ctnna1 | 9,21E-11 | 0,573735056 | 0,919 | 0,687 | 1,46E-06 | T_CLEC | Ctnna1 |
| Cnn31 | 9,57E-11 | 0,59158725 | 0,944 | 0,534 | 1,52E-06 | T_CLEC | Cnn3 |
| Ctla2a1 | 1,17E-10 | 0,520003758 | 0,969 | 0,534 | 1,86E-06 | T_CLEC | Ctla2a |
| Calcrl | 1,25E-10 | 0,412099698 | 0,731 | 0,356 | 1,98E-06 | T_CLEC | Calcrl |
| Kcnj8 | 1,66E-10 | 1,028139413 | 0,35 | 0,067 | 2,63E-06 | T_CLEC | Kcnj8 |
| Piezo2 | 1,86E-10 | 0,60540364 | 0,688 | 0,362 | 2,94E-06 | T_CLEC | Piezo2 |
| Myct1 | 2,05E-10 | 0,592769189 | 0,65 | 0,319 | 3,25E-06 | T_CLEC | Myct1 |
| Shank3 | 2,09E-10 | 0,25947851 | 0,569 | 0,264 | 3,31E-06 | T_CLEC | Shank3 |
| Kcne3 | 2,15E-10 | 0,829505865 | 0,638 | 0,325 | 3,41E-06 | T_CLEC | Kcne3 |
| S100a61 | 2,77E-10 | 0,415508649 | 0,938 | 0,712 | 4,39E-06 | T_CLEC | S100a6 |
| Prkcdbp1 | 2,81E-10 | 0,455123312 | 0,95 | 0,503 | 4,45E-06 | T_CLEC | Prkcdbp |
| Tshz2 | 3,03E-10 | 0,519575791 | 0,612 | 0,282 | 4,81E-06 | T_CLEC | Tshz2 |
| Tie11 | 3,05E-10 | 0,455866077 | 0,844 | 0,436 | 4,83E-06 | T_CLEC | Tie1 |
| Emp1 | 3,61E-10 | 0,556113899 | 0,888 | 0,62 | 5,71E-06 | T_CLEC | Emp1 |
| Gnb1 | 3,72E-10 | 0,533255432 | 1 | 0,994 | 5,90E-06 | T_CLEC | Gnb1 |
| Ptrf1 | 5,62E-10 | 0,475468692 | 0,9 | 0,509 | 8,91E-06 | T_CLEC | Ptrf |
| Sptan1 | 5,83E-10 | 0,64124894 | 0,825 | 0,656 | 9,24E-06 | T_CLEC | Sptan1 |
| Cdc42bpa | 5,83E-10 | 0,55402809 | 0,512 | 0,202 | 9,25E-06 | T_CLEC | Cdc42bpa |
| Dock9 | 9,42E-10 | 0,421594028 | 0,781 | 0,411 | 1,49E-05 | T_CLEC | Dock9 |
| Akap2 | 9,84E-10 | 0,363527232 | 0,562 | 0,233 | 1,56E-05 | T_CLEC | Akap2 |
| Cd9 | 1,08E-09 | 0,529739087 | 0,925 | 0,804 | 1,72E-05 | T_CLEC | Cd9 |
| Rapgef5 | 1,11E-09 | 0,563746171 | 0,744 | 0,423 | 1,76E-05 | T_CLEC | Rapgef5 |
| Tes | 1,16E-09 | 0,410223125 | 0,875 | 0,571 | 1,83E-05 | T_CLEC | Tes |
| Slc9a3r21 | 1,36E-09 | 0,538724534 | 0,819 | 0,436 | 2,16E-05 | T_CLEC | Slc9a3r2 |
| Ehd2 | 1,44E-09 | 0,422677526 | 0,706 | 0,368 | 2,28E-05 | T_CLEC | Ehd2 |
| Inhbb | 1,66E-09 | 0,767264461 | 0,606 | 0,313 | 2,64E-05 | T_CLEC | Inhbb |
| Nav1 | 1,73E-09 | 0,521633644 | 0,612 | 0,307 | 2,74E-05 | T_CLEC | Nav1 |
| Ptprg | 1,80E-09 | 0,510299193 | 0,569 | 0,245 | 2,86E-05 | T_CLEC | Ptprg |
| Msn1 | 1,82E-09 | 0,385111029 | 0,975 | 0,957 | 2,89E-05 | T_CLEC | Msn |
| Wwtr1 | 1,88E-09 | 0,586644888 | 0,669 | 0,374 | 2,99E-05 | T_CLEC | Wwtr1 |
| Adamts4 | 1,94E-09 | 0,448218883 | 0,612 | 0,27 | 3,08E-05 | T_CLEC | Adamts4 |
| Vwf1 | 2,55E-09 | 0,50608674 | 0,881 | 0,503 | 4,04E-05 | T_CLEC | Vwf |
| Trp53i111 | 2,95E-09 | 0,372028125 | 0,894 | 0,448 | 4,67E-05 | T_CLEC | Trp53i11 |
| Nfia | 3,07E-09 | 0,39731878 | 0,862 | 0,601 | 4,86E-05 | T_CLEC | Nfia |
| Lars2 | 3,42E-09 | 0,332652188 | 1 | 1 | 5,42E-05 | T_CLEC | Lars2 |
| Creb3l2 | 3,42E-09 | 0,560611454 | 0,731 | 0,411 | 5,43E-05 | T_CLEC | Creb3l2 |
| Amotl1 | 3,55E-09 | 0,529617208 | 0,681 | 0,374 | 5,63E-05 | T_CLEC | Amotl1 |
| Mast4 | 3,57E-09 | 0,471999109 | 0,706 | 0,38 | 5,67E-05 | T_CLEC | Mast4 |
| Rgcc | 3,70E-09 | 0,656720591 | 0,625 | 0,313 | 5,87E-05 | T_CLEC | Rgcc |
| Cd341 | 4,02E-09 | 0,467712368 | 0,975 | 0,521 | 6,37E-05 | T_CLEC | Cd34 |
| Adam10 | 4,33E-09 | 0,53684086 | 0,95 | 0,896 | 6,86E-05 | T_CLEC | Adam10 |
| S100a161 | 4,63E-09 | 0,419885059 | 0,888 | 0,472 | 7,34E-05 | T_CLEC | S100a16 |
| Entpd1 | 5,06E-09 | 0,778588113 | 0,894 | 0,804 | 8,01E-05 | T_CLEC | Entpd1 |
| Hif1a | 5,38E-09 | 0,606392857 | 0,869 | 0,748 | 8,53E-05 | T_CLEC | Hif1a |
| Adgrl41 | 5,56E-09 | 0,370640001 | 0,931 | 0,491 | 8,82E-05 | T_CLEC | Adgrl4 |
| Lpp | 6,35E-09 | 0,488106425 | 0,781 | 0,546 | 0,000100717 | T_CLEC | Lpp |
| Sept11 | 6,67E-09 | 0,506017925 | 0,706 | 0,423 | 0,000105675 | T_CLEC | Sept11 |
| Limch1 | 6,92E-09 | 0,434889136 | 0,45 | 0,153 | 0,000109704 | T_CLEC | Limch1 |
| Nedd41 | 7,75E-09 | 0,436678927 | 0,9 | 0,54 | 0,000122917 | T_CLEC | Nedd4 |
| Sox71 | 8,72E-09 | 0,372202396 | 0,769 | 0,387 | 0,000138223 | T_CLEC | Sox7 |
| Prss231 | 1,03E-08 | 0,479028051 | 0,794 | 0,448 | 0,000164046 | T_CLEC | Prss23 |
| mt-Rnr2 | 1,26E-08 | 0,324476157 | 1 | 1 | 0,00020029 | T_CLEC | mt-Rnr2 |
| Nid21 | 1,30E-08 | 0,411046429 | 0,794 | 0,46 | 0,0002058 | T_CLEC | Nid2 |
| Prex2 | 1,38E-08 | 0,447512014 | 0,888 | 0,767 | 0,000218643 | T_CLEC | Prex2 |
| Dusp3 | 1,76E-08 | 0,431520966 | 0,944 | 0,736 | 0,000278843 | T_CLEC | Dusp3 |
| Myo10 | 1,78E-08 | 0,474682462 | 0,625 | 0,313 | 0,000282687 | T_CLEC | Myo10 |
| Plod1 | 2,19E-08 | 0,524606928 | 0,912 | 0,84 | 0,000347438 | T_CLEC | Plod1 |
| Tspan18 | 2,66E-08 | 0,395971183 | 0,725 | 0,393 | 0,000421922 | T_CLEC | Tspan18 |
| Pxdn | 2,87E-08 | 0,630546431 | 0,662 | 0,405 | 0,000454231 | T_CLEC | Pxdn |
| Ly6c11 | 3,09E-08 | 0,439665396 | 0,825 | 0,466 | 0,000490411 | T_CLEC | Ly6c1 |
| Pde2a | 3,53E-08 | 0,666786236 | 0,55 | 0,264 | 0,000560328 | T_CLEC | Pde2a |
| Egfl71 | 3,82E-08 | 0,381312833 | 0,994 | 0,521 | 0,000605749 | T_CLEC | Egfl7 |
| Tacr1 | 4,35E-08 | 0,485585682 | 0,362 | 0,11 | 0,000690148 | T_CLEC | Tacr1 |
| Ptprm | 4,37E-08 | 0,64089967 | 0,631 | 0,374 | 0,000693188 | T_CLEC | Ptprm |
| Aplp2 | 4,41E-08 | 0,480608179 | 0,931 | 0,724 | 0,000699754 | T_CLEC | Aplp2 |
| Dchs1 | 4,56E-08 | 0,494273353 | 0,538 | 0,27 | 0,000722452 | T_CLEC | Dchs1 |
| Sema6d1 | 4,87E-08 | 0,479445226 | 0,781 | 0,472 | 0,000771421 | T_CLEC | Sema6d |
| Nckap1 | 5,17E-08 | 0,455293514 | 0,6 | 0,301 | 0,000820011 | T_CLEC | Nckap1 |
| Apold11 | 5,21E-08 | 0,461193513 | 0,712 | 0,405 | 0,000826382 | T_CLEC | Apold1 |
| Tram2 | 5,38E-08 | 0,489531164 | 0,756 | 0,503 | 0,000852528 | T_CLEC | Tram2 |
| Tmem2041 | 5,48E-08 | 0,483931678 | 0,712 | 0,393 | 0,000868976 | T_CLEC | Tmem204 |
| Lmcd1 | 7,07E-08 | 0,416878844 | 0,369 | 0,11 | 0,001120249 | T_CLEC | Lmcd1 |
| Tnfrsf22 | 7,11E-08 | 0,311811701 | 0,569 | 0,313 | 0,00112757 | T_CLEC | Tnfrsf22 |
| Ipo11 | 7,74E-08 | 0,519142216 | 0,706 | 0,417 | 0,001226855 | T_CLEC | Ipo11 |
| Plpp31 | 8,24E-08 | 0,440212384 | 0,712 | 0,387 | 0,001306401 | T_CLEC | Plpp3 |
| Adamts11 | 9,00E-08 | 0,318124149 | 0,744 | 0,399 | 0,001426296 | T_CLEC | Adamts1 |
| Nrp2 | 9,09E-08 | 0,411429453 | 0,644 | 0,368 | 0,001441453 | T_CLEC | Nrp2 |
| Aplnr1 | 9,24E-08 | 0,52895075 | 0,725 | 0,405 | 0,001464347 | T_CLEC | Aplnr |
| Fermt21 | 1,00E-07 | 0,393614194 | 0,775 | 0,436 | 0,001585906 | T_CLEC | Fermt2 |
| Exoc3l2 | 1,05E-07 | 0,435076218 | 0,419 | 0,16 | 0,001664559 | T_CLEC | Exoc3l2 |
| Endod1 | 1,17E-07 | 0,636327292 | 0,675 | 0,448 | 0,001858669 | T_CLEC | Endod1 |
| Nes | 1,42E-07 | 0,406529788 | 0,494 | 0,227 | 0,002253319 | T_CLEC | Nes |
| Wscd1 | 1,46E-07 | 0,390682328 | 0,5 | 0,209 | 0,002307544 | T_CLEC | Wscd1 |
| Plxna2 | 1,65E-07 | 0,38691773 | 0,538 | 0,252 | 0,002608806 | T_CLEC | Plxna2 |
| Rhoc1 | 1,66E-07 | 0,323659819 | 0,9 | 0,564 | 0,00263729 | T_CLEC | Rhoc |
| Upp11 | 1,70E-07 | 0,377127429 | 0,8 | 0,46 | 0,002690943 | T_CLEC | Upp1 |
| Tuba1a1 | 1,84E-07 | 0,439805885 | 0,838 | 0,607 | 0,002914621 | T_CLEC | Tuba1a |
| Pls3 | 1,90E-07 | 0,555165381 | 0,731 | 0,448 | 0,003014822 | T_CLEC | Pls3 |
| Atp1b3 | 2,14E-07 | 0,414615259 | 0,95 | 0,883 | 0,003384723 | T_CLEC | Atp1b3 |
| Pcdh17 | 2,15E-07 | 0,323101218 | 0,544 | 0,27 | 0,003402902 | T_CLEC | Pcdh17 |
| Igfbp31 | 2,22E-07 | 0,254115632 | 0,988 | 0,577 | 0,003519978 | T_CLEC | Igfbp3 |
| Cd300lg | 2,23E-07 | 0,705789112 | 0,4 | 0,153 | 0,003532496 | T_CLEC | Cd300lg |
| Tmem2 | 2,38E-07 | 0,350574728 | 0,675 | 0,393 | 0,003768197 | T_CLEC | Tmem2 |
| Ext1 | 2,68E-07 | 0,487593642 | 0,606 | 0,362 | 0,004253865 | T_CLEC | Ext1 |
| Tjp1 | 2,75E-07 | 0,462888529 | 0,625 | 0,356 | 0,00435334 | T_CLEC | Tjp1 |
| Dbn11 | 3,15E-07 | 0,395763082 | 0,662 | 0,368 | 0,00499851 | T_CLEC | Dbn1 |
| Robo1 | 3,25E-07 | 0,322589725 | 0,331 | 0,104 | 0,005149752 | T_CLEC | Robo1 |
| Rasip11 | 3,29E-07 | 0,355251192 | 0,762 | 0,436 | 0,005220946 | T_CLEC | Rasip1 |
| Lama5 | 3,56E-07 | 0,605059017 | 0,55 | 0,282 | 0,005636044 | T_CLEC | Lama5 |
| Tmem881 | 3,57E-07 | 0,281579671 | 0,9 | 0,497 | 0,005664112 | T_CLEC | Tmem88 |
| Ackr3 | 3,87E-07 | 0,493080336 | 0,556 | 0,288 | 0,006129477 | T_CLEC | Ackr3 |
| Apbb2 | 3,91E-07 | 0,447588406 | 0,744 | 0,454 | 0,006202247 | T_CLEC | Apbb2 |
| Crebbp | 3,92E-07 | 0,38415043 | 0,738 | 0,509 | 0,006215271 | T_CLEC | Crebbp |
| Gm4202 | 3,96E-07 | 0,337360389 | 0,888 | 0,822 | 0,006282333 | T_CLEC | Gm4202 |
| Nudt4 | 4,29E-07 | 0,576522187 | 0,519 | 0,307 | 0,00680542 | T_CLEC | Nudt4 |
| Sorbs31 | 4,44E-07 | 0,368552575 | 0,819 | 0,558 | 0,007043186 | T_CLEC | Sorbs3 |
| Sept10 | 4,61E-07 | 0,2848015 | 0,794 | 0,595 | 0,007307698 | T_CLEC | Sept10 |
| Elk31 | 4,73E-07 | 0,398070525 | 0,912 | 0,73 | 0,007496624 | T_CLEC | Elk3 |
| Rell1 | 5,15E-07 | 0,374930016 | 0,644 | 0,362 | 0,008166109 | T_CLEC | Rell1 |
| Bmpr2 | 5,17E-07 | 0,455420354 | 0,731 | 0,46 | 0,008194341 | T_CLEC | Bmpr2 |
| Ptpn14 | 5,63E-07 | 0,353107047 | 0,394 | 0,147 | 0,008931849 | T_CLEC | Ptpn14 |
| Dysf | 6,04E-07 | 0,279951075 | 0,544 | 0,245 | 0,009570432 | T_CLEC | Dysf |
| Arhgef12 | 6,30E-07 | 0,343851622 | 0,575 | 0,313 | 0,009988153 | T_CLEC | Arhgef12 |
| Tm4sf11 | 6,85E-07 | 0,273426204 | 0,938 | 0,521 | 0,010857072 | T_CLEC | Tm4sf1 |
| Emid1 | 7,29E-07 | 0,350637623 | 0,456 | 0,19 | 0,01156158 | T_CLEC | Emid1 |
| Zcchc14 | 8,34E-07 | 0,370947528 | 0,488 | 0,221 | 0,01321559 | T_CLEC | Zcchc14 |

| | | | | | | | |
|---|---|---|---|---|---|---|---|
| Ldlrad4 | 2,67E-05 | 0,325079327 | 0,27 | 0,085 | 0,423861651 | 2 | Ldlrad4 |
| Arhgap29 | 2,74E-05 | 0,303959697 | 0,825 | 0,492 | 0,434061364 | 2 | Arhgap29 |
| Myo10 | 2,74E-05 | 0,295507415 | 0,698 | 0,412 | 0,434261934 | 2 | Myo10 |
| Ubqln1 | 2,82E-05 | 0,359294559 | 0,651 | 0,373 | 0,447369862 | 2 | Ubqln1 |
| Plcg1 | 2,84E-05 | 0,272970098 | 0,746 | 0,427 | 0,450461884 | 2 | Plcg1 |
| Pttg1ip | 2,85E-05 | 0,329400411 | 0,841 | 0,6 | 0,451346328 | 2 | Pttg1ip |
| Bace2 | 2,91E-05 | 0,285489038 | 0,492 | 0,223 | 0,461845009 | 2 | Bace2 |
| Rnf14 | 2,92E-05 | 0,307000502 | 0,587 | 0,3 | 0,463461248 | 2 | Rnf14 |
| Ppp5c | 3,07E-05 | 0,284879757 | 0,54 | 0,285 | 0,487032549 | 2 | Ppp5c |
| Eif4g2 | 3,10E-05 | 0,280701533 | 1 | 0,881 | 0,490671772 | 2 | Eif4g2 |
| Zfand5 | 3,18E-05 | 0,331005636 | 0,873 | 0,635 | 0,503774664 | 2 | Zfand5 |
| Eva1a | 3,37E-05 | 0,312475751 | 0,365 | 0,135 | 0,533575688 | 2 | Eva1a |
| Gja11 | 3,46E-05 | 0,254429688 | 0,921 | 0,596 | 0,547950488 | 2 | Gja1 |
| Mapk11 | 3,64E-05 | 0,313322283 | 0,413 | 0,173 | 0,576521451 | 2 | Mapk11 |
| Tmem51 | 3,65E-05 | 0,298664687 | 0,381 | 0,15 | 0,579214568 | 2 | Tmem51 |
| Trim16 | 3,98E-05 | 0,409703461 | 0,444 | 0,2 | 0,630706461 | 2 | Trim16 |
| Litaf | 4,04E-05 | 0,342179417 | 0,984 | 0,877 | 0,640829488 | 2 | Litaf |
| Jup1 | 4,10E-05 | 0,288233819 | 0,952 | 0,681 | 0,649602537 | 2 | Jup |
| Timp31 | 4,13E-05 | 0,388462592 | 0,952 | 0,577 | 0,655357399 | 2 | Timp3 |
| Lims1 | 4,23E-05 | 0,39515137 | 0,841 | 0,615 | 0,671278127 | 2 | Lims1 |
| Hsp90ab1 | 4,27E-05 | 0,25546771 | 1 | 0,985 | 0,676996723 | 2 | Hsp90ab1 |
| Cd109 | 4,33E-05 | 0,35593718 | 0,667 | 0,381 | 0,687175401 | 2 | Cd109 |
| Sar1a | 4,59E-05 | 0,372640327 | 0,746 | 0,542 | 0,72786196 | 2 | Sar1a |
| Rnf122 | 4,79E-05 | 0,259157579 | 0,476 | 0,235 | 0,759915703 | 2 | Rnf122 |
| Akap12 | 4,91E-05 | 0,313113454 | 0,587 | 0,308 | 0,778239627 | 2 | Akap12 |
| Anxa21 | 5,32E-05 | 0,348460659 | 1 | 0,838 | 0,843254514 | 2 | Anxa2 |
| Arrdc3 | 5,70E-05 | 0,409646776 | 0,571 | 0,308 | 0,903953053 | 2 | Arrdc3 |
| Tmed4 | 5,78E-05 | 0,260018788 | 0,651 | 0,369 | 0,917019517 | 2 | Tmed4 |
| H2-Q6 | 5,82E-05 | 0,433271083 | 0,683 | 0,392 | 0,922324191 | 2 | H2-Q6 |
| Rpl15-ps3 | 5,91E-05 | 0,262594626 | 0,984 | 0,896 | 0,937609819 | 2 | Rpl15-ps3 |
| Flnb | 5,93E-05 | 0,316108978 | 0,794 | 0,485 | 0,939972835 | 2 | Flnb |
| Tax1bp3 | 6,02E-05 | 0,280461639 | 0,921 | 0,7 | 0,953510661 | 2 | Tax1bp3 |
| Skp1a | 6,18E-05 | 0,325870733 | 0,952 | 0,796 | 0,979866696 | 2 | Skp1a |
| Gorasp2 | 6,42E-05 | 0,395895664 | 0,857 | 0,604 | 1 | 2 | Gorasp2 |
| Psmd12 | 6,53E-05 | 0,294246532 | 0,778 | 0,535 | 1 | 2 | Psmd12 |
| Oasl2 | 7,34E-05 | 0,30173647 | 0,397 | 0,162 | 1 | 2 | Oasl2 |
| Sec61a1 | 7,69E-05 | 0,356034579 | 0,905 | 0,738 | 1 | 2 | Sec61a1 |
| Adamts4 | 7,87E-05 | 0,449881274 | 0,635 | 0,392 | 1 | 2 | Adamts4 |
| Igfbp71 | 7,91E-05 | 0,289923498 | 1 | 0,712 | 1 | 2 | Igfbp7 |
| Tie11 | 8,13E-05 | 0,350239306 | 0,873 | 0,581 | 1 | 2 | Tie1 |
| Fkbp10 | 8,14E-05 | 0,275998325 | 0,746 | 0,427 | 1 | 2 | Fkbp10 |
| Plscr3 | 8,24E-05 | 0,31477345 | 0,778 | 0,546 | 1 | 2 | Plscr3 |
| Scfd1 | 8,49E-05 | 0,264762905 | 0,556 | 0,292 | 1 | 2 | Scfd1 |
| Rab11b | 8,62E-05 | 0,316850029 | 0,937 | 0,765 | 1 | 2 | Rab11b |
| Igfbp31 | 8,66E-05 | 0,308324232 | 1 | 0,727 | 1 | 2 | Igfbp3 |
| Ipo11 | 8,87E-05 | 0,340098742 | 0,778 | 0,508 | 1 | 2 | Ipo11 |
| Cnn2 | 9,23E-05 | 0,392742393 | 0,794 | 0,596 | 1 | 2 | Cnn2 |
| Mapk12 | 9,34E-05 | 0,267734061 | 0,381 | 0,162 | 1 | 2 | Mapk12 |
| Eif4ebp1 | 9,95E-05 | 0,250442941 | 0,905 | 0,638 | 1 | 2 | Eif4ebp1 |
| Puf60 | 1,00E-04 | 0,282214734 | 0,81 | 0,538 | 1 | 2 | Puf60 |
| Cd931 | 0,000105976 | 0,297686894 | 1 | 0,912 | 1 | 2 | Cd93 |
| Rce1 | 0,000108524 | 0,296598193 | 0,46 | 0,231 | 1 | 2 | Rce1 |
| Prelid1 | 0,000109858 | 0,294233473 | 0,952 | 0,877 | 1 | 2 | Prelid1 |
| Mapre1 | 0,000118292 | 0,286383819 | 0,873 | 0,708 | 1 | 2 | Mapre1 |
| Tpi11 | 0,000126117 | 0,400472802 | 0,921 | 0,727 | 1 | 2 | Tpi1 |
| Itga3 | 0,000137188 | 0,408685328 | 0,556 | 0,338 | 1 | 2 | Itga3 |
| Slc1a5 | 0,000137808 | 0,3946415 | 0,667 | 0,423 | 1 | 2 | Slc1a5 |
| Icam21 | 0,000138475 | 0,307224233 | 0,889 | 0,588 | 1 | 2 | Icam2 |
| Ctnnb1 | 0,000150393 | 0,314007873 | 0,968 | 0,796 | 1 | 2 | Ctnnb1 |
| Cnbp | 0,000150498 | 0,335774234 | 1 | 0,881 | 1 | 2 | Cnbp |
| Phb | 0,000151769 | 0,261295331 | 0,794 | 0,519 | 1 | 2 | Phb |
| Slc44a1 | 0,000152894 | 0,32236558 | 0,667 | 0,415 | 1 | 2 | Slc44a1 |
| Rars | 0,000156118 | 0,266989582 | 0,556 | 0,296 | 1 | 2 | Rars |
| Pdlim5 | 0,000158187 | 0,314667671 | 0,651 | 0,427 | 1 | 2 | Pdlim5 |
| Dpysl3 | 0,000164816 | 0,290976172 | 0,54 | 0,288 | 1 | 2 | Dpysl3 |
| Raph1 | 0,000170537 | 0,271033359 | 0,667 | 0,408 | 1 | 2 | Raph1 |
| Ero1l | 0,000170921 | 0,394840425 | 0,556 | 0,312 | 1 | 2 | Ero1l |
| Csde1 | 0,000179158 | 0,33381198 | 0,921 | 0,769 | 1 | 2 | Csde1 |
| Tubb2a | 0,000181953 | 0,300269071 | 0,683 | 0,423 | 1 | 2 | Tubb2a |
| Map4k4 | 0,000182022 | 0,305539449 | 0,762 | 0,527 | 1 | 2 | Map4k4 |
| Egfl71 | 0,000183399 | 0,312124328 | 1 | 0,696 | 1 | 2 | Egfl7 |
| Ralgds | 0,00018586 | 0,39621313 | 0,54 | 0,315 | 1 | 2 | Ralgds |
| Tmem43 | 0,000186193 | 0,327904174 | 0,524 | 0,281 | 1 | 2 | Tmem43 |
| Nck11 | 0,000192073 | 0,253591456 | 0,587 | 0,323 | 1 | 2 | Nck1 |
| Pecam11 | 0,000192161 | 0,303133634 | 0,984 | 0,719 | 1 | 2 | Pecam1 |
| Dusp31 | 0,000203101 | 0,327587663 | 0,937 | 0,815 | 1 | 2 | Dusp3 |
| Lrp10 | 0,000204669 | 0,262320713 | 0,81 | 0,6 | 1 | 2 | Lrp10 |
| Ints11 | 0,000211207 | 0,266814136 | 0,444 | 0,215 | 1 | 2 | Ints11 |
| Jag1 | 0,000214525 | 0,265368524 | 0,683 | 0,419 | 1 | 2 | Jag1 |
| Prss231 | 0,000215859 | 0,39314849 | 0,857 | 0,562 | 1 | 2 | Prss23 |
| Slc25a25 | 0,000217255 | 0,254339051 | 0,286 | 0,108 | 1 | 2 | Slc25a25 |
| Ywhaq | 0,000228035 | 0,272150685 | 0,952 | 0,812 | 1 | 2 | Ywhaq |
| Mrpl3 | 0,000228551 | 0,332458319 | 0,444 | 0,227 | 1 | 2 | Mrpl3 |
| Itpa | 0,000231721 | 0,266992117 | 0,54 | 0,304 | 1 | 2 | Itpa |
| Tspan6 | 0,000233759 | 0,283448476 | 0,54 | 0,292 | 1 | 2 | Tspan6 |
| Ldha | 0,000239298 | 0,356746348 | 0,968 | 0,919 | 1 | 2 | Ldha |
| Angptl4 | 0,000242757 | 0,407751725 | 0,397 | 0,177 | 1 | 2 | Angptl4 |
| Ppp1r7 | 0,000245433 | 0,273874657 | 0,492 | 0,273 | 1 | 2 | Ppp1r7 |
| Pigx | 0,000246937 | 0,294990697 | 0,476 | 0,265 | 1 | 2 | Pigx |
| Pfkl | 0,000248767 | 0,40833334 | 0,667 | 0,462 | 1 | 2 | Pfkl |
| Fam149a | 0,000250533 | 0,37710233 | 0,444 | 0,231 | 1 | 2 | Fam149a |
| Sec23ip | 0,000269772 | 0,355909118 | 0,46 | 0,25 | 1 | 2 | Sec23ip |
| Raet1e | 0,000285256 | 0,264759375 | 0,413 | 0,204 | 1 | 2 | Raet1e |
| Slco2b1 | 0,000287068 | 0,331180301 | 0,635 | 0,408 | 1 | 2 | Slco2b1 |
| Tmem184b | 0,000288858 | 0,278736499 | 0,746 | 0,508 | 1 | 2 | Tmem184b |
| Mxra81 | 0,000302407 | 0,297777116 | 0,635 | 0,365 | 1 | 2 | Mxra8 |
| Dcbld1 | 0,000332522 | 0,263490722 | 0,508 | 0,269 | 1 | 2 | Dcbld1 |
| Anxa5 | 0,000332585 | 0,309311708 | 0,984 | 0,827 | 1 | 2 | Anxa5 |
| Saraf | 0,000339971 | 0,291378792 | 0,778 | 0,558 | 1 | 2 | Saraf |
| Gdpd5 | 0,000342931 | 0,300236509 | 0,349 | 0,158 | 1 | 2 | Gdpd5 |
| Ssr2 | 0,000348702 | 0,258707594 | 0,841 | 0,592 | 1 | 2 | Ssr2 |
| Pcdhgc4 | 0,000356297 | 0,265208204 | 0,635 | 0,419 | 1 | 2 | Pcdhgc4 |
| Nfkb2 | 0,000360127 | 0,258669954 | 0,381 | 0,185 | 1 | 2 | Nfkb2 |
| Irgm1 | 0,000373773 | 0,545149084 | 0,556 | 0,331 | 1 | 2 | Irgm1 |
| Slc38a2 | 0,000374384 | 0,316459147 | 0,825 | 0,612 | 1 | 2 | Slc38a2 |
| Apln | 0,00037566 | 0,387801029 | 0,587 | 0,331 | 1 | 2 | Apln |
| Sqstm1 | 0,000382441 | 0,315471653 | 0,952 | 0,735 | 1 | 2 | Sqstm1 |
| Clec14a1 | 0,000388297 | 0,311000109 | 0,81 | 0,492 | 1 | 2 | Clec14a |
| Sh3gl1 | 0,000395907 | 0,308168945 | 0,651 | 0,404 | 1 | 2 | Sh3gl1 |
| Gata2 | 0,00039599 | 0,379569464 | 0,429 | 0,227 | 1 | 2 | Gata2 |
| Strn4 | 0,000410951 | 0,289244948 | 0,603 | 0,396 | 1 | 2 | Strn4 |
| Wdr1 | 0,00042272 | 0,291305155 | 0,952 | 0,892 | 1 | 2 | Wdr1 |
| Ldlrap1 | 0,000428537 | 0,253736836 | 0,524 | 0,308 | 1 | 2 | Ldlrap1 |
| Rnf185 | 0,000432565 | 0,258139365 | 0,365 | 0,169 | 1 | 2 | Rnf185 |
| Nrbp1 | 0,000439978 | 0,308060261 | 0,762 | 0,558 | 1 | 2 | Nrbp1 |
| Nap1l4 | 0,000443562 | 0,284599041 | 0,762 | 0,581 | 1 | 2 | Nap1l4 |
| Aaed1 | 0,000444134 | 0,293386425 | 0,603 | 0,381 | 1 | 2 | Aaed1 |
| Adamts11 | 0,00044758 | 0,406854835 | 0,794 | 0,515 | 1 | 2 | Adamts1 |
| Crip21 | 0,000454608 | 0,296642658 | 0,984 | 0,665 | 1 | 2 | Crip2 |
| Il2rg | 0,000467553 | 0,391777061 | 0,825 | 0,665 | 1 | 2 | Il2rg |
| Tomm70a | 0,000482529 | 0,408721516 | 0,524 | 0,323 | 1 | 2 | Tomm70a |

| | | | | | | | |
|---|---|---|---|---|---|---|---|
| Arhgap291 | 8,43E-07 | 0,322314858 | 0,706 | 0,411 | 0,013364444 | T_CLEC | Arhgap29 |
| Raph1 | 8,46E-07 | 0,353535721 | 0,588 | 0,331 | 0,01341054 | T_CLEC | Raph1 |
| Mfge8 | 9,16E-07 | 0,484461709 | 0,7 | 0,491 | 0,014518521 | T_CLEC | Mfge8 |
| Fzd41 | 9,63E-07 | 0,290232661 | 0,656 | 0,344 | 0,015259416 | T_CLEC | Fzd4 |
| Tspan7 | 9,73E-07 | 0,537500582 | 0,456 | 0,215 | 0,015428625 | T_CLEC | Tspan7 |
| Flnb1 | 9,94E-07 | 0,392830408 | 0,694 | 0,399 | 0,015754972 | T_CLEC | Flnb |
| 9430020K01Rik1 | 1,04E-06 | 0,468932086 | 0,725 | 0,46 | 0,016535451 | T_CLEC | 9430020K01Rik |
| Plk21 | 1,09E-06 | 0,398675858 | 0,881 | 0,601 | 0,017320809 | T_CLEC | Plk2 |
| Calm1 | 1,11E-06 | 0,297401531 | 0,994 | 0,994 | 0,017544249 | T_CLEC | Calm1 |
| Adamts7 | 1,21E-06 | 0,397200534 | 0,312 | 0,098 | 0,01917214 | T_CLEC | Adamts7 |
| Icam21 | 1,31E-06 | 0,387508757 | 0,812 | 0,485 | 0,020794087 | T_CLEC | Icam2 |
| Notch1 | 1,35E-06 | 0,554049696 | 0,744 | 0,613 | 0,021360814 | T_CLEC | Notch1 |
| Ctnnb11 | 1,35E-06 | 0,449752016 | 0,888 | 0,773 | 0,02145486 | T_CLEC | Ctnnb1 |
| Igfbp71 | 1,41E-06 | 0,319106097 | 1 | 0,54 | 0,022319507 | T_CLEC | Igfbp7 |
| Gm10800 | 1,47E-06 | 0,292839568 | 0,825 | 0,742 | 0,023244586 | T_CLEC | Gm10800 |
| Chst15 | 1,55E-06 | 0,363228046 | 0,394 | 0,153 | 0,02457498 | T_CLEC | Chst15 |
| Erg | 1,56E-06 | 0,269671205 | 0,481 | 0,239 | 0,024788101 | T_CLEC | Erg |
| Asap1 | 1,64E-06 | 0,416355304 | 0,594 | 0,325 | 0,026019815 | T_CLEC | Asap1 |
| Map4k4 | 1,64E-06 | 0,384637389 | 0,694 | 0,454 | 0,026045483 | T_CLEC | Map4k4 |
| Ramp21 | 1,78E-06 | 0,279894024 | 0,906 | 0,503 | 0,028140588 | T_CLEC | Ramp2 |
| Gimap41 | 1,78E-06 | 0,39508009 | 0,769 | 0,46 | 0,028237952 | T_CLEC | Gimap4 |
| Cd276 | 1,80E-06 | 0,432632487 | 0,475 | 0,239 | 0,028474311 | T_CLEC | Cd276 |
| Cgnl1 | 1,80E-06 | 0,373096176 | 0,438 | 0,19 | 0,028549819 | T_CLEC | Cgnl1 |
| 2900026A02Rik | 1,87E-06 | 0,414067982 | 0,6 | 0,356 | 0,029669416 | T_CLEC | 2900026A02Rik |
| Cdr2l | 1,97E-06 | 0,452572483 | 0,569 | 0,325 | 0,031305566 | T_CLEC | Cdr2l |
| Sept7 | 2,04E-06 | 0,367320999 | 0,881 | 0,718 | 0,032275244 | T_CLEC | Sept7 |
| Nr2f2 | 2,13E-06 | 0,332175729 | 0,5 | 0,258 | 0,033834534 | T_CLEC | Nr2f2 |
| Apln | 2,17E-06 | 0,55548212 | 0,506 | 0,258 | 0,034446058 | T_CLEC | Apln |
| Oaf | 2,53E-06 | 0,431973659 | 0,556 | 0,313 | 0,040154223 | T_CLEC | Oaf |
| Shroom2 | 2,60E-06 | 0,289953807 | 0,612 | 0,344 | 0,041268517 | T_CLEC | Shroom2 |
| Thbd | 2,73E-06 | 0,407947998 | 0,488 | 0,233 | 0,043211033 | T_CLEC | Thbd |
| Ralb | 2,80E-06 | 0,432192449 | 0,894 | 0,785 | 0,044424464 | T_CLEC | Ralb |
| Caskin2 | 3,03E-06 | 0,36127263 | 0,556 | 0,325 | 0,048005108 | T_CLEC | Caskin2 |
| Yes1 | 3,18E-06 | 0,252326328 | 0,538 | 0,27 | 0,050389403 | T_CLEC | Yes1 |
| Rai14 | 3,31E-06 | 0,445859702 | 0,644 | 0,411 | 0,052405136 | T_CLEC | Rai14 |
| Ltbp4 | 3,67E-06 | 0,448630604 | 0,481 | 0,239 | 0,058140697 | T_CLEC | Ltbp4 |
| Dpysl2 | 3,80E-06 | 0,587771162 | 0,85 | 0,791 | 0,060168713 | T_CLEC | Dpysl2 |
| Dpysl3 | 3,81E-06 | 0,296128482 | 0,456 | 0,221 | 0,060329772 | T_CLEC | Dpysl3 |
| Snrk1 | 4,18E-06 | 0,269390272 | 0,831 | 0,546 | 0,066235162 | T_CLEC | Snrk |
| Ephb4 | 4,42E-06 | 0,305961374 | 0,594 | 0,307 | 0,070094723 | T_CLEC | Ephb4 |
| Gnai2 | 4,59E-06 | 0,29156889 | 0,95 | 0,988 | 0,072754636 | T_CLEC | Gnai2 |
| Maged1 | 4,66E-06 | 0,563422566 | 0,631 | 0,399 | 0,073874162 | T_CLEC | Maged1 |
| Ccser2 | 4,71E-06 | 0,386287456 | 0,55 | 0,307 | 0,074622841 | T_CLEC | Ccser2 |
| Kif5b | 4,94E-06 | 0,484361897 | 0,812 | 0,712 | 0,078347341 | T_CLEC | Kif5b |
| Pitpnm2 | 5,22E-06 | 0,297188444 | 0,425 | 0,202 | 0,082809562 | T_CLEC | Pitpnm2 |
| Adamts9 | 5,27E-06 | 0,378531904 | 0,606 | 0,337 | 0,083471699 | T_CLEC | Adamts9 |
| Cav11 | 5,38E-06 | 0,28562025 | 0,7 | 0,38 | 0,085225311 | T_CLEC | Cav1 |
| Mprip1 | 5,48E-06 | 0,364892068 | 0,744 | 0,503 | 0,086945939 | T_CLEC | Mprip |
| H1f0 | 5,52E-06 | 0,541239849 | 0,831 | 0,712 | 0,087533416 | T_CLEC | H1f0 |
| Adgrg3 | 5,95E-06 | 0,48918233 | 0,319 | 0,117 | 0,094368219 | T_CLEC | Adgrg3 |
| Gsk3b | 6,00E-06 | 0,39794201 | 0,85 | 0,773 | 0,095098536 | T_CLEC | Gsk3b |
| Snn | 6,37E-06 | 0,282652417 | 0,512 | 0,294 | 0,100934515 | T_CLEC | Snn |
| Cdc42ep1 | 6,37E-06 | 0,474478453 | 0,575 | 0,344 | 0,100936066 | T_CLEC | Cdc42ep1 |
| Schip1 | 6,38E-06 | 0,393211964 | 0,681 | 0,448 | 0,101096861 | T_CLEC | Schip1 |
| Cav21 | 6,45E-06 | 0,382057044 | 0,694 | 0,442 | 0,102273907 | T_CLEC | Cav2 |
| Fyn | 6,51E-06 | 0,295902753 | 0,6 | 0,325 | 0,103179885 | T_CLEC | Fyn |
| Tek1 | 6,52E-06 | 0,447967541 | 0,644 | 0,393 | 0,103373296 | T_CLEC | Tek |
| Pde4d | 6,63E-06 | 0,27645824 | 0,388 | 0,16 | 0,105087899 | T_CLEC | Pde4d |
| B3gnt31 | 6,80E-06 | 0,375189007 | 0,694 | 0,436 | 0,10783154 | T_CLEC | B3gnt3 |
| Npnt | 6,93E-06 | 0,397939618 | 0,256 | 0,074 | 0,109861533 | T_CLEC | Npnt |
| Unc5b | 7,18E-06 | 0,274147362 | 0,35 | 0,135 | 0,113803748 | T_CLEC | Unc5b |
| Lrrc58 | 7,26E-06 | 0,402139 | 0,925 | 0,926 | 0,115120819 | T_CLEC | Lrrc58 |
| Vim1 | 7,42E-06 | 0,286958509 | 1 | 0,957 | 0,117559721 | T_CLEC | Vim |
| Map1b | 7,62E-06 | 0,629684416 | 0,562 | 0,35 | 0,120771978 | T_CLEC | Map1b |
| Robo41 | 7,65E-06 | 0,377454752 | 0,738 | 0,448 | 0,121306366 | T_CLEC | Robo4 |
| Mkl2 | 8,94E-06 | 0,517858393 | 0,7 | 0,491 | 0,141686502 | T_CLEC | Mkl2 |
| Arhgef5 | 9,18E-06 | 0,277292166 | 0,338 | 0,135 | 0,145499543 | T_CLEC | Arhgef5 |
| Atp9a | 9,34E-06 | 0,318112685 | 0,4 | 0,166 | 0,148130551 | T_CLEC | Atp9a |
| Pcdh1 | 9,98E-06 | 0,27460531 | 0,569 | 0,319 | 0,15820676 | T_CLEC | Pcdh1 |
| Cyb5a | 1,04E-05 | 0,388940292 | 0,831 | 0,699 | 0,164262652 | T_CLEC | Cyb5a |
| Calu1 | 1,05E-05 | 0,376528737 | 0,838 | 0,675 | 0,166564507 | T_CLEC | Calu |
| Klk8 | 1,06E-05 | 0,330994938 | 0,312 | 0,117 | 0,168811384 | T_CLEC | Klk8 |
| Arhgap31 | 1,23E-05 | 0,484390556 | 0,7 | 0,515 | 0,195143003 | T_CLEC | Arhgap31 |
| Spns2 | 1,32E-05 | 0,357718842 | 0,388 | 0,178 | 0,209301082 | T_CLEC | Spns2 |
| Plat | 1,43E-05 | 0,540920463 | 0,519 | 0,301 | 0,225980978 | T_CLEC | Plat |
| Synpo | 1,47E-05 | 0,443768521 | 0,625 | 0,393 | 0,232856329 | T_CLEC | Synpo |
| Adarb1 | 1,50E-05 | 0,273835448 | 0,406 | 0,19 | 0,237031023 | T_CLEC | Adarb1 |
| Dock6 | 1,52E-05 | 0,453504286 | 0,525 | 0,307 | 0,241712547 | T_CLEC | Dock6 |
| Fkbp1a1 | 1,54E-05 | 0,304168938 | 0,981 | 0,877 | 0,244602687 | T_CLEC | Fkbp1a |
| Slc12a2 | 1,66E-05 | 0,38788938 | 0,388 | 0,184 | 0,263666786 | T_CLEC | Slc12a2 |
| Ushbp11 | 1,68E-05 | 0,300426686 | 0,712 | 0,393 | 0,265802811 | T_CLEC | Ushbp1 |
| Eml1 | 1,73E-05 | 0,422985318 | 0,562 | 0,337 | 0,274054438 | T_CLEC | Eml1 |
| Afap1l11 | 1,74E-05 | 0,34078627 | 0,65 | 0,374 | 0,27555647 | T_CLEC | Afap1l1 |
| Gm20721 | 1,82E-05 | 0,54745896 | 0,669 | 0,528 | 0,288943511 | T_CLEC | Gm20721 |
| Notch4 | 1,90E-05 | 0,398394947 | 0,581 | 0,35 | 0,300810505 | T_CLEC | Notch4 |
| Ddah1 | 1,91E-05 | 0,359916768 | 0,425 | 0,209 | 0,303042903 | T_CLEC | Ddah1 |
| Palm | 1,92E-05 | 0,33056359 | 0,494 | 0,258 | 0,304683637 | T_CLEC | Palm |
| Arap3 | 1,94E-05 | 0,270792825 | 0,625 | 0,38 | 0,307119083 | T_CLEC | Arap3 |
| Smtn1 | 2,04E-05 | 0,356602198 | 0,712 | 0,46 | 0,322748663 | T_CLEC | Smtn |
| Dll41 | 2,16E-05 | 0,397273437 | 0,594 | 0,356 | 0,343086463 | T_CLEC | Dll4 |
| Atrx | 2,17E-05 | 0,475844661 | 0,706 | 0,552 | 0,343937093 | T_CLEC | Atrx |
| Card10 | 2,32E-05 | 0,37321236 | 0,35 | 0,141 | 0,367405655 | T_CLEC | Card10 |
| Akap12 | 2,38E-05 | 0,411040744 | 0,469 | 0,258 | 0,376591011 | T_CLEC | Akap12 |
| Prkar1a1 | 2,41E-05 | 0,30579563 | 0,938 | 0,859 | 0,382774675 | T_CLEC | Prkar1a |
| Mn1 | 2,43E-05 | 0,283743154 | 0,275 | 0,092 | 0,385750079 | T_CLEC | Mn1 |
| Myo1b1 | 2,45E-05 | 0,336252049 | 0,7 | 0,436 | 0,388229717 | T_CLEC | Myo1b |
| Angpt2 | 2,46E-05 | 0,684426824 | 0,375 | 0,166 | 0,39023607 | T_CLEC | Angpt2 |
| Degs1 | 2,51E-05 | 0,457088662 | 0,85 | 0,73 | 0,397945737 | T_CLEC | Degs1 |
| Stc1 | 2,52E-05 | 0,548975058 | 0,481 | 0,27 | 0,398712712 | T_CLEC | Stc1 |
| Map4 | 2,64E-05 | 0,395101722 | 0,688 | 0,564 | 0,418588519 | T_CLEC | Map4 |
| Tsc22d11 | 2,64E-05 | 0,279643106 | 0,725 | 0,466 | 0,419245587 | T_CLEC | Tsc22d1 |
| Uaca | 2,77E-05 | 0,38068016 | 0,575 | 0,337 | 0,438561093 | T_CLEC | Uaca |
| Fam167b1 | 2,78E-05 | 0,303037354 | 0,681 | 0,429 | 0,441286441 | T_CLEC | Fam167b |
| Mpzl11 | 3,07E-05 | 0,367327427 | 0,731 | 0,448 | 0,486453928 | T_CLEC | Mpzl1 |
| Fam65a | 3,32E-05 | 0,422203985 | 0,594 | 0,423 | 0,526028531 | T_CLEC | Fam65a |
| Fgd5 | 3,37E-05 | 0,290994808 | 0,431 | 0,227 | 0,534976884 | T_CLEC | Fgd5 |
| Nus1 | 3,42E-05 | 0,331581029 | 0,65 | 0,436 | 0,542271371 | T_CLEC | Nus1 |
| Id31 | 3,70E-05 | 0,343105616 | 0,675 | 0,448 | 0,58656728 | T_CLEC | Id3 |
| Myadm1 | 3,73E-05 | 0,31658463 | 0,862 | 0,675 | 0,591468373 | T_CLEC | Myadm |
| Svil | 3,80E-05 | 0,319210406 | 0,575 | 0,399 | 0,602495538 | T_CLEC | Svil |
| Igfbp5 | 3,80E-05 | 0,630153377 | 0,438 | 0,233 | 0,602574787 | T_CLEC | Igfbp5 |
| Dock1 | 3,96E-05 | 0,395184558 | 0,6 | 0,436 | 0,628247021 | T_CLEC | Dock1 |
| Ptn | 4,14E-05 | 0,711101555 | 0,294 | 0,11 | 0,656200831 | T_CLEC | Ptn |
| Pcdh12 | 4,15E-05 | 0,358425113 | 0,362 | 0,172 | 0,657072681 | T_CLEC | Pcdh12 |
| Rasgrp3 | 4,35E-05 | 0,353877829 | 0,519 | 0,294 | 0,689551174 | T_CLEC | Rasgrp3 |
| Erc1 | 4,44E-05 | 0,375883732 | 0,369 | 0,172 | 0,704024949 | T_CLEC | Erc1 |
| Ctnnd1 | 4,57E-05 | 0,341421011 | 0,644 | 0,423 | 0,724001593 | T_CLEC | Ctnnd1 |
| Vamp5 | 4,82E-05 | 0,316526275 | 0,612 | 0,417 | 0,764657649 | T_CLEC | Vamp5 |
| Cd1091 | 5,08E-05 | 0,301602953 | 0,556 | 0,319 | 0,805402779 | T_CLEC | Cd109 |
| Pdzd8 | 5,48E-05 | 0,34348895 | 0,481 | 0,294 | 0,867972173 | T_CLEC | Pdzd8 |
| Ptp4a2 | 5,53E-05 | 0,440453467 | 0,9 | 0,896 | 0,877212856 | T_CLEC | Ptp4a2 |

| | | | | | | | |
|---|---|---|---|---|---|---|---|
| Ica1 | 0,000501236 | 0,312823794 | 0,571 | 0,365 | 1 | 2 | Ica1 |
| Rlim | 0,000512622 | 0,253499994 | 0,667 | 0,431 | 1 | 2 | Rlim |
| Dpp3 | 0,000540095 | 0,343477923 | 0,429 | 0,238 | 1 | 2 | Dpp3 |
| Ergic3 | 0,000548746 | 0,318855904 | 0,73 | 0,554 | 1 | 2 | Ergic3 |
| Arcn1 | 0,000558171 | 0,352191232 | 0,746 | 0,6 | 1 | 2 | Arcn1 |
| Rps6ka2 | 0,000567751 | 0,253263655 | 0,444 | 0,238 | 1 | 2 | Rps6ka2 |
| Nrros | 0,000576855 | 0,423874023 | 0,778 | 0,577 | 1 | 2 | Nrros |
| Ppp1r13b | 0,000585686 | 0,265474825 | 0,619 | 0,381 | 1 | 2 | Ppp1r13b |
| Bid | 0,000602011 | 0,300284287 | 0,286 | 0,119 | 1 | 2 | Bid |
| Eif2s1 | 0,000638563 | 0,251532026 | 0,762 | 0,531 | 1 | 2 | Eif2s1 |
| Frmd8 | 0,000666395 | 0,338226655 | 0,444 | 0,25 | 1 | 2 | Frmd8 |
| Copb1 | 0,000673552 | 0,464644151 | 0,635 | 0,458 | 1 | 2 | Copb1 |
| Ctnnbip1 | 0,000700214 | 0,326421378 | 0,714 | 0,508 | 1 | 2 | Ctnnbip1 |
| Tfg | 0,000703169 | 0,274631239 | 0,635 | 0,392 | 1 | 2 | Tfg |
| Ap1s1 | 0,00070982 | 0,279626891 | 0,698 | 0,492 | 1 | 2 | Ap1s1 |
| Scarf11 | 0,000777111 | 0,259479954 | 0,587 | 0,362 | 1 | 2 | Scarf1 |
| Fn1 | 0,000786449 | 0,293011267 | 0,905 | 0,708 | 1 | 2 | Fn1 |
| Col4a1 | 0,000816841 | 0,291130114 | 1 | 0,723 | 1 | 2 | Col4a1 |
| Plaur | 0,000887263 | 0,326886316 | 0,825 | 0,612 | 1 | 2 | Plaur |
| Ifnar11 | 0,000892599 | 0,350591882 | 0,683 | 0,465 | 1 | 2 | Ifnar1 |
| Gcsh | 0,0009079 | 0,26717742 | 0,524 | 0,327 | 1 | 2 | Gcsh |
| Stam2 | 0,000944102 | 0,330646138 | 0,381 | 0,204 | 1 | 2 | Stam2 |
| Pkig | 0,00095317 | 0,252950602 | 0,889 | 0,665 | 1 | 2 | Pkig |
| Actr3 | 0,001009826 | 0,279568204 | 0,952 | 0,838 | 1 | 2 | Actr3 |
| Eif1a | 0,001015621 | 0,310145379 | 0,635 | 0,423 | 1 | 2 | Eif1a |
| Plod1 | 0,001025489 | 0,251163884 | 0,968 | 0,854 | 1 | 2 | Plod1 |
| Hmgcl | 0,001070552 | 0,258642746 | 0,619 | 0,381 | 1 | 2 | Hmgcl |
| Cd1d1 | 0,001073726 | 0,287109194 | 0,302 | 0,135 | 1 | 2 | Cd1d1 |
| Gfm1 | 0,001083428 | 0,290374258 | 0,349 | 0,173 | 1 | 2 | Gfm1 |
| Cald11 | 0,001096403 | 0,256769132 | 0,841 | 0,612 | 1 | 2 | Cald1 |
| Cct3 | 0,001104677 | 0,272771362 | 0,794 | 0,558 | 1 | 2 | Cct3 |
| Uba1 | 0,001114744 | 0,370875214 | 0,825 | 0,681 | 1 | 2 | Uba1 |
| Tada1 | 0,001139856 | 0,2511441 | 0,413 | 0,212 | 1 | 2 | Tada1 |
| Prr5l | 0,001145221 | 0,283013404 | 0,333 | 0,158 | 1 | 2 | Prr5l |
| Anxa7 | 0,001197035 | 0,267261701 | 0,762 | 0,558 | 1 | 2 | Anxa7 |
| 9530068E07Rik | 0,0012259 | 0,353260871 | 0,635 | 0,435 | 1 | 2 | 9530068E07Rik |
| Myo1b1 | 0,001225908 | 0,280529103 | 0,746 | 0,523 | 1 | 2 | Myo1b |
| Anxa6 | 0,00123916 | 0,369563472 | 0,667 | 0,492 | 1 | 2 | Anxa6 |
| Ttyh2 | 0,00124198 | 0,315549565 | 0,556 | 0,358 | 1 | 2 | Ttyh2 |
| Rnf19a | 0,001250889 | 0,257914426 | 0,429 | 0,238 | 1 | 2 | Rnf19a |
| Crtap | 0,00126765 | 0,261187317 | 0,667 | 0,435 | 1 | 2 | Crtap |
| Tmem214 | 0,001345724 | 0,346861693 | 0,381 | 0,204 | 1 | 2 | Tmem214 |
| Poldip3 | 0,001406184 | 0,330855842 | 0,476 | 0,296 | 1 | 2 | Poldip3 |
| Casp6 | 0,001422868 | 0,290528512 | 0,492 | 0,3 | 1 | 2 | Casp6 |
| Kctd5 | 0,001528174 | 0,291078897 | 0,444 | 0,258 | 1 | 2 | Kctd5 |
| Yes1 | 0,001544918 | 0,287316984 | 0,556 | 0,365 | 1 | 2 | Yes1 |
| Hmces | 0,001572261 | 0,275356051 | 0,365 | 0,192 | 1 | 2 | Hmces |
| Ptbp1 | 0,001579548 | 0,306071968 | 0,714 | 0,542 | 1 | 2 | Ptbp1 |
| Traf7 | 0,001581325 | 0,272882145 | 0,571 | 0,362 | 1 | 2 | Traf7 |
| Arhgef7 | 0,001605248 | 0,251359794 | 0,698 | 0,485 | 1 | 2 | Arhgef7 |
| Cops5 | 0,001642125 | 0,267492941 | 0,619 | 0,408 | 1 | 2 | Cops5 |
| Ehd31 | 0,00167387 | 0,263667266 | 0,556 | 0,331 | 1 | 2 | Ehd3 |
| Cdc16 | 0,001708008 | 0,280168077 | 0,492 | 0,292 | 1 | 2 | Cdc16 |
| Eif4h | 0,001799799 | 0,278372575 | 0,857 | 0,727 | 1 | 2 | Eif4h |
| Tapbp | 0,001813701 | 0,261460989 | 0,857 | 0,712 | 1 | 2 | Tapbp |
| Ccnd31 | 0,0022267 | 0,296985461 | 0,746 | 0,577 | 1 | 2 | Ccnd3 |
| Il11ra1 | 0,00222695 | 0,31249889 | 0,381 | 0,212 | 1 | 2 | Il11ra1 |
| Klhdc2 | 0,002313322 | 0,257486497 | 0,365 | 0,196 | 1 | 2 | Klhdc2 |
| Mapk7 | 0,002391025 | 0,257415361 | 0,27 | 0,123 | 1 | 2 | Mapk7 |
| Get4 | 0,002485012 | 0,269071463 | 0,603 | 0,396 | 1 | 2 | Get4 |
| Edem2 | 0,002490567 | 0,354644381 | 0,444 | 0,277 | 1 | 2 | Edem2 |
| Nckipsd | 0,002557388 | 0,325251997 | 0,381 | 0,219 | 1 | 2 | Nckipsd |
| Dars | 0,002708542 | 0,348392756 | 0,508 | 0,335 | 1 | 2 | Dars |
| Slc20a1 | 0,002754092 | 0,270345847 | 0,524 | 0,338 | 1 | 2 | Slc20a1 |
| P4hb | 0,003111229 | 0,28552238 | 0,952 | 0,942 | 1 | 2 | P4hb |
| Hspa5 | 0,003115925 | 0,376603781 | 1 | 0,862 | 1 | 2 | Hspa5 |
| 1110059E24Rik | 0,003248623 | 0,302676532 | 0,444 | 0,269 | 1 | 2 | 1110059E24Rik |
| Ube2l6 | 0,003311118 | 0,292013546 | 0,413 | 0,227 | 1 | 2 | Ube2l6 |
| Nfkbia | 0,003798666 | 0,274761512 | 0,905 | 0,835 | 1 | 2 | Nfkbia |
| Bcl10 | 0,003881843 | 0,293626817 | 0,651 | 0,465 | 1 | 2 | Bcl10 |
| Rnf141 | 0,003951183 | 0,274339192 | 0,556 | 0,369 | 1 | 2 | Rnf141 |
| Cct7 | 0,004006103 | 0,251547579 | 0,794 | 0,623 | 1 | 2 | Cct7 |
| Fam32a | 0,004131337 | 0,271559096 | 0,667 | 0,454 | 1 | 2 | Fam32a |
| Eno1 | 0,004160975 | 0,321435903 | 0,984 | 0,923 | 1 | 2 | Eno1 |
| Adipor1 | 0,00418974 | 0,251024454 | 0,921 | 0,792 | 1 | 2 | Adipor1 |
| Ifi47 | 0,004427702 | 0,326187529 | 0,54 | 0,362 | 1 | 2 | Ifi47 |
| Ogfr | 0,004527322 | 0,279453336 | 0,413 | 0,254 | 1 | 2 | Ogfr |
| Prcp | 0,004559406 | 0,299286087 | 0,778 | 0,685 | 1 | 2 | Prcp |
| Cr1l | 0,004702558 | 0,259384954 | 0,746 | 0,546 | 1 | 2 | Cr1l |
| H2-K1 | 0,004709882 | 0,29746071 | 1 | 0,965 | 1 | 2 | H2-K1 |
| Gbp3 | 0,004805358 | 0,269875592 | 0,476 | 0,296 | 1 | 2 | Gbp3 |
| Ch25h | 0,004819591 | 0,403986702 | 0,413 | 0,227 | 1 | 2 | Ch25h |
| Xbp1 | 0,004919947 | 0,287820552 | 0,889 | 0,8 | 1 | 2 | Xbp1 |
| Bcl2l11 | 0,005228182 | 0,25316221 | 0,397 | 0,235 | 1 | 2 | Bcl2l11 |
| Ackr2 | 0,005537675 | 0,266799576 | 0,46 | 0,288 | 1 | 2 | Ackr2 |
| Tmem185b | 0,005608089 | 0,25460657 | 0,349 | 0,196 | 1 | 2 | Tmem185b |
| Sft2d2 | 0,00561381 | 0,284257045 | 0,508 | 0,335 | 1 | 2 | Sft2d2 |
| Ctgf | 0,006103878 | 0,531702143 | 0,429 | 0,265 | 1 | 2 | Ctgf |
| Atp6v0a2 | 0,006329497 | 0,313036485 | 0,492 | 0,327 | 1 | 2 | Atp6v0a2 |
| Capn2 | 0,006909175 | 0,274536265 | 0,603 | 0,458 | 1 | 2 | Capn2 |
| Tacc1 | 0,007724965 | 0,259041108 | 0,841 | 0,769 | 1 | 2 | Tacc1 |
| Cdk2 | 0,00813524 | 0,264808794 | 0,46 | 0,315 | 1 | 2 | Cdk2 |
| Atg4b | 0,008226669 | 0,304807963 | 0,413 | 0,265 | 1 | 2 | Atg4b |
| Tram1 | 0,008330761 | 0,271963147 | 0,81 | 0,7 | 1 | 2 | Tram1 |
| Errfi1 | 0,008592131 | 0,274909699 | 0,302 | 0,158 | 1 | 2 | Errfi1 |
| Tap1 | 0,008605911 | 0,387636961 | 0,556 | 0,4 | 1 | 2 | Tap1 |
| Ssr3 | 0,009078385 | 0,273748097 | 0,921 | 0,758 | 1 | 2 | Ssr3 |
| Cyr61 | 0,009335657 | 0,590767429 | 0,349 | 0,204 | 1 | 2 | Cyr61 |
| Fabp4 | 8,09E-24 | 0,969905905 | 0,655 | 0,087 | 1,28E-19 | 3 | Fabp4 |
| Kcnj8 | 5,31E-22 | 1,397253048 | 0,655 | 0,109 | 8,42E-18 | 3 | Kcnj8 |
| Gpihbp1 | 1,04E-21 | 1,656019103 | 0,931 | 0,442 | 1,65E-17 | 3 | Gpihbp1 |
| Abcc9 | 8,57E-19 | 1,238559926 | 0,862 | 0,317 | 1,36E-14 | 3 | Abcc9 |
| Kcne31 | 1,48E-17 | 1,075913637 | 0,914 | 0,385 | 2,35E-13 | 3 | Kcne3 |
| Angpt21 | 4,51E-17 | 1,470629012 | 0,69 | 0,177 | 7,15E-13 | 3 | Angpt2 |
| Trp53i111 | 1,42E-16 | 1,018308562 | 0,983 | 0,6 | 2,25E-12 | 3 | Trp53i11 |
| Adm1 | 2,51E-16 | 1,065130965 | 0,759 | 0,234 | 3,98E-12 | 3 | Adm |
| Kit1 | 1,07E-15 | 1,044057715 | 0,81 | 0,302 | 1,70E-11 | 3 | Kit |
| Aqp1 | 3,93E-15 | 0,858404122 | 0,552 | 0,106 | 6,23E-11 | 3 | Aqp1 |
| Gng11 | 4,98E-15 | 0,84990562 | 0,966 | 0,702 | 7,89E-11 | 3 | Gng11 |
| Maged1 | 8,68E-15 | 1,577206711 | 0,845 | 0,442 | 1,38E-10 | 3 | Maged1 |
| Sparc1 | 4,32E-14 | 0,701997591 | 1 | 0,747 | 6,85E-10 | 3 | Sparc |
| Apln1 | 4,55E-14 | 1,332193731 | 0,741 | 0,302 | 7,22E-10 | 3 | Apln |
| Adgrf5 | 1,13E-13 | 0,908757319 | 0,966 | 0,619 | 1,80E-09 | 3 | Adgrf5 |
| Lamb11 | 1,59E-13 | 0,961999459 | 0,931 | 0,604 | 2,52E-09 | 3 | Lamb1 |
| Sparcl11 | 1,67E-13 | 0,816852055 | 1 | 0,664 | 2,65E-09 | 3 | Sparcl1 |
| Cdh13 | 1,70E-13 | 0,94501568 | 0,966 | 0,577 | 2,70E-09 | 3 | Cdh13 |
| Cdh5 | 3,11E-13 | 0,689653775 | 1 | 0,675 | 4,92E-09 | 3 | Cdh5 |
| Col4a21 | 5,86E-13 | 0,725621836 | 1 | 0,717 | 9,29E-09 | 3 | Col4a2 |
| Pecam12 | 1,61E-12 | 0,653040642 | 1 | 0,721 | 2,55E-08 | 3 | Pecam1 |
| Flt4 | 2,73E-12 | 0,774541339 | 0,931 | 0,585 | 4,33E-08 | 3 | Flt4 |
| Inhbb | 2,99E-12 | 0,801169776 | 0,828 | 0,377 | 4,73E-08 | 3 | Inhbb |

| | | | | | | | |
|---|---|---|---|---|---|---|---|
| Bvht | 5,62E-05 | 0,294822414 | 0,562 | 0,344 | 0,891612379 | T_CLEC | Bvht |
| Pcdhgc4 | 5,95E-05 | 0,363977925 | 0,562 | 0,362 | 0,943006612 | T_CLEC | Pcdhgc4 |
| Pakap | 6,13E-05 | 0,38214312 | 0,319 | 0,135 | 0,971257862 | T_CLEC | Pakap |
| Adgrl2 | 6,32E-05 | 0,390200115 | 0,519 | 0,307 | 1 | T_CLEC | Adgrl2 |
| Slc29a1 | 6,64E-05 | 0,505126739 | 0,688 | 0,577 | 1 | T_CLEC | Slc29a1 |
| Ndfip1 | 7,28E-05 | 0,36986945 | 0,8 | 0,712 | 1 | T_CLEC | Ndfip1 |
| Zbtb20 | 7,60E-05 | 0,409627354 | 0,656 | 0,491 | 1 | T_CLEC | Zbtb20 |
| Afdn | 8,20E-05 | 0,340616712 | 0,625 | 0,429 | 1 | T_CLEC | Afdn |
| Sh3tc1 | 8,55E-05 | 0,380923808 | 0,356 | 0,166 | 1 | T_CLEC | Sh3tc1 |
| Bmp1 | 8,67E-05 | 0,260051547 | 0,419 | 0,215 | 1 | T_CLEC | Bmp1 |
| Arhgap26 | 8,74E-05 | 0,339685257 | 0,519 | 0,325 | 1 | T_CLEC | Arhgap26 |
| Ly6a1 | 8,83E-05 | 0,293208337 | 0,906 | 0,656 | 1 | T_CLEC | Ly6a |
| Itgb3 | 9,35E-05 | 0,320069191 | 0,469 | 0,276 | 1 | T_CLEC | Itgb3 |
| Tspan12 | 9,60E-05 | 0,483135922 | 0,531 | 0,337 | 1 | T_CLEC | Tspan12 |
| Parva | 9,89E-05 | 0,301882302 | 0,45 | 0,245 | 1 | T_CLEC | Parva |
| Reep3 | 9,94E-05 | 0,320475597 | 0,675 | 0,509 | 1 | T_CLEC | Reep3 |
| Crmp1 | 0,000102717 | 0,310282239 | 0,288 | 0,11 | 1 | T_CLEC | Crmp1 |
| Pik3c2b | 0,000111801 | 0,319685118 | 0,588 | 0,405 | 1 | T_CLEC | Pik3c2b |
| Chst1 | 0,000114525 | 0,303281245 | 0,388 | 0,19 | 1 | T_CLEC | Chst1 |
| Actn41 | 0,000114741 | 0,301318744 | 0,9 | 0,773 | 1 | T_CLEC | Actn4 |
| Kcnq1 | 0,000118255 | 0,430897395 | 0,531 | 0,35 | 1 | T_CLEC | Kcnq1 |
| Tnfaip11 | 0,000124338 | 0,31830248 | 0,8 | 0,564 | 1 | T_CLEC | Tnfaip1 |
| S100a13 | 0,000131118 | 0,268095184 | 0,812 | 0,706 | 1 | T_CLEC | S100a13 |
| Cda | 0,000132932 | 0,27808401 | 0,35 | 0,178 | 1 | T_CLEC | Cda |
| Lmna1 | 0,000137631 | 0,340117405 | 0,788 | 0,663 | 1 | T_CLEC | Lmna |
| Myo6 | 0,000138192 | 0,336224915 | 0,269 | 0,11 | 1 | T_CLEC | Myo6 |
| Kit1 | 0,000145496 | 0,415920687 | 0,494 | 0,294 | 1 | T_CLEC | Kit |
| Arl4a | 0,000146001 | 0,410487366 | 0,544 | 0,356 | 1 | T_CLEC | Arl4a |
| Macf1 | 0,000152552 | 0,344420381 | 0,8 | 0,706 | 1 | T_CLEC | Macf1 |
| Clic41 | 0,000152929 | 0,276785766 | 0,906 | 0,798 | 1 | T_CLEC | Clic4 |
| Dok4 | 0,000155742 | 0,364853341 | 0,356 | 0,172 | 1 | T_CLEC | Dok4 |
| Sox13 | 0,000178885 | 0,280631472 | 0,338 | 0,166 | 1 | T_CLEC | Sox13 |
| Gnas1 | 0,000179892 | 0,312170038 | 0,938 | 0,859 | 1 | T_CLEC | Gnas |
| Gnb41 | 0,000180478 | 0,312900797 | 0,562 | 0,337 | 1 | T_CLEC | Gnb4 |
| Nhsl1 | 0,000186466 | 0,326264137 | 0,45 | 0,27 | 1 | T_CLEC | Nhsl1 |
| Cep170 | 0,000207056 | 0,287051096 | 0,438 | 0,276 | 1 | T_CLEC | Cep170 |
| Pde4b | 0,000207417 | 0,414544573 | 0,7 | 0,528 | 1 | T_CLEC | Pde4b |
| Slc44a1 | 0,000207879 | 0,298591531 | 0,569 | 0,362 | 1 | T_CLEC | Slc44a1 |
| Antxr1 | 0,000210919 | 0,250844934 | 0,369 | 0,196 | 1 | T_CLEC | Antxr1 |
| Utrn | 0,00021538 | 0,31240315 | 0,538 | 0,356 | 1 | T_CLEC | Utrn |
| Xiap | 0,000216332 | 0,354682988 | 0,706 | 0,558 | 1 | T_CLEC | Xiap |
| Hes11 | 0,00021634 | 0,302590425 | 0,694 | 0,466 | 1 | T_CLEC | Hes1 |
| Anxa1 | 0,000223214 | 0,370474088 | 0,775 | 0,632 | 1 | T_CLEC | Anxa1 |
| Gm28875 | 0,000224252 | 0,278585464 | 0,325 | 0,153 | 1 | T_CLEC | Gm28875 |
| Fgfr1 | 0,000236604 | 0,322882105 | 0,35 | 0,178 | 1 | T_CLEC | Fgfr1 |
| Knop1 | 0,00024405 | 0,289053655 | 0,562 | 0,362 | 1 | T_CLEC | Knop1 |
| N4bp3 | 0,000252129 | 0,434468526 | 0,506 | 0,325 | 1 | T_CLEC | N4bp3 |
| Tmed9 | 0,000256131 | 0,356753475 | 0,675 | 0,546 | 1 | T_CLEC | Tmed9 |
| mt-Cytb | 0,000263912 | 0,288288319 | 1 | 1 | 1 | T_CLEC | mt-Cytb |
| Lxn1 | 0,00026563 | 0,303804144 | 0,725 | 0,552 | 1 | T_CLEC | Lxn |
| Chst2 | 0,000265848 | 0,264674026 | 0,331 | 0,16 | 1 | T_CLEC | Chst2 |
| Igfbp41 | 0,000269983 | 0,284714353 | 0,844 | 0,724 | 1 | T_CLEC | Igfbp4 |
| Itsn2 | 0,000280444 | 0,345078851 | 0,694 | 0,515 | 1 | T_CLEC | Itsn2 |
| Efnb2 | 0,000286665 | 0,344939004 | 0,344 | 0,178 | 1 | T_CLEC | Efnb2 |
| Gprc5b | 0,00029027 | 0,277617374 | 0,275 | 0,123 | 1 | T_CLEC | Gprc5b |
| Wwc2 | 0,000290484 | 0,266180351 | 0,381 | 0,202 | 1 | T_CLEC | Wwc2 |
| Serpinb6a | 0,000290698 | 0,379676675 | 0,669 | 0,515 | 1 | T_CLEC | Serpinb6a |
| Tspan6 | 0,000297723 | 0,39338599 | 0,431 | 0,252 | 1 | T_CLEC | Tspan6 |
| Kmt2a | 0,000318277 | 0,319749982 | 0,4 | 0,227 | 1 | T_CLEC | Kmt2a |
| Fam212a | 0,000324941 | 0,272117349 | 0,412 | 0,233 | 1 | T_CLEC | Fam212a |
| Myo18a | 0,000326891 | 0,304157189 | 0,462 | 0,288 | 1 | T_CLEC | Myo18a |
| Slc43a3 | 0,000340239 | 0,388144511 | 0,5 | 0,319 | 1 | T_CLEC | Slc43a3 |
| Nxn | 0,000340697 | 0,331971491 | 0,338 | 0,178 | 1 | T_CLEC | Nxn |
| Luzp1 | 0,000347366 | 0,417206723 | 0,712 | 0,595 | 1 | T_CLEC | Luzp1 |
| Myh9 | 0,000350885 | 0,344124233 | 0,975 | 0,975 | 1 | T_CLEC | Myh9 |
| Pitpnc1 | 0,000354424 | 0,331671698 | 0,681 | 0,54 | 1 | T_CLEC | Pitpnc1 |
| Ywhah | 0,000359595 | 0,370578185 | 0,9 | 0,859 | 1 | T_CLEC | Ywhah |
| Atrn | 0,000364932 | 0,336049233 | 0,375 | 0,209 | 1 | T_CLEC | Atrn |
| Wasf2 | 0,000368648 | 0,379431035 | 0,806 | 0,834 | 1 | T_CLEC | Wasf2 |
| Tmem263 | 0,000369705 | 0,287203999 | 0,519 | 0,325 | 1 | T_CLEC | Tmem263 |
| Sema6a | 0,000375916 | 0,252664637 | 0,5 | 0,313 | 1 | T_CLEC | Sema6a |
| BC028528 | 0,000392859 | 0,367619491 | 0,856 | 0,785 | 1 | T_CLEC | BC028528 |
| Tgfb1i1 | 0,000394662 | 0,309433652 | 0,462 | 0,27 | 1 | T_CLEC | Tgfb1i1 |
| Ptpn12 | 0,000415412 | 0,288709055 | 0,531 | 0,325 | 1 | T_CLEC | Ptpn12 |
| Pkp4 | 0,00049107 | 0,335522694 | 0,412 | 0,245 | 1 | T_CLEC | Pkp4 |
| Hyal2 | 0,000501786 | 0,321001844 | 0,525 | 0,325 | 1 | T_CLEC | Hyal2 |
| Ptk2 | 0,000502855 | 0,320442871 | 0,506 | 0,294 | 1 | T_CLEC | Ptk2 |
| Kdm5b | 0,000507986 | 0,36887685 | 0,294 | 0,147 | 1 | T_CLEC | Kdm5b |
| Cers4 | 0,000518262 | 0,27065144 | 0,319 | 0,147 | 1 | T_CLEC | Cers4 |
| Plcb1 | 0,000532715 | 0,309519361 | 0,3 | 0,147 | 1 | T_CLEC | Plcb1 |
| Eif3a | 0,000544827 | 0,341909235 | 0,694 | 0,601 | 1 | T_CLEC | Eif3a |
| Gm10925 | 0,000551859 | 0,33313548 | 0,962 | 0,957 | 1 | T_CLEC | Gm10925 |
| Magi1 | 0,000557588 | 0,334726877 | 0,331 | 0,178 | 1 | T_CLEC | Magi1 |
| Myo1d | 0,000560493 | 0,302865678 | 0,281 | 0,129 | 1 | T_CLEC | Myo1d |
| Pcnp | 0,000566622 | 0,484362351 | 0,738 | 0,613 | 1 | T_CLEC | Pcnp |
| Plcg1 | 0,000603761 | 0,276300851 | 0,581 | 0,399 | 1 | T_CLEC | Plcg1 |
| Slc44a2 | 0,000625927 | 0,253523199 | 0,506 | 0,301 | 1 | T_CLEC | Slc44a2 |
| Lrrc32 | 0,000637726 | 0,260191584 | 0,312 | 0,147 | 1 | T_CLEC | Lrrc32 |
| Ndrg1 | 0,000640727 | 0,406177704 | 0,538 | 0,356 | 1 | T_CLEC | Ndrg1 |
| Gna11 | 0,000651048 | 0,284936103 | 0,644 | 0,46 | 1 | T_CLEC | Gna11 |
| Cd151 | 0,000659299 | 0,343540265 | 0,556 | 0,399 | 1 | T_CLEC | Cd151 |
| Clec1a | 0,000660807 | 0,331238128 | 0,5 | 0,307 | 1 | T_CLEC | Clec1a |
| CT010467.1 | 0,000663896 | 0,330126578 | 1 | 1 | 1 | T_CLEC | CT010467.1 |
| Jam2 | 0,000664565 | 0,413803251 | 0,469 | 0,276 | 1 | T_CLEC | Jam2 |
| Pea15a | 0,000722173 | 0,260030501 | 0,875 | 0,828 | 1 | T_CLEC | Pea15a |
| Sdpr | 0,000725861 | 0,342258114 | 0,369 | 0,196 | 1 | T_CLEC | Sdpr |
| Selenom | 0,000743172 | 0,288480793 | 0,619 | 0,442 | 1 | T_CLEC | Selenom |
| 2810025M15Rik | 0,00082117 | 0,279793196 | 0,538 | 0,38 | 1 | T_CLEC | 2810025M15Rik |
| Rock2 | 0,000821921 | 0,305713567 | 0,562 | 0,387 | 1 | T_CLEC | Rock2 |
| Sh2d3c1 | 0,000855761 | 0,258887412 | 0,731 | 0,497 | 1 | T_CLEC | Sh2d3c |
| Kdelr2 | 0,000874899 | 0,368023934 | 0,925 | 0,89 | 1 | T_CLEC | Kdelr2 |
| Nostrin1 | 0,000882154 | 0,324043712 | 0,65 | 0,472 | 1 | T_CLEC | Nostrin |
| Pja2 | 0,000947355 | 0,292688817 | 0,5 | 0,313 | 1 | T_CLEC | Pja2 |
| 9530082P21Rik | 0,000954694 | 0,317869124 | 0,606 | 0,436 | 1 | T_CLEC | 9530082P21Rik |
| Gimap1 | 0,000968009 | 0,315140841 | 0,519 | 0,362 | 1 | T_CLEC | Gimap1 |
| Cdc42ep31 | 0,000977205 | 0,303534927 | 0,544 | 0,368 | 1 | T_CLEC | Cdc42ep3 |
| Ccdc88a | 0,000995849 | 0,309125864 | 0,394 | 0,221 | 1 | T_CLEC | Ccdc88a |
| Cracr2b | 0,001051215 | 0,262238606 | 0,6 | 0,423 | 1 | T_CLEC | Cracr2b |
| Nudt3 | 0,001094877 | 0,392298334 | 0,55 | 0,454 | 1 | T_CLEC | Nudt3 |
| Cmip | 0,001142589 | 0,444285402 | 0,631 | 0,515 | 1 | T_CLEC | Cmip |
| Abcg1 | 0,00116459 | 0,442718814 | 0,744 | 0,62 | 1 | T_CLEC | Abcg1 |
| Ablim1 | 0,001251842 | 0,262806264 | 0,475 | 0,301 | 1 | T_CLEC | Ablim1 |
| Prkd3 | 0,001281146 | 0,267707643 | 0,381 | 0,233 | 1 | T_CLEC | Prkd3 |
| Itga6 | 0,001356501 | 0,305633704 | 0,788 | 0,699 | 1 | T_CLEC | Itga6 |
| Ralgapa1 | 0,001363984 | 0,339566881 | 0,331 | 0,19 | 1 | T_CLEC | Ralgapa1 |
| Klc1 | 0,001385996 | 0,311567039 | 0,531 | 0,356 | 1 | T_CLEC | Klc1 |
| Dcbld1 | 0,001390359 | 0,389347912 | 0,394 | 0,239 | 1 | T_CLEC | Dcbld1 |
| Cds2 | 0,00141512 | 0,294881811 | 0,556 | 0,387 | 1 | T_CLEC | Cds2 |
| Plekhg1 | 0,001527563 | 0,282052799 | 0,481 | 0,301 | 1 | T_CLEC | Plekhg1 |
| Slc39a10 | 0,001670643 | 0,276357912 | 0,394 | 0,227 | 1 | T_CLEC | Slc39a10 |
| Ttc3 | 0,001682459 | 0,26267439 | 0,456 | 0,294 | 1 | T_CLEC | Ttc3 |
| Alg14 | 0,001694156 | 0,254388604 | 0,375 | 0,221 | 1 | T_CLEC | Alg14 |

| | | | | | | | |
|---|---|---|---|---|---|---|---|
| Esm1 | 3,36E-12 | 1,376155896 | 0,448 | 0,094 | 5,32E-08 | 3 | Esm1 |
| Col4a11 | 5,60E-12 | 0,692083609 | 1 | 0,728 | 8,88E-08 | 3 | Col4a1 |
| Creb3l21 | 2,20E-11 | 0,843799329 | 0,862 | 0,506 | 3,49E-07 | 3 | Creb3l2 |
| S1pr11 | 2,89E-11 | 0,706335588 | 1 | 0,717 | 4,57E-07 | 3 | S1pr1 |
| Kcnj2 | 3,71E-11 | 0,357473388 | 0,483 | 0,109 | 5,88E-07 | 3 | Kcnj2 |
| Aplp2 | 6,22E-11 | 0,781074018 | 0,983 | 0,792 | 9,86E-07 | 3 | Aplp2 |
| Map4k41 | 6,70E-11 | 0,529558759 | 0,897 | 0,502 | 1,06E-06 | 3 | Map4k4 |
| Pde4b1 | 9,68E-11 | 0,915846342 | 0,845 | 0,562 | 1,54E-06 | 3 | Pde4b |
| Ackr31 | 9,83E-11 | 0,768548551 | 0,759 | 0,347 | 1,56E-06 | 3 | Ackr3 |
| Rell1 | 1,45E-10 | 0,714731043 | 0,828 | 0,43 | 2,30E-06 | 3 | Rell1 |
| Chst11 | 1,71E-10 | 0,468512773 | 0,638 | 0,211 | 2,71E-06 | 3 | Chst1 |
| Adamts71 | 3,22E-10 | 0,518987746 | 0,5 | 0,14 | 5,11E-06 | 3 | Adamts7 |
| Hspg2 | 3,61E-10 | 0,664022381 | 1 | 0,642 | 5,72E-06 | 3 | Hspg2 |
| Anxa61 | 6,16E-10 | 0,873153497 | 0,793 | 0,468 | 9,76E-06 | 3 | Anxa6 |
| Pde2a1 | 6,44E-10 | 0,801106378 | 0,741 | 0,332 | 1,02E-05 | 3 | Pde2a |
| Lama41 | 7,01E-10 | 0,738664974 | 0,948 | 0,562 | 1,11E-05 | 3 | Lama4 |
| Lamc1 | 7,78E-10 | 0,676173522 | 0,966 | 0,623 | 1,23E-05 | 3 | Lamc1 |
| Mical2 | 9,41E-10 | 0,291049144 | 0,31 | 0,053 | 1,49E-05 | 3 | Mical2 |
| Plod11 | 1,42E-09 | 0,674670787 | 0,948 | 0,86 | 2,25E-05 | 3 | Plod1 |
| Cd2761 | 2,25E-09 | 0,643714681 | 0,655 | 0,291 | 3,56E-05 | 3 | Cd276 |
| Pxdn1 | 2,32E-09 | 0,588696607 | 0,845 | 0,464 | 3,67E-05 | 3 | Pxdn |
| Tpm41 | 2,60E-09 | 0,568893369 | 1 | 0,909 | 4,12E-05 | 3 | Tpm4 |
| Ednrb | 3,16E-09 | 0,747293799 | 0,776 | 0,381 | 5,01E-05 | 3 | Ednrb |
| 2900026A02Rik | 5,37E-09 | 0,605388828 | 0,759 | 0,415 | 8,51E-05 | 3 | 2900026A02Rik |
| Sema3f | 5,81E-09 | 0,605443848 | 0,879 | 0,562 | 9,22E-05 | 3 | Sema3f |
| Plvap1 | 6,00E-09 | 0,75715801 | 0,931 | 0,634 | 9,50E-05 | 3 | Plvap |
| Ptprm | 7,60E-09 | 0,804944265 | 0,776 | 0,442 | 0,00012052 | 3 | Ptprm |
| 8430408G22Rik | 8,36E-09 | 0,712118736 | 0,534 | 0,189 | 0,000132548 | 3 | 8430408G22Rik |
| Lrrc8a1 | 1,06E-08 | 0,615771295 | 0,966 | 0,747 | 0,00016779 | 3 | Lrrc8a |
| N4bp31 | 1,13E-08 | 0,646835642 | 0,724 | 0,347 | 0,000178778 | 3 | N4bp3 |
| Rbp11 | 1,15E-08 | 0,584863456 | 0,914 | 0,562 | 0,000182269 | 3 | Rbp1 |
| Fscn11 | 1,20E-08 | 0,612728377 | 0,983 | 0,653 | 0,000190516 | 3 | Fscn1 |
| Gja12 | 2,75E-08 | 0,594702008 | 0,948 | 0,596 | 0,00043664 | 3 | Gja1 |
| Ramp3 | 2,83E-08 | 0,908758173 | 0,603 | 0,253 | 0,000448478 | 3 | Ramp3 |
| Cdc42ep31 | 3,28E-08 | 0,595088115 | 0,741 | 0,392 | 0,000520267 | 3 | Cdc42ep3 |
| Spata13 | 3,40E-08 | 0,499982282 | 0,776 | 0,415 | 0,000539012 | 3 | Spata13 |
| Serpinh11 | 3,72E-08 | 0,44905406 | 0,983 | 0,66 | 0,000590096 | 3 | Serpinh1 |
| Rgcc | 5,01E-08 | 0,815358527 | 0,724 | 0,411 | 0,00079445 | 3 | Rgcc |
| Sema6d | 5,12E-08 | 0,582095825 | 0,914 | 0,562 | 0,000811806 | 3 | Sema6d |
| Pkm1 | 6,07E-08 | 0,463136301 | 1 | 0,977 | 0,000961625 | 3 | Pkm |
| Nudt4 | 6,44E-08 | 0,623730451 | 0,672 | 0,355 | 0,001021371 | 3 | Nudt4 |
| Adgrl41 | 6,90E-08 | 0,521242532 | 0,966 | 0,653 | 0,001094349 | 3 | Adgrl4 |
| Nid1 | 7,51E-08 | 0,682575264 | 0,897 | 0,642 | 0,001190012 | 3 | Nid1 |
| Dll4 | 1,05E-07 | 0,657501471 | 0,741 | 0,415 | 0,001667413 | 3 | Dll4 |
| Pcsk6 | 1,35E-07 | 0,368099139 | 0,293 | 0,06 | 0,002140745 | 3 | Pcsk6 |
| Spsb1 | 1,36E-07 | 0,713386301 | 0,69 | 0,385 | 0,002148329 | 3 | Spsb1 |
| Gprc5b | 1,40E-07 | 0,496803969 | 0,431 | 0,147 | 0,002226356 | 3 | Gprc5b |
| Tcf41 | 1,44E-07 | 0,530500596 | 1 | 0,857 | 0,002287798 | 3 | Tcf4 |
| Nid21 | 1,61E-07 | 0,540913702 | 0,879 | 0,57 | 0,002549226 | 3 | Nid2 |
| Ptrf1 | 1,62E-07 | 0,636042805 | 0,948 | 0,649 | 0,002567039 | 3 | Ptrf |
| Olfml2a1 | 1,68E-07 | 0,52066864 | 0,638 | 0,279 | 0,002670216 | 3 | Olfml2a |
| Gnb41 | 1,79E-07 | 0,548724218 | 0,759 | 0,381 | 0,002834947 | 3 | Gnb4 |
| Rps6ka21 | 1,89E-07 | 0,367755982 | 0,552 | 0,219 | 0,002996121 | 3 | Rps6ka2 |
| Tram2 | 1,93E-07 | 0,527643723 | 0,862 | 0,577 | 0,003060765 | 3 | Tram2 |
| Robo4 | 2,00E-07 | 0,766382474 | 0,845 | 0,536 | 0,00317268 | 3 | Robo4 |
| Ptgr1 | 2,17E-07 | 0,621187991 | 0,776 | 0,494 | 0,003434609 | 3 | Ptgr1 |
| Pcdh11 | 2,54E-07 | 0,386285021 | 0,724 | 0,381 | 0,004033063 | 3 | Pcdh1 |
| Mest | 3,17E-07 | 0,546267195 | 0,569 | 0,226 | 0,005018842 | 3 | Mest |
| Mxra7 | 3,23E-07 | 0,27406386 | 0,517 | 0,192 | 0,005121795 | 3 | Mxra7 |
| Mcam1 | 3,33E-07 | 0,407333317 | 0,983 | 0,675 | 0,005280405 | 3 | Mcam |
| Xylt1 | 3,65E-07 | 0,260737935 | 0,431 | 0,147 | 0,005789712 | 3 | Xylt1 |
| Cav1 | 4,30E-07 | 0,590668038 | 0,828 | 0,475 | 0,006820451 | 3 | Cav1 |
| Dapk2 | 4,33E-07 | 0,504449028 | 0,328 | 0,083 | 0,006863138 | 3 | Dapk2 |
| Angptl41 | 4,34E-07 | 0,543385049 | 0,466 | 0,166 | 0,006880168 | 3 | Angptl4 |
| Ppfibp1 | 5,56E-07 | 0,465915435 | 0,879 | 0,619 | 0,008809813 | 3 | Ppfibp1 |
| Degs1 | 5,66E-07 | 0,612494462 | 0,897 | 0,766 | 0,008973043 | 3 | Degs1 |
| Rnf1221 | 6,06E-07 | 0,313026462 | 0,534 | 0,226 | 0,009604474 | 3 | Rnf122 |
| Lrrc58 | 6,30E-07 | 0,497730158 | 1 | 0,909 | 0,009994412 | 3 | Lrrc58 |
| Gnai2 | 6,92E-07 | 0,336338789 | 0,966 | 0,97 | 0,010968462 | 3 | Gnai2 |
| Oaf1 | 8,16E-07 | 0,492410034 | 0,69 | 0,377 | 0,012935696 | 3 | Oaf |
| Sptbn11 | 8,39E-07 | 0,45706517 | 1 | 0,762 | 0,013307017 | 3 | Sptbn1 |
| Thbs1 | 8,43E-07 | 1,409836225 | 0,448 | 0,181 | 0,013367529 | 3 | Thbs1 |
| Myo101 | 9,18E-07 | 0,465753901 | 0,759 | 0,404 | 0,01455439 | 3 | Myo10 |
| Piezo21 | 1,04E-06 | 0,531940705 | 0,759 | 0,472 | 0,016531079 | 3 | Piezo2 |
| Dysf | 1,28E-06 | 0,489094254 | 0,672 | 0,332 | 0,020327345 | 3 | Dysf |
| Ackr21 | 1,39E-06 | 0,553875976 | 0,586 | 0,264 | 0,022037103 | 3 | Ackr2 |
| Efna11 | 1,41E-06 | 0,44384338 | 0,897 | 0,592 | 0,022310465 | 3 | Efna1 |
| 9430020K01Rik1 | 1,44E-06 | 0,568857518 | 0,862 | 0,532 | 0,022885822 | 3 | 9430020K01Rik |
| Arhgef28 | 1,55E-06 | 0,395165925 | 0,362 | 0,109 | 0,024504098 | 3 | Arhgef28 |
| Arhgef12 | 1,61E-06 | 0,48368336 | 0,672 | 0,392 | 0,025557631 | 3 | Arhgef12 |
| Gm373761 | 1,68E-06 | 0,484141794 | 1 | 0,985 | 0,026685063 | 3 | Gm37376 |
| Enpp6 | 1,75E-06 | 0,537073644 | 0,293 | 0,075 | 0,027704823 | 3 | Enpp6 |
| Raph11 | 1,85E-06 | 0,540594491 | 0,69 | 0,408 | 0,029302157 | 3 | Raph1 |
| Flt11 | 2,24E-06 | 0,607902349 | 0,966 | 0,675 | 0,035478083 | 3 | Flt1 |
| Emcn1 | 2,30E-06 | 0,486871812 | 0,914 | 0,679 | 0,036402344 | 3 | Emcn |
| Malat11 | 2,40E-06 | 0,458353376 | 1 | 0,996 | 0,037992959 | 3 | Malat1 |
| Ccser1 | 2,46E-06 | 0,338932031 | 0,259 | 0,06 | 0,039051068 | 3 | Ccser1 |
| Ctgf1 | 3,05E-06 | 0,740867501 | 0,517 | 0,249 | 0,048347371 | 3 | Ctgf |
| Cav2 | 3,73E-06 | 0,55815938 | 0,776 | 0,521 | 0,059166985 | 3 | Cav2 |
| Ift122 | 3,77E-06 | 0,486274584 | 0,362 | 0,121 | 0,059691298 | 3 | Ift122 |
| Wwtr1 | 4,20E-06 | 0,539827994 | 0,759 | 0,468 | 0,066597675 | 3 | Wwtr1 |
| Fgfr1 | 4,24E-06 | 0,413810149 | 0,5 | 0,211 | 0,067142515 | 3 | Fgfr1 |
| Calm11 | 4,25E-06 | 0,353584366 | 1 | 0,992 | 0,067377053 | 3 | Calm1 |
| Prkcdbp1 | 4,41E-06 | 0,455858681 | 0,948 | 0,675 | 0,069882188 | 3 | Prkcdbp |
| Phactr4 | 4,64E-06 | 0,316629831 | 0,724 | 0,408 | 0,073598613 | 3 | Phactr4 |
| Kctd11 | 4,65E-06 | 0,301232014 | 0,31 | 0,087 | 0,073654529 | 3 | Kctd11 |
| mt-Rnr1 | 4,72E-06 | 0,341656302 | 1 | 1 | 0,074767244 | 3 | mt-Rnr1 |
| Ece12 | 4,99E-06 | 0,4637667 | 1 | 0,717 | 0,079060402 | 3 | Ece1 |
| Man1a | 5,17E-06 | 0,671014666 | 0,707 | 0,472 | 0,081875706 | 3 | Man1a |
| Cd9 | 5,58E-06 | 0,414616883 | 0,948 | 0,845 | 0,08849186 | 3 | Cd9 |
| Itga31 | 5,67E-06 | 0,449878866 | 0,638 | 0,325 | 0,089942209 | 3 | Itga3 |
| Pcdh17 | 6,91E-06 | 0,318821048 | 0,655 | 0,351 | 0,109533413 | 3 | Pcdh17 |
| Myct1 | 8,24E-06 | 0,567302605 | 0,707 | 0,434 | 0,130630998 | 3 | Myct1 |
| Sptan1 | 8,70E-06 | 0,504589065 | 0,862 | 0,713 | 0,137978521 | 3 | Sptan1 |
| Cd47 | 9,78E-06 | 0,518648941 | 0,845 | 0,83 | 0,154995378 | 3 | Cd47 |
| Emid1 | 1,06E-05 | 0,42261645 | 0,569 | 0,268 | 0,167365225 | 3 | Emid1 |
| Dchs1 | 1,06E-05 | 0,550473853 | 0,621 | 0,355 | 0,168186871 | 3 | Dchs1 |
| Vim1 | 1,24E-05 | 0,346000984 | 1 | 0,974 | 0,196434886 | 3 | Vim |
| Alg14 | 1,29E-05 | 0,329324973 | 0,534 | 0,245 | 0,204330193 | 3 | Alg14 |
| Fstl11 | 1,31E-05 | 0,401058262 | 0,828 | 0,468 | 0,207502636 | 3 | Fstl1 |
| Abcg1 | 1,43E-05 | 0,66606024 | 0,845 | 0,645 | 0,226865554 | 3 | Abcg1 |
| Slc12a2 | 1,44E-05 | 0,340417512 | 0,517 | 0,234 | 0,227922541 | 3 | Slc12a2 |
| Yes11 | 1,73E-05 | 0,268978541 | 0,655 | 0,347 | 0,273702506 | 3 | Yes1 |
| Tjp1 | 1,79E-05 | 0,486023432 | 0,724 | 0,438 | 0,283925567 | 3 | Tjp1 |
| Shroom2 | 1,91E-05 | 0,414569017 | 0,707 | 0,426 | 0,30207398 | 3 | Shroom2 |
| Tmem255a | 1,92E-05 | 0,708219196 | 0,293 | 0,102 | 0,305022895 | 3 | Tmem255a |
| Gsap | 2,19E-05 | 0,259281053 | 0,345 | 0,117 | 0,347016086 | 3 | Gsap |
| Tfpi2 | 2,23E-05 | 0,357054687 | 0,379 | 0,14 | 0,353877881 | 3 | Tfpi2 |
| Ptk7 | 2,31E-05 | 0,283481231 | 0,276 | 0,083 | 0,366505621 | 3 | Ptk7 |
| Anxa22 | 2,32E-05 | 0,423693391 | 0,983 | 0,845 | 0,367791439 | 3 | Anxa2 |
| Ahnak1 | 2,47E-05 | 0,472234415 | 0,897 | 0,649 | 0,391598146 | 3 | Ahnak |
| Litaf1 | 2,50E-05 | 0,501054874 | 0,948 | 0,887 | 0,396887302 | 3 | Litaf |

| | | | | | | | |
|---|---|---|---|---|---|---|---|
| Pkn3 | 0,001749733 | 0,275428701 | 0,462 | 0,276 | 1 | T_CLEC | Pkn3 |
| Pde8a | 0,00175401 | 0,268780404 | 0,381 | 0,221 | 1 | T_CLEC | Pde8a |
| Snapc3 | 0,001760945 | 0,25791918 | 0,269 | 0,135 | 1 | T_CLEC | Snapc3 |
| Itga5 | 0,001794426 | 0,357844365 | 0,681 | 0,577 | 1 | T_CLEC | Itga5 |
| Ctdsp2 | 0,001812582 | 0,262783983 | 0,581 | 0,436 | 1 | T_CLEC | Ctdsp2 |
| Hk1 | 0,001891011 | 0,289653142 | 0,5 | 0,344 | 1 | T_CLEC | Hk1 |
| Tinagl1 | 0,001920636 | 0,353216326 | 0,294 | 0,153 | 1 | T_CLEC | Tinagl1 |
| Lats2 | 0,002018519 | 0,305548173 | 0,481 | 0,337 | 1 | T_CLEC | Lats2 |
| Nrep | 0,002044581 | 0,408732183 | 0,55 | 0,472 | 1 | T_CLEC | Nrep |
| Srrm2 | 0,002260807 | 0,366194375 | 0,762 | 0,687 | 1 | T_CLEC | Srrm2 |
| Zdhhc20 | 0,002471077 | 0,343752124 | 0,731 | 0,65 | 1 | T_CLEC | Zdhhc20 |
| Tmem254c | 0,002497832 | 0,331700681 | 0,544 | 0,405 | 1 | T_CLEC | Tmem254c |
| Stim2 | 0,002514976 | 0,256867114 | 0,394 | 0,245 | 1 | T_CLEC | Stim2 |
| Ehd31 | 0,002639697 | 0,29335219 | 0,456 | 0,294 | 1 | T_CLEC | Ehd3 |
| Fkbp7 | 0,002645022 | 0,259691075 | 0,375 | 0,233 | 1 | T_CLEC | Fkbp7 |
| Git1 | 0,002700754 | 0,310582965 | 0,525 | 0,393 | 1 | T_CLEC | Git1 |
| Nras | 0,00277803 | 0,377061434 | 0,762 | 0,73 | 1 | T_CLEC | Nras |
| Tm9sf3 | 0,003041149 | 0,36264105 | 0,688 | 0,656 | 1 | T_CLEC | Tm9sf3 |
| Qk | 0,003104941 | 0,35752943 | 0,831 | 0,822 | 1 | T_CLEC | Qk |
| Pmp22 | 0,003119302 | 0,262236614 | 0,319 | 0,184 | 1 | T_CLEC | Pmp22 |
| Ktn1 | 0,003155091 | 0,384753554 | 0,594 | 0,479 | 1 | T_CLEC | Ktn1 |
| Csgalnact1 | 0,003224025 | 0,258687835 | 0,275 | 0,141 | 1 | T_CLEC | Csgalnact1 |
| Kbtbd11 | 0,003336894 | 0,398340367 | 0,319 | 0,202 | 1 | T_CLEC | Kbtbd11 |
| Ywhaq | 0,003396847 | 0,302274585 | 0,856 | 0,822 | 1 | T_CLEC | Ywhaq |
| Cd81 | 0,003717476 | 0,266026593 | 0,956 | 1 | 1 | T_CLEC | Cd81 |
| Hnrnpa0 | 0,003806744 | 0,272475952 | 0,612 | 0,528 | 1 | T_CLEC | Hnrnpa0 |
| Baz2b | 0,003838272 | 0,294742479 | 0,488 | 0,344 | 1 | T_CLEC | Baz2b |
| Rsu1 | 0,003907325 | 0,340664423 | 0,769 | 0,712 | 1 | T_CLEC | Rsu1 |
| Tspan151 | 0,004113667 | 0,273670603 | 0,45 | 0,288 | 1 | T_CLEC | Tspan15 |
| Calm2 | 0,004116502 | 0,325087898 | 0,944 | 0,975 | 1 | T_CLEC | Calm2 |
| Lrrc8b | 0,004325636 | 0,282599113 | 0,388 | 0,258 | 1 | T_CLEC | Lrrc8b |
| Clip1 | 0,004609183 | 0,262473195 | 0,4 | 0,258 | 1 | T_CLEC | Clip1 |
| Tex2 | 0,004630845 | 0,266233961 | 0,275 | 0,147 | 1 | T_CLEC | Tex2 |
| Arglu1 | 0,004786393 | 0,431641344 | 0,888 | 0,883 | 1 | T_CLEC | Arglu1 |
| Etnk1 | 0,004899447 | 0,252834252 | 0,406 | 0,264 | 1 | T_CLEC | Etnk1 |
| Man1a | 0,004973127 | 0,340541532 | 0,575 | 0,454 | 1 | T_CLEC | Man1a |
| Pgm2l1 | 0,004997479 | 0,274242155 | 0,338 | 0,215 | 1 | T_CLEC | Pgm2l1 |
| C1qtnf1 | 0,005279843 | 0,362553092 | 0,4 | 0,27 | 1 | T_CLEC | C1qtnf1 |
| Exoc3l | 0,005310583 | 0,350765273 | 0,406 | 0,276 | 1 | T_CLEC | Exoc3l |
| Spsb1 | 0,005373574 | 0,406374226 | 0,5 | 0,38 | 1 | T_CLEC | Spsb1 |
| Rgs3 | 0,005383153 | 0,278570909 | 0,45 | 0,313 | 1 | T_CLEC | Rgs3 |
| Smurf1 | 0,005489984 | 0,304696081 | 0,262 | 0,147 | 1 | T_CLEC | Smurf1 |
| Cpne8 | 0,005907166 | 0,263450321 | 0,538 | 0,429 | 1 | T_CLEC | Cpne8 |
| Spata13 | 0,006225216 | 0,393009621 | 0,519 | 0,442 | 1 | T_CLEC | Spata13 |
| Tmod3 | 0,006360464 | 0,324660589 | 0,806 | 0,761 | 1 | T_CLEC | Tmod3 |
| Map1lc3b | 0,006674199 | 0,359498121 | 0,925 | 0,896 | 1 | T_CLEC | Map1lc3b |
| Gbas | 0,006767079 | 0,303243236 | 0,656 | 0,564 | 1 | T_CLEC | Gbas |
| Arf2 | 0,006940577 | 0,306738716 | 0,569 | 0,466 | 1 | T_CLEC | Arf2 |
| Anxa6 | 0,006970939 | 0,4633684 | 0,569 | 0,485 | 1 | T_CLEC | Anxa6 |
| 4931406P16Rik | 0,006972815 | 0,256221479 | 0,588 | 0,436 | 1 | T_CLEC | 4931406P16Rik |
| C330006A16Rik | 0,007122215 | 0,365214361 | 0,681 | 0,613 | 1 | T_CLEC | C330006A16Rik |
| Cyb5r31 | 0,007323879 | 0,306025389 | 0,75 | 0,706 | 1 | T_CLEC | Cyb5r3 |
| Procr | 0,007782965 | 0,250135015 | 0,362 | 0,233 | 1 | T_CLEC | Procr |
| Sptlc1 | 0,007783153 | 0,257727944 | 0,35 | 0,221 | 1 | T_CLEC | Sptlc1 |
| Csnk1a1 | 0,008848546 | 0,270394507 | 0,781 | 0,724 | 1 | T_CLEC | Csnk1a1 |
| Stap2 | 0,008985051 | 0,280054259 | 0,462 | 0,344 | 1 | T_CLEC | Stap2 |
| Zhx3 | 0,009298644 | 0,263782778 | 0,356 | 0,239 | 1 | T_CLEC | Zhx3 |
| Akr1b8 | 0,009748903 | 0,331724812 | 0,444 | 0,319 | 1 | T_CLEC | Akr1b8 |
| Golm1 | 0,009858126 | 0,371577359 | 0,538 | 0,423 | 1 | T_CLEC | Golm1 |

| | | | | | | | |
|---|---|---|---|---|---|---|---|
| Ldb2 | 2,51E-05 | 0,432260169 | 0,845 | 0,577 | 0,397233775 | 3 | Ldb2 |
| Ehd21 | 2,92E-05 | 0,393546713 | 0,776 | 0,483 | 0,463260525 | 3 | Ehd2 |
| Cyr611 | 3,06E-05 | 1,106653475 | 0,431 | 0,189 | 0,484827988 | 3 | Cyr61 |
| Kdr1 | 3,21E-05 | 0,513869233 | 0,862 | 0,619 | 0,509284034 | 3 | Kdr |
| Ndrg1 | 3,37E-05 | 0,896623439 | 0,621 | 0,408 | 0,534213286 | 3 | Ndrg1 |
| Lpp | 3,42E-05 | 0,446363199 | 0,81 | 0,63 | 0,54199634 | 3 | Lpp |
| Tspan151 | 3,82E-05 | 0,449564908 | 0,586 | 0,321 | 0,605543235 | 3 | Tspan15 |
| Hif1a | 4,00E-05 | 0,572910499 | 0,931 | 0,781 | 0,634745764 | 3 | Hif1a |
| Akap2 | 4,26E-05 | 0,39523042 | 0,621 | 0,347 | 0,675633443 | 3 | Akap2 |
| Grb10 | 4,27E-05 | 0,548075186 | 0,638 | 0,404 | 0,676848723 | 3 | Grb10 |
| Plec1 | 4,32E-05 | 0,469254189 | 0,931 | 0,747 | 0,684916766 | 3 | Plec |
| Igfbp32 | 4,35E-05 | 0,392919314 | 1 | 0,732 | 0,689539659 | 3 | Igfbp3 |
| Chst15 | 4,37E-05 | 0,42248181 | 0,483 | 0,226 | 0,692850343 | 3 | Chst15 |
| Col18a11 | 4,40E-05 | 0,575193072 | 0,81 | 0,547 | 0,697529911 | 3 | Col18a1 |
| Map4k51 | 4,42E-05 | 0,43126603 | 0,707 | 0,438 | 0,70088377 | 3 | Map4k5 |
| C77080 | 4,71E-05 | 0,346218342 | 0,448 | 0,211 | 0,746629243 | 3 | C77080 |
| Kdelr31 | 4,75E-05 | 0,475691384 | 0,414 | 0,174 | 0,753149499 | 3 | Kdelr3 |
| Plk21 | 5,10E-05 | 0,46790768 | 0,931 | 0,698 | 0,808948698 | 3 | Plk2 |
| Fam43a1 | 5,38E-05 | 0,375549168 | 0,655 | 0,389 | 0,852291522 | 3 | Fam43a |
| Chst2 | 5,39E-05 | 0,413297352 | 0,431 | 0,204 | 0,854948742 | 3 | Chst2 |
| Akap121 | 5,61E-05 | 0,336329347 | 0,586 | 0,313 | 0,888603297 | 3 | Akap12 |
| Itgb31 | 5,81E-05 | 0,503568213 | 0,586 | 0,325 | 0,921506972 | 3 | Itgb3 |
| Micall2 | 6,14E-05 | 0,309037834 | 0,517 | 0,268 | 0,97299287 | 3 | Micall2 |
| Prss232 | 6,38E-05 | 0,527591969 | 0,845 | 0,57 | 1 | 3 | Prss23 |
| Timp32 | 6,44E-05 | 0,313600744 | 0,879 | 0,6 | 1 | 3 | Timp3 |
| S100a161 | 6,59E-05 | 0,381437219 | 0,931 | 0,623 | 1 | 3 | S100a16 |
| Sh2d51 | 6,61E-05 | 0,446086491 | 0,362 | 0,147 | 1 | 3 | Sh2d5 |
| Arhgap291 | 7,02E-05 | 0,428095998 | 0,776 | 0,509 | 1 | 3 | Arhgap29 |
| Cda | 7,30E-05 | 0,445817914 | 0,431 | 0,226 | 1 | 3 | Cda |
| Mast41 | 7,69E-05 | 0,424376561 | 0,759 | 0,494 | 1 | 3 | Mast4 |
| Adamts41 | 8,12E-05 | 0,294134862 | 0,672 | 0,389 | 1 | 3 | Adamts4 |
| Notch4 | 8,41E-05 | 0,491441442 | 0,655 | 0,423 | 1 | 3 | Notch4 |
| Tmem2 | 9,05E-05 | 0,297996431 | 0,759 | 0,483 | 1 | 3 | Tmem2 |
| Dbn11 | 9,44E-05 | 0,445566515 | 0,707 | 0,472 | 1 | 3 | Dbn1 |
| Bnip31 | 9,47E-05 | 0,797228542 | 0,379 | 0,181 | 1 | 3 | Bnip3 |
| Plxna21 | 9,72E-05 | 0,348934404 | 0,621 | 0,343 | 1 | 3 | Plxna2 |
| Tm9sf3 | 0,000100406 | 0,536085472 | 0,776 | 0,649 | 1 | 3 | Tm9sf3 |
| Ctnna11 | 0,000104287 | 0,357271233 | 0,948 | 0,77 | 1 | 3 | Ctnna1 |
| Gpr41 | 0,000105723 | 0,250781709 | 0,845 | 0,555 | 1 | 3 | Gpr4 |
| Jup2 | 0,00010842 | 0,414918748 | 0,914 | 0,694 | 1 | 3 | Jup |
| Rhoc1 | 0,000112437 | 0,295921044 | 0,948 | 0,683 | 1 | 3 | Rhoc |
| Nrep1 | 0,000121754 | 0,375001321 | 0,672 | 0,475 | 1 | 3 | Nrep |
| Chsy1 | 0,000123329 | 0,299096569 | 0,552 | 0,302 | 1 | 3 | Chsy1 |
| Pgk1 | 0,000127007 | 0,551791572 | 0,845 | 0,687 | 1 | 3 | Pgk1 |
| Ppp1r13b1 | 0,00012809 | 0,305353374 | 0,638 | 0,381 | 1 | 3 | Ppp1r13b |
| Itga1 | 0,000128243 | 0,30881509 | 0,5 | 0,253 | 1 | 3 | Itga1 |
| Reep1 | 0,000129011 | 0,306825567 | 0,379 | 0,158 | 1 | 3 | Reep1 |
| Rgs3 | 0,000133909 | 0,426540657 | 0,586 | 0,336 | 1 | 3 | Rgs3 |
| Myo1d | 0,000139647 | 0,452280739 | 0,379 | 0,166 | 1 | 3 | Myo1d |
| Amotl1 | 0,000140639 | 0,435119742 | 0,707 | 0,487 | 1 | 3 | Amotl1 |
| Flnb1 | 0,000149751 | 0,385152641 | 0,776 | 0,494 | 1 | 3 | Flnb |
| Nectin21 | 0,000154654 | 0,430202506 | 0,621 | 0,4 | 1 | 3 | Nectin2 |
| Icam22 | 0,000158273 | 0,401703475 | 0,862 | 0,6 | 1 | 3 | Icam2 |
| Tuba1a1 | 0,000169744 | 0,403501793 | 0,914 | 0,679 | 1 | 3 | Tuba1a |
| Gas7 | 0,000171968 | 0,421004561 | 0,414 | 0,196 | 1 | 3 | Gas7 |
| Coro1c | 0,000173984 | 0,408631047 | 0,862 | 0,721 | 1 | 3 | Coro1c |
| Itgb11 | 0,000174651 | 0,31365646 | 0,983 | 0,932 | 1 | 3 | Itgb1 |
| AC107851.2 | 0,000183049 | 0,295286361 | 0,259 | 0,087 | 1 | 3 | AC107851.2 |
| Tmem47 | 0,000185825 | 0,29575561 | 0,483 | 0,23 | 1 | 3 | Tmem47 |
| Pcdhgc41 | 0,00019611 | 0,416164948 | 0,655 | 0,419 | 1 | 3 | Pcdhgc4 |
| Plau | 0,00021215 | 0,256471971 | 0,5 | 0,268 | 1 | 3 | Plau |
| Bsg | 0,000231421 | 0,304444128 | 0,983 | 0,917 | 1 | 3 | Bsg |
| Tshz21 | 0,000236386 | 0,316968992 | 0,655 | 0,4 | 1 | 3 | Tshz2 |
| Schip11 | 0,000239855 | 0,304511868 | 0,776 | 0,517 | 1 | 3 | Schip1 |
| Myo1b2 | 0,000277628 | 0,274669282 | 0,793 | 0,517 | 1 | 3 | Myo1b |
| Tmem2041 | 0,000286395 | 0,419805915 | 0,776 | 0,502 | 1 | 3 | Tmem204 |
| Nox41 | 0,000296618 | 0,29943725 | 0,345 | 0,147 | 1 | 3 | Nox4 |
| Sept2 | 0,00029712 | 0,377572936 | 1 | 0,951 | 1 | 3 | Sept2 |
| Sept11 | 0,00029906 | 0,388273735 | 0,741 | 0,525 | 1 | 3 | Sept11 |
| Mpzl11 | 0,000314681 | 0,450637915 | 0,776 | 0,547 | 1 | 3 | Mpzl1 |
| Sacs | 0,000316536 | 0,38574343 | 0,293 | 0,117 | 1 | 3 | Sacs |
| Fermt21 | 0,000317809 | 0,279084842 | 0,845 | 0,551 | 1 | 3 | Fermt2 |
| Cald12 | 0,000320291 | 0,360772912 | 0,828 | 0,619 | 1 | 3 | Cald1 |
| S100a61 | 0,000330296 | 0,306424646 | 0,948 | 0,796 | 1 | 3 | S100a6 |
| Gm4202 | 0,000337858 | 0,281781909 | 0,966 | 0,83 | 1 | 3 | Gm4202 |
| Fam212a | 0,000340777 | 0,294926903 | 0,517 | 0,279 | 1 | 3 | Fam212a |
| Card10 | 0,000342272 | 0,307244952 | 0,431 | 0,204 | 1 | 3 | Card10 |
| Calu1 | 0,000343173 | 0,417823106 | 0,862 | 0,732 | 1 | 3 | Calu |
| Map1b1 | 0,000349102 | 0,522196968 | 0,638 | 0,415 | 1 | 3 | Map1b |
| Snrk1 | 0,000350288 | 0,385360136 | 0,862 | 0,649 | 1 | 3 | Snrk |
| Emp11 | 0,000377214 | 0,347787377 | 0,879 | 0,725 | 1 | 3 | Emp1 |
| Farp1 | 0,000383477 | 0,280904517 | 0,293 | 0,117 | 1 | 3 | Farp1 |
| Pja2 | 0,000384205 | 0,389043706 | 0,603 | 0,362 | 1 | 3 | Pja2 |
| Klk8 | 0,000400105 | 0,343688866 | 0,379 | 0,177 | 1 | 3 | Klk8 |
| Pxdc1 | 0,000417009 | 0,351687886 | 0,345 | 0,162 | 1 | 3 | Pxdc1 |
| Slc16a3 | 0,000427246 | 0,645245654 | 0,431 | 0,245 | 1 | 3 | Slc16a3 |
| Tspan91 | 0,000428171 | 0,381794086 | 0,741 | 0,525 | 1 | 3 | Tspan9 |
| Prnp1 | 0,000429409 | 0,420991842 | 0,845 | 0,604 | 1 | 3 | Prnp |
| Rftn2 | 0,000455553 | 0,293200794 | 0,259 | 0,091 | 1 | 3 | Rftn2 |
| Smad11 | 0,000458816 | 0,251622813 | 0,81 | 0,547 | 1 | 3 | Smad1 |
| Cnn32 | 0,000472289 | 0,356296727 | 0,966 | 0,687 | 1 | 3 | Cnn3 |
| Tgfb1i11 | 0,00047717 | 0,453044359 | 0,552 | 0,325 | 1 | 3 | Tgfb1i1 |
| Sox182 | 0,000479187 | 0,317646262 | 0,81 | 0,623 | 1 | 3 | Sox18 |
| Ctnnb11 | 0,000503652 | 0,404104588 | 0,879 | 0,819 | 1 | 3 | Ctnnb1 |
| F11r1 | 0,000516095 | 0,364828327 | 0,948 | 0,868 | 1 | 3 | F11r |
| Tspan61 | 0,000521663 | 0,476449465 | 0,517 | 0,302 | 1 | 3 | Tspan6 |
| Csgalnact1 | 0,000548811 | 0,251860579 | 0,379 | 0,17 | 1 | 3 | Csgalnact1 |
| Prkar1a1 | 0,000582908 | 0,27824844 | 0,948 | 0,887 | 1 | 3 | Prkar1a |
| Clec1a1 | 0,0006165 | 0,35420117 | 0,621 | 0,355 | 1 | 3 | Clec1a |
| Wscd1 | 0,000616662 | 0,43404773 | 0,534 | 0,313 | 1 | 3 | Wscd1 |
| Ccdc85b1 | 0,00062958 | 0,283114377 | 0,741 | 0,517 | 1 | 3 | Ccdc85b |
| Plxnd11 | 0,000637811 | 0,361415617 | 0,914 | 0,811 | 1 | 3 | Plxnd1 |
| Tgfbr2 | 0,000645221 | 0,422105299 | 0,776 | 0,679 | 1 | 3 | Tgfbr2 |
| Plp2 | 0,000695493 | 0,375752147 | 0,759 | 0,596 | 1 | 3 | Plp2 |
| Itga51 | 0,000699346 | 0,368560739 | 0,776 | 0,596 | 1 | 3 | Itga5 |
| Slc38a21 | 0,00073758 | 0,42363948 | 0,793 | 0,623 | 1 | 3 | Slc38a2 |
| Ets11 | 0,000744706 | 0,371682076 | 0,948 | 0,687 | 1 | 3 | Ets1 |
| Adam10 | 0,000755681 | 0,403552887 | 0,948 | 0,917 | 1 | 3 | Adam10 |
| Cd932 | 0,000764717 | 0,3031472 | 0,983 | 0,917 | 1 | 3 | Cd93 |
| AA467197 | 0,000767421 | 0,285603515 | 0,741 | 0,558 | 1 | 3 | AA467197 |
| Nrarp | 0,000768973 | 0,377776744 | 0,397 | 0,204 | 1 | 3 | Nrarp |
| Tnfrsf22 | 0,000781165 | 0,340240703 | 0,603 | 0,404 | 1 | 3 | Tnfrsf22 |
| Cracr2b | 0,00078593 | 0,39627664 | 0,69 | 0,472 | 1 | 3 | Cracr2b |
| Wbp1l | 0,000793379 | 0,448849342 | 0,534 | 0,313 | 1 | 3 | Wbp1l |
| Ppic2 | 0,000803972 | 0,338534532 | 0,931 | 0,675 | 1 | 3 | Ppic |
| Cdc42ep11 | 0,000820696 | 0,425410393 | 0,603 | 0,426 | 1 | 3 | Cdc42ep1 |
| Arhgap181 | 0,000834732 | 0,428705775 | 0,621 | 0,419 | 1 | 3 | Arhgap18 |
| Cd300lg1 | 0,000852077 | 0,373589175 | 0,448 | 0,238 | 1 | 3 | Cd300lg |
| Prex21 | 0,000856792 | 0,432568703 | 0,914 | 0,808 | 1 | 3 | Prex2 |
| Actn41 | 0,000863303 | 0,374886788 | 0,897 | 0,823 | 1 | 3 | Actn4 |
| Dock91 | 0,00086791 | 0,382096454 | 0,793 | 0,551 | 1 | 3 | Dock9 |
| Afap1l11 | 0,000895802 | 0,427358434 | 0,69 | 0,472 | 1 | 3 | Afap1l1 |

| | | | | | | | |
|---|---|---|---|---|---|---|---|
| Pdgfb | 0,000944547 | 0,312291818 | 0,69 | 0,479 | 1 | 3 | Pdgfb |
| Tnfrsf23 | 0,000971934 | 0,265025841 | 0,483 | 0,272 | 1 | 3 | Tnfrsf23 |
| Bmp1 | 0,000974187 | 0,351060694 | 0,5 | 0,275 | 1 | 3 | Bmp1 |
| Ddah1 | 0,000993968 | 0,435381861 | 0,466 | 0,283 | 1 | 3 | Ddah1 |
| Stim2 | 0,001031484 | 0,366436406 | 0,483 | 0,283 | 1 | 3 | Stim2 |
| Egfl72 | 0,001055202 | 0,303468524 | 1 | 0,702 | 1 | 3 | Egfl7 |
| Cab39 | 0,001074917 | 0,364153199 | 0,655 | 0,479 | 1 | 3 | Cab39 |
| Rapgef1 | 0,001083779 | 0,396293802 | 0,707 | 0,54 | 1 | 3 | Rapgef1 |
| Pcdh12 | 0,001098048 | 0,382261641 | 0,431 | 0,23 | 1 | 3 | Pcdh12 |
| Cgnl1 | 0,001101658 | 0,284972747 | 0,483 | 0,275 | 1 | 3 | Cgnl1 |
| Apbb2 | 0,001107668 | 0,350592752 | 0,793 | 0,555 | 1 | 3 | Apbb2 |
| Ctsl | 0,001133371 | 0,379948332 | 0,931 | 0,879 | 1 | 3 | Ctsl |
| Nos31 | 0,001141743 | 0,268868625 | 0,81 | 0,547 | 1 | 3 | Nos3 |
| Eif4ebp11 | 0,001225365 | 0,396275632 | 0,793 | 0,668 | 1 | 3 | Eif4ebp1 |
| Smtn1 | 0,001228176 | 0,339102774 | 0,793 | 0,54 | 1 | 3 | Smtn |
| Tex2 | 0,001242305 | 0,348890717 | 0,362 | 0,177 | 1 | 3 | Tex2 |
| Aldoa1 | 0,001262816 | 0,306311441 | 0,948 | 0,958 | 1 | 3 | Aldoa |
| Ecscr2 | 0,001278057 | 0,368761622 | 0,966 | 0,679 | 1 | 3 | Ecscr |
| Pmepa11 | 0,001326507 | 0,301137701 | 0,724 | 0,558 | 1 | 3 | Pmepa1 |
| Dpysl2 | 0,001450044 | 0,438183477 | 0,897 | 0,804 | 1 | 3 | Dpysl2 |
| Nes1 | 0,001490347 | 0,288065634 | 0,534 | 0,321 | 1 | 3 | Nes |
| Fkbp101 | 0,001503805 | 0,325568774 | 0,69 | 0,445 | 1 | 3 | Fkbp10 |
| Notch11 | 0,001534654 | 0,282516187 | 0,759 | 0,66 | 1 | 3 | Notch1 |
| Bnip3l | 0,001573063 | 0,39230095 | 0,759 | 0,615 | 1 | 3 | Bnip3l |
| Pear1 | 0,001583851 | 0,371427981 | 0,466 | 0,264 | 1 | 3 | Pear1 |
| Tax1bp31 | 0,00162776 | 0,407033562 | 0,828 | 0,725 | 1 | 3 | Tax1bp3 |
| Mpdz | 0,001689672 | 0,289325579 | 0,276 | 0,125 | 1 | 3 | Mpdz |
| Zdhhc20 | 0,001735884 | 0,399607297 | 0,793 | 0,668 | 1 | 3 | Zdhhc20 |
| Nfib | 0,001813381 | 0,344254162 | 0,828 | 0,577 | 1 | 3 | Nfib |
| Tspan14 | 0,001838071 | 0,428666983 | 0,69 | 0,566 | 1 | 3 | Tspan14 |
| Pde4d | 0,001852124 | 0,267564727 | 0,431 | 0,238 | 1 | 3 | Pde4d |
| Ets2 | 0,001860973 | 0,628923639 | 0,793 | 0,668 | 1 | 3 | Ets2 |
| Lmna | 0,001930236 | 0,391904439 | 0,828 | 0,702 | 1 | 3 | Lmna |
| Myadm1 | 0,001945166 | 0,274220634 | 0,897 | 0,74 | 1 | 3 | Myadm |
| Plcb4 | 0,002011766 | 0,293087297 | 0,414 | 0,234 | 1 | 3 | Plcb4 |
| Tes1 | 0,002038068 | 0,269651113 | 0,862 | 0,691 | 1 | 3 | Tes |
| F2rl31 | 0,00217384 | 0,294260582 | 0,293 | 0,132 | 1 | 3 | F2rl3 |
| Bcl2l111 | 0,002209318 | 0,318354077 | 0,414 | 0,234 | 1 | 3 | Bcl2l11 |
| Synpo1 | 0,002255859 | 0,301748759 | 0,707 | 0,464 | 1 | 3 | Synpo |
| Trib2 | 0,002263953 | 0,379135638 | 0,414 | 0,223 | 1 | 3 | Trib2 |
| Elk31 | 0,002313529 | 0,308299249 | 0,914 | 0,8 | 1 | 3 | Elk3 |
| Kif5b | 0,002348049 | 0,369527302 | 0,862 | 0,74 | 1 | 3 | Kif5b |
| Lxn1 | 0,002362752 | 0,278122501 | 0,776 | 0,608 | 1 | 3 | Lxn |
| Tet3 | 0,00236872 | 0,350991983 | 0,466 | 0,298 | 1 | 3 | Tet3 |
| Lrrc8c1 | 0,002424881 | 0,27194102 | 0,828 | 0,57 | 1 | 3 | Lrrc8c |
| Map4 | 0,002490074 | 0,30531265 | 0,707 | 0,608 | 1 | 3 | Map4 |
| Sh3tc1 | 0,002578883 | 0,291163494 | 0,414 | 0,226 | 1 | 3 | Sh3tc1 |
| Rnd3 | 0,002592621 | 0,335827562 | 0,31 | 0,151 | 1 | 3 | Rnd3 |
| Bag3 | 0,002748816 | 0,290495021 | 0,517 | 0,355 | 1 | 3 | Bag3 |
| Chst71 | 0,002754489 | 0,540510176 | 0,31 | 0,155 | 1 | 3 | Chst7 |
| Cd1091 | 0,002940072 | 0,336980259 | 0,603 | 0,4 | 1 | 3 | Cd109 |
| Rapgef5 | 0,002971746 | 0,390590744 | 0,741 | 0,547 | 1 | 3 | Rapgef5 |
| Smad4 | 0,003070981 | 0,352269932 | 0,414 | 0,242 | 1 | 3 | Smad4 |
| Eno11 | 0,0031242 | 0,354367117 | 0,966 | 0,928 | 1 | 3 | Eno1 |
| Sh3bp5 | 0,003188319 | 0,299779994 | 0,621 | 0,4 | 1 | 3 | Sh3bp5 |
| Plat1 | 0,00319656 | 0,508113893 | 0,569 | 0,374 | 1 | 3 | Plat |
| Fzd41 | 0,003246114 | 0,36272458 | 0,655 | 0,464 | 1 | 3 | Fzd4 |
| Ptp4a31 | 0,003380654 | 0,404581999 | 0,707 | 0,592 | 1 | 3 | Ptp4a3 |
| Srp54a | 0,003459275 | 0,297699677 | 0,724 | 0,608 | 1 | 3 | Srp54a |
| Igfbp41 | 0,003482129 | 0,427392857 | 0,828 | 0,774 | 1 | 3 | Igfbp4 |
| Hilpda | 0,003564763 | 0,479749658 | 0,466 | 0,309 | 1 | 3 | Hilpda |
| Fam102b | 0,003828728 | 0,318660427 | 0,362 | 0,189 | 1 | 3 | Fam102b |
| Sept9 | 0,003876296 | 0,50169177 | 0,724 | 0,687 | 1 | 3 | Sept9 |
| Sept7 | 0,004060394 | 0,276601652 | 0,914 | 0,774 | 1 | 3 | Sept7 |
| Tnfaip8l11 | 0,004072692 | 0,303362059 | 0,517 | 0,309 | 1 | 3 | Tnfaip8l1 |
| Eif2s3x | 0,004131063 | 0,393082136 | 0,569 | 0,438 | 1 | 3 | Eif2s3x |
| Abhd17b | 0,004167875 | 0,315320661 | 0,414 | 0,238 | 1 | 3 | Abhd17b |
| Tmed3 | 0,004190276 | 0,354403726 | 0,569 | 0,411 | 1 | 3 | Tmed3 |
| Spry4 | 0,004228872 | 0,366766871 | 0,431 | 0,257 | 1 | 3 | Spry4 |
| Cds2 | 0,004230396 | 0,25458473 | 0,621 | 0,438 | 1 | 3 | Cds2 |
| Arhgap311 | 0,004367297 | 0,278810944 | 0,759 | 0,574 | 1 | 3 | Arhgap31 |
| Dock61 | 0,004402664 | 0,314318685 | 0,586 | 0,377 | 1 | 3 | Dock6 |
| Nckap1 | 0,004606661 | 0,330338966 | 0,621 | 0,411 | 1 | 3 | Nckap1 |
| 2810403A07Rik | 0,004719022 | 0,256123028 | 0,741 | 0,543 | 1 | 3 | 2810403A07Rik |
| Fam63b | 0,004769266 | 0,252595345 | 0,466 | 0,294 | 1 | 3 | Fam63b |
| Nol4l | 0,004857328 | 0,489743887 | 0,466 | 0,325 | 1 | 3 | Nol4l |
| Gnb11 | 0,004899326 | 0,264829844 | 1 | 0,996 | 1 | 3 | Gnb1 |
| Zfc3h1 | 0,004907091 | 0,287839724 | 0,397 | 0,238 | 1 | 3 | Zfc3h1 |
| Higd1a | 0,004941721 | 0,367323532 | 0,759 | 0,649 | 1 | 3 | Higd1a |
| Tns1 | 0,004980424 | 0,292436637 | 0,638 | 0,445 | 1 | 3 | Tns1 |
| Slc23a2 | 0,005060104 | 0,319457436 | 0,517 | 0,343 | 1 | 3 | Slc23a2 |
| Sh2d3c1 | 0,00516631 | 0,338882456 | 0,793 | 0,574 | 1 | 3 | Sh2d3c |
| Atp2b4 | 0,005185612 | 0,270015404 | 0,362 | 0,223 | 1 | 3 | Atp2b4 |
| Magi1 | 0,005295225 | 0,337581505 | 0,379 | 0,226 | 1 | 3 | Magi1 |
| S100a10 | 0,005337662 | 0,349838029 | 0,845 | 0,74 | 1 | 3 | S100a10 |
| Endod1 | 0,005411465 | 0,315389151 | 0,707 | 0,528 | 1 | 3 | Endod1 |
| Grhpr | 0,005450773 | 0,34089466 | 0,276 | 0,136 | 1 | 3 | Grhpr |
| Ankrd46 | 0,005759027 | 0,336810756 | 0,362 | 0,204 | 1 | 3 | Ankrd46 |
| Myzap | 0,005805447 | 0,327854074 | 0,534 | 0,362 | 1 | 3 | Myzap |
| Sypl | 0,006162549 | 0,312606564 | 0,845 | 0,634 | 1 | 3 | Sypl |
| Sec31a | 0,00617988 | 0,256740923 | 0,672 | 0,494 | 1 | 3 | Sec31a |
| Mgll | 0,006417103 | 0,276450013 | 0,345 | 0,185 | 1 | 3 | Mgll |
| Lnx2 | 0,006487352 | 0,278718525 | 0,345 | 0,196 | 1 | 3 | Lnx2 |
| Dock1 | 0,006761869 | 0,36372258 | 0,621 | 0,494 | 1 | 3 | Dock1 |
| St8sia4 | 0,006938461 | 0,407663635 | 0,483 | 0,34 | 1 | 3 | St8sia4 |
| Pink1 | 0,00700386 | 0,290357012 | 0,534 | 0,404 | 1 | 3 | Pink1 |
| Apaf1 | 0,007198136 | 0,271328609 | 0,397 | 0,234 | 1 | 3 | Apaf1 |
| Taok1 | 0,007535507 | 0,381840199 | 0,655 | 0,475 | 1 | 3 | Taok1 |
| Dnajc18 | 0,00765824 | 0,272186117 | 0,362 | 0,204 | 1 | 3 | Dnajc18 |
| Ywhah | 0,007690824 | 0,373322975 | 0,897 | 0,875 | 1 | 3 | Ywhah |
| Spry1 | 0,007745054 | 0,323670878 | 0,362 | 0,208 | 1 | 3 | Spry1 |
| Asap1 | 0,007810466 | 0,359740282 | 0,569 | 0,434 | 1 | 3 | Asap1 |
| Stx2 | 0,007858203 | 0,434358597 | 0,552 | 0,426 | 1 | 3 | Stx2 |
| Limch11 | 0,008196317 | 0,308569088 | 0,431 | 0,272 | 1 | 3 | Limch1 |
| Slc35g2 | 0,008244252 | 0,305152035 | 0,31 | 0,158 | 1 | 3 | Slc35g2 |
| Gng12 | 0,008536954 | 0,321492636 | 0,741 | 0,596 | 1 | 3 | Gng12 |
| Hk1 | 0,008603847 | 0,288725874 | 0,552 | 0,392 | 1 | 3 | Hk1 |
| Clic41 | 0,008854345 | 0,280210938 | 0,897 | 0,842 | 1 | 3 | Clic4 |
| Ddah21 | 0,009343444 | 0,270194143 | 0,569 | 0,43 | 1 | 3 | Ddah2 |
| CT010467.1 | 0,009814881 | 0,282403316 | 1 | 1 | 1 | 3 | CT010467.1 |
| Tgoln1 | 0,009841817 | 0,363091357 | 0,621 | 0,491 | 1 | 3 | Tgoln1 |
| Adgrg3 | 0,009901968 | 0,456801486 | 0,345 | 0,189 | 1 | 3 | Adgrg3 |
| Pbk | 3,21E-37 | 1,269710446 | 0,808 | 0,037 | 5,09E-33 | 4 | Pbk |
| Ube2c | 1,12E-33 | 1,525661484 | 0,846 | 0,057 | 1,77E-29 | 4 | Ube2c |
| Kif4 | 4,75E-33 | 0,832240641 | 0,654 | 0,02 | 7,53E-29 | 4 | Kif4 |
| Bub1 | 1,02E-31 | 1,01445025 | 0,692 | 0,03 | 1,61E-27 | 4 | Bub1 |
| Prc1 | 1,89E-31 | 1,38620898 | 0,769 | 0,047 | 2,99E-27 | 4 | Prc1 |
| Hmmr | 1,31E-29 | 0,932048556 | 0,615 | 0,024 | 2,07E-25 | 4 | Hmmr |
| Fam64a | 6,72E-29 | 1,002942037 | 0,654 | 0,034 | 1,07E-24 | 4 | Fam64a |
| Top2a | 8,27E-29 | 1,667162445 | 0,846 | 0,077 | 1,31E-24 | 4 | Top2a |
| Cep55 | 1,07E-28 | 1,001202466 | 0,654 | 0,034 | 1,69E-24 | 4 | Cep55 |
| Kif22 | 2,80E-28 | 1,452274372 | 0,808 | 0,074 | 4,43E-24 | 4 | Kif22 |

| | | | | | | | |
|---|---|---|---|---|---|---|---|
| Birc5 | 3,40E-28 | 1,478725718 | 0,962 | 0,125 | 5,38E-24 | 4 | Birc5 |
| Ccna2 | 4,26E-28 | 1,423895551 | 0,846 | 0,084 | 6,76E-24 | 4 | Ccna2 |
| Bub1b | 4,79E-28 | 1,01170232 | 0,692 | 0,04 | 7,59E-24 | 4 | Bub1b |
| Cdca8 | 9,49E-28 | 1,061920837 | 0,808 | 0,071 | 1,50E-23 | 4 | Cdca8 |
| Sgol1 | 1,22E-25 | 0,406791901 | 0,538 | 0,02 | 1,94E-21 | 4 | Sgol1 |
| Foxm1 | 1,64E-25 | 0,36970371 | 0,731 | 0,054 | 2,59E-21 | 4 | Foxm1 |
| Ska1 | 1,84E-25 | 0,710366291 | 0,577 | 0,027 | 2,91E-21 | 4 | Ska1 |
| Nusap1 | 2,54E-24 | 1,250959781 | 0,692 | 0,061 | 4,03E-20 | 4 | Nusap1 |
| Pclaf | 5,46E-24 | 1,283233822 | 0,846 | 0,104 | 8,66E-20 | 4 | Pclaf |
| Gtse1 | 5,56E-24 | 0,332274009 | 0,615 | 0,037 | 8,81E-20 | 4 | Gtse1 |
| Ncapd2 | 6,06E-24 | 0,903172408 | 0,846 | 0,104 | 9,61E-20 | 4 | Ncapd2 |
| Smc2 | 9,01E-24 | 1,392430413 | 0,962 | 0,155 | 1,43E-19 | 4 | Smc2 |
| Esco2 | 2,23E-23 | 0,435580149 | 0,5 | 0,02 | 3,53E-19 | 4 | Esco2 |
| Mki67 | 3,84E-23 | 1,551404858 | 0,846 | 0,111 | 6,09E-19 | 4 | Mki67 |
| Cdk1 | 6,79E-23 | 1,643562807 | 0,923 | 0,158 | 1,08E-18 | 4 | Cdk1 |
| Mastl | 2,88E-22 | 0,634122042 | 0,577 | 0,04 | 4,56E-18 | 4 | Mastl |
| Ccnb1 | 7,19E-22 | 1,268324015 | 0,654 | 0,057 | 1,14E-17 | 4 | Ccnb1 |
| Rad51 | 8,62E-22 | 0,809118367 | 0,615 | 0,047 | 1,37E-17 | 4 | Rad51 |
| Hist1h1b | 8,65E-22 | 0,5182486 | 0,538 | 0,034 | 1,37E-17 | 4 | Hist1h1b |
| Ckap2l | 1,26E-21 | 0,883147164 | 0,654 | 0,057 | 1,99E-17 | 4 | Ckap2l |
| Cdc20 | 1,36E-21 | 1,445883981 | 0,692 | 0,071 | 2,15E-17 | 4 | Cdc20 |
| Cenpq | 2,75E-21 | 0,940303482 | 0,692 | 0,071 | 4,36E-17 | 4 | Cenpq |
| Racgap1 | 4,82E-21 | 1,151625755 | 0,769 | 0,101 | 7,64E-17 | 4 | Racgap1 |
| Cenpe | 5,24E-21 | 0,930164371 | 0,615 | 0,051 | 8,31E-17 | 4 | Cenpe |
| Rrm2 | 6,94E-21 | 1,487023673 | 0,731 | 0,088 | 1,10E-16 | 4 | Rrm2 |
| Gm15452 | 9,24E-21 | 0,304126414 | 0,846 | 0,152 | 1,46E-16 | 4 | Gm15452 |
| Anln | 9,56E-21 | 0,811562746 | 0,692 | 0,074 | 1,51E-16 | 4 | Anln |
| Cdca5 | 1,04E-20 | 0,662803118 | 0,5 | 0,027 | 1,65E-16 | 4 | Cdca5 |
| Tacc3 | 1,11E-20 | 1,028113113 | 0,769 | 0,104 | 1,76E-16 | 4 | Tacc3 |
| Nuf2 | 1,47E-20 | 0,728976062 | 0,538 | 0,037 | 2,34E-16 | 4 | Nuf2 |
| Mis18bp1 | 2,15E-20 | 1,057098114 | 0,5 | 0,03 | 3,41E-16 | 4 | Mis18bp1 |
| Ckap2 | 2,49E-20 | 1,138358823 | 0,692 | 0,081 | 3,95E-16 | 4 | Ckap2 |
| Melk | 3,33E-20 | 0,832611343 | 0,462 | 0,024 | 5,28E-16 | 4 | Melk |
| Kif20b | 3,52E-20 | 0,684130263 | 0,577 | 0,044 | 5,58E-16 | 4 | Kif20b |
| Cenpw | 3,65E-20 | 0,399875956 | 0,885 | 0,141 | 5,79E-16 | 4 | Cenpw |
| Depdc1a | 3,98E-20 | 0,658023772 | 0,423 | 0,017 | 6,31E-16 | 4 | Depdc1a |
| Sapcd2 | 6,83E-20 | 0,421673576 | 0,423 | 0,017 | 1,08E-15 | 4 | Sapcd2 |
| Cks2 | 1,46E-19 | 1,007286954 | 0,885 | 0,205 | 2,31E-15 | 4 | Cks2 |
| Aspm | 2,39E-19 | 0,483630971 | 0,5 | 0,03 | 3,79E-15 | 4 | Aspm |
| Cdca2 | 2,94E-19 | 0,588538092 | 0,538 | 0,04 | 4,66E-15 | 4 | Cdca2 |
| Cenpf | 2,99E-19 | 1,250365416 | 0,615 | 0,067 | 4,73E-15 | 4 | Cenpf |
| Diaph3 | 6,50E-19 | 0,35645805 | 0,615 | 0,061 | 1,03E-14 | 4 | Diaph3 |
| H2afx | 9,18E-19 | 1,310240662 | 0,962 | 0,249 | 1,46E-14 | 4 | H2afx |
| Shcbp1 | 9,57E-19 | 0,923034978 | 0,577 | 0,057 | 1,52E-14 | 4 | Shcbp1 |
| Gm44346 | 1,05E-18 | 0,547028278 | 0,885 | 0,189 | 1,66E-14 | 4 | Gm44346 |
| Neil3 | 1,13E-18 | 0,717101947 | 0,5 | 0,037 | 1,79E-14 | 4 | Neil3 |
| Hist2h3c2 | 1,18E-18 | 0,265532945 | 0,5 | 0,037 | 1,87E-14 | 4 | Hist2h3c2 |
| Kif15 | 1,27E-18 | 0,636665048 | 0,538 | 0,044 | 2,02E-14 | 4 | Kif15 |
| Tk1 | 1,50E-18 | 1,141687722 | 0,846 | 0,155 | 2,37E-14 | 4 | Tk1 |
| Kif20a | 1,64E-18 | 0,90245883 | 0,692 | 0,091 | 2,60E-14 | 4 | Kif20a |
| Cdca31 | 1,97E-18 | 0,66965847 | 0,769 | 0,101 | 3,12E-14 | 4 | Cdca3 |
| Spc25 | 2,05E-18 | 0,978054555 | 0,692 | 0,098 | 3,25E-14 | 4 | Spc25 |
| Hist1h2ae | 3,34E-18 | 0,622840972 | 0,538 | 0,047 | 5,30E-14 | 4 | Hist1h2ae |
| Plk1 | 3,59E-18 | 0,944143814 | 0,5 | 0,037 | 5,69E-14 | 4 | Plk1 |
| Hist1h2ap | 8,53E-18 | 2,399714515 | 0,769 | 0,141 | 1,35E-13 | 4 | Hist1h2ap |
| Prr11 | 8,60E-18 | 0,330938653 | 0,423 | 0,024 | 1,36E-13 | 4 | Prr11 |
| Kif11 | 9,86E-18 | 0,758370748 | 0,654 | 0,084 | 1,56E-13 | 4 | Kif11 |
| Kif23 | 1,23E-17 | 0,96629397 | 0,577 | 0,061 | 1,95E-13 | 4 | Kif23 |
| Gen1 | 1,26E-17 | 0,570386356 | 0,423 | 0,024 | 2,00E-13 | 4 | Gen1 |
| Ube2t | 1,85E-17 | 0,603181895 | 0,423 | 0,024 | 2,93E-13 | 4 | Ube2t |
| Plk4 | 3,93E-17 | 0,828775251 | 0,692 | 0,101 | 6,22E-13 | 4 | Plk4 |
| Stmn1 | 4,41E-17 | 1,469048631 | 1 | 0,38 | 6,99E-13 | 4 | Stmn1 |
| Mxd3 | 5,43E-17 | 0,66924051 | 0,731 | 0,141 | 8,60E-13 | 4 | Mxd3 |
| Ect2 | 7,08E-17 | 0,606030735 | 0,423 | 0,027 | 1,12E-12 | 4 | Ect2 |
| Asf1b | 7,95E-17 | 0,694388822 | 0,654 | 0,084 | 1,26E-12 | 4 | Asf1b |
| Cdc25c | 1,33E-16 | 0,303139035 | 0,346 | 0,013 | 2,11E-12 | 4 | Cdc25c |
| Knl1 | 1,63E-16 | 0,490644218 | 0,5 | 0,044 | 2,58E-12 | 4 | Knl1 |
| Fam83d | 1,65E-16 | 0,649122812 | 0,346 | 0,013 | 2,61E-12 | 4 | Fam83d |
| Spag5 | 1,67E-16 | 0,304234281 | 0,423 | 0,027 | 2,65E-12 | 4 | Spag5 |
| Iqgap3 | 2,01E-16 | 0,715603075 | 0,615 | 0,077 | 3,18E-12 | 4 | Iqgap3 |
| Ccnb2 | 2,42E-16 | 1,167869659 | 0,769 | 0,138 | 3,84E-12 | 4 | Ccnb2 |
| Tmpo | 2,91E-16 | 1,294744956 | 0,923 | 0,246 | 4,61E-12 | 4 | Tmpo |
| Knstrn | 3,81E-16 | 0,727992106 | 0,615 | 0,077 | 6,05E-12 | 4 | Knstrn |
| Hist2h3b | 4,44E-16 | 0,505354075 | 0,538 | 0,061 | 7,03E-12 | 4 | Hist2h3b |
| Tyms | 9,19E-16 | 1,228054569 | 0,923 | 0,279 | 1,46E-11 | 4 | Tyms |
| Tpx2 | 1,02E-15 | 0,95568362 | 0,692 | 0,114 | 1,61E-11 | 4 | Tpx2 |
| Nek2 | 1,11E-15 | 0,723696851 | 0,423 | 0,03 | 1,77E-11 | 4 | Nek2 |
| Ncaph | 2,11E-15 | 0,920147008 | 0,692 | 0,111 | 3,35E-11 | 4 | Ncaph |
| Aurkb | 2,88E-15 | 0,722936134 | 0,5 | 0,051 | 4,57E-11 | 4 | Aurkb |
| Cenpa | 3,60E-15 | 1,13029627 | 0,731 | 0,148 | 5,71E-11 | 4 | Cenpa |
| Ndc80 | 4,18E-15 | 0,604995604 | 0,577 | 0,074 | 6,62E-11 | 4 | Ndc80 |
| AC153546.1 | 6,26E-15 | 1,108688914 | 1 | 0,411 | 9,93E-11 | 4 | AC153546.1 |
| Tuba1b | 7,32E-15 | 1,483416472 | 1 | 0,825 | 1,16E-10 | 4 | Tuba1b |
| Aurka | 1,11E-14 | 0,911676968 | 0,462 | 0,044 | 1,77E-10 | 4 | Aurka |
| Uhrf1 | 1,58E-14 | 0,631182053 | 0,731 | 0,135 | 2,51E-10 | 4 | Uhrf1 |
| Trim59 | 1,90E-14 | 0,610409404 | 0,538 | 0,071 | 3,01E-10 | 4 | Trim59 |
| Ncapg21 | 1,95E-14 | 0,485939196 | 0,769 | 0,138 | 3,09E-10 | 4 | Ncapg2 |
| C330027C09Rik | 2,01E-14 | 0,483674735 | 0,423 | 0,037 | 3,19E-10 | 4 | C330027C09Rik |
| Ska3 | 2,64E-14 | 0,723338712 | 0,423 | 0,037 | 4,19E-10 | 4 | Ska3 |
| Smc4 | 2,85E-14 | 0,92828622 | 0,962 | 0,303 | 4,51E-10 | 4 | Smc4 |
| Sgol2a | 3,35E-14 | 0,674300786 | 0,538 | 0,071 | 5,31E-10 | 4 | Sgol2a |
| 2700099C18Rik | 4,37E-14 | 0,386862666 | 0,423 | 0,037 | 6,92E-10 | 4 | 2700099C18Rik |
| Cenpl | 5,50E-14 | 0,546665018 | 0,423 | 0,037 | 8,72E-10 | 4 | Cenpl |
| Phgdh | 5,81E-14 | 1,044926619 | 0,462 | 0,051 | 9,21E-10 | 4 | Phgdh |
| Hmgb2 | 6,07E-14 | 1,166247407 | 1 | 0,481 | 9,63E-10 | 4 | Hmgb2 |
| Stil | 8,16E-14 | 0,419650351 | 0,385 | 0,03 | 1,29E-09 | 4 | Stil |
| Tubb5 | 1,18E-13 | 1,243625236 | 1 | 0,963 | 1,87E-09 | 4 | Tubb5 |
| Kif2c | 1,84E-13 | 0,641907486 | 0,346 | 0,024 | 2,92E-09 | 4 | Kif2c |
| Hmgn2 | 1,93E-13 | 1,340366955 | 1 | 0,751 | 3,06E-09 | 4 | Hmgn2 |
| Pif1 | 2,04E-13 | 0,37839855 | 0,269 | 0,01 | 3,23E-09 | 4 | Pif1 |
| Cenpm | 2,19E-13 | 0,453523313 | 0,538 | 0,071 | 3,47E-09 | 4 | Cenpm |
| H2afz | 6,09E-13 | 1,218802222 | 1 | 0,862 | 9,66E-09 | 4 | H2afz |
| Pkmyt1 | 7,01E-13 | 0,410437798 | 0,577 | 0,088 | 1,11E-08 | 4 | Pkmyt1 |
| Spc24 | 7,39E-13 | 0,832891789 | 0,615 | 0,118 | 1,17E-08 | 4 | Spc24 |
| Traip | 7,97E-13 | 0,280981174 | 0,308 | 0,017 | 1,26E-08 | 4 | Traip |
| Hmgb3 | 1,54E-12 | 0,923106567 | 0,885 | 0,283 | 2,45E-08 | 4 | Hmgb3 |
| Ccdc34 | 2,51E-12 | 0,575517539 | 0,692 | 0,155 | 3,98E-08 | 4 | Ccdc34 |
| Mad2l1 | 3,43E-12 | 0,721640359 | 0,692 | 0,141 | 5,44E-08 | 4 | Mad2l1 |
| 1700066M21Rik | 3,50E-12 | 0,287364314 | 0,346 | 0,027 | 5,56E-08 | 4 | 1700066M21Rik |
| Cenpi | 4,03E-12 | 0,414876992 | 0,385 | 0,037 | 6,39E-08 | 4 | Cenpi |
| Cks1b | 4,95E-12 | 0,930532596 | 0,846 | 0,286 | 7,84E-08 | 4 | Cks1b |
| E2f7 | 5,49E-12 | 0,473927396 | 0,577 | 0,104 | 8,70E-08 | 4 | E2f7 |
| Wdr62 | 7,18E-12 | 0,489063553 | 0,385 | 0,037 | 1,14E-07 | 4 | Wdr62 |
| Cit | 7,26E-12 | 0,307331805 | 0,654 | 0,135 | 1,15E-07 | 4 | Cit |
| Kpna2 | 7,64E-12 | 1,577052896 | 0,808 | 0,276 | 1,21E-07 | 4 | Kpna2 |
| Rrm1 | 9,40E-12 | 0,973829217 | 0,808 | 0,219 | 1,49E-07 | 4 | Rrm1 |
| Usp1 | 1,35E-11 | 0,664820092 | 0,692 | 0,158 | 2,14E-07 | 4 | Usp1 |
| Trip13 | 1,92E-11 | 0,342248934 | 0,385 | 0,04 | 3,04E-07 | 4 | Trip13 |
| Gm15387 | 2,95E-11 | 0,779180568 | 1 | 0,872 | 4,68E-07 | 4 | Gm15387 |
| Zwilch | 3,62E-11 | 0,258900111 | 0,346 | 0,03 | 5,73E-07 | 4 | Zwilch |
| Troap | 4,20E-11 | 0,285653281 | 0,385 | 0,04 | 6,66E-07 | 4 | Troap |

| | | | | | | | |
|---|---|---|---|---|---|---|---|
| Gm10282 | 4,24E-11 | 0,638447306 | 0,962 | 0,613 | 6,72E-07 | 4 | Gm10282 |
| Rangap1 | 4,56E-11 | 0,705105789 | 0,692 | 0,165 | 7,23E-07 | 4 | Rangap1 |
| Incenp1 | 4,83E-11 | 0,331255745 | 0,654 | 0,125 | 7,65E-07 | 4 | Incenp |
| Dek | 6,82E-11 | 1,118746785 | 0,962 | 0,572 | 1,08E-06 | 4 | Dek |
| Cdkn2c | 8,46E-11 | 0,496338592 | 0,538 | 0,094 | 1,34E-06 | 4 | Cdkn2c |
| Lin9 | 1,18E-10 | 0,438373827 | 0,385 | 0,047 | 1,87E-06 | 4 | Lin9 |
| Nucks1 | 1,38E-10 | 0,664779679 | 1 | 0,421 | 2,18E-06 | 4 | Nucks1 |
| Tubb4b | 1,72E-10 | 1,140108372 | 0,962 | 0,532 | 2,73E-06 | 4 | Tubb4b |
| Ttk | 2,47E-10 | 0,398857616 | 0,308 | 0,027 | 3,92E-06 | 4 | Ttk |
| Ccnf | 2,53E-10 | 0,450320176 | 0,346 | 0,037 | 4,00E-06 | 4 | Ccnf |
| Bora | 2,76E-10 | 0,379170224 | 0,5 | 0,084 | 4,37E-06 | 4 | Bora |
| Ckap5 | 2,92E-10 | 0,650632723 | 0,769 | 0,209 | 4,63E-06 | 4 | Ckap5 |
| Ncapg | 3,45E-10 | 0,581105125 | 0,462 | 0,077 | 5,47E-06 | 4 | Ncapg |
| Phf19 | 3,59E-10 | 0,294042319 | 0,269 | 0,02 | 5,69E-06 | 4 | Phf19 |
| Katnbl1 | 3,93E-10 | 0,61488563 | 0,654 | 0,158 | 6,22E-06 | 4 | Katnbl1 |
| Prim1 | 4,00E-10 | 0,496564708 | 0,5 | 0,088 | 6,35E-06 | 4 | Prim1 |
| Cbx5 | 5,16E-10 | 0,726183632 | 0,731 | 0,195 | 8,18E-06 | 4 | Cbx5 |
| Ulbp1 | 5,58E-10 | 0,436158317 | 0,538 | 0,101 | 8,85E-06 | 4 | Ulbp1 |
| Gm10184 | 6,81E-10 | 0,463597121 | 0,538 | 0,118 | 1,08E-05 | 4 | Gm10184 |
| Cdc25b | 8,03E-10 | 0,383233594 | 0,346 | 0,04 | 1,27E-05 | 4 | Cdc25b |
| Mcm2 | 9,71E-10 | 0,845001077 | 0,654 | 0,162 | 1,54E-05 | 4 | Mcm2 |
| Fbxo5 | 1,07E-09 | 0,635077427 | 0,308 | 0,03 | 1,70E-05 | 4 | Fbxo5 |
| Ptma | 1,28E-09 | 0,639702691 | 1 | 0,99 | 2,03E-05 | 4 | Ptma |
| Ska2 | 1,31E-09 | 0,277888466 | 0,538 | 0,098 | 2,07E-05 | 4 | Ska2 |
| Pole | 1,31E-09 | 0,375641398 | 0,385 | 0,051 | 2,07E-05 | 4 | Pole |
| Haus6 | 1,38E-09 | 0,441428499 | 0,462 | 0,077 | 2,19E-05 | 4 | Haus6 |
| Dbf4 | 1,48E-09 | 0,343602866 | 0,615 | 0,138 | 2,34E-05 | 4 | Dbf4 |
| Dnajc9 | 1,50E-09 | 0,702340808 | 0,615 | 0,152 | 2,39E-05 | 4 | Dnajc9 |
| Hmgb1 | 2,03E-09 | 0,71949138 | 1 | 0,953 | 3,22E-05 | 4 | Hmgb1 |
| Rfc5 | 2,63E-09 | 0,704570181 | 0,577 | 0,135 | 4,17E-05 | 4 | Rfc5 |
| Ncapd3 | 3,03E-09 | 0,616941391 | 0,615 | 0,162 | 4,81E-05 | 4 | Ncapd3 |
| Dlgap5 | 3,34E-09 | 0,329967132 | 0,423 | 0,067 | 5,30E-05 | 4 | Dlgap5 |
| Aplf | 3,34E-09 | 0,318709548 | 0,423 | 0,067 | 5,30E-05 | 4 | Aplf |
| Calm2 | 3,61E-09 | 0,794629084 | 1 | 0,956 | 5,72E-05 | 4 | Calm2 |
| Sae1 | 3,89E-09 | 0,913062525 | 0,846 | 0,387 | 6,17E-05 | 4 | Sae1 |
| G2e3 | 4,24E-09 | 0,568742155 | 0,538 | 0,121 | 6,71E-05 | 4 | G2e3 |
| Fam111a | 4,97E-09 | 0,555000718 | 0,692 | 0,195 | 7,87E-05 | 4 | Fam111a |
| Cmc2 | 5,13E-09 | 0,325082507 | 0,577 | 0,135 | 8,14E-05 | 4 | Cmc2 |
| Hist1h4d | 7,99E-09 | 0,429153174 | 0,654 | 0,189 | 0,000126717 | 4 | Hist1h4d |
| Fen1 | 8,00E-09 | 0,669178921 | 0,577 | 0,145 | 0,000126816 | 4 | Fen1 |
| Gm21596 | 8,57E-09 | 0,523905108 | 1 | 0,818 | 0,000135843 | 4 | Gm21596 |
| D030056L22Rik | 9,52E-09 | 0,285061332 | 0,423 | 0,071 | 0,000150869 | 4 | D030056L22Rik |
| Gm10182 | 9,59E-09 | 0,269051803 | 0,846 | 0,505 | 0,000152053 | 4 | Gm10182 |
| Anp32b | 9,93E-09 | 0,723577323 | 1 | 0,562 | 0,000157429 | 4 | Anp32b |
| Tubg1 | 1,18E-08 | 0,821767419 | 0,615 | 0,182 | 0,000187579 | 4 | Tubg1 |
| Cenpn | 1,33E-08 | 0,319375839 | 0,269 | 0,027 | 0,000211051 | 4 | Cenpn |
| Cenph | 1,42E-08 | 0,442774958 | 0,346 | 0,051 | 0,000224658 | 4 | Cenph |
| Lmnb1 | 1,42E-08 | 0,399215627 | 0,423 | 0,074 | 0,000225106 | 4 | Lmnb1 |
| Ncl | 1,59E-08 | 0,828180881 | 1 | 0,828 | 0,000251995 | 4 | Ncl |
| Rfc4 | 1,68E-08 | 0,293665465 | 0,423 | 0,071 | 0,000266531 | 4 | Rfc4 |
| Ercc6l | 2,17E-08 | 0,251016055 | 0,269 | 0,027 | 0,000343933 | 4 | Ercc6l |
| Nup35 | 2,32E-08 | 0,416976506 | 0,462 | 0,094 | 0,00036805 | 4 | Nup35 |
| Ppa1 | 2,46E-08 | 0,535311028 | 0,654 | 0,182 | 0,00039011 | 4 | Ppa1 |
| Haus3 | 2,83E-08 | 0,37309893 | 0,462 | 0,091 | 0,000449116 | 4 | Haus3 |
| Rpa1 | 3,13E-08 | 0,734355602 | 0,731 | 0,232 | 0,000495757 | 4 | Rpa1 |
| Ran | 3,72E-08 | 0,721700226 | 0,962 | 0,801 | 0,000589447 | 4 | Ran |
| Tipin | 3,73E-08 | 0,977828851 | 0,692 | 0,269 | 0,000591178 | 4 | Tipin |
| Cep192 | 4,22E-08 | 0,368462114 | 0,577 | 0,141 | 0,000669087 | 4 | Cep192 |
| Tubb61 | 4,26E-08 | 0,916146969 | 0,962 | 0,721 | 0,000675327 | 4 | Tubb6 |
| Exo1 | 4,59E-08 | 0,39412967 | 0,269 | 0,03 | 0,00072748 | 4 | Exo1 |
| Anp32e | 5,06E-08 | 0,712714411 | 0,731 | 0,32 | 0,000802813 | 4 | Anp32e |
| Slfn9 | 5,54E-08 | 0,416107597 | 0,462 | 0,094 | 0,000878243 | 4 | Slfn9 |
| Tcf19 | 6,05E-08 | 0,748047094 | 0,5 | 0,114 | 0,000958678 | 4 | Tcf19 |
| Tfdp1 | 7,48E-08 | 0,700586407 | 0,769 | 0,296 | 0,001185197 | 4 | Tfdp1 |
| Cdc7 | 7,50E-08 | 0,460521844 | 0,346 | 0,054 | 0,001188398 | 4 | Cdc7 |
| Ccne1 | 7,70E-08 | 0,520768445 | 0,346 | 0,054 | 0,001219993 | 4 | Ccne1 |
| Mybl2 | 1,06E-07 | 0,358220062 | 0,308 | 0,044 | 0,001675062 | 4 | Mybl2 |
| Atad2 | 1,25E-07 | 0,48858619 | 0,538 | 0,128 | 0,001976058 | 4 | Atad2 |
| Uchl3 | 1,58E-07 | 0,57898849 | 0,731 | 0,279 | 0,002512012 | 4 | Uchl3 |
| Lig1 | 1,83E-07 | 0,702644314 | 0,615 | 0,205 | 0,00289698 | 4 | Lig1 |
| Arhgef39 | 1,86E-07 | 0,415187935 | 0,269 | 0,034 | 0,002951399 | 4 | Arhgef39 |
| Gm4617 | 2,53E-07 | 0,283484913 | 1 | 0,909 | 0,004003921 | 4 | Gm4617 |
| Brd8 | 3,18E-07 | 0,448883265 | 0,808 | 0,306 | 0,005041627 | 4 | Brd8 |
| Dnmt1 | 3,21E-07 | 0,535578319 | 0,615 | 0,205 | 0,005091902 | 4 | Dnmt1 |
| Ppia | 3,53E-07 | 0,381666639 | 1 | 1 | 0,005588328 | 4 | Ppia |
| Prim2 | 4,50E-07 | 0,596661314 | 0,423 | 0,098 | 0,007134256 | 4 | Prim2 |
| Pold1 | 4,54E-07 | 0,553074722 | 0,5 | 0,128 | 0,007198786 | 4 | Pold1 |
| Exosc8 | 4,71E-07 | 0,564256909 | 0,769 | 0,283 | 0,007459604 | 4 | Exosc8 |
| Nasp | 4,77E-07 | 0,428814723 | 0,731 | 0,269 | 0,007562497 | 4 | Nasp |
| Rad21 | 4,99E-07 | 0,523340154 | 0,846 | 0,384 | 0,007915625 | 4 | Rad21 |
| Adk | 5,11E-07 | 0,287462961 | 0,462 | 0,111 | 0,008106769 | 4 | Adk |
| Gm8203 | 5,42E-07 | 0,642127713 | 0,962 | 0,744 | 0,008586522 | 4 | Gm8203 |
| Cntln | 8,23E-07 | 0,504729721 | 0,5 | 0,141 | 0,013049529 | 4 | Cntln |
| Psat1 | 9,05E-07 | 0,565452006 | 0,385 | 0,081 | 0,014341541 | 4 | Psat1 |
| Cenpk | 9,59E-07 | 0,314108334 | 0,308 | 0,051 | 0,01519965 | 4 | Cenpk |
| Gm10123 | 1,11E-06 | 0,392234729 | 1 | 0,987 | 0,017533526 | 4 | Gm10123 |
| H1f0 | 1,16E-06 | 0,815045746 | 1 | 0,751 | 0,018453479 | 4 | H1f0 |
| Dut | 1,43E-06 | 0,503888515 | 0,615 | 0,212 | 0,022605642 | 4 | Dut |
| Odf2 | 1,53E-06 | 0,45954781 | 0,577 | 0,172 | 0,024198958 | 4 | Odf2 |
| Bhlhb9 | 1,58E-06 | 0,490160504 | 0,269 | 0,04 | 0,024998711 | 4 | Bhlhb9 |
| Brca2 | 1,62E-06 | 0,311029773 | 0,346 | 0,067 | 0,025614114 | 4 | Brca2 |
| Cenpc1 | 1,80E-06 | 0,322965201 | 0,423 | 0,098 | 0,028526902 | 4 | Cenpc1 |
| Gpsm2 | 1,80E-06 | 0,32091444 | 0,269 | 0,04 | 0,02857664 | 4 | Gpsm2 |
| Set | 1,95E-06 | 0,816804128 | 1 | 0,828 | 0,030919391 | 4 | Set |
| Cdkn2d | 1,96E-06 | 0,58040359 | 0,462 | 0,128 | 0,031014249 | 4 | Cdkn2d |
| 2810025M15Rik | 2,00E-06 | 0,526446039 | 0,846 | 0,424 | 0,031669295 | 4 | 2810025M15Rik |
| Ppih | 2,39E-06 | 0,361961051 | 0,462 | 0,125 | 0,037818898 | 4 | Ppih |
| Mettl14 | 3,07E-06 | 0,257688059 | 0,615 | 0,215 | 0,048644351 | 4 | Mettl14 |
| Myef2 | 3,17E-06 | 0,676979879 | 0,846 | 0,556 | 0,050323348 | 4 | Myef2 |
| Siva11 | 3,32E-06 | 0,350346364 | 0,577 | 0,195 | 0,052577635 | 4 | Siva1 |
| Tinagl1 | 3,43E-06 | 0,414131365 | 0,615 | 0,189 | 0,054432595 | 4 | Tinagl1 |
| Lmnb2 | 3,48E-06 | 0,472239852 | 0,385 | 0,088 | 0,055243977 | 4 | Lmnb2 |
| Dsn1 | 3,62E-06 | 0,342155673 | 0,269 | 0,044 | 0,057408545 | 4 | Dsn1 |
| Rpa3 | 3,65E-06 | 0,340361174 | 0,615 | 0,222 | 0,05793786 | 4 | Rpa3 |
| Rnd1 | 3,69E-06 | 0,308951376 | 0,385 | 0,088 | 0,058550056 | 4 | Rnd1 |
| Cep57 | 4,09E-06 | 0,486050595 | 0,577 | 0,199 | 0,064764541 | 4 | Cep57 |
| Sdpr1 | 4,19E-06 | 0,589268179 | 0,654 | 0,249 | 0,066457352 | 4 | Sdpr |
| Papd7 | 5,07E-06 | 0,339197896 | 0,385 | 0,094 | 0,080303336 | 4 | Papd7 |
| Hells | 5,20E-06 | 0,317289822 | 0,5 | 0,145 | 0,082414181 | 4 | Hells |
| Ube2s | 5,73E-06 | 0,377991188 | 0,769 | 0,327 | 0,090786881 | 4 | Ube2s |
| Rbbp7 | 5,99E-06 | 0,724985184 | 0,808 | 0,569 | 0,094961744 | 4 | Rbbp7 |
| Ppil1 | 6,07E-06 | 0,547546478 | 0,692 | 0,279 | 0,096270965 | 4 | Ppil1 |
| Tuba1c1 | 6,10E-06 | 0,833093829 | 0,962 | 0,623 | 0,096707504 | 4 | Tuba1c |
| Myh10 | 6,27E-06 | 0,521059837 | 0,769 | 0,354 | 0,09935838 | 4 | Myh10 |
| Ssrp1 | 6,37E-06 | 0,566264087 | 0,808 | 0,438 | 0,101032881 | 4 | Ssrp1 |
| Kpnb1 | 6,64E-06 | 0,505049791 | 0,923 | 0,535 | 0,105225033 | 4 | Kpnb1 |
| Slf1 | 6,91E-06 | 0,45842811 | 0,423 | 0,114 | 0,109599114 | 4 | Slf1 |
| E2f8 | 7,33E-06 | 0,583942341 | 0,269 | 0,047 | 0,116148312 | 4 | E2f8 |
| Hn1l | 7,35E-06 | 0,91683137 | 0,769 | 0,468 | 0,116448316 | 4 | Hn1l |
| Nop14 | 7,55E-06 | 0,406773681 | 0,462 | 0,138 | 0,11972018 | 4 | Nop14 |

| | | | | | | | |
|---|---|---|---|---|---|---|---|
| Pfdn1 | 7,89E-06 | 0,498098467 | 0,885 | 0,508 | 0,125055565 | 4 | Pfdn1 |
| Selenoh | 8,38E-06 | 0,562693444 | 0,769 | 0,438 | 0,132865215 | 4 | Selenoh |
| Donson | 9,66E-06 | 0,535580749 | 0,423 | 0,128 | 0,153115175 | 4 | Donson |
| Zfp367 | 1,01E-05 | 0,513407297 | 0,385 | 0,101 | 0,160164565 | 4 | Zfp367 |
| Gm10131 | 1,08E-05 | 0,409412711 | 0,962 | 0,768 | 0,171323289 | 4 | Gm10131 |
| Casp12 | 1,15E-05 | 0,314038287 | 0,5 | 0,145 | 0,18194618 | 4 | Casp12 |
| Hmgn1 | 1,18E-05 | 0,656858087 | 0,923 | 0,667 | 0,187459126 | 4 | Hmgn1 |
| Cdk5rap2 | 1,19E-05 | 0,382279663 | 0,423 | 0,125 | 0,188056088 | 4 | Cdk5rap2 |
| Tjp2 | 1,20E-05 | 0,546958966 | 0,577 | 0,212 | 0,190488284 | 4 | Tjp2 |
| Ppp1cb | 1,23E-05 | 0,587600243 | 0,846 | 0,438 | 0,194645546 | 4 | Ppp1cb |
| Arhgap11a | 1,34E-05 | 0,26297295 | 0,346 | 0,081 | 0,21165257 | 4 | Arhgap11a |
| Cep295 | 1,36E-05 | 0,254078243 | 0,5 | 0,148 | 0,21501547 | 4 | Cep295 |
| Hint1 | 1,40E-05 | 0,430294334 | 1 | 0,848 | 0,222536992 | 4 | Hint1 |
| Cnot9 | 1,43E-05 | 0,61995085 | 0,731 | 0,35 | 0,227301447 | 4 | Cnot9 |
| 4930579G24Rik | 1,48E-05 | 0,281283833 | 0,308 | 0,064 | 0,234003977 | 4 | 4930579G24Rik |
| Impdh2 | 1,48E-05 | 0,555583253 | 0,846 | 0,431 | 0,234860552 | 4 | Impdh2 |
| Ranbp1 | 1,54E-05 | 0,570451777 | 0,962 | 0,731 | 0,243524797 | 4 | Ranbp1 |
| Cep57l1 | 1,56E-05 | 0,370069344 | 0,269 | 0,051 | 0,247218398 | 4 | Cep57l1 |
| Rnaseh2b | 1,57E-05 | 0,349690294 | 0,385 | 0,101 | 0,248173355 | 4 | Rnaseh2b |
| Fignl1 | 1,57E-05 | 0,377717708 | 0,308 | 0,064 | 0,248995714 | 4 | Fignl1 |
| Gm28875 | 1,66E-05 | 0,396149089 | 0,577 | 0,209 | 0,263585626 | 4 | Gm28875 |
| Mcm4 | 1,67E-05 | 0,583308996 | 0,538 | 0,172 | 0,264808677 | 4 | Mcm4 |
| Slc43a3 | 1,75E-05 | 0,470707757 | 0,808 | 0,374 | 0,277433902 | 4 | Slc43a3 |
| Nup205 | 1,75E-05 | 0,27235962 | 0,5 | 0,145 | 0,277773758 | 4 | Nup205 |
| Acp1 | 1,80E-05 | 0,478821432 | 0,731 | 0,333 | 0,284926822 | 4 | Acp1 |
| Gtf2e2 | 1,93E-05 | 0,499844605 | 0,769 | 0,357 | 0,305913883 | 4 | Gtf2e2 |
| Psip1 | 1,98E-05 | 0,54426426 | 0,538 | 0,185 | 0,313913907 | 4 | Psip1 |
| Mcm5 | 2,05E-05 | 0,71324147 | 0,538 | 0,192 | 0,324302052 | 4 | Mcm5 |
| Rpa2 | 2,15E-05 | 0,403810614 | 0,5 | 0,155 | 0,341578177 | 4 | Rpa2 |
| Erh | 2,20E-05 | 0,367701952 | 0,962 | 0,768 | 0,347971754 | 4 | Erh |
| Zfp948 | 2,33E-05 | 0,347876822 | 0,538 | 0,168 | 0,369500827 | 4 | Zfp948 |
| Lancl2 | 2,34E-05 | 0,417513217 | 0,308 | 0,067 | 0,371249977 | 4 | Lancl2 |
| Fech | 2,39E-05 | 0,515959592 | 0,538 | 0,195 | 0,378519208 | 4 | Fech |
| Bgn | 2,40E-05 | 0,481443016 | 0,5 | 0,162 | 0,380772815 | 4 | Bgn |
| Ahctf1 | 2,44E-05 | 0,595541604 | 0,769 | 0,327 | 0,386141939 | 4 | Ahctf1 |
| Ccne2 | 2,44E-05 | 0,518734604 | 0,308 | 0,067 | 0,386344755 | 4 | Ccne2 |
| Dynlt1f | 2,58E-05 | 0,413552815 | 0,846 | 0,542 | 0,408553524 | 4 | Dynlt1f |
| Brip1os | 2,69E-05 | 0,424299804 | 0,5 | 0,178 | 0,426100806 | 4 | Brip1os |
| Nxt1 | 2,70E-05 | 0,294623324 | 0,577 | 0,209 | 0,427839983 | 4 | Nxt1 |
| H2afv | 2,72E-05 | 0,504410643 | 0,846 | 0,451 | 0,431199871 | 4 | H2afv |
| Polr3k | 2,88E-05 | 0,467203388 | 0,615 | 0,273 | 0,457155657 | 4 | Polr3k |
| 4930523C07Rik1 | 2,91E-05 | 0,301157817 | 0,538 | 0,182 | 0,460818918 | 4 | 4930523C07Rik |
| Prdx4 | 3,10E-05 | 0,501212599 | 0,846 | 0,508 | 0,491277699 | 4 | Prdx4 |
| Gmds | 3,11E-05 | 0,257662846 | 0,462 | 0,138 | 0,493398401 | 4 | Gmds |
| Dclk2 | 3,15E-05 | 0,263907276 | 0,308 | 0,071 | 0,498821837 | 4 | Dclk2 |
| Msh6 | 3,30E-05 | 0,257805638 | 0,385 | 0,104 | 0,522521489 | 4 | Msh6 |
| Trp531 | 3,42E-05 | 0,527405236 | 0,885 | 0,515 | 0,542722734 | 4 | Trp53 |
| Rfwd3 | 3,43E-05 | 0,286222941 | 0,423 | 0,118 | 0,544209803 | 4 | Rfwd3 |
| Mcm7 | 3,64E-05 | 0,823066025 | 0,538 | 0,215 | 0,576751429 | 4 | Mcm7 |
| Acot7 | 3,66E-05 | 0,370329991 | 0,654 | 0,283 | 0,579883364 | 4 | Acot7 |
| Banf1 | 3,85E-05 | 0,543904683 | 0,885 | 0,68 | 0,611003266 | 4 | Banf1 |
| Tdp1 | 4,00E-05 | 0,289904006 | 0,346 | 0,088 | 0,633624857 | 4 | Tdp1 |
| Nup85 | 4,13E-05 | 0,380563456 | 0,577 | 0,226 | 0,654338473 | 4 | Nup85 |
| Enpp4 | 4,34E-05 | 0,29452798 | 0,462 | 0,145 | 0,687824915 | 4 | Enpp4 |
| Cdh2 | 4,39E-05 | 0,407385366 | 0,5 | 0,172 | 0,69620362 | 4 | Cdh2 |
| Smc1a | 4,52E-05 | 0,698912133 | 0,769 | 0,455 | 0,71713066 | 4 | Smc1a |
| Dph5 | 4,73E-05 | 0,274621252 | 0,462 | 0,145 | 0,749820867 | 4 | Dph5 |
| Rbbp4 | 4,74E-05 | 0,587764481 | 0,808 | 0,512 | 0,751883889 | 4 | Rbbp4 |
| Gmnn | 4,93E-05 | 0,598456243 | 0,5 | 0,168 | 0,780873705 | 4 | Gmnn |
| Tardbp | 4,96E-05 | 0,273993866 | 0,808 | 0,384 | 0,786490013 | 4 | Tardbp |
| Cdca4 | 5,15E-05 | 0,446720146 | 0,577 | 0,236 | 0,815613174 | 4 | Cdca4 |
| Lyar | 5,39E-05 | 0,449731627 | 0,577 | 0,242 | 0,854436985 | 4 | Lyar |
| Srsf1 | 5,41E-05 | 0,485920574 | 0,731 | 0,387 | 0,85715908 | 4 | Srsf1 |
| Ddx39 | 5,44E-05 | 0,550272185 | 0,846 | 0,549 | 0,862435228 | 4 | Ddx39 |
| Fam92a | 5,63E-05 | 0,407656017 | 0,615 | 0,249 | 0,892909956 | 4 | Fam92a |
| Cdc23 | 6,02E-05 | 0,28019063 | 0,5 | 0,162 | 0,953827792 | 4 | Cdc23 |
| Prkar2a | 6,35E-05 | 0,436198025 | 0,5 | 0,185 | 1 | 4 | Prkar2a |
| Fubp1 | 6,78E-05 | 0,528877363 | 0,808 | 0,478 | 1 | 4 | Fubp1 |
| Pmf1 | 7,47E-05 | 0,446933901 | 0,615 | 0,279 | 1 | 4 | Pmf1 |
| Cbx3 | 7,47E-05 | 0,625209774 | 1 | 0,949 | 1 | 4 | Cbx3 |
| Mbd4 | 7,61E-05 | 0,462076543 | 0,346 | 0,094 | 1 | 4 | Mbd4 |
| Dck | 8,35E-05 | 0,729953749 | 0,538 | 0,229 | 1 | 4 | Dck |
| Ppp1r71 | 8,68E-05 | 0,425322885 | 0,654 | 0,286 | 1 | 4 | Ppp1r7 |
| Cdk41 | 8,73E-05 | 0,615033022 | 0,923 | 0,65 | 1 | 4 | Cdk4 |
| Tgtp1 | 8,86E-05 | 0,310962199 | 0,615 | 0,242 | 1 | 4 | Tgtp1 |
| Haus5 | 8,89E-05 | 0,417424847 | 0,308 | 0,077 | 1 | 4 | Haus5 |
| Sf3a3 | 9,17E-05 | 0,676753716 | 0,654 | 0,323 | 1 | 4 | Sf3a3 |
| Mrpl28 | 9,36E-05 | 0,438033552 | 0,769 | 0,401 | 1 | 4 | Mrpl28 |
| Wdr76 | 9,68E-05 | 0,318570824 | 0,423 | 0,141 | 1 | 4 | Wdr76 |
| Hnrnpa1 | 0,000104114 | 0,457422509 | 1 | 0,909 | 1 | 4 | Hnrnpa1 |
| Timm50 | 0,000107137 | 0,39321214 | 0,692 | 0,296 | 1 | 4 | Timm50 |
| Prpf38a | 0,000116261 | 0,348603153 | 0,577 | 0,226 | 1 | 4 | Prpf38a |
| Pcna | 0,000117463 | 0,983670287 | 0,731 | 0,508 | 1 | 4 | Pcna |
| Anapc5 | 0,000118892 | 0,541835028 | 0,923 | 0,66 | 1 | 4 | Anapc5 |
| Ctc1 | 0,000126728 | 0,508182481 | 0,385 | 0,114 | 1 | 4 | Ctc1 |
| Bcar1 | 0,000129959 | 0,334081427 | 0,769 | 0,401 | 1 | 4 | Bcar1 |
| Tpr | 0,000139638 | 0,591573209 | 0,923 | 0,569 | 1 | 4 | Tpr |
| Gm14305 | 0,000142963 | 0,294869652 | 0,462 | 0,168 | 1 | 4 | Gm14305 |
| Rhno1 | 0,000149168 | 0,336319586 | 0,462 | 0,165 | 1 | 4 | Rhno1 |
| Gm13461 | 0,000150773 | 0,289361946 | 0,885 | 0,667 | 1 | 4 | Gm13461 |
| Xrcc6 | 0,000153627 | 0,337948796 | 0,385 | 0,118 | 1 | 4 | Xrcc6 |
| Hnrnpf | 0,000155572 | 0,546345809 | 1 | 0,872 | 1 | 4 | Hnrnpf |
| Hmgn5 | 0,000156701 | 0,318002762 | 0,615 | 0,242 | 1 | 4 | Hmgn5 |
| Txndc12 | 0,000162136 | 0,392613169 | 0,885 | 0,498 | 1 | 4 | Txndc12 |
| Chek2 | 0,000168048 | 0,462210019 | 0,423 | 0,165 | 1 | 4 | Chek2 |
| Mtf2 | 0,00016841 | 0,294667847 | 0,577 | 0,226 | 1 | 4 | Mtf2 |
| Rdx | 0,000170685 | 0,418417939 | 0,885 | 0,556 | 1 | 4 | Rdx |
| Nt5dc2 | 0,000174307 | 0,338769861 | 0,5 | 0,195 | 1 | 4 | Nt5dc2 |
| Miip | 0,000176942 | 0,287919625 | 0,462 | 0,162 | 1 | 4 | Miip |
| E2f2 | 0,000177868 | 0,273550842 | 0,269 | 0,064 | 1 | 4 | E2f2 |
| Nop10 | 0,000178185 | 0,330525047 | 0,885 | 0,596 | 1 | 4 | Nop10 |
| Esyt2 | 0,000179942 | 0,342910096 | 0,462 | 0,168 | 1 | 4 | Esyt2 |
| Dtd1 | 0,00018031 | 0,300435798 | 0,385 | 0,118 | 1 | 4 | Dtd1 |
| Dpy30 | 0,000185969 | 0,393343232 | 0,692 | 0,364 | 1 | 4 | Dpy30 |
| Sumo2 | 0,000188772 | 0,44133607 | 0,962 | 0,842 | 1 | 4 | Sumo2 |
| Apex1 | 0,000189698 | 0,579217962 | 0,615 | 0,293 | 1 | 4 | Apex1 |
| Nde1 | 0,000190305 | 0,476867852 | 0,654 | 0,276 | 1 | 4 | Nde1 |
| Gatsl2 | 0,000194376 | 0,257101957 | 0,385 | 0,114 | 1 | 4 | Gatsl2 |
| Dctpp1 | 0,000198697 | 0,390079737 | 0,577 | 0,263 | 1 | 4 | Dctpp1 |
| Tmem97 | 0,000199441 | 0,276611554 | 0,423 | 0,135 | 1 | 4 | Tmem97 |
| Prps1 | 0,000199495 | 0,306836053 | 0,5 | 0,178 | 1 | 4 | Prps1 |
| Tdp2 | 0,000202744 | 0,332806819 | 0,5 | 0,189 | 1 | 4 | Tdp2 |
| Tagln21 | 0,000203499 | 0,449074454 | 0,962 | 0,855 | 1 | 4 | Tagln2 |
| Rad18 | 0,000205756 | 0,338347113 | 0,423 | 0,145 | 1 | 4 | Rad18 |
| Sox11 | 0,000206854 | 0,266347463 | 0,308 | 0,084 | 1 | 4 | Sox11 |
| Cep135 | 0,000207739 | 0,501403172 | 0,269 | 0,067 | 1 | 4 | Cep135 |
| Nudcd2 | 0,000211114 | 0,535596092 | 0,692 | 0,347 | 1 | 4 | Nudcd2 |
| Ssna1 | 0,00021314 | 0,381332243 | 0,885 | 0,522 | 1 | 4 | Ssna1 |
| Nckap11 | 0,000214831 | 0,419330724 | 0,808 | 0,418 | 1 | 4 | Nckap1 |
| Rbm3 | 0,000217076 | 0,356568858 | 0,962 | 0,939 | 1 | 4 | Rbm3 |

| | | | | | | | |
|---|---|---|---|---|---|---|---|
| Ncbp1 | 0,00022413 | 0,416315048 | 0,5 | 0,195 | 1 | 4 | Ncbp1 |
| Idh3a | 0,000224678 | 0,497839398 | 0,577 | 0,276 | 1 | 4 | Idh3a |
| Lrrc40 | 0,000242927 | 0,344953422 | 0,308 | 0,088 | 1 | 4 | Lrrc40 |
| Larp7 | 0,000247074 | 0,33470495 | 0,615 | 0,296 | 1 | 4 | Larp7 |
| Hn11 | 0,000249792 | 0,632570775 | 0,885 | 0,596 | 1 | 4 | Hn1 |
| Trim28 | 0,000257412 | 0,456979432 | 0,731 | 0,37 | 1 | 4 | Trim28 |
| Bola3 | 0,000261828 | 0,283762393 | 0,808 | 0,431 | 1 | 4 | Bola3 |
| Tmem106c | 0,00027184 | 0,268957955 | 0,385 | 0,125 | 1 | 4 | Tmem106c |
| Usp39 | 0,000279898 | 0,398464934 | 0,538 | 0,212 | 1 | 4 | Usp39 |
| Cdc27 | 0,000288168 | 0,308313308 | 0,538 | 0,222 | 1 | 4 | Cdc27 |
| Snw1 | 0,000308765 | 0,346572547 | 0,692 | 0,364 | 1 | 4 | Snw1 |
| Nop56 | 0,000316798 | 0,374620778 | 0,731 | 0,38 | 1 | 4 | Nop56 |
| Slbp | 0,000317923 | 0,641312922 | 0,769 | 0,418 | 1 | 4 | Slbp |
| Smc3 | 0,000324836 | 0,498648935 | 0,654 | 0,327 | 1 | 4 | Smc3 |
| Hist1h1c | 0,000335367 | 0,325555966 | 0,538 | 0,212 | 1 | 4 | Hist1h1c |
| Cnot6l | 0,000339392 | 0,382876817 | 0,538 | 0,232 | 1 | 4 | Cnot6l |
| Bace1 | 0,000340854 | 0,261591894 | 0,346 | 0,108 | 1 | 4 | Bace1 |
| Pole4 | 0,000347369 | 0,33195957 | 0,692 | 0,306 | 1 | 4 | Pole4 |
| Ptges3 | 0,000352079 | 0,543383273 | 0,846 | 0,825 | 1 | 4 | Ptges3 |
| Xpo1 | 0,000356219 | 0,484626335 | 0,615 | 0,276 | 1 | 4 | Xpo1 |
| Metap2 | 0,000364247 | 0,543135496 | 0,885 | 0,684 | 1 | 4 | Metap2 |
| Snrpb | 0,000372579 | 0,474389466 | 0,923 | 0,801 | 1 | 4 | Snrpb |
| Slc24a5 | 0,000376477 | 0,341554625 | 0,692 | 0,401 | 1 | 4 | Slc24a5 |
| Ccser2 | 0,000393399 | 0,443681071 | 0,731 | 0,401 | 1 | 4 | Ccser2 |
| Sept10 | 0,00039833 | 0,31038034 | 0,923 | 0,673 | 1 | 4 | Sept10 |
| Hist1h1e | 0,000407433 | 0,332611685 | 0,346 | 0,118 | 1 | 4 | Hist1h1e |
| Nutf2-ps1 | 0,000411214 | 0,40904806 | 0,846 | 0,589 | 1 | 4 | Nutf2-ps1 |
| Ncaph2 | 0,000414616 | 0,480982658 | 0,692 | 0,407 | 1 | 4 | Ncaph2 |
| Spns21 | 0,000414665 | 0,339942746 | 0,577 | 0,256 | 1 | 4 | Spns2 |
| Pola1 | 0,000416241 | 0,496754721 | 0,308 | 0,094 | 1 | 4 | Pola1 |
| Pfas | 0,000427941 | 0,380629259 | 0,615 | 0,306 | 1 | 4 | Pfas |
| Ddx42 | 0,000457848 | 0,346627899 | 0,654 | 0,337 | 1 | 4 | Ddx42 |
| Rfc1 | 0,000461611 | 0,395100585 | 0,462 | 0,195 | 1 | 4 | Rfc1 |
| Hjurp | 0,000464213 | 0,359318525 | 0,692 | 0,387 | 1 | 4 | Hjurp |
| Rfc3 | 0,000464527 | 0,367144803 | 0,423 | 0,155 | 1 | 4 | Rfc3 |
| Cct71 | 0,000468691 | 0,595654946 | 0,808 | 0,643 | 1 | 4 | Cct7 |
| Pebp1 | 0,000469899 | 0,387713185 | 0,885 | 0,572 | 1 | 4 | Pebp1 |
| Sorbs21 | 0,000477685 | 0,33515424 | 0,731 | 0,38 | 1 | 4 | Sorbs2 |
| Ddx39b | 0,000491796 | 0,492938786 | 0,885 | 0,643 | 1 | 4 | Ddx39b |
| Gart | 0,000523688 | 0,41853281 | 0,5 | 0,212 | 1 | 4 | Gart |
| Senp1 | 0,000528564 | 0,434907678 | 0,385 | 0,148 | 1 | 4 | Senp1 |
| Skiv2l2 | 0,000536157 | 0,425415069 | 0,615 | 0,323 | 1 | 4 | Skiv2l2 |
| Rbm8a | 0,000557748 | 0,460357628 | 0,885 | 0,606 | 1 | 4 | Rbm8a |
| Saal1 | 0,000573156 | 0,283504226 | 0,308 | 0,091 | 1 | 4 | Saal1 |
| Uck2 | 0,000578526 | 0,262687369 | 0,731 | 0,391 | 1 | 4 | Uck2 |
| Cul5 | 0,000578578 | 0,522879288 | 0,615 | 0,31 | 1 | 4 | Cul5 |
| Pou2f1 | 0,000610584 | 0,322102867 | 0,346 | 0,111 | 1 | 4 | Pou2f1 |
| Hnrnpm | 0,000623899 | 0,331302863 | 0,962 | 0,569 | 1 | 4 | Hnrnpm |
| Setx | 0,000638483 | 0,388236528 | 0,615 | 0,32 | 1 | 4 | Setx |
| Aldh18a1 | 0,000643311 | 0,551416312 | 0,346 | 0,114 | 1 | 4 | Aldh18a1 |
| Fh1 | 0,000664059 | 0,285745824 | 0,615 | 0,293 | 1 | 4 | Fh1 |
| Nudc | 0,000665619 | 0,411958196 | 0,923 | 0,593 | 1 | 4 | Nudc |
| Vars | 0,000687327 | 0,437876554 | 0,615 | 0,33 | 1 | 4 | Vars |
| Reep4 | 0,000689693 | 0,399982094 | 0,385 | 0,138 | 1 | 4 | Reep4 |
| Lsm6 | 0,000696128 | 0,262241704 | 0,846 | 0,465 | 1 | 4 | Lsm6 |
| Mdm2 | 0,000715952 | 0,325098634 | 0,615 | 0,34 | 1 | 4 | Mdm2 |
| Bag5 | 0,000729645 | 0,316043906 | 0,577 | 0,263 | 1 | 4 | Bag5 |
| 1810037I17Rik | 0,00075201 | 0,351855739 | 0,885 | 0,65 | 1 | 4 | 1810037I17Rik |
| Cpne81 | 0,00076461 | 0,354042751 | 0,808 | 0,455 | 1 | 4 | Cpne8 |
| Cdt11 | 0,000778736 | 0,599044634 | 0,385 | 0,145 | 1 | 4 | Cdt1 |
| Oat | 0,000797705 | 0,503235437 | 0,5 | 0,222 | 1 | 4 | Oat |
| Anapc4 | 0,000804846 | 0,437028176 | 0,615 | 0,323 | 1 | 4 | Anapc4 |
| Magoh | 0,000815194 | 0,382714653 | 0,731 | 0,418 | 1 | 4 | Magoh |
| Xpo4 | 0,000828364 | 0,405159809 | 0,346 | 0,121 | 1 | 4 | Xpo4 |
| Fundc2 | 0,000841178 | 0,270797338 | 0,769 | 0,424 | 1 | 4 | Fundc2 |
| Ilf2 | 0,00085154 | 0,658099914 | 0,654 | 0,407 | 1 | 4 | Ilf2 |
| Ptgfrn | 0,000857349 | 0,66437979 | 0,5 | 0,242 | 1 | 4 | Ptgfrn |
| Bcl6b1 | 0,000863145 | 0,301857172 | 0,692 | 0,32 | 1 | 4 | Bcl6b |
| Abce1 | 0,000873304 | 0,526261667 | 0,615 | 0,347 | 1 | 4 | Abce1 |
| Cse1l | 0,00087961 | 0,356467456 | 0,538 | 0,236 | 1 | 4 | Cse1l |
| Pmp221 | 0,000895072 | 0,261674867 | 0,538 | 0,226 | 1 | 4 | Pmp22 |
| Hnrnpab | 0,000895355 | 0,383211953 | 0,962 | 0,737 | 1 | 4 | Hnrnpab |
| Adss | 0,000902004 | 0,381331164 | 0,692 | 0,35 | 1 | 4 | Adss |
| Nedd42 | 0,000903078 | 0,332474485 | 1 | 0,694 | 1 | 4 | Nedd4 |
| Prpf4 | 0,000909132 | 0,397377924 | 0,423 | 0,165 | 1 | 4 | Prpf4 |
| Eif4e | 0,000921791 | 0,536239494 | 0,731 | 0,485 | 1 | 4 | Eif4e |
| Cdc123 | 0,000927381 | 0,274664708 | 0,731 | 0,377 | 1 | 4 | Cdc123 |
| Nsmce4a | 0,000969876 | 0,371593141 | 0,5 | 0,226 | 1 | 4 | Nsmce4a |
| Dtymk | 0,000985871 | 0,288603889 | 0,731 | 0,394 | 1 | 4 | Dtymk |
| Csnk1a1 | 0,000996366 | 0,359474098 | 1 | 0,731 | 1 | 4 | Csnk1a1 |
| Snrpa1 | 0,001003124 | 0,34220417 | 0,769 | 0,391 | 1 | 4 | Snrpa1 |
| Hcfc1 | 0,001034694 | 0,327986452 | 0,654 | 0,34 | 1 | 4 | Hcfc1 |
| Irf1 | 0,001040847 | 0,296978351 | 0,692 | 0,367 | 1 | 4 | Irf1 |
| Lrrc581 | 0,001069773 | 0,396857753 | 1 | 0,919 | 1 | 4 | Lrrc58 |
| Abcb7 | 0,001081405 | 0,304647804 | 0,269 | 0,077 | 1 | 4 | Abcb7 |
| Arhgef151 | 0,001100182 | 0,321226437 | 0,885 | 0,559 | 1 | 4 | Arhgef15 |
| Eif4a3 | 0,001101363 | 0,571396802 | 0,769 | 0,535 | 1 | 4 | Eif4a3 |
| Rbmx | 0,001147673 | 0,27752192 | 0,385 | 0,145 | 1 | 4 | Rbmx |
| Eed | 0,001157582 | 0,360644324 | 0,462 | 0,205 | 1 | 4 | Eed |
| Sqle | 0,001175399 | 0,38789123 | 0,385 | 0,145 | 1 | 4 | Sqle |
| Haus4 | 0,001187806 | 0,266914668 | 0,423 | 0,158 | 1 | 4 | Haus4 |
| Mre11a | 0,001222093 | 0,386164001 | 0,308 | 0,098 | 1 | 4 | Mre11a |
| Eef1d | 0,001281902 | 0,378673966 | 0,885 | 0,613 | 1 | 4 | Eef1d |
| Nrm | 0,00128435 | 0,288128806 | 0,346 | 0,121 | 1 | 4 | Nrm |
| Idh2 | 0,001319593 | 0,462580693 | 0,615 | 0,316 | 1 | 4 | Idh2 |
| Lmf2 | 0,001342014 | 0,383687249 | 0,731 | 0,391 | 1 | 4 | Lmf2 |
| Haus81 | 0,001347587 | 0,356659299 | 0,346 | 0,125 | 1 | 4 | Haus8 |
| Gm5641 | 0,001351571 | 0,343903079 | 0,731 | 0,455 | 1 | 4 | Gm5641 |
| Chaf1b | 0,001381375 | 0,558915521 | 0,308 | 0,104 | 1 | 4 | Chaf1b |
| Tm9sf1 | 0,001382721 | 0,492197662 | 0,692 | 0,374 | 1 | 4 | Tm9sf1 |
| Syncrip | 0,001433413 | 0,455820735 | 0,731 | 0,471 | 1 | 4 | Syncrip |
| Rcan3 | 0,001468182 | 0,397667588 | 0,5 | 0,242 | 1 | 4 | Rcan3 |
| Hsp90aa1 | 0,001484324 | 0,425807751 | 0,962 | 0,778 | 1 | 4 | Hsp90aa1 |
| Mcm6 | 0,001501957 | 0,53403897 | 0,615 | 0,343 | 1 | 4 | Mcm6 |
| Traf71 | 0,00151551 | 0,387780986 | 0,692 | 0,377 | 1 | 4 | Traf7 |
| Arpp19 | 0,001548831 | 0,286308722 | 1 | 0,828 | 1 | 4 | Arpp19 |
| Paics | 0,001550086 | 0,38285376 | 0,808 | 0,471 | 1 | 4 | Paics |
| Npm1 | 0,001605372 | 0,507958893 | 0,962 | 0,902 | 1 | 4 | Npm1 |
| Sigirr | 0,001651044 | 0,388500438 | 0,577 | 0,283 | 1 | 4 | Sigirr |
| Cbx6 | 0,001651255 | 0,393694629 | 0,346 | 0,121 | 1 | 4 | Cbx6 |
| Ndufs1 | 0,001663712 | 0,292822433 | 0,615 | 0,293 | 1 | 4 | Ndufs1 |
| Ndc1 | 0,001664646 | 0,291321355 | 0,308 | 0,104 | 1 | 4 | Ndc1 |
| Eri1 | 0,001748155 | 0,25342687 | 0,423 | 0,165 | 1 | 4 | Eri1 |
| 4932438A13Rik | 0,001769152 | 0,387592432 | 0,731 | 0,401 | 1 | 4 | 4932438A13Rik |
| Ptbp11 | 0,001776573 | 0,439735785 | 0,769 | 0,559 | 1 | 4 | Ptbp1 |
| Nfib1 | 0,001777176 | 0,408571372 | 0,885 | 0,599 | 1 | 4 | Nfib |
| AI506816 | 0,001780105 | 0,559389826 | 0,692 | 0,441 | 1 | 4 | AI506816 |
| Mcm3 | 0,001834633 | 0,400354958 | 0,538 | 0,273 | 1 | 4 | Mcm3 |
| Ak6 | 0,001955284 | 0,453708553 | 0,846 | 0,559 | 1 | 4 | Ak6 |
| Dennd5b | 0,001967123 | 0,270686452 | 0,692 | 0,377 | 1 | 4 | Dennd5b |
| Morf4l21 | 0,001981969 | 0,371298597 | 1 | 0,795 | 1 | 4 | Morf4l2 |

| | | | | | | | |
|---|---|---|---|---|---|---|---|
| Pds5a | 0,002040127 | 0,349488827 | 0,577 | 0,316 | 1 | 4 | Pds5a |
| Nhp2 | 0,00207112 | 0,276073779 | 0,808 | 0,505 | 1 | 4 | Nhp2 |
| Commd10 | 0,002074028 | 0,323562754 | 0,692 | 0,407 | 1 | 4 | Commd10 |
| Phc1 | 0,002098622 | 0,386422285 | 0,346 | 0,125 | 1 | 4 | Phc1 |
| Ifrd1 | 0,00212525 | 0,254676111 | 0,577 | 0,293 | 1 | 4 | Ifrd1 |
| Ptprk1 | 0,002125596 | 0,353991562 | 0,731 | 0,441 | 1 | 4 | Ptprk |
| Nup93 | 0,002176561 | 0,331913446 | 0,5 | 0,226 | 1 | 4 | Nup93 |
| Suv39h1 | 0,00218385 | 0,281481603 | 0,346 | 0,128 | 1 | 4 | Suv39h1 |
| Hnrnpk | 0,002249413 | 0,361775651 | 0,962 | 0,97 | 1 | 4 | Hnrnpk |
| Bub3 | 0,002270754 | 0,448098862 | 0,808 | 0,556 | 1 | 4 | Bub3 |
| Emg1 | 0,00230779 | 0,31678819 | 0,769 | 0,478 | 1 | 4 | Emg1 |
| Snrpe | 0,002348526 | 0,251665876 | 0,885 | 0,71 | 1 | 4 | Snrpe |
| Rsl1d1 | 0,002362606 | 0,456837652 | 0,615 | 0,357 | 1 | 4 | Rsl1d1 |
| Hnrnpu | 0,002414952 | 0,400419297 | 0,923 | 0,774 | 1 | 4 | Hnrnpu |
| Btbd10 | 0,00245016 | 0,262688953 | 0,385 | 0,155 | 1 | 4 | Btbd10 |
| Ddx50 | 0,002490292 | 0,436017263 | 0,692 | 0,441 | 1 | 4 | Ddx50 |
| Tmem2522 | 0,002522726 | 0,427904483 | 0,923 | 0,599 | 1 | 4 | Tmem252 |
| Hspd1 | 0,002524513 | 0,550103867 | 0,692 | 0,401 | 1 | 4 | Hspd1 |
| Sf3a1 | 0,002538131 | 0,260714237 | 0,538 | 0,283 | 1 | 4 | Sf3a1 |
| Fem1c | 0,00255264 | 0,345014732 | 0,5 | 0,246 | 1 | 4 | Fem1c |
| Hnrnpa2b1 | 0,002629211 | 0,307344815 | 1 | 0,97 | 1 | 4 | Hnrnpa2b1 |
| Mis12 | 0,002640357 | 0,273810482 | 0,346 | 0,131 | 1 | 4 | Mis12 |
| Tmem98 | 0,002740893 | 0,401047086 | 0,5 | 0,253 | 1 | 4 | Tmem98 |
| Ewsr1 | 0,00277803 | 0,436253573 | 0,923 | 0,7 | 1 | 4 | Ewsr1 |
| Foxp11 | 0,002884621 | 0,397266785 | 0,808 | 0,603 | 1 | 4 | Foxp1 |
| Actn42 | 0,002949146 | 0,328135609 | 0,962 | 0,825 | 1 | 4 | Actn4 |
| Mtmr2 | 0,002959692 | 0,270310615 | 0,923 | 0,68 | 1 | 4 | Mtmr2 |
| Yaf2 | 0,003033766 | 0,371827324 | 0,654 | 0,357 | 1 | 4 | Yaf2 |
| Eif4g21 | 0,003064878 | 0,305951544 | 1 | 0,896 | 1 | 4 | Eif4g2 |
| Ckap4 | 0,003098924 | 0,275315001 | 0,654 | 0,367 | 1 | 4 | Ckap4 |
| Nt5c3 | 0,003291074 | 0,309085254 | 0,423 | 0,189 | 1 | 4 | Nt5c3 |
| Psmb7 | 0,003377212 | 0,351383049 | 0,808 | 0,616 | 1 | 4 | Psmb7 |
| Hdac3 | 0,00338766 | 0,607822481 | 0,462 | 0,266 | 1 | 4 | Hdac3 |
| Pfdn6 | 0,003522117 | 0,323592012 | 0,654 | 0,394 | 1 | 4 | Pfdn6 |
| Pbdc1 | 0,003555042 | 0,270102768 | 0,654 | 0,37 | 1 | 4 | Pbdc1 |
| Cggbp1 | 0,003697607 | 0,262346918 | 0,769 | 0,488 | 1 | 4 | Cggbp1 |
| Nme1 | 0,003755208 | 0,419369076 | 0,885 | 0,66 | 1 | 4 | Nme1 |
| Atrx | 0,003834 | 0,596193858 | 0,885 | 0,606 | 1 | 4 | Atrx |
| Cops7a | 0,003885847 | 0,314004403 | 0,731 | 0,424 | 1 | 4 | Cops7a |
| Sf3b3 | 0,003910793 | 0,499686723 | 0,654 | 0,38 | 1 | 4 | Sf3b3 |
| Sac3d1 | 0,003920886 | 0,278733932 | 0,423 | 0,195 | 1 | 4 | Sac3d1 |
| Cdkn2aipnl | 0,004075115 | 0,312102192 | 0,731 | 0,478 | 1 | 4 | Cdkn2aipnl |
| Mad1l1 | 0,004082832 | 0,331017573 | 0,385 | 0,165 | 1 | 4 | Mad1l1 |
| Scoc | 0,004100055 | 0,40817843 | 0,615 | 0,377 | 1 | 4 | Scoc |
| Gm20390 | 0,00415017 | 0,348010676 | 0,846 | 0,576 | 1 | 4 | Gm20390 |
| Rnf19a1 | 0,004277098 | 0,363343588 | 0,5 | 0,256 | 1 | 4 | Rnf19a |
| Kdr2 | 0,004298628 | 0,274964298 | 0,962 | 0,636 | 1 | 4 | Kdr |
| Mzt1 | 0,004325405 | 0,322366126 | 0,577 | 0,333 | 1 | 4 | Mzt1 |
| Ube2i | 0,004372632 | 0,290576301 | 0,923 | 0,781 | 1 | 4 | Ube2i |
| Pnpt1 | 0,004430852 | 0,278530868 | 0,423 | 0,199 | 1 | 4 | Pnpt1 |
| Acadl | 0,004652178 | 0,345571067 | 0,769 | 0,512 | 1 | 4 | Acadl |
| Thrap3 | 0,004693192 | 0,37812433 | 0,808 | 0,549 | 1 | 4 | Thrap3 |
| Smad5 | 0,004741123 | 0,272226562 | 0,692 | 0,414 | 1 | 4 | Smad5 |
| Cyc1 | 0,004748444 | 0,412402023 | 0,846 | 0,586 | 1 | 4 | Cyc1 |
| Prpf18 | 0,004761196 | 0,360208663 | 0,423 | 0,202 | 1 | 4 | Prpf18 |
| Ppfibp11 | 0,004780677 | 0,452431256 | 0,885 | 0,646 | 1 | 4 | Ppfibp1 |
| Prdx6 | 0,004898488 | 0,419756939 | 0,769 | 0,535 | 1 | 4 | Prdx6 |
| Sept71 | 0,005173022 | 0,430881839 | 1 | 0,781 | 1 | 4 | Sept7 |
| Abcf2 | 0,005331874 | 0,325272912 | 0,5 | 0,259 | 1 | 4 | Abcf2 |
| Psmd1 | 0,005365597 | 0,332615028 | 0,885 | 0,586 | 1 | 4 | Psmd1 |
| Mum1 | 0,005445942 | 0,26306029 | 0,5 | 0,242 | 1 | 4 | Mum1 |
| Dnajc3 | 0,005497515 | 0,449844542 | 0,808 | 0,545 | 1 | 4 | Dnajc3 |
| Mrpl171 | 0,005533366 | 0,294240962 | 0,923 | 0,7 | 1 | 4 | Mrpl17 |
| Bzw1 | 0,005601851 | 0,29585303 | 0,962 | 0,774 | 1 | 4 | Bzw1 |
| G3bp1 | 0,005659104 | 0,396282059 | 0,846 | 0,64 | 1 | 4 | G3bp1 |
| Nol11 | 0,005693551 | 0,432694068 | 0,346 | 0,152 | 1 | 4 | Nol11 |
| Ddb1 | 0,005779438 | 0,306228124 | 0,846 | 0,569 | 1 | 4 | Ddb1 |
| Ptpre | 0,005895568 | 0,335414972 | 0,731 | 0,468 | 1 | 4 | Ptpre |
| Eif2s11 | 0,005931635 | 0,488885085 | 0,769 | 0,559 | 1 | 4 | Eif2s1 |
| Lrch3 | 0,005966452 | 0,380040031 | 0,5 | 0,253 | 1 | 4 | Lrch3 |
| Lpp1 | 0,006033822 | 0,400011311 | 0,885 | 0,643 | 1 | 4 | Lpp |
| Rps6ka3 | 0,00606259 | 0,318835224 | 0,615 | 0,35 | 1 | 4 | Rps6ka3 |
| Ruvbl2 | 0,006097777 | 0,391555534 | 0,538 | 0,316 | 1 | 4 | Ruvbl2 |
| Lias | 0,006099305 | 0,296332785 | 0,462 | 0,226 | 1 | 4 | Lias |
| C1qtnf1 | 0,006146375 | 0,3013892 | 0,577 | 0,313 | 1 | 4 | C1qtnf1 |
| Cnih1 | 0,00624765 | 0,29594002 | 0,731 | 0,451 | 1 | 4 | Cnih1 |
| Lsm4 | 0,00632255 | 0,326615896 | 0,731 | 0,576 | 1 | 4 | Lsm4 |
| Vangl1 | 0,006365817 | 0,357804081 | 0,577 | 0,31 | 1 | 4 | Vangl1 |
| Tmco1 | 0,006383131 | 0,355091008 | 0,885 | 0,579 | 1 | 4 | Tmco1 |
| Hnrnpr | 0,00639145 | 0,255004229 | 0,731 | 0,397 | 1 | 4 | Hnrnpr |
| Cnot7 | 0,006394931 | 0,270583374 | 0,577 | 0,293 | 1 | 4 | Cnot7 |
| Uchl5 | 0,006471404 | 0,494299285 | 0,5 | 0,306 | 1 | 4 | Uchl5 |
| Rsrc1 | 0,006530706 | 0,311052972 | 0,5 | 0,283 | 1 | 4 | Rsrc1 |
| Vdac3 | 0,006677273 | 0,403151214 | 0,808 | 0,62 | 1 | 4 | Vdac3 |
| Tpm42 | 0,006757866 | 0,315619232 | 1 | 0,919 | 1 | 4 | Tpm4 |
| Exoc4 | 0,006765523 | 0,40916785 | 0,5 | 0,232 | 1 | 4 | Exoc4 |
| Nr3c1 | 0,006781526 | 0,330522511 | 0,5 | 0,266 | 1 | 4 | Nr3c1 |
| Psmc3 | 0,006824978 | 0,30128153 | 0,923 | 0,754 | 1 | 4 | Psmc3 |
| Rock2 | 0,006931161 | 0,269567968 | 0,769 | 0,448 | 1 | 4 | Rock2 |
| Atp5o | 0,007018681 | 0,301417137 | 0,885 | 0,785 | 1 | 4 | Atp5o |
| Cul1 | 0,007020667 | 0,310678348 | 0,654 | 0,438 | 1 | 4 | Cul1 |
| Bex3 | 0,007063181 | 0,421220817 | 0,5 | 0,306 | 1 | 4 | Bex3 |
| Mdp1 | 0,007071239 | 0,330180917 | 0,462 | 0,239 | 1 | 4 | Mdp1 |
| Rrp9 | 0,007162519 | 0,302912567 | 0,385 | 0,172 | 1 | 4 | Rrp9 |
| Vbp1 | 0,007191328 | 0,298587321 | 0,654 | 0,394 | 1 | 4 | Vbp1 |
| Atad3a | 0,007332203 | 0,401567459 | 0,385 | 0,175 | 1 | 4 | Atad3a |
| Maoa1 | 0,007348431 | 0,251256031 | 0,462 | 0,229 | 1 | 4 | Maoa |
| Cbfb | 0,007367904 | 0,305743682 | 0,577 | 0,343 | 1 | 4 | Cbfb |
| Fam57a | 0,007370972 | 0,283321063 | 0,538 | 0,283 | 1 | 4 | Fam57a |
| Tmem263 | 0,007475196 | 0,34073191 | 0,654 | 0,401 | 1 | 4 | Tmem263 |
| Txnl1 | 0,007483034 | 0,261356067 | 0,731 | 0,441 | 1 | 4 | Txnl1 |
| Prdx1 | 0,007511631 | 0,276283305 | 1 | 0,933 | 1 | 4 | Prdx1 |
| Terf1 | 0,007543319 | 0,30525091 | 0,385 | 0,195 | 1 | 4 | Terf1 |
| Stip1 | 0,007639978 | 0,411542538 | 0,654 | 0,401 | 1 | 4 | Stip1 |
| Cdv3 | 0,007779264 | 0,252263766 | 0,731 | 0,475 | 1 | 4 | Cdv3 |
| Bok1 | 0,007939839 | 0,455440041 | 0,654 | 0,374 | 1 | 4 | Bok |
| Pnn | 0,008052137 | 0,321902291 | 0,615 | 0,367 | 1 | 4 | Pnn |
| Stard4 | 0,008261766 | 0,341964547 | 0,5 | 0,269 | 1 | 4 | Stard4 |
| Srsf7 | 0,008329221 | 0,499520473 | 0,808 | 0,62 | 1 | 4 | Srsf7 |
| Polr2a | 0,008454682 | 0,370306873 | 0,731 | 0,475 | 1 | 4 | Polr2a |
| Pcnp | 0,008515155 | 0,313707087 | 0,962 | 0,65 | 1 | 4 | Pcnp |
| Serbp1 | 0,008559509 | 0,348017357 | 0,885 | 0,835 | 1 | 4 | Serbp1 |
| Eif3b | 0,008685734 | 0,257394744 | 0,769 | 0,485 | 1 | 4 | Eif3b |
| Pdlim12 | 0,008764796 | 0,305473059 | 1 | 0,68 | 1 | 4 | Pdlim1 |
| Nop58 | 0,008935347 | 0,314659671 | 0,538 | 0,306 | 1 | 4 | Nop58 |
| Psmd11 | 0,00899522 | 0,325887091 | 0,731 | 0,478 | 1 | 4 | Psmd11 |
| Med21 | 0,00905472 | 0,289840737 | 0,654 | 0,374 | 1 | 4 | Med21 |
| Nelfcd | 0,009115361 | 0,386697596 | 0,423 | 0,212 | 1 | 4 | Nelfcd |
| Clcc1 | 0,009144606 | 0,287964744 | 0,5 | 0,256 | 1 | 4 | Clcc1 |
| Gabpb1 | 0,009159407 | 0,30169468 | 0,5 | 0,279 | 1 | 4 | Gabpb1 |
| Abcc41 | 0,009182116 | 0,267470491 | 0,654 | 0,411 | 1 | 4 | Abcc4 |
| Nnt | 0,009377173 | 0,295580726 | 0,308 | 0,131 | 1 | 4 | Nnt |
| Acat1 | 0,009533531 | 0,278417034 | 0,692 | 0,418 | 1 | 4 | Acat1 |
| Eif61 | 0,00954632 | 0,276613927 | 0,846 | 0,596 | 1 | 4 | Eif6 |
| Nsmce2 | 0,009564087 | 0,307072352 | 0,538 | 0,316 | 1 | 4 | Nsmce2 |
| Topbp1 | 0,00980864 | 0,314917924 | 0,462 | 0,242 | 1 | 4 | Topbp1 |
| Kars | 0,009981205 | 0,35528808 | 0,808 | 0,593 | 1 | 4 | Kars |
| Mrpl51 | 0,009992646 | 0,250728522 | 0,769 | 0,569 | 1 | 4 | Mrpl51 |
