## Supplementary table 1 for "Emerging single cell endothelial heterogeneity supports sprouting tumour angiogenesis and growth"

### Supplementary Table 2

| p_val | avg_logFC | pct.1 | pct.2 | p_val_adj | cluster | gene | log2fc | log2pct |
|---|---|---|---|---|---|---|---|---|
| 1,59E-112 | 0,87802355 | 0,953 | 0,622 | 5,60E-108 | C1_KCNE3 | FLT1 | 1,26672022 | 0,61156021 |
| 7,93E-108 | 0,846246705 | 0,87 | 0,554 | 2,79E-103 | C1_KCNE3 | HTRA1 | 1,22087592 | 0,64643473 |
| 5,73E-99 | 1,00221056 | 0,931 | 0,611 | 2,01E-94 | C1_KCNE3 | INSR | 1,4458842 | 0,60357818 |
| 5,03E-93 | 0,769270394 | 0,903 | 0,577 | 1,77E-88 | C1_KCNE3 | VWA1 | 1,10982258 | 0,64167314 |
| 1,58E-91 | 1,587201504 | 0,454 | 0,088 | 5,54E-87 | C1_KCNE3 | ESM1 | 2,28984774 | 2,30319153 |
| 8,39E-86 | 0,746708624 | 0,818 | 0,469 | 2,95E-81 | C1_KCNE3 | KDR | 1,07727283 | 0,79600537 |
| 6,95E-78 | 0,887854634 | 0,69 | 0,351 | 2,45E-73 | C1_KCNE3 | LINC00152 | 1,28090348 | 0,96513574 |
| 2,42E-71 | 0,633814295 | 0,433 | 0,086 | 8,51E-67 | C1_KCNE3 | GABRD | 0,91440074 | 2,26699242 |
| 5,71E-70 | 0,593903839 | 0,463 | 0,131 | 2,01E-65 | C1_KCNE3 | COL13A1 | 0,85682212 | 1,78290188 |
| 6,65E-67 | 0,685821719 | 0,846 | 0,557 | 2,34E-62 | C1_KCNE3 | TMEM204 | 0,98943159 | 0,598589 |
| 3,73E-66 | 0,622761943 | 0,719 | 0,358 | 1,31E-61 | C1_KCNE3 | HECW2 | 0,89845557 | 0,99602015 |
| 2,63E-64 | 0,576362347 | 0,506 | 0,132 | 9,25E-60 | C1_KCNE3 | EDIL3 | 0,8315151 | 1,8991474 |
| 4,28E-64 | 0,59301742 | 0,78 | 0,462 | 1,50E-59 | C1_KCNE3 | MLEC | 0,85554329 | 0,7492701 |
| 3,02E-63 | 0,493853512 | 0,43 | 0,097 | 1,06E-58 | C1_KCNE3 | ADAMTSL2 | 0,71248001 | 2,09244625 |
| 6,17E-61 | 0,435978375 | 0,336 | 0,055 | 2,17E-56 | C1_KCNE3 | ITGA8 | 0,62898384 | 2,50673733 |
| 1,85E-59 | 0,601928682 | 0,78 | 0,443 | 6,52E-55 | C1_KCNE3 | ITGA1 | 0,86839953 | 0,80919392 |
| 2,95E-59 | 0,667206633 | 0,541 | 0,182 | 1,04E-54 | C1_KCNE3 | EDNRB | 0,9625757 | 1,54586268 |
| 2,63E-58 | 0,601855227 | 0,468 | 0,127 | 9,24E-54 | C1_KCNE3 | UNC5B | 0,86829355 | 1,84130225 |
| 4,34E-56 | 0,845373429 | 0,463 | 0,161 | 1,52E-51 | C1_KCNE3 | CA2 | 1,21961605 | 1,49532529 |
| 5,63E-55 | 0,673657323 | 0,52 | 0,216 | 1,98E-50 | C1_KCNE3 | MIR4435-1H( | 0,97188208 | 1,24827105 |
| 1,34E-54 | 0,525687935 | 0,674 | 0,344 | 4,72E-50 | C1_KCNE3 | PINK1 | 0,75840738 | 0,96018454 |
| 6,06E-54 | 0,571016751 | 0,596 | 0,247 | 2,13E-49 | C1_KCNE3 | PDGFD | 0,82380303 | 1,25394126 |
| 7,19E-54 | 0,722655084 | 0,681 | 0,442 | 2,53E-49 | C1_KCNE3 | PPAP2B | 1,04257091 | 0,61793374 |
| 2,64E-53 | 0,577797715 | 0,712 | 0,354 | 9,27E-49 | C1_KCNE3 | LAMA4 | 0,8335859 | 0,99798927 |
| 7,36E-53 | 1,056536355 | 0,735 | 0,374 | 2,59E-48 | C1_KCNE3 | IGFBP3 | 1,52425976 | 0,96532742 |
| 2,97E-45 | 0,537242979 | 0,456 | 0,173 | 1,05E-40 | C1_KCNE3 | CPM | 0,77507778 | 1,37288951 |
| 3,49E-45 | 0,584129368 | 0,522 | 0,22 | 1,23E-40 | C1_KCNE3 | LBH | 0,84272054 | 1,22787796 |
| 7,50E-45 | 0,481220259 | 0,742 | 0,431 | 2,64E-40 | C1_KCNE3 | PTPRG | 0,69425408 | 0,77678011 |
| 8,96E-45 | 0,646316267 | 0,723 | 0,417 | 3,15E-40 | C1_KCNE3 | IVNS1ABP | 0,93243727 | 0,78669545 |
| 1,70E-43 | 0,51650935 | 0,591 | 0,303 | 5,96E-39 | C1_KCNE3 | PMEPA1 | 0,74516548 | 0,95238198 |
| 7,20E-43 | 0,575370801 | 0,239 | 0,042 | 2,53E-38 | C1_KCNE3 | KCNE3 | 0,8300846 | 2,37614849 |
| 6,67E-42 | 0,471656107 | 0,634 | 0,335 | 2,35E-37 | C1_KCNE3 | DYSF | 0,68045593 | 0,91028118 |
| 3,30E-41 | 0,878495504 | 0,423 | 0,158 | 1,16E-36 | C1_KCNE3 | ANGPT2 | 1,26740111 | 1,39273883 |
| 1,92E-40 | 0,461466877 | 0,589 | 0,268 | 6,76E-36 | C1_KCNE3 | CDA | 0,66575597 | 1,12156198 |
| 1,56E-39 | 0,469998415 | 0,676 | 0,402 | 5,49E-35 | C1_KCNE3 | GRB10 | 0,67806438 | 0,742626 |
| 2,97E-39 | 0,507460943 | 0,634 | 0,33 | 1,05E-34 | C1_KCNE3 | NRP2 | 0,73211139 | 0,93165484 |
| 3,86E-39 | 0,506416454 | 0,487 | 0,192 | 1,36E-34 | C1_KCNE3 | ANGPTL2 | 0,73060451 | 1,32046269 |
| 1,11E-38 | 0,497387291 | 0,693 | 0,395 | 3,89E-34 | C1_KCNE3 | MYO1B | 0,71757818 | 0,80322704 |
| 5,26E-37 | 0,426138742 | 0,683 | 0,363 | 1,85E-32 | C1_KCNE3 | RASGRP3 | 0,61478825 | 0,9027028 |
| 5,61E-37 | 0,58821679 | 0,605 | 0,291 | 1,97E-32 | C1_KCNE3 | TNFRSF4 | 0,84861745 | 1,04321207 |
| 4,81E-172 | 1,412837547 | 0,699 | 0,159 | 1,69E-167 | C2_NID2 | PGF | 2,03829372 | 2,10187961 |
| 8,96E-118 | 0,885838766 | 0,493 | 0,062 | 3,15E-113 | C2_NID2 | LOX | 1,27799519 | 2,89391274 |
| 9,61E-112 | 1,31965574 | 0,344 | 0,019 | 3,38E-107 | C2_NID2 | LY6H | 1,90386079 | 3,86212073 |
| 2,90E-111 | 1,234329222 | 0,599 | 0,26 | 1,02E-106 | C2_NID2 | APOD | 1,78076065 | 1,18855619 |
| 1,21E-110 | 0,981425447 | 0,713 | 0,322 | 4,27E-106 | C2_NID2 | PXDN | 1,41589763 | 1,13469321 |
| 2,15E-87 | 0,928723284 | 0,628 | 0,225 | 7,55E-83 | C2_NID2 | TP53I11 | 1,33986448 | 1,46057164 |
| 1,07E-76 | 0,769417077 | 0,748 | 0,443 | 3,77E-72 | C2_NID2 | FSCN1 | 1,1100342 | 0,74915113 |
| 1,85E-72 | 0,704964287 | 0,398 | 0,097 | 6,50E-68 | C2_NID2 | CHST1 | 1,01704848 | 1,98221069 |
| 2,09E-67 | 0,713558486 | 0,347 | 0,073 | 7,36E-63 | C2_NID2 | NID2 | 1,02944729 | 2,1740294 |
| 1,03E-60 | 0,764619016 | 0,708 | 0,358 | 3,63E-56 | C2_NID2 | LAMA4 | 1,10311206 | 0,97393253 |
| 1,46E-60 | 0,636086182 | 0,579 | 0,239 | 5,12E-56 | C2_NID2 | TNFAIP8L1 | 0,91767838 | 1,25908722 |
| 1,70E-59 | 0,525821292 | 0,441 | 0,118 | 5,98E-55 | C2_NID2 | CHST15 | 0,75859977 | 1,85838539 |
| 9,54E-59 | 0,452132461 | 0,264 | 0,023 | 3,36E-54 | C2_NID2 | KIT | 0,65228926 | 3,26410744 |
| 2,43E-58 | 0,714665312 | 0,585 | 0,253 | 8,54E-54 | C2_NID2 | CTHRC1 | 1,0310441 | 1,19334389 |
| 1,16E-57 | 0,646485289 | 0,55 | 0,192 | 4,07E-53 | C2_NID2 | ANGPTL2 | 0,93268112 | 1,49429214 |
| 4,32E-52 | 0,709587423 | 0,53 | 0,187 | 1,52E-47 | C2_NID2 | GPIHBP1 | 1,02371826 | 1,47843258 |
| 5,55E-52 | 0,585649907 | 0,567 | 0,266 | 1,95E-47 | C2_NID2 | LXN | 0,84491422 | 1,0777223 |
| 6,15E-47 | 0,788509302 | 0,602 | 0,294 | 2,16E-42 | C2_NID2 | TNFRSF4 | 1,13757846 | 1,02155103 |
| 1,14E-46 | 0,743682341 | 0,736 | 0,475 | 4,00E-42 | C2_NID2 | MCAM | 1,07290682 | 0,62643914 |
| 1,22E-46 | 0,45917978 | 0,347 | 0,091 | 4,31E-42 | C2_NID2 | FAM129A | 0,66245639 | 1,87446912 |
| 4,96E-44 | 0,455732281 | 0,387 | 0,11 | 1,74E-39 | C2_NID2 | HOMER3 | 0,6574827 | 1,76921979 |
| 9,13E-42 | 0,548905258 | 0,679 | 0,418 | 3,21E-37 | C2_NID2 | LAMC1 | 0,79190289 | 0,69333866 |
| 3,89E-40 | 0,771655 | 0,622 | 0,3 | 1,37E-35 | C2_NID2 | ADM | 1,11326284 | 1,0396562 |
| 6,93E-40 | 0,579205078 | 0,39 | 0,107 | 2,44E-35 | C2_NID2 | ACKR3 | 0,83561629 | 1,81835392 |
| 1,59E-38 | 0,550958099 | 0,178 | 0,018 | 5,60E-34 | C2_NID2 | PRND | 0,79486452 | 2,99213788 |
| 2,56E-38 | 0,412932734 | 0,341 | 0,1 | 8,99E-34 | C2_NID2 | WFS1 | 0,59573601 | 1,72038271 |
| 5,47E-38 | 0,546326724 | 0,521 | 0,254 | 1,93E-33 | C2_NID2 | SMTN | 0,78818285 | 1,0221107 |
| 1,11E-37 | 0,491106106 | 0,576 | 0,283 | 3,91E-33 | C2_NID2 | NDST1 | 0,70851634 | 1,01246935 |
| 8,21E-36 | 0,920726047 | 0,453 | 0,208 | 2,89E-31 | C2_NID2 | IGF2 | 1,3283269 | 1,10449417 |
| 5,32E-35 | 0,591488465 | 0,736 | 0,459 | 1,87E-30 | C2_NID2 | MMP2 | 0,85333747 | 0,67534874 |
| 6,84E-35 | 0,673068554 | 0,602 | 0,375 | 2,41E-30 | C2_NID2 | RGS3 | 0,97103266 | 0,6756971 |
| 1,15E-34 | 0,464814333 | 0,493 | 0,224 | 4,04E-30 | C2_NID2 | TSPAN15 | 0,67058533 | 1,12079814 |
| 4,73E-34 | 0,552199209 | 0,742 | 0,475 | 1,66E-29 | C2_NID2 | MARCKS | 0,79665506 | 0,63807384 |
| 1,61E-31 | 0,569190172 | 0,401 | 0,179 | 5,66E-27 | C2_NID2 | SLC4A7 | 0,82116784 | 1,14177396 |
| 1,97E-29 | 0,44229115 | 0,564 | 0,292 | 6,94E-25 | C2_NID2 | ARHGAP18 | 0,63809125 | 0,93796572 |
| 6,22E-29 | 0,476507401 | 0,599 | 0,349 | 2,19E-24 | C2_NID2 | FILIP1 | 0,68745486 | 0,77079919 |
| 1,65E-28 | 0,683983839 | 0,562 | 0,352 | 5,79E-24 | C2_NID2 | CXCR4 | 0,98678009 | 0,66742466 |
| 6,91E-27 | 0,425463643 | 0,567 | 0,314 | 2,43E-22 | C2_NID2 | JUP | 0,61381429 | 0,84245872 |
| 2,93E-26 | 0,424137675 | 0,516 | 0,286 | 1,03E-21 | C2_NID2 | SMAD1 | 0,61190132 | 0,84026422 |
| 3,04E-25 | 0,445174347 | 0,544 | 0,307 | 1,07E-20 | C2_NID2 | FAM43A | 0,64225082 | 0,81526012 |
| 1,14E-124 | 0,405871345 | 0,692 | 0,565 | 4,02E-120 | C3_CA4 | TMEM88 | 0,58554858 | 0,29019674 |
| 1,00E-117 | 0,607016872 | 0,771 | 0,623 | 3,53E-113 | C3_CA4 | RGCC | 0,87574023 | 0,30529209 |
| 4,50E-100 | 0,825398491 | 0,2 | 0,044 | 1,58E-95 | C3_CA4 | CA4 | 1,19079831 | 2,06477026 |
| 1,88E-99 | 0,554741767 | 0,601 | 0,498 | 6,60E-95 | C3_CA4 | ID2 | 0,8003232 | 0,26875939 |
| 1,87E-97 | 0,570135696 | 0,462 | 0,285 | 6,56E-93 | C3_CA4 | CD36 | 0,82253194 | 0,68736965 |
| 2,18E-97 | 0,351539733 | 0,725 | 0,676 | 7,65E-93 | C3_CA4 | PRMT1 | 0,50716463 | 0,10024167 |
| 8,43E-90 | 0,754884257 | 0,587 | 0,536 | 2,97E-85 | C3_CA4 | MT1X | 1,08906777 | 0,12996858 |
| 1,01E-86 | 0,408480983 | 0,676 | 0,616 | 3,56E-82 | C3_CA4 | SEPP1 | 0,58931349 | 0,13306153 |
| 9,74E-86 | 0,375144262 | 0,837 | 0,737 | 3,43E-81 | C3_CA4 | VAMP5 | 0,54121877 | 0,18240105 |
| 1,58E-84 | 0,399719918 | 0,568 | 0,446 | 5,55E-80 | C3_CA4 | GYPC | 0,57667394 | 0,34540771 |
| 8,24E-79 | 0,387575752 | 0,669 | 0,583 | 2,90E-74 | C3_CA4 | SGK1 | 0,55915362 | 0,19693244 |
| 1,98E-76 | 0,430631559 | 0,384 | 0,301 | 6,97E-72 | C3_CA4 | C11orf96 | 0,62127001 | 0,3462385 |
| 3,48E-75 | 0,758771462 | 0,402 | 0,287 | 1,22E-70 | C3_CA4 | MT1M | 1,09467582 | 0,47906043 |
| 1,48E-73 | 0,331238224 | 0,826 | 0,745 | 5,21E-69 | C3_CA4 | RAMP3 | 0,47787574 | 0,14795788 |
| 4,22E-70 | 0,327334107 | 0,364 | 0,26 | 1,49E-65 | C3_CA4 | OSBPL1A | 0,47224329 | 0,47762846 |
| 4,05E-67 | 0,439675396 | 0,643 | 0,611 | 1,42E-62 | C3_CA4 | CD320 | 0,63431751 | 0,07306346 |
| 1,90E-66 | 0,649046431 | 0,519 | 0,475 | 6,66E-62 | C3_CA4 | MT1E | 0,93637607 | 0,12653241 |
| 1,04E-65 | 0,393496097 | 0,723 | 0,632 | 3,65E-61 | C3_CA4 | ITM2A | 0,56769487 | 0,19264508 |
| 3,24E-64 | 0,325688047 | 0,582 | 0,466 | 1,14E-59 | C3_CA4 | TM4SF18 | 0,46986853 | 0,31763344 |
| 4,37E-64 | 0,816742241 | 0,386 | 0,22 | 1,54E-59 | C3_CA4 | FCN3 | 1,17830998 | 0,79724361 |
| 1,34E-61 | 0,38727918 | 0,182 | 0,067 | 4,70E-57 | C3_CA4 | PRX | 0,55872575 | 1,37696946 |
| 6,70E-61 | 0,361302748 | 0,497 | 0,399 | 2,35E-56 | C3_CA4 | F2RL3 | 0,52124968 | 0,31333207 |
| 3,68E-52 | 0,601528252 | 0,498 | 0,416 | 1,29E-47 | C3_CA4 | STC1 | 0,86782183 | 0,25673817 |
| 1,55E-48 | 0,424880485 | 0,6 | 0,547 | 5,44E-44 | C3_CA4 | HES1 | 0,61297297 | 0,13226688 |
| 1,77E-10 | 0,380932225 | 0,187 | 0,153 | 6,22E-06 | C3_CA4 | PLPP1 | 0,54956903 | 0,28118175 |
| 0 | 1,836346096 | 0,893 | 0,342 | 0 | C4_ACKR1 | ACKR1 | 2,64928741 | 1,37177978 |
| 1,63E-292 | 0,987387542 | 0,524 | 0,114 | 5,75E-288 | C4_ACKR1 | C7 | 1,42449911 | 2,15230615 |
| 8,47E-265 | 0,753370751 | 0,567 | 0,129 | 2,98E-260 | C4_ACKR1 | SELP | 1,08688425 | 2,09378215 |
| 6,02E-263 | 1,241354085 | 0,745 | 0,41 | 2,12E-258 | C4_ACKR1 | CLU | 1,79089538 | 0,85377926 |
| 8,75E-254 | 1,216702867 | 0,486 | 0,15 | 3,08E-249 | C4_ACKR1 | CPE | 1,75533119 | 1,66345481 |
| 2,26E-252 | 0,738843021 | 0,49 | 0,087 | 7,93E-248 | C4_ACKR1 | MMRN1 | 1,06592516 | 2,42772276 |
| 2,23E-245 | 0,521969116 | 0,362 | 0,037 | 7,85E-241 | C4_ACKR1 | RAB3C | 0,75304225 | 3,12731883 |
| 5,20E-226 | 0,874800109 | 0,698 | 0,302 | 1,83E-221 | C4_ACKR1 | CCL14 | 1,26206978 | 1,19528603 |
| 1,16E-197 | 0,641612977 | 0,566 | 0,206 | 4,07E-193 | C4_ACKR1 | IL1R1 | 0,92565186 | 1,43624775 |
| 5,03E-191 | 1,038491823 | 0,498 | 0,147 | 1,77E-186 | C4_ACKR1 | POSTN | 1,498227 | 1,72648708 |
| 4,04E-188 | 0,530209655 | 0,451 | 0,114 | 1,42E-183 | C4_ACKR1 | LRRC1 | 0,76493084 | 1,93807225 |
| 1,57E-179 | 0,55883308 | 0,519 | 0,176 | 5,53E-175 | C4_ACKR1 | HAPLN3 | 0,80622571 | 1,53357711 |
| 1,70E-166 | 0,423582189 | 0,27 | 0,035 | 5,99E-162 | C4_ACKR1 | GPR126 | 0,61109992 | 2,78135971 |
| 7,20E-159 | 0,631768676 | 0,665 | 0,406 | 2,53E-154 | C4_ACKR1 | CTSC | 0,91144954 | 0,7050227 |
| 5,55E-155 | 0,690924797 | 0,31 | 0,054 | 1,95E-150 | C4_ACKR1 | CCL23 | 0,99679378 | 2,41656497 |
| 6,01E-149 | 0,571248156 | 0,684 | 0,362 | 2,11E-144 | C4_ACKR1 | ZNF385D | 0,82413688 | 0,90872392 |
| 1,34E-148 | 0,525694721 | 0,762 | 0,437 | 4,71E-144 | C4_ACKR1 | TFPI | 0,75841717 | 0,79518021 |
| 3,55E-141 | 0,506306813 | 0,785 | 0,504 | 1,25E-136 | C4_ACKR1 | ACTN1 | 0,73044633 | 0,634187 |
| 2,51E-135 | 0,646481035 | 0,7 | 0,464 | 8,83E-131 | C4_ACKR1 | NCOA7 | 0,93267498 | 0,58803533 |
| 1,31E-123 | 0,602826642 | 0,599 | 0,301 | 4,62E-119 | C4_ACKR1 | NR2F2 | 0,86969501 | 0,9810169 |
| 1,30E-120 | 0,479135438 | 0,518 | 0,22 | 4,57E-116 | C4_ACKR1 | HLA-DQA1 | 0,69124632 | 1,21688595 |
| 2,97E-120 | 0,651205683 | 0,28 | 0,065 | 1,04E-115 | C4_ACKR1 | MADCAM1 | 0,93949121 | 2,02553509 |
| 2,29E-117 | 0,596157198 | 0,38 | 0,167 | 8,05E-113 | C4_ACKR1 | TMEM176A | 0,86007303 | 1,16244988 |
| 1,03E-116 | 0,440081504 | 0,7 | 0,38 | 3,63E-112 | C4_ACKR1 | IL33 | 0,6349034 | 0,87276481 |

| | | | | | | | | |
|---|---|---|---|---|---|---|---|---|
| 1,82E-112 | 0,574972438 | 0,391 | 0,185 | 6,41E-108 | C4_ACKR1 | TMEM176B | 0,82950989 | 1,05950101 |
| 1,48E-95 | 0,71002946 | 0,392 | 0,172 | 5,20E-91 | C4_ACKR1 | PTGDS | 1,02435598 | 1,16538965 |
| 3,00E-86 | 0,438067143 | 0,453 | 0,215 | 1,06E-81 | C4_ACKR1 | CNKSR3 | 0,63199729 | 1,05784407 |
| 3,34E-84 | 0,826268016 | 0,426 | 0,206 | 1,18E-79 | C4_ACKR1 | AKAP12 | 1,19205277 | 1,03044487 |
| 3,93E-74 | 0,507814774 | 0,309 | 0,108 | 1,38E-69 | C4_ACKR1 | VCAM1 | 0,73262186 | 1,47444179 |
| 5,27E-55 | 0,644786248 | 0,24 | 0,083 | 1,85E-50 | C4_ACKR1 | SELE | 0,93022992 | 1,47720632 |
| 8,02E-50 | 0,661410683 | 0,218 | 0,091 | 2,82E-45 | C4_ACKR1 | IL6 | 0,95421391 | 1,2159374 |
| 4,18E-30 | 0,484274517 | 0,104 | 0,04 | 1,47E-25 | C4_ACKR1 | CSF3 | 0,69866044 | 1,27633123 |
| 4,57E-19 | 0,413509518 | 0,135 | 0,062 | 1,61E-14 | C4_ACKR1 | EFEMP1 | 0,59656813 | 1,06319383 |
| 0 | 1,442011763 | 0,69 | 0,075 | 0 | C5_FBLN5 | GJA5 | 2,08038322 | 3,11894107 |
| 0 | 1,439522595 | 0,708 | 0,06 | 0 | C5_FBLN5 | FBLN5 | 2,07679211 | 3,45539045 |
| 0 | 1,100438656 | 0,704 | 0,103 | 0 | C5_FBLN5 | SEMA3G | 1,58759739 | 2,71475432 |
| 1,56E-226 | 0,87567591 | 0,683 | 0,292 | 5,50E-222 | C5_FBLN5 | NUDT4 | 1,26333329 | 1,21194563 |
| 1,85E-226 | 1,144443104 | 0,722 | 0,222 | 6,50E-222 | C5_FBLN5 | GJA4 | 1,65108239 | 1,67926307 |
| 8,09E-224 | 1,48168079 | 0,789 | 0,341 | 2,84E-219 | C5_FBLN5 | IGFBP3 | 2,13761353 | 1,19836697 |
| 1,02E-212 | 0,794654213 | 0,729 | 0,289 | 3,60E-208 | C5_FBLN5 | ARL15 | 1,14644369 | 1,31996391 |
| 6,48E-203 | 0,927972849 | 0,742 | 0,258 | 2,28E-198 | C5_FBLN5 | HEY1 | 1,33878183 | 1,50604544 |
| 2,28E-199 | 0,61496184 | 0,423 | 0,039 | 8,01E-195 | C5_FBLN5 | TSPAN2 | 0,8872024 | 3,28203537 |
| 5,79E-198 | 0,985925755 | 0,746 | 0,399 | 2,04E-193 | C5_FBLN5 | KCTD12 | 1,4223902 | 0,89445761 |
| 2,43E-194 | 0,90295831 | 0,809 | 0,533 | 8,53E-190 | C5_FBLN5 | STMN1 | 1,30269348 | 0,59742262 |
| 1,43E-189 | 0,908383393 | 0,435 | 0,065 | 5,03E-185 | C5_FBLN5 | PCSK5 | 1,31052022 | 2,6520767 |
| 1,63E-186 | 0,912845685 | 0,51 | 0,134 | 5,73E-182 | C5_FBLN5 | LTBP4 | 1,31695794 | 1,88948755 |
| 2,62E-186 | 0,797155617 | 0,821 | 0,452 | 9,22E-182 | C5_FBLN5 | EFNB2 | 1,15005246 | 0,85394762 |
| 8,85E-169 | 0,564427968 | 0,49 | 0,083 | 3,11E-164 | C5_FBLN5 | PLLP | 0,81429743 | 2,4918531 |
| 5,66E-159 | 0,663362671 | 0,729 | 0,444 | 1,99E-154 | C5_FBLN5 | PDCD4 | 0,95703004 | 0,70906462 |
| 4,09E-156 | 0,686613142 | 0,738 | 0,445 | 1,44E-151 | C5_FBLN5 | OCIAD2 | 0,99057337 | 0,72343721 |
| 6,73E-148 | 1,16314629 | 0,612 | 0,247 | 2,37E-143 | C5_FBLN5 | ENPP2 | 1,67806538 | 1,29184676 |
| 2,82E-147 | 0,612571146 | 0,545 | 0,164 | 9,93E-143 | C5_FBLN5 | SYNJ2 | 0,88375336 | 1,70240837 |
| 2,68E-135 | 0,760971068 | 0,317 | 0,052 | 9,41E-131 | C5_FBLN5 | SULF1 | 1,09784919 | 2,49802686 |
| 9,35E-133 | 0,660160339 | 0,674 | 0,339 | 3,29E-128 | C5_FBLN5 | MECOM | 0,95241005 | 0,98100301 |
| 5,56E-130 | 0,863719325 | 0,506 | 0,144 | 1,96E-125 | C5_FBLN5 | FBLN2 | 1,24608359 | 1,77801096 |
| 4,57E-123 | 0,915097833 | 0,762 | 0,444 | 1,61E-118 | C5_FBLN5 | ADAMTS1 | 1,32020711 | 0,77251113 |
| 3,49E-115 | 0,832496511 | 0,673 | 0,299 | 1,23E-110 | C5_FBLN5 | CXCL12 | 1,20103859 | 1,15721395 |
| 3,34E-113 | 0,827829421 | 0,322 | 0,061 | 1,17E-108 | C5_FBLN5 | SERPINE2 | 1,1943054 | 2,30875271 |
| 6,47E-109 | 0,786683248 | 0,182 | 0,008 | 2,27E-104 | C5_FBLN5 | DKK2 | 1,13494402 | 3,84645474 |
| 2,01E-106 | 0,572885309 | 0,477 | 0,153 | 7,06E-102 | C5_FBLN5 | VEGFC | 0,82649879 | 1,60910859 |
| 7,46E-106 | 0,638759212 | 0,505 | 0,176 | 2,62E-101 | C5_FBLN5 | PPP1R14A | 0,92153475 | 1,49450755 |
| 1,99E-103 | 0,609689837 | 0,434 | 0,151 | 7,00E-99 | C5_FBLN5 | ELN | 0,8795965 | 1,49267491 |
| 4,33E-99 | 0,704060039 | 0,582 | 0,324 | 1,52E-94 | C5_FBLN5 | JAG1 | 1,01574393 | 0,83527292 |
| 2,62E-94 | 0,435810636 | 0,329 | 0,072 | 9,23E-90 | C5_FBLN5 | BMX | 0,62874184 | 2,11691775 |
| 1,85E-93 | 0,5825975 | 0,613 | 0,293 | 6,49E-89 | C5_FBLN5 | DLL4 | 0,84051052 | 1,05229451 |
| 1,42E-91 | 0,584455302 | 0,446 | 0,171 | 4,98E-87 | C5_FBLN5 | FILIP1L | 0,84319077 | 1,357552 |
| 9,46E-90 | 0,585493126 | 0,272 | 0,047 | 3,33E-85 | C5_FBLN5 | TMEM100 | 0,84468803 | 2,41330245 |
| 1,42E-87 | 0,499763729 | 0,649 | 0,375 | 5,01E-83 | C5_FBLN5 | EMP3 | 0,72100665 | 0,78329122 |
| 1,67E-85 | 0,588259473 | 0,54 | 0,312 | 5,89E-81 | C5_FBLN5 | SLC26A2 | 0,84867902 | 0,78177339 |
| 1,65E-84 | 0,476765859 | 0,451 | 0,167 | 5,81E-80 | C5_FBLN5 | FBLIM1 | 0,68782774 | 1,40662526 |
| 1,63E-81 | 0,506290938 | 0,421 | 0,14 | 5,74E-77 | C5_FBLN5 | SLC14A1 | 0,73042343 | 1,55480053 |
| 4,03E-81 | 0,4227705 | 0,421 | 0,141 | 1,42E-76 | C5_FBLN5 | ARHGAP4 | 0,6099289 | 1,54488506 |
| 3,32E-79 | 0,632987329 | 0,505 | 0,181 | 1,17E-74 | C5_FBLN5 | IGF2 | 0,91320768 | 1,45519463 |
| 5,33E-84 | 0,5219189 | 0,792 | 0,608 | 1,87E-79 | C6_APLNR | PRCP | 0,75296981 | 0,37869265 |
| 1,12E-76 | 0,611587711 | 0,559 | 0,338 | 3,93E-72 | C6_APLNR | APLNR | 0,88233456 | 0,71748659 |
| 4,21E-73 | 0,538102412 | 0,736 | 0,576 | 1,48E-68 | C6_APLNR | PCDH17 | 0,77631768 | 0,35093538 |
| 3,73E-66 | 0,43226026 | 0,805 | 0,725 | 1,31E-61 | C6_APLNR | ZFP36L1 | 0,62361973 | 0,15002544 |
| 1,68E-56 | 0,884080181 | 0,363 | 0,212 | 5,92E-52 | C6_APLNR | IGFBP5 | 1,27545809 | 0,76201072 |
| 1,26E-51 | 0,426558799 | 0,688 | 0,585 | 4,43E-47 | C6_APLNR | TGM2 | 0,61539426 | 0,2321404 |
| 1,95E-48 | 0,366870332 | 0,638 | 0,483 | 6,85E-44 | C6_APLNR | TMEM2 | 0,52928201 | 0,39793759 |
| 2,42E-41 | 0,333363843 | 0,604 | 0,468 | 8,52E-37 | C6_APLNR | RAI14 | 0,48094236 | 0,36460204 |
| 5,04E-40 | 0,35155988 | 0,657 | 0,552 | 1,77E-35 | C6_APLNR | TSHZ2 | 0,50719369 | 0,24915389 |
| 2,36E-39 | 0,331175412 | 0,647 | 0,526 | 8,32E-35 | C6_APLNR | ENTPD1 | 0,47778512 | 0,2961601 |
| 5,32E-24 | 0,419246011 | 0,696 | 0,611 | 1,87E-19 | C6_APLNR | MT-ND4L | 0,60484414 | 0,18648409 |
| 4,07E-15 | 0,365774598 | 0,479 | 0,368 | 1,43E-10 | C6_APLNR | ICAM1 | 0,5277012 | 0,37583142 |
| 3,91E-08 | 0,678724188 | 0,213 | 0,17 | 0,00137517 | C6_APLNR | MTRNR2L12 | 0,97919202 | 0,31697321 |
| 1,97E-07 | 0,400166433 | 0,18 | 0,138 | 0,00691432 | C6_APLNR | MT-ATP8 | 0,57731813 | 0,37151012 |
| 3,38E-156 | 0,971215363 | 0,925 | 0,567 | 1,19E-151 | C7_SSUH2 | GLUL | 1,40116759 | 0,70121557 |
| 3,28E-132 | 0,904751418 | 0,471 | 0,042 | 1,15E-127 | C7_SSUH2 | SSUH2 | 1,30528038 | 3,34022891 |
| 1,57E-115 | 1,194482251 | 0,826 | 0,312 | 5,52E-111 | C7_SSUH2 | CXCL12 | 1,72327362 | 1,39036564 |
| 4,90E-106 | 1,531357554 | 0,753 | 0,324 | 1,72E-101 | C7_SSUH2 | FABP4 | 2,20928195 | 1,20411026 |
| 4,70E-85 | 0,831845823 | 0,654 | 0,206 | 1,65E-80 | C7_SSUH2 | SLC6A6 | 1,20009984 | 1,64303547 |
| 3,05E-83 | 0,872465825 | 0,601 | 0,183 | 1,07E-78 | C7_SSUH2 | ADAMTS6 | 1,25870212 | 1,68858513 |
| 4,70E-80 | 0,816321408 | 0,736 | 0,252 | 1,65E-75 | C7_SSUH2 | GJA4 | 1,17770285 | 1,52770518 |
| 1,89E-79 | 0,614998449 | 0,731 | 0,265 | 6,65E-75 | C7_SSUH2 | AIF1L | 0,88725521 | 1,44674636 |
| 4,76E-77 | 0,648429004 | 0,665 | 0,253 | 1,67E-72 | C7_SSUH2 | CLDN15 | 0,93548531 | 1,37679003 |
| 7,26E-74 | 0,973561527 | 0,879 | 0,574 | 2,55E-69 | C7_SSUH2 | RBP7 | 1,40455239 | 0,61048302 |
| 6,39E-65 | 0,559176539 | 0,795 | 0,368 | 2,25E-60 | C7_SSUH2 | BTNL9 | 0,80672122 | 1,10082437 |
| 4,16E-60 | 0,522662291 | 0,707 | 0,321 | 1,46E-55 | C7_SSUH2 | EFNB1 | 0,7540423 | 1,12700528 |
| 7,63E-57 | 0,437926531 | 0,269 | 0,029 | 2,68E-52 | C7_SSUH2 | TMEM178A | 0,63179443 | 3,01056924 |
| 2,70E-55 | 0,817217254 | 0,573 | 0,197 | 9,50E-51 | C7_SSUH2 | IGF2 | 1,17899528 | 1,5167142 |
| 3,47E-49 | 0,529625336 | 0,789 | 0,508 | 1,22E-44 | C7_SSUH2 | PLS3 | 0,76408785 | 0,63018018 |
| 4,36E-48 | 0,541338972 | 0,681 | 0,292 | 1,53E-43 | C7_SSUH2 | HEY1 | 0,78098705 | 1,20774565 |
| 5,16E-47 | 0,514015622 | 0,628 | 0,28 | 1,81E-42 | C7_SSUH2 | ASRGL1 | 0,74156779 | 1,15124358 |
| 7,13E-47 | 0,588681064 | 0,52 | 0,208 | 2,51E-42 | C7_SSUH2 | ATP13A3 | 0,84928725 | 1,30146399 |
| 1,26E-36 | 0,456391797 | 0,744 | 0,415 | 4,42E-32 | C7_SSUH2 | ASS1 | 0,65843418 | 0,83457639 |
| 1,30E-36 | 0,423136093 | 0,458 | 0,179 | 4,56E-32 | C7_SSUH2 | PPY | 0,61045634 | 1,33130643 |
| 1,78E-35 | 0,436678448 | 0,802 | 0,506 | 6,27E-31 | C7_SSUH2 | UACA | 0,62999383 | 0,65924538 |
| 2,50E-35 | 0,492739033 | 0,797 | 0,509 | 8,78E-31 | C7_SSUH2 | MAST4 | 0,71087216 | 0,64183388 |
| 3,52E-28 | 0,509048056 | 0,711 | 0,397 | 1,24E-23 | C7_SSUH2 | GPR116 | 0,73440111 | 0,83276409 |
| 8,89E-28 | 0,442580376 | 0,434 | 0,161 | 3,13E-23 | C7_SSUH2 | ALPL | 0,63850851 | 1,4030377 |
| 1,34E-27 | 0,496186393 | 0,463 | 0,199 | 4,72E-23 | C7_SSUH2 | PPP1R14A | 0,71584565 | 1,19793938 |
| 3,80E-24 | 0,444216183 | 0,637 | 0,329 | 1,34E-19 | C7_SSUH2 | CRIP1 | 0,64086848 | 0,94272519 |
| 5,53E-23 | 0,433588548 | 0,615 | 0,307 | 1,95E-18 | C7_SSUH2 | IGLL5 | 0,62553605 | 0,99072219 |
| 6,47E-17 | 0,515877827 | 0,696 | 0,46 | 2,28E-12 | C7_SSUH2 | RCAN1 | 0,74425438 | 0,59218373 |
| 3,33E-228 | 4,842770513 | 0,874 | 0,103 | 1,17E-223 | C8_PROX1 | CCL21 | 6,986641 | 3,02483185 |
| 6,02E-135 | 2,301967455 | 0,825 | 0,27 | 2,12E-130 | C8_PROX1 | TFF3 | 3,32103703 | 1,59367972 |
| 8,38E-92 | 1,699831067 | 0,602 | 0,012 | 2,95E-87 | C8_PROX1 | PDPN | 2,45233785 | 5,15808987 |
| 3,78E-76 | 1,600600652 | 0,883 | 0,499 | 1,33E-71 | C8_PROX1 | TFPI | 2,30917862 | 0,81713594 |
| 9,89E-70 | 1,333049876 | 0,544 | 0,296 | 3,48E-65 | C8_PROX1 | RBP1 | 1,92318445 | 0,86704266 |
| 1,87E-67 | 1,323076728 | 0,524 | 0,039 | 6,58E-63 | C8_PROX1 | PROX1 | 1,90879623 | 3,58769229 |
| 1,23E-58 | 0,999248136 | 0,359 | 0,003 | 4,34E-54 | C8_PROX1 | RELN | 1,44161033 | 5,50779464 |
| 2,51E-57 | 1,052830023 | 0,427 | 0,016 | 8,82E-53 | C8_PROX1 | IGF1 | 1,51891265 | 4,36257008 |
| 1,92E-56 | 1,638117797 | 0,738 | 0,162 | 6,76E-52 | C8_PROX1 | MMRN1 | 2,36330442 | 2,15351411 |
| 1,46E-53 | 1,101210429 | 0,631 | 0,322 | 5,14E-49 | C8_PROX1 | LAPTM5 | 1,58871082 | 0,95973613 |
| 2,50E-49 | 0,880249274 | 0,194 | 0,051 | 8,80E-45 | C8_PROX1 | MFAP4 | 1,26993126 | 1,8292697 |
| 4,29E-49 | 0,854422078 | 0,427 | 0,029 | 1,51E-44 | C8_PROX1 | SEMA3A | 1,23267049 | 3,66742466 |
| 5,17E-48 | 2,221045301 | 0,524 | 0,071 | 1,82E-43 | C8_PROX1 | EFEMP1 | 3,20429104 | 2,7991964 |
| 2,53E-46 | 1,076794565 | 0,748 | 0,458 | 8,90E-42 | C8_PROX1 | PPFIBP1 | 1,55348618 | 0,70163767 |
| 1,66E-45 | 0,922705559 | 0,379 | 0,013 | 5,84E-41 | C8_PROX1 | TBX1 | 1,33118273 | 4,4150375 |
| 7,68E-41 | 0,422888411 | 0,252 | 0,002 | 2,70E-36 | C8_PROX1 | ACKR2 | 0,61009901 | 5,19826963 |
| 8,69E-41 | 0,970324452 | 0,398 | 0,024 | 3,06E-36 | C8_PROX1 | MRC1 | 1,39988227 | 3,79665503 |
| 1,61E-38 | 0,94520386 | 0,709 | 0,457 | 5,67E-34 | C8_PROX1 | LIMS1 | 1,36364092 | 0,62803122 |
| 1,21E-34 | 0,567930264 | 0,272 | 0,009 | 4,26E-30 | C8_PROX1 | PARD6G | 0,81935017 | 4,30638724 |
| 1,56E-34 | 0,700244121 | 0,301 | 0,015 | 5,50E-30 | C8_PROX1 | EFNA5 | 1,01023872 | 3,93545975 |
| 8,01E-33 | 0,819405788 | 0,369 | 0,039 | 2,82E-28 | C8_PROX1 | NR2F1 | 1,18215267 | 3,08746284 |
| 1,78E-32 | 0,474650357 | 0,252 | 0,005 | 6,25E-28 | C8_PROX1 | SEMA3D | 0,68477572 | 4,68369645 |
| 7,45E-32 | 0,750464643 | 0,408 | 0,055 | 2,62E-27 | C8_PROX1 | TSPAN5 | 1,08269162 | 2,78310738 |
| 1,46E-30 | 0,742875022 | 0,32 | 0,082 | 5,15E-26 | C8_PROX1 | ITM2C | 1,07174211 | 1,90135241 |
| 2,26E-30 | 0,477899104 | 0,301 | 0,018 | 7,96E-26 | C8_PROX1 | RP11-322E11 | 0,68946267 | 3,73382589 |
| 3,79E-30 | 0,721332497 | 0,388 | 0,074 | 1,33E-25 | C8_PROX1 | SLC38A1 | 1,04066282 | 2,31460475 |
| 1,24E-29 | 0,563781086 | 0,194 | 0,002 | 4,37E-25 | C8_PROX1 | ADGRG3 | 0,81336418 | 4,8292697 |
| 5,17E-28 | 0,971636149 | 0,388 | 0,126 | 1,82E-23 | C8_PROX1 | MAF | 1,40177465 | 1,5849625 |
| 6,77E-28 | 0,764338766 | 0,476 | 0,171 | 2,38E-23 | C8_PROX1 | LAYN | 1,10270775 | 1,45046147 |
| 1,24E-27 | 0,817861311 | 0,437 | 0,13 | 4,34E-23 | C8_PROX1 | CCDC80 | 1,17992446 | 1,71108696 |
| 3,47E-27 | 0,696313937 | 0,301 | 0,125 | 1,22E-22 | C8_PROX1 | MFAP2 | 1,00456866 | 1,23502003 |
| 5,92E-27 | 0,600297627 | 0,272 | 0,035 | 2,08E-22 | C8_PROX1 | SCN3B | 0,86604641 | 2,79181407 |
| 4,68E-26 | 0,635733101 | 0,291 | 0,038 | 1,65E-21 | C8_PROX1 | DOCK5 | 0,91716899 | 2,78318861 |
| 9,40E-25 | 0,797885537 | 0,515 | 0,287 | 3,30E-20 | C8_PROX1 | TSTA3 | 1,15110551 | 0,83254325 |
| 1,29E-24 | 0,509563384 | 0,282 | 0,044 | 4,55E-20 | C8_PROX1 | SLC24A1 | 0,73514457 | 2,55019708 |
| 4,02E-24 | 0,726197397 | 0,282 | 0,063 | 1,41E-19 | C8_PROX1 | C17orf67 | 1,04768138 | 2,07744409 |
| 4,22E-24 | 0,538549868 | 0,282 | 0,022 | 1,48E-19 | C8_PROX1 | COLEC12 | 0,77696322 | 3,41001942 |
| 7,16E-24 | 0,739007804 | 0,398 | 0,176 | 2,52E-19 | C8_PROX1 | RARRES2 | 1,06616289 | 1,15479014 |
| 5,04E-21 | 0,55229425 | 0,301 | 0,047 | 1,77E-16 | C8_PROX1 | DHODH | 0,79679218 | 2,55694812 |
| 5,78E-21 | 0,615347042 | 0,447 | 0,274 | 2,03E-16 | C8_PROX1 | MYL6B | 0,88775813 | 0,69605765 |
