## Supplementary table 2 for "Emerging single cell endothelial heterogeneity supports sprouting tumour angiogenesis and growth"

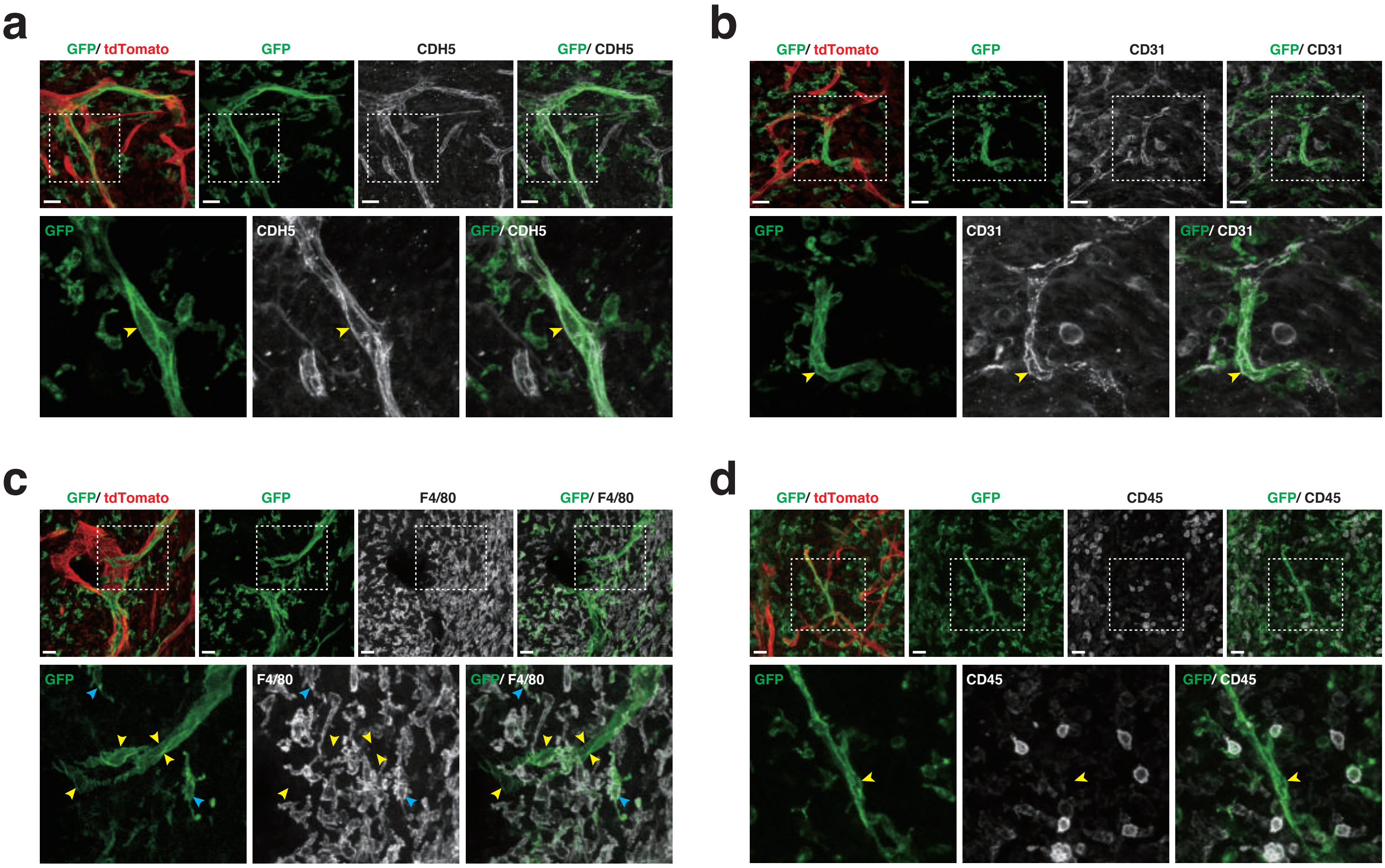


**Extended Data Fig. 1: CLECs in mouse glioma express endothelial cell markers, but lack macrophage and myeloid cell markers.** Counterstaining on PFA-fixed 4 weeks CT2A glioma sections of *Csf1r-Mer2.Cre*$^{mTmG}$ with indicated antibodies. Lower panels show the detail images of outlined squares in upper panels.Yellow arrowheads, CLECs; blue arrowheads, macrophages. Scale bar: 25 $\mu$m.

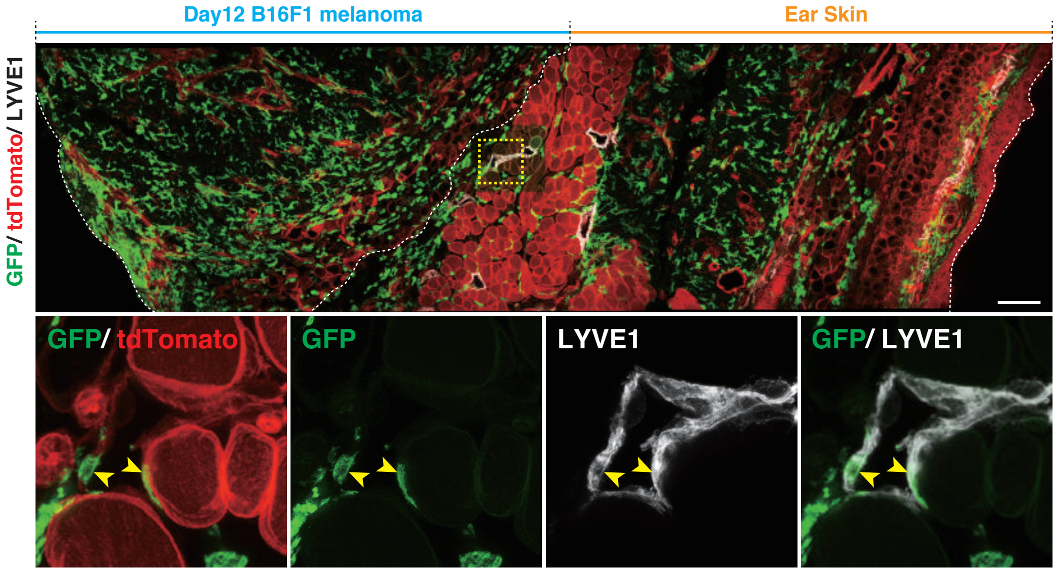


**Extended Data Fig. 2: CLECs contribute to lymphatic endothelium in the tumour environment.** Fluorescent images of counterstaining on PFA-fixed section of day 12 B16F1 melanoma under ear skin of *Csf1r-Mer2.Cre*$^{mTmG}$ mice using LYVE1 antibody. Lower panel indicates detail image of yellow outlined square in upper panel. Yellow arrow heads, CLECs. Scale bar: 100 $\mu$m.

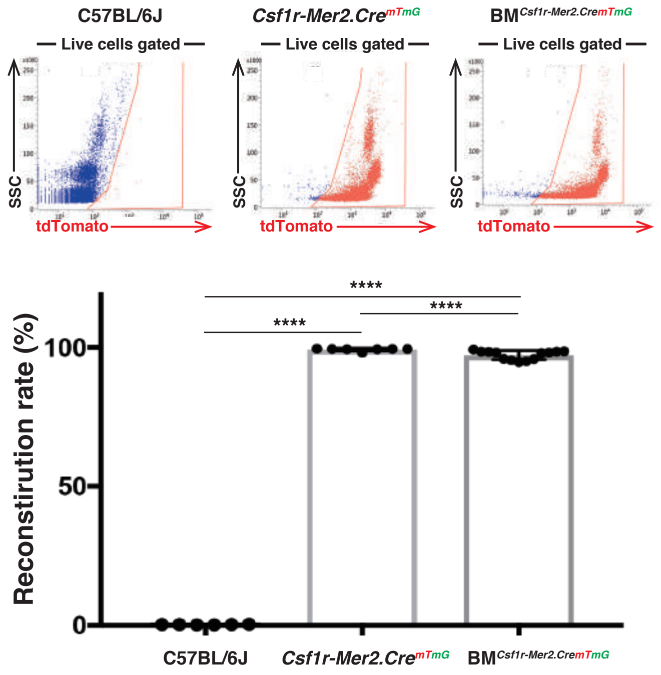


**Extended Data Fig. 3: Reconstitution rate in bone marrow chimeras.** Reconstitution rate was determined in peripheral blood 8 weeks after bone marrow transplantation. TdTomato-positive cells in blood of: C57BL/6J (0.3 ± 0.1%, n= 6), *Csf1r-Mer2.Cre*$^{mTmG}$ (99.2 ± 0.5%, n= 7), BM$^{Csf1r\text{-}Mer2.CremTmG}$ (97.3 ± 1.6%, n= 13). Bars represent mean ± s.d. ****P<0.0001. Two-tailed unpaired Mann-Whitney’ s U test.

**Matsumoto K et al., Extended Data Figure 3**

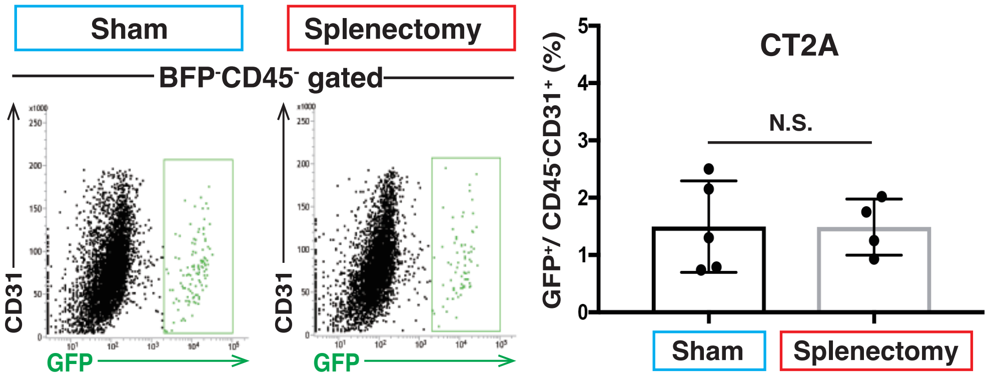


**Extended Data Fig. 4: CLECs do not originate from spleen.** Surgical removal of spleen (splenectomy) was performed 3 weeks before glioma injection. CLECs isolated from CT2A glioma of sham control *Csf1r-Mer2.Cre*$^{mTmG}$ mice (n= 5) constitute 1.5 ± 0.8 % of total tumour endothelial cell population, compared to 1.5 ± 0.5% found in *Csf1r-Mer2.Cre*$^{mTmG}$ with splenectomy (n= 4). Bars represent mean ± s.d. Two-tailed unpaired Mann-Whitney's U test.

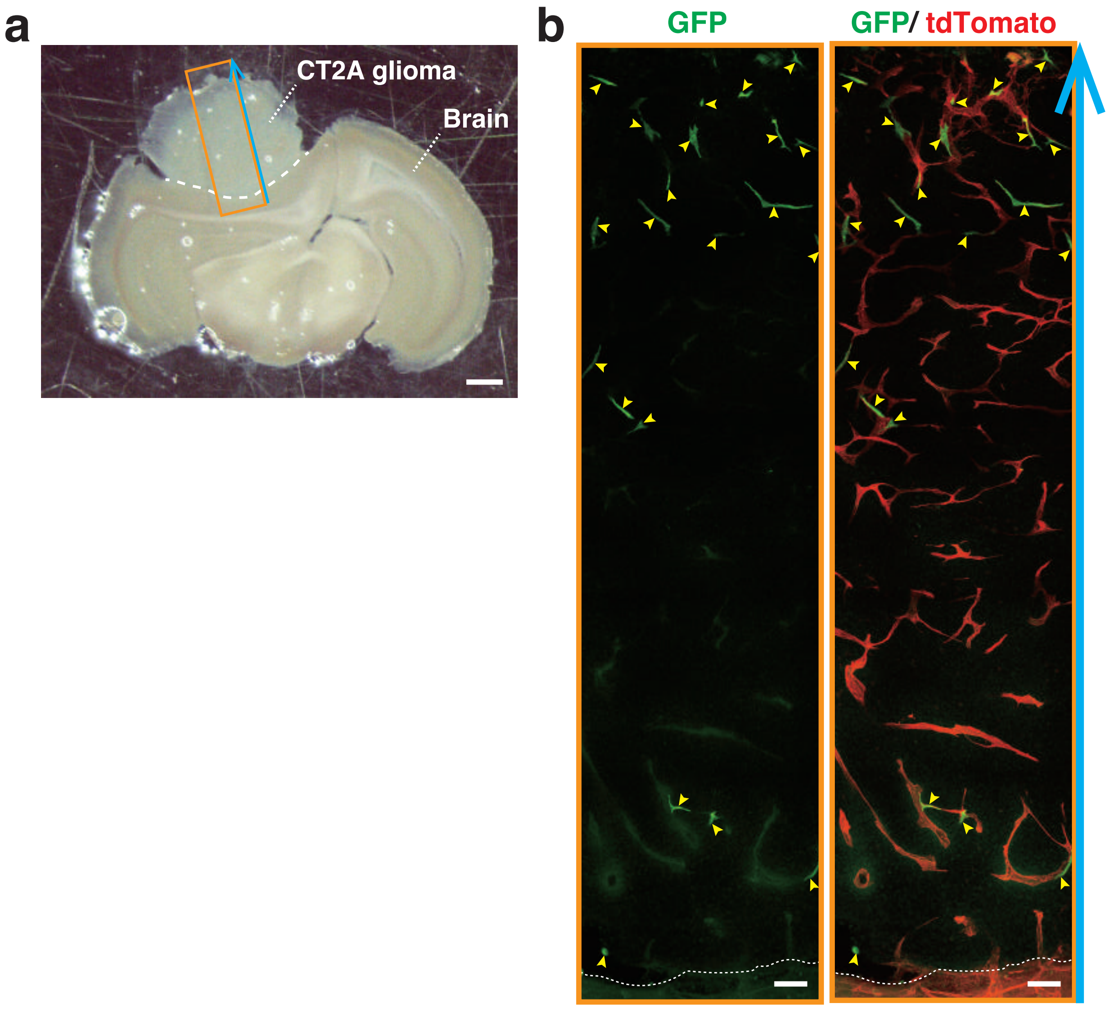


**Extended Data Fig. 5: CLECs locate at peripheral area of mouse glioma.** a, Vibratome section (200 $\mu$m-thick) of 4 weeks CT2A glioma of *Csf1r-Mer2.Cre*$^{mTmG}$:: $BM^{WT}$ mice. Scale bar: 1 mm. b, Fluorescent images of area outlined in a. Yellow arrowheads, CLECs. Scale bar: 100 $\mu$m.

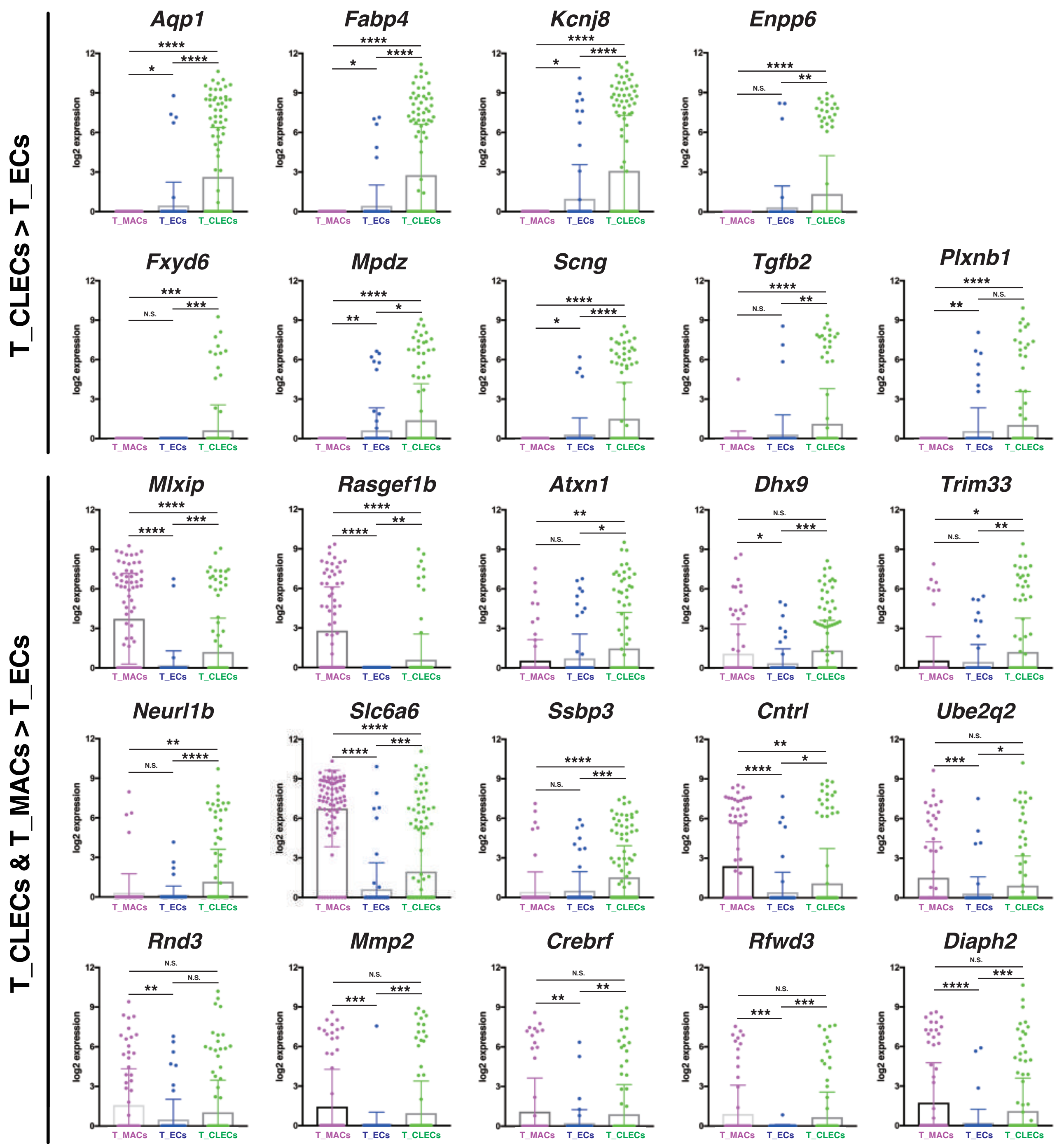


**Extended Data Fig. 6: 24 tumour CLECs markers in mouse glioma.** SCDE analysis of scRNA-seq data of tumour macrophages (T_MACs: $CD45^{+}GFP^{+}$, n= 79), tumour endothelial cells (T_ECs: $CD45^{-}CD31^{+}GFP^{-}$, n= 78), tumour CLECs (T_CLECs: $CD45^{-}CD31^{+}GFP^{+}$, n= 141) in 4 weeks CT2A glioma in *Csf1r-Mer2.Cre*$^{mTmG}$ mice. Bars represent mean ± s.d. *P<0.05,**P<0.01,***P<0.001, ****P<0.0001. Two-tailed unpaired t-test with Welch correction.

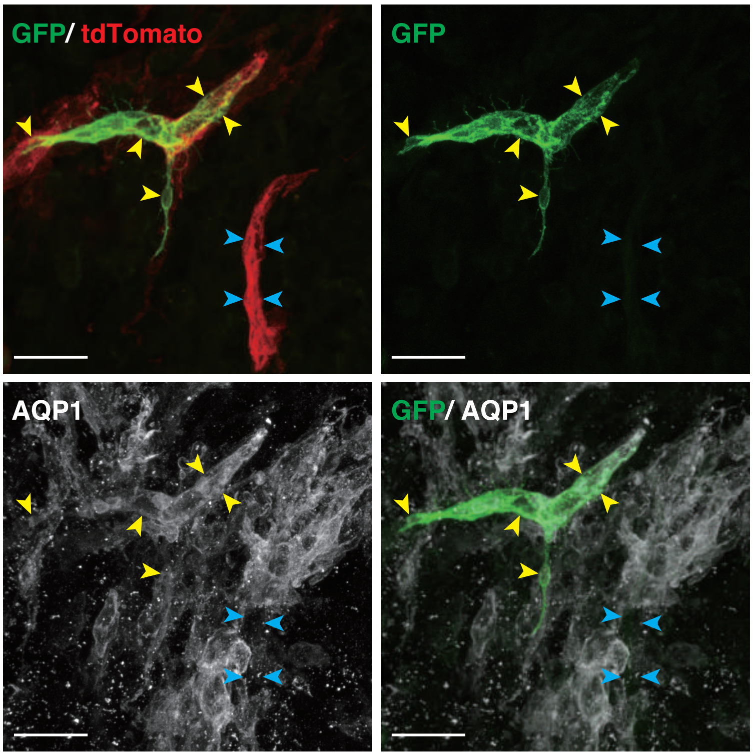


**Extended Data Fig. 7: CLECs in glioma express AQP1.** Counterstaining on PFA-fixed CT2A glioma of *Csf1r-Mer2.Cre*$^{mTmG}$::BM$^{WT}$ mice using AQP1 antibody. Yellow arrowheads, T_CLECs; blue arrowheads, T_ECs. Scale bar: 25 $\mu$m.

**Matsumoto K et al., Extended Data Figure 7**

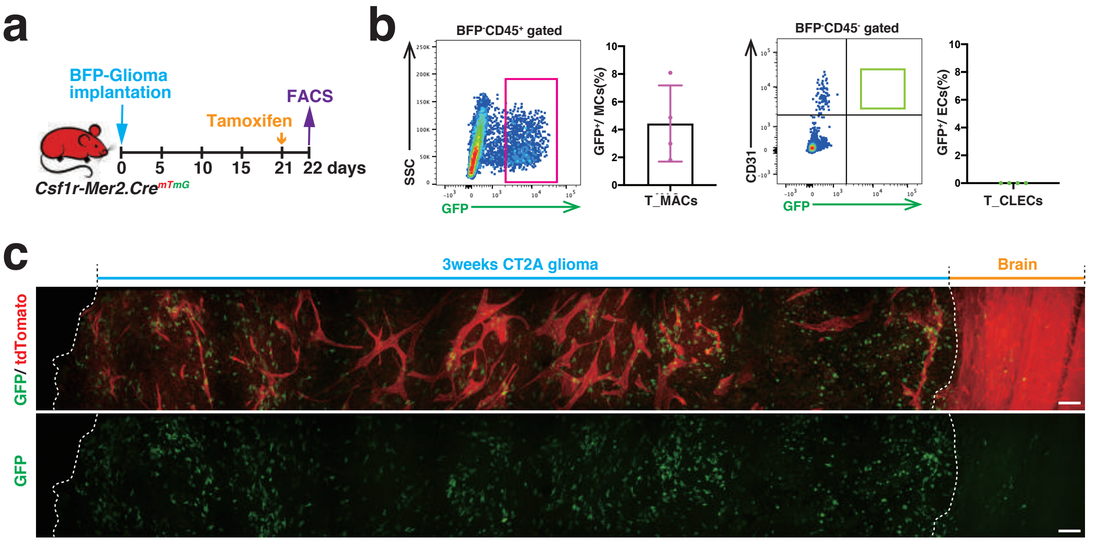


**Extended Data Fig. 8: CLECs in glioma cannot be labelled by GFP within 24 hours after tamoxifen induction.** a-b, Tamoxifen induction timing. b, Quantification of the population of tumour macrophagens (T_MACs: $CD45^{+}GFP^{+}$) and tumour CLECs (T_CLECs: $CD45^{-}CD31^{+}GFP^{+}$) by flow cytometric analysis in tumour samples (n=4). Bars represent mean ± s.d. c, Fluorescent images of PFA-fixed 3 weeks CT2A glioma de of *Csf1r-Mer2.Cre*$^{mTmG}$ mice within 24 hours after tamoxifen induction. Scale bar: 100 $\mu$m.

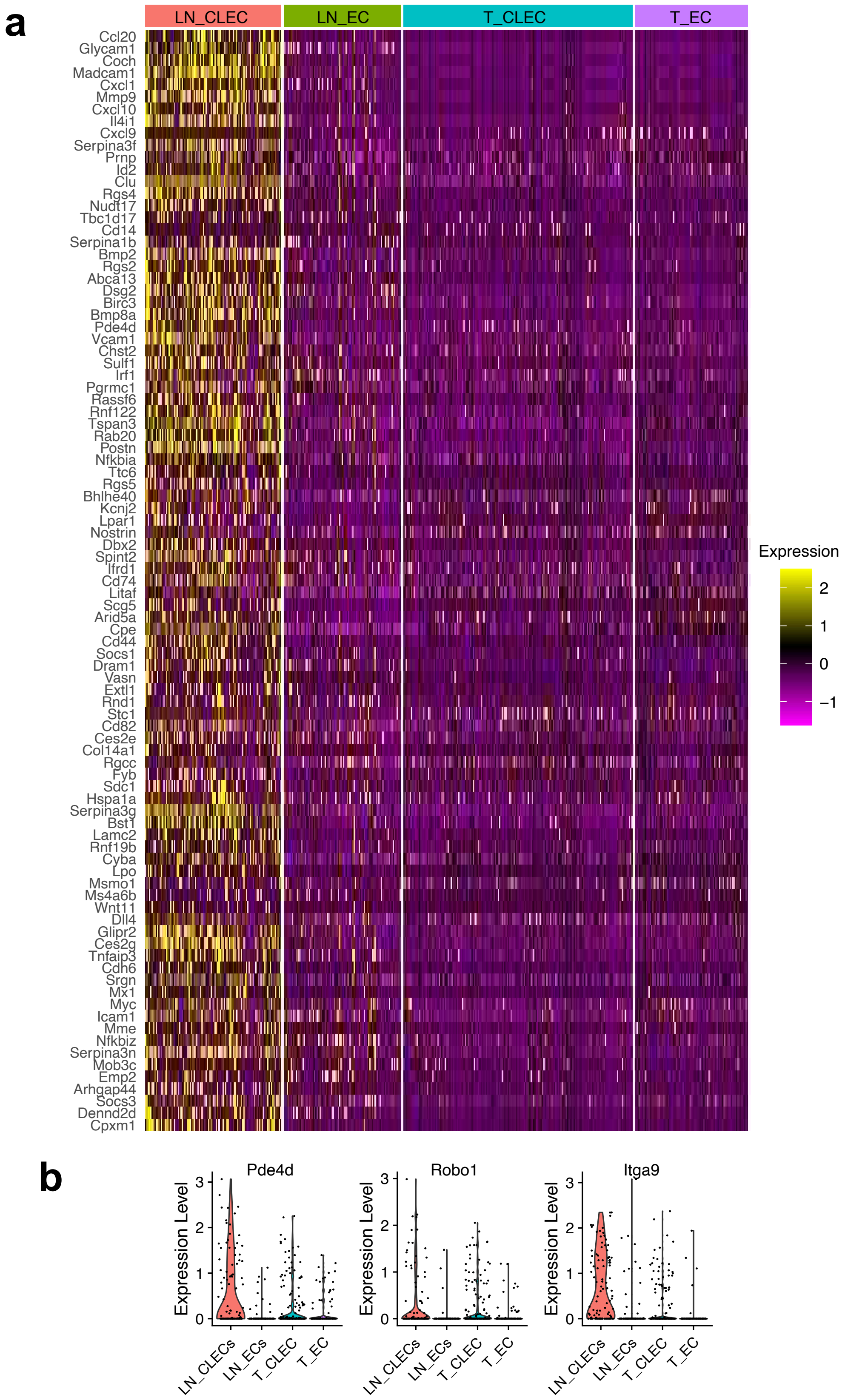


**Extended Data Fig. 9: CLEC and EC heterogeneity in tumor and lymph node** .

a, Heatmap shows the expression of n=91 fLECs markers in LN_CLEC, LN_EC, T_CLEC and T_EC single cells. b, Violin plots show the expression of 3 CLEC markers: Pde4d, Robo1 and Itga9

**Matsumoto K et al., Extended Data Figure 9**
